## Supplemental table 2 for "Digestive Dimensions of Autism: A Multiscale Exploration of Gut-Brain Interactions"

| Pathway | Total | Expected | Hits | P.Value | FDR | Genes |
|---|---|---|---|---|---|---|
| Neuron projection | 685 | 15.9 | 38 | 6.36E-07 | 0.000143 | SLC8A3,PTPRN2,SYNGAP1,SEMA[illegible] |
| Dendrite | 342 | 7.94 | 23 | 5.39E-06 | 0.000607 | SLC8A3,SYNGAP1,SEMA3A,GRIN[illegible] |
| Cell projection part | 686 | 15.9 | 32 | 0.000148 | 0.0111 | SLC8A3,PTPRN2,SYNGAP1,GRIN[illegible] |
| Synapse | 558 | 13 | 26 | 0.000626 | 0.0352 | PTPRN2,GRIN2A,DMXL2,CHRNA5,[illegible] |
| Synapse part | 396 | 9.19 | 20 | 0.001 | 0.044 | PTPRN2,GRIN2A,DMXL2,CHRNA5,[illegible] |
| Growth cone | 92 | 2.14 | 8 | 0.00134 | 0.044 | MAPT,EXOC4,NGEF,TWF2,TSHZ3 |
| Cell projection | 1410 | 32.7 | 50 | 0.00166 | 0.044 | SLC8A3,MAGI1,PTPRN2,SYNGAP1 |
| Integral to endoplasm ic reticulum membrane | 118 | 2.74 | 9 | 0.00173 | 0.044 | HLA-B,RTN1,SPPL3,EXT1,HLA-DQ[illegible] |
| Site of polarized growth | 96 | 2.23 | 8 | 0.00176 | 0.044 | MAPT,EXOC4,NGEF,TWF2,TSHZ3 |
| Intrinsic to endoplasm ic reticulum membrane | 135 | 3.13 | 9 | 0.00428 | 0.0964 | HLA-B,RTN1,SPPL3,EXT1,HLA-DQ[illegible] |
| Voltage_ga ted calcium channel complex | 33 | 0.766 | 4 | 0.00687 | 0.13 | CACNA2D2,CACNA1C,CACNB2,C[illegible] |
| Trans_Gol gi network | 120 | 2.79 | 8 | 0.00691 | 0.13 | PRKD1,ARFGEF2,HLA-DQA1,FUR[illegible] |
| Nuclear speck | 152 | 3.53 | 9 | 0.00914 | 0.154 | KMT2E,RSRC1,CTR9,ZMIZ1,ZNF6[illegible] |
| Lamellipodi um | 127 | 2.95 | 8 | 0.0096 | 0.154 | DPP4,DYSF,CARMIL1,CTNND1,CT[illegible] |
| Transport vesicle | 158 | 3.67 | 9 | 0.0116 | 0.174 | PTPRN2,HLA-B,HLA-DQA1,SYT1,F[illegible] |
| Cell junction | 847 | 19.7 | 30 | 0.0144 | 0.202 | MAGI1,GRIN2A,DMXL2,PRKD1,DP[illegible] |
| Myofibril | 197 | 4.57 | 10 | 0.017 | 0.225 | HDAC4,FXR1,CACNA1C,CALD1,C[illegible] |
| Apical junction complex | 123 | 2.86 | 7 | 0.0247 | 0.308 | MAGI1,MAGI2,CTNNB1,ANK3,PAR[illegible] |
| Contractile fiber | 214 | 4.97 | 10 | 0.0281 | 0.333 | HDAC4,FXR1,CACNA1C,CALD1,C[illegible] |
| Cell_cell junction | 346 | 8.03 | 14 | 0.0319 | 0.346 | MAGI1,PRKD1,DPP4,PKP4,LDB1,D[illegible] |
| Cell leading edge | 283 | 6.57 | 12 | 0.0332 | 0.346 | DPP4,CNTNAP2,DYSF,CARMIL1,C[illegible] |

| | | | | | | |
|---|---|---|---|---|---|---|
| Nucleoplasm part | 910 | 21.1 | 30 | 0.0339 | 0.346 | KMT2D,KMT2A,MSL2,KMT2E,RERI |
| Integral to organelle membrane | 231 | 5.36 | 10 | 0.0438 | 0.428 | HLA-B,RTN1,SPPL3,EXT1,HLA-DQ |
| Axon | 269 | 6.24 | 11 | 0.05 | 0.469 | PTPRN2,SEMA3A,FEZ1,MAPT,CNT |
| Mediator complex | 37 | 0.859 | 3 | 0.054 | 0.486 | MED27,MED8,GLI3 |
| Acetylcholine_gated channel complex | 17 | 0.395 | 2 | 0.0581 | 0.503 | CHRNA5,CHRNA3 |
| Contractile fiber part | 187 | 4.34 | 8 | 0.0706 | 0.579 | HDAC4,FXR1,CACNA1C,CTNNB1,/ |
| Receptor complex | 189 | 4.39 | 8 | 0.0741 | 0.579 | GRIN2A,CHRNA5,GABBR1,ITGA11 |
| Intrinsic to organelle membrane | 255 | 5.92 | 10 | 0.0746 | 0.579 | HLA-B,RTN1,SPPL3,EXT1,HLA-DQ |
| Clathrin_coated vesicle | 224 | 5.2 | 9 | 0.0782 | 0.586 | DMXL2,PCLO,HLA-DQA1,SYT1,FU |
| Nuclear body | 295 | 6.85 | 11 | 0.0838 | 0.59 | KMT2E,RSRC1,CTR9,ANKS1B,ZMI |
| Coated vesicle membrane | 162 | 3.76 | 7 | 0.0839 | 0.59 | HLA-B,DMXL2,HLA-DQA1,SYT1,HL |
| Perinuclear region of cytoplasm | 475 | 11 | 16 | 0.0885 | 0.603 | DGKI,PTCH1,PKP4,DYNC1I1,HFE,I |
| Kinesin complex | 24 | 0.557 | 2 | 0.106 | 0.692 | KLC1,KIF5C |
| Replication fork | 50 | 1.16 | 3 | 0.11 | 0.692 | RPA3,RFC2,ZMIZ2 |
| Microtubule associated complex | 141 | 3.27 | 6 | 0.111 | 0.692 | MAPT,DYNC1I1,KLC1,KIF5C,KIF21 |
| Coated vesicle | 283 | 6.57 | 10 | 0.124 | 0.739 | HLA-B,DMXL2,PCLO,HLA-DQA1,SY |
| Histone deacetylase complex | 53 | 1.23 | 3 | 0.125 | 0.739 | RERE,HDAC4,HDAC9 |
| Nuclear replication fork | 28 | 0.65 | 2 | 0.137 | 0.766 | RPA3,ZMIZ2 |
| Cell body | 290 | 6.73 | 10 | 0.139 | 0.766 | SLC8A3,CNTNAP2,CHRNA3,CACN |
| Endocytic vesicle | 187 | 4.34 | 7 | 0.146 | 0.766 | HLA-B,DPP4,HLA-DQA1,SYT1,HLA |
| Lysosomal membrane | 153 | 3.55 | 6 | 0.146 | 0.766 | HLA-DMA,HLA-DQA1,ABCB9,HLA-I |

| | | | | | | |
|---|---|---|---|---|---|---|
| Cytoplasmic vesicle membrane | 403 | 9.36 | 13 | 0.146 | 0.766 | PTPRN2,HLA-B,SPPL3,DMXL2,DYS |
| Membrane raft | 189 | 4.39 | 7 | 0.151 | 0.774 | PTCH1,DPP4,CACNA1C,CD55,FUF |
| Nucleoplasm | 1820 | 42.3 | 49 | 0.155 | 0.776 | KMT2D,KMT2A,MSL2,RPA3,KMT2E |
| Transcription factor complex | 303 | 7.03 | 10 | 0.169 | 0.812 | FOXP1,TCF4,LDB1,XRCC6,CTNNE |
| Golgi membrane | 605 | 14 | 18 | 0.17 | 0.812 | GALNT12,HLA-B,SPPL3,CLSTN2,M |
| Vesicle membrane | 417 | 9.68 | 13 | 0.174 | 0.816 | PTPRN2,HLA-B,SPPL3,DMXL2,DYS |
| Sarcomere | 163 | 3.78 | 6 | 0.179 | 0.822 | HDAC4,CACNA1C,CTNNB1,KCNN2 |
| Microtubule | 352 | 8.17 | 11 | 0.197 | 0.888 | SLC8A3,FEZ1,MAPT,SPAG16,DYN |
| Spindle pole | 101 | 2.34 | 4 | 0.208 | 0.917 | PKP4,DYNC1I1,CUL3,CTNNB1 |
| Microvillus | 70 | 1.63 | 3 | 0.222 | 0.932 | CTNNB1,TWF2,DOCK4 |
| Synaptic vesicle | 104 | 2.41 | 4 | 0.222 | 0.932 | DMXL2,PCLO,SYT1,CTTNBP2 |
| Tight junction | 105 | 2.44 | 4 | 0.227 | 0.932 | MAGI1,MAGI2,ANK3,PARD3B |
| Voltage_gated potassium channel complex | 71 | 1.65 | 3 | 0.228 | 0.932 | KCNG2,CNTNAP2,KCNB1 |
| Golgi apparatus part | 761 | 17.7 | 21 | 0.237 | 0.951 | GALNT12,HLA-B,SPPL3,F2,CLSTN |
| Cytoplasmic vesicle part | 462 | 10.7 | 13 | 0.278 | 1 | PTPRN2,HLA-B,SPPL3,DMXL2,DYS |
| DNA_directed RNA polymerase II, core complex | 14 | 0.325 | 1 | 0.28 | 1 | POLR2E |
| Intercalated disc | 45 | 1.04 | 2 | 0.281 | 1 | CTNNB1,ANK3 |
| Golgi stack | 117 | 2.72 | 4 | 0.288 | 1 | ST3GAL3,GBF1,ZFYVE1,GALNT1 |
| Cell cortex | 195 | 4.53 | 6 | 0.3 | 1 | PRKD1,EXOC4,MYO9B,CALD1,CTI |
| Protein serine/threonine phosphatase complex | 48 | 1.11 | 2 | 0.307 | 1 | PPP2R2B,PPP2R5C |
| Cell surface | 518 | 12 | 14 | 0.318 | 1 | ANXA9,GRIN2A,DPP4,CNTNAP2,P |

| | | | | | | |
|---|---|---|---|---|---|---|
| Extrinsic to plasma membrane | 87 | 2.02 | 3 | 0.329 | 1 | GNAI1,CTNNB1,RGS6 |
| Vacuolar membrane | 206 | 4.78 | 6 | 0.345 | 1 | HLA-DMA,HLA-DQA1,ABCB9,HLA-I |
| Apical part of cell | 289 | 6.71 | 8 | 0.357 | 1 | DPP4,HFE,CD55,CTNNB1,SRR,PTI |
| Late endosome | 175 | 4.06 | 5 | 0.384 | 1 | HLA-DMA,HLA-DRB5,HLA-DRB1,W |
| Endoplasmic reticulum lumen | 175 | 4.06 | 5 | 0.384 | 1 | PTPRN2,F2,COL11A1,CALU,GBF1 |
| Anchored to membrane | 147 | 3.41 | 4 | 0.446 | 1 | CNTN4,CD55,NTM,NEGR1 |
| Lytic vacuole | 401 | 9.31 | 10 | 0.454 | 1 | HLA-DMA,USP4,RPTOR,HLA-DQA1 |
| Lysosome | 401 | 9.31 | 10 | 0.454 | 1 | HLA-DMA,USP4,RPTOR,HLA-DQA1 |
| Nuclear ubiquitin ligase complex | 26 | 0.604 | 1 | 0.457 | 1 | ANAPC4 |
| Trans_Golgi network transport vesicle | 26 | 0.604 | 1 | 0.457 | 1 | FURIN |
| Golgi_associated vesicle | 70 | 1.63 | 2 | 0.486 | 1 | SPPL3,FURIN |
| Plasma membrane part | 2320 | 53.7 | 54 | 0.508 | 1 | PTPRN2,HLA-B,GNAI1,SYNGAP1,H |
| Myosin complex | 75 | 1.74 | 2 | 0.523 | 1 | MYO18A,MYO9B |
| Protein complex | 4050 | 94.1 | 94 | 0.523 | 1 | SLC8A3,KMT2D,MYO18A,KMT2A,E |
| Integrin complex | 32 | 0.743 | 1 | 0.529 | 1 | ITGA11 |
| Actin cytoskeleton | 430 | 9.98 | 10 | 0.543 | 1 | MYO18A,MYO9B,HDAC4,MAD1L1,I |
| Nuclear lumen | 2690 | 62.5 | 62 | 0.548 | 1 | KMT2D,KMT2A,MSL2,RPA3,KMT2E |
| Apical plasma membrane | 216 | 5.01 | 5 | 0.565 | 1 | DPP4,CD55,PTPRO,CLCN3,ATP2B |
| Integral to plasma membrane | 1270 | 29.5 | 29 | 0.566 | 1 | PTPRN2,HLA-B,GRIN2A,FADS2,PF |
| Early endosome | 217 | 5.04 | 5 | 0.569 | 1 | HLA-B,HFE,CNTNAP2,WNT3,CLCN |

| | | | | | | |
|---|---|---|---|---|---|---|
| Endoplasmic reticulum part | 1060 | 24.5 | 24 | 0.572 | 1 | PTPRN2,ERLIN1,HLA-B,CISD2,RTN |
| Vacuole | 486 | 11.3 | 11 | 0.577 | 1 | HLA-DMA,USP4,RPTOR,HLA-DQA1 |
| Cortical actin cytoskeleton | 37 | 0.859 | 1 | 0.581 | 1 | CALD1 |
| Intrinsic to plasma membrane | 1320 | 30.7 | 30 | 0.582 | 1 | PTPRN2,HLA-B,SYNGAP1,GRIN2A |
| Nuclear pore | 86 | 2 | 2 | 0.597 | 1 | MAD1L1,POM121C |
| Nuclear outer membrane_endoplasmic reticulum membrane network | 894 | 20.8 | 20 | 0.601 | 1 | ERLIN1,HLA-B,CISD2,RTN1,VRK2, |
| Extrinsic to membrane | 135 | 3.13 | 3 | 0.61 | 1 | GNAI1,CTNNB1,RGS6 |
| Microtubule organizing center | 543 | 12.6 | 12 | 0.611 | 1 | GNAI1,FEZ1,WRN,KIZ,BBS9,MPHO |
| Centrosome | 412 | 9.56 | 9 | 0.621 | 1 | GNAI1,FEZ1,WRN,KIZ,MPHOSPH9 |
| Vacuolar part | 279 | 6.48 | 6 | 0.632 | 1 | HLA-DMA,HLA-DQA1,ABCB9,HLA-[illegible] |
| Spindle microtubule | 43 | 0.998 | 1 | 0.636 | 1 | CUL3 |
| Golgi apparatus | 1400 | 32.5 | 31 | 0.638 | 1 | GALNT12,HLA-B,SPPL3,PTCH1,PR |
| Collagen | 93 | 2.16 | 2 | 0.64 | 1 | COL11A1, TNXB |
| DNA_directed RNA polymerase II, holoenzyme | 93 | 2.16 | 2 | 0.64 | 1 | CTR9,POLR2E |
| Endoplasmic reticulum membrane | 872 | 20.2 | 19 | 0.646 | 1 | ERLIN1,HLA-B,CISD2,RTN1,VRK2, |
| Nucleolus | 652 | 15.1 | 14 | 0.656 | 1 | WRN,MRM2,PPM1E,FXR1,MED27, |
| Cell cortex part | 97 | 2.25 | 2 | 0.662 | 1 | EXOC4,CALD1 |

| | | | | | | |
|---|---|---|---|---|---|---|
| Integral to Golgi membrane | 47 | 1.09 | 1 | 0.669 | 1 | ST3GAL3 |
| Chromosome, centromeric region | 198 | 4.6 | 4 | 0.678 | 1 | DYNC1I1,MAD1L1,STAG1,PPP2R5 |
| Intrinsic to Golgi membrane | 50 | 1.16 | 1 | 0.692 | 1 | ST3GAL3 |
| Pore complex | 103 | 2.39 | 2 | 0.694 | 1 | MAD1L1,POM121C |
| Cytoskeletal part | 1570 | 36.5 | 34 | 0.698 | 1 | SLC8A3,MYO18A,GNAI1,GRIN2A,F |
| Actin filament | 51 | 1.18 | 1 | 0.699 | 1 | MYO9B |
| DNA_directed RNA polymerase complex | 105 | 2.44 | 2 | 0.704 | 1 | CTR9,POLR2E |
| Nuclear DNA_directed RNA polymerase complex | 105 | 2.44 | 2 | 0.704 | 1 | CTR9,POLR2E |
| Organelle outer membrane | 156 | 3.62 | 3 | 0.706 | 1 | CISD2,PPP2R2B,SYNE1 |
| RNA polymerase complex | 106 | 2.46 | 2 | 0.709 | 1 | CTR9,POLR2E |
| Spindle | 261 | 6.06 | 5 | 0.728 | 1 | PKP4,DYNC1I1,CUL3,MAD1L1,CTN |
| Basolateral plasma membrane | 162 | 3.76 | 3 | 0.73 | 1 | CTNNB1,ANK3,CNNM2 |
| Outer membrane | 163 | 3.78 | 3 | 0.733 | 1 | CISD2,PPP2R2B,SYNE1 |
| Cytoplasmic vesicle | 1110 | 25.7 | 23 | 0.743 | 1 | PTPRN2,HLA-B,SPPL3,DMXL2,DP |
| Cytoplasmic membrane_bounded vesicle | 1020 | 23.7 | 21 | 0.752 | 1 | PTPRN2,HLA-B,SPPL3,DMXL2,DP |
| Cortical cytoskeleton | 60 | 1.39 | 1 | 0.756 | 1 | CALD1 |
| Microtubule cytoskeleton | 1120 | 26 | 23 | 0.757 | 1 | SLC8A3,GNAI1,FEZ1,MAPT,WRN, |
| Nuclear chromosome part | 273 | 6.34 | 5 | 0.763 | 1 | RPA3,LDB1,MSH5,XRCC6,ZMIZ2 |

| | | | | | | |
|---|---|---|---|---|---|---|
| Peroxisomal membrane | 71 | 1.65 | 1 | 0.812 | 1 | CNOT1 |
| Microbody membrane | 71 | 1.65 | 1 | 0.812 | 1 | CNOT1 |
| Vesicle | 1210 | 28.1 | 24 | 0.816 | 1 | PTPRN2,HLA-B,SPPL3,DMXL2,DPI |
| Organelle lumen | 3380 | 78.6 | 72 | 0.817 | 1 | PTPRN2,KMT2D,KMT2A,MSL2,RPA |
| Mitochondrial outer membrane | 133 | 3.09 | 2 | 0.818 | 1 | CISD2,PPP2R2B |
| Condensed nuclear chromosome | 73 | 1.69 | 1 | 0.821 | 1 | MSH5 |
| Condensed chromosome | 193 | 4.48 | 3 | 0.829 | 1 | DYNC1I1,MSH5,MAD1L1 |
| Cell_substrate adherens junction | 138 | 3.2 | 2 | 0.834 | 1 | CTNNB1,SORBS1 |
| Endoplasmic reticulum_Golgi intermediate compartment | 77 | 1.79 | 1 | 0.837 | 1 | MYO18A |
| PML body | 77 | 1.79 | 1 | 0.837 | 1 | TENM2 |
| Cytoskeleton | 2200 | 51 | 45 | 0.84 | 1 | SLC8A3,MYO18A,GNAI1,GRIN2A,F |
| Nuclear part | 3330 | 77.4 | 70 | 0.844 | 1 | KMT2D,KMT2A,MSL2,RPA3,AKAP6 |
| Membrane_bounded vesicle | 1100 | 25.5 | 21 | 0.852 | 1 | PTPRN2,HLA-B,SPPL3,DMXL2,DPI |
| Extracellular matrix part | 204 | 4.74 | 3 | 0.856 | 1 | THSD4,COL11A1,TNXB |
| Cell_substrate junction | 147 | 3.41 | 2 | 0.859 | 1 | CTNNB1,SORBS1 |
| Membrane_enclosed lumen | 3440 | 80 | 72 | 0.859 | 1 | PTPRN2,KMT2D,KMT2A,MSL2,RPA |
| Nuclear membrane | 207 | 4.81 | 3 | 0.863 | 1 | AKAP6,POM121C,SYNE1 |
| Kinetochore | 149 | 3.46 | 2 | 0.864 | 1 | DYNC1I1,MAD1L1 |

| | | | | | | |
|---|---|---|---|---|---|---|
| Nuclear chromosome | 320 | 7.43 | 5 | 0.868 | 1 | RPA3,LDB1,MSH5,XRCC6,ZMIZ2 |
| Macromolecular complex | 4800 | 111 | 102 | 0.869 | 1 | SLC8A3,KMT2D,MYO18A,KMT2A,E |
| Nuclear matrix | 87 | 2.02 | 1 | 0.871 | 1 | SORBS1 |
| Adherens junction | 212 | 4.92 | 3 | 0.873 | 1 | CTNND1,CTNNB1,SORBS1 |
| Mitochondrial respiratory chain complex I | 89 | 2.07 | 1 | 0.877 | 1 | NDUFA2 |
| Membrane coat | 89 | 2.07 | 1 | 0.877 | 1 | SNAP91 |
| NADH dehydrogenase complex | 89 | 2.07 | 1 | 0.877 | 1 | NDUFA2 |
| Respiratory chain complex I | 89 | 2.07 | 1 | 0.877 | 1 | NDUFA2 |
| Coated membrane | 89 | 2.07 | 1 | 0.877 | 1 | SNAP91 |
| Ubiquitin ligase complex | 165 | 3.83 | 2 | 0.899 | 1 | ANAPC4,CUL3 |
| Microtubule organizing center part | 100 | 2.32 | 1 | 0.905 | 1 | MPHOSPH9 |
| Endomembrane system | 2160 | 50.1 | 42 | 0.909 | 1 | PTPRN2,GALNT12,ERLIN1,HLA-B,/ |
| Chromosomal part | 670 | 15.6 | 11 | 0.913 | 1 | RPA3,DYNC1I1,CTR9,LDB1,MSH5, |
| Microbody part | 106 | 2.46 | 1 | 0.918 | 1 | CNOT1 |
| Peroxisomal part | 106 | 2.46 | 1 | 0.918 | 1 | CNOT1 |
| Endosome | 683 | 15.9 | 11 | 0.924 | 1 | HLA-B,HLA-DMA,HFE,CNTNAP2,AI |
| Peroxisome | 181 | 4.2 | 2 | 0.926 | 1 | CNOT1,GBF1 |
| Microbody | 181 | 4.2 | 2 | 0.926 | 1 | CNOT1,GBF1 |
| Cytosol | 2660 | 61.8 | 52 | 0.929 | 1 | PDE1C,ANXA9,ARHGAP15,PSMA5 |
| Mitochondrial respiratory chain | 116 | 2.69 | 1 | 0.935 | 1 | NDUFA2 |

| | | | | | | |
|---|---|---|---|---|---|---|
| Proteasome complex | 117 | 2.72 | 1 | 0.937 | 1 | PSMA5 |
| External side of plasma membrane | 204 | 4.74 | 2 | 0.952 | 1 | HLA-DRB5,HLA-DRB1 |
| Focal adhesion | 132 | 3.06 | 1 | 0.956 | 1 | SORBS1 |
| Proteinaceous extracellular matrix | 398 | 9.24 | 5 | 0.956 | 1 | THSD4,COL11A1,MMP16,TNXB,WI |
| Ruffle | 135 | 3.13 | 1 | 0.959 | 1 | RHOA |
| Endoplasmic reticulum | 1660 | 38.5 | 29 | 0.962 | 1 | FKBP9,PTPRN2,ERLIN1,HLA-B,AK/ |
| Mitochondrial membrane part | 217 | 5.04 | 2 | 0.963 | 1 | IMMP2L,NDUFA2 |
| Chromosome | 784 | 18.2 | 11 | 0.976 | 1 | RPA3,DYNC1I1,CTR9,LDB1,MSH5, |
| Nuclear chromatin | 159 | 3.69 | 1 | 0.977 | 1 | LDB1 |
| Spliceosomal complex | 161 | 3.74 | 1 | 0.978 | 1 | PRPF3 |
| Nuclear envelope | 387 | 8.98 | 4 | 0.981 | 1 | AKAP6,MAD1L1,POM121C,SYNE1 |
| Ribosome | 249 | 5.78 | 2 | 0.981 | 1 | MRPL33,RPS6KL1 |
| Chromatin | 326 | 7.57 | 3 | 0.983 | 1 | CTR9,LDB1,STAG1 |
| Mitochondrial matrix | 329 | 7.64 | 3 | 0.983 | 1 | TFB1M,IVD,PPA2 |
| Mitochondrial matrix | 329 | 7.64 | 3 | 0.983 | 1 | TFB1M,IVD,PPA2 |
| Secretory granule | 276 | 6.41 | 2 | 0.989 | 1 | PTPRN2,SYT1 |
| Plasma membrane | 5500 | 128 | 107 | 0.99 | 1 | SLC8A3,MAGI1,GPR26,PTPRN2,HI |
| Ribonucleoprotein complex | 681 | 15.8 | 8 | 0.991 | 1 | MRPL33,TNRC6A,SND1,FXR1,PRF |
| Non_membrane_bounded organelle | 3940 | 91.5 | 73 | 0.991 | 1 | SLC8A3,MYO18A,RPA3,GNAI1,MR |
| Intracellular non_membrane_bounded organelle | 3940 | 91.5 | 73 | 0.991 | 1 | SLC8A3,MYO18A,RPA3,GNAI1,MR |
| Cytosolic part | 204 | 4.74 | 1 | 0.992 | 1 | PIK3R2 |

| | | | | | | |
|---|---|---|---|---|---|---|
| Mitochondrial membrane | 684 | 15.9 | 7 | 0.997 | 1 | CISD2,VRK2,SPNS1,GPD2,PPP2R2 |
| Extracellular matrix | 570 | 13.2 | 5 | 0.997 | 1 | THSD4,COL11A1,MMP16,TNXB,WI |
| Mitochondrial inner membrane | 504 | 11.7 | 4 | 0.998 | 1 | SPNS1,GPD2,IMMP2L,NDUFA2 |
| Intrinsic to membrane | 5760 | 134 | 108 | 0.998 | 1 | GRAMD1B,SLC8A3,GPR26,PTPRN |
| Integral to membrane | 5590 | 130 | 104 | 0.998 | 1 | GRAMD1B,SLC8A3,GPR26,PTPRN |
| Mitochondrial envelope | 729 | 16.9 | 7 | 0.998 | 1 | CISD2,VRK2,SPNS1,GPD2,PPP2R2 |
| Organelle inner membrane | 537 | 12.5 | 4 | 0.999 | 1 | SPNS1,GPD2,IMMP2L,NDUFA2 |
| Extracellular space | 901 | 20.9 | 9 | 0.999 | 1 | SEMA3E,ENOX1,PXDNL,F2,OLFM4 |
| Mitochondrion | 2060 | 47.7 | 29 | 0.999 | 1 | MTHFD1L,SLC8A3,MSRA,MRPL33, |
| Envelope | 1130 | 26.2 | 12 | 1 | 1 | AKAP6,CISD2,GRIN2A,VRK2,SPNS |
| Organelle membrane | 3020 | 70.1 | 46 | 1 | 1 | PTPRN2,GALNT12,ERLIN1,HLA-B,/ |
| Mitochondrial part | 1040 | 24.1 | 10 | 1 | 1 | CISD2,VRK2,SPNS1,GPD2,TFB1M, |
| Organelle envelope | 1110 | 25.9 | 11 | 1 | 1 | AKAP6,CISD2,VRK2,SPNS1,GPD2, |
| Extracellular region part | 1320 | 30.6 | 13 | 1 | 1 | SEMA3E,THSD4,ENOX1,PXDNL,DI |
| Nucleus | 7600 | 177 | 137 | 1 | 1 | KDM5D,NREP,NPAS3,KMT2D,KMT |
| Organelle part | 8790 | 204 | 149 | 1 | 1 | SLC8A3,PTPRN2,GALNT12,KMT2D |
| Membrane part | 7520 | 175 | 120 | 1 | 1 | GRAMD1B,SLC8A3,GPR26,PTPRN |
| Intracellular organelle part | 8620 | 200 | 144 | 1 | 1 | SLC8A3,PTPRN2,GALNT12,KMT2D |
| Extracellular region | 2860 | 66.3 | 28 | 1 | 1 | SEMA3E,SEMA3A,THSD4,ENOX1,I |
| Cytoplasmic part | 9740 | 226 | 150 | 1 | 1 | FKBP9,MTHFD1L,SLC8A3,PDE1C,I |
| Cytoplasm | 13100 | 305 | 215 | 1 | 1 | FKBP9,MTHFD1L,NREP,SLC8A3,M |
| Membrane | 11700 | 272 | 170 | 1 | 1 | GRAMD1B,FKBP9,SLC8A3,MAGI1, |

3A,GRIN2A,FEZ1,MAPT,EXOC4,CNTNAP2,BSN,CNTN4,ANKS1B,KIF5C,ARFGEF2,CHRNA3,CACNA1C,TT

2A,FEZ1,CNTNAP2,BSN,ANKS1B,ARFGEF2,CHRNA3,CACNA1C,TTLL7,GRIK1,DBN1,MAGI2,CTNNB1,KCN

2A,MAPT,EXOC4,DPP4,BBS9,SPAG16,CNTNAP2,BSN,ANKS1B,KIF5C,ARFGEF2,CHRNA3,CACNA1C,TTLL

,CLSTN2,PCLO,HDAC4,GABBR1,BSN,ANKS1B,ARFGEF2,CHRNA3,CACNA1C,GRIK1,MAGI2,CADM2,CTN

,CLSTN2,PCLO,GABBR1,BSN,ANKS1B,CHRNA3,CACNA1C,GRIK1,MAGI2,SYT1,KCNB1,ANK3,CTTNBP2,

,SEMA3A,GRIN2A,FEZ1,MAPT,EXOC4,DPP4,BBS9,SPAG16,CNTNAP2,BSN,DYSF,CNTN4,ANKS1B,KIF5C

P4,PKP4,CHRNA5,PCLO,GABBR1,BSN,ANKS1B,ARFGEF2,CHRNA3,LDB1,GRIK1,DBN1,MAGI2,CADM2,C [illegible],FOXP1,RSRC1,TCF4,CTR9,SGF29,HDAC4,ANKS1B,ZMIZ1,LDB1,ZNF638,PRPF3,XRCC6,MEF2C,CTNN[illegible]

E,RERE,FOXP1,PSMA5,RSRC1,ESR2,ESRRG,WRN,TAF1C,ANAPC4,TCF4,CTR9,SGF29,HDAC4,ZFPM2,N

IPHOSPH9,EXT1,ARFGEF2,HLA-DQA1,MGAT5B,ST3GAL3,FURIN,GBF1,HLA-DRB5,HLA-DRB1,GALNT1,G

2,MPHOSPH9,MMP16,EXT1,ARFGEF2,HLA-DQA1,MGAT5B,ST3GAL3,FURIN,GBF1,HLA-DRB5,ZFYVE1,H ıLA-DMA,GRIN2A,FADS2,PTCH1,PRKD1,EXOC4,DPP4,PKP4,BBS9,CHRNA5,CACNA2D2,HFE,KCNG2,SL(

ERLIN1,MSL2,HLA-B,DGKI,RPA3,AKAP6,KMT2E,GNAI1,RERE,CISD2,FOXP1,HLA-DMA,GRIN2A,FEZ1,PSM

E,RERE,FOXP1,PSMA5,RSRC1,ESR2,MAPT,ESRRG,WRN,MRM2,TAF1C,ANAPC4,TCF4,CTR9,SGF29,HD/

RKD1,CHRNA5,HFE,KCNG2,SLC6A9,CNTNAP2,GABBR1,MMP16,ITGA11,ATP2A2,CHRNA3,FGFR1,HLA-D(

N1,VRK2,AGMO,SPPL3,FADS2,F2,CLSTN2,COL11A1,EXT1,CALU,ATP2A2,HLA-DQA1,ABCB9,DDN,POM12

,FADS2,PRKD1,CHRNA5,HFE,KCNG2,SLC6A9,CNTNAP2,GABBR1,MMP16,ITGA11,ATP2A2,CHRNA3,FGF

AGMO,SPPL3,FADS2,CLSTN2,EXT1,ATP2A2,HLA-DQA1,ABCB9,DDN,POM121C,HLA-DRB5,HLA-DRB1,SY

KD1,F2,CLSTN2,CUL3,CNTNAP2,MPHOSPH9,SEMA6D,MMP16,EXT1,CALU,ARFGEF2,HLA-DQA1,MGAT5

AGMO,SPPL3,FADS2,CLSTN2,EXT1,ATP2A2,HLA-DQA1,ABCB9,DDN,POM121C,HLA-DRB5,HLA-DRB1,HL [illegible]EZ1,MAPT,WRN,KIZ,PKP4,BBS9,SPAG16,DYNC1I1,KLC1,MYO9B,CUL3,HDAC4,MPHOSPH9,ANKS1B,KIF[illegible]

[illegible]P4,HFE,PCLO,DYSF,SND1,CALU,ARFGEF2,FGFR1,HLA-DQA1,SYT1,FURIN,CTTNBP2,CADPS,HLA-DRB5

[illegible]P4,PCLO,DYSF,SND1,CALU,ARFGEF2,FGFR1,HLA-DQA1,SYT1,FURIN,CTTNBP2,CADPS,HLA-DRB5,HLA-[illegible]

[illegible]KIZ,PKP4,BBS9,SPAG16,DYNC1I1,KLC1,CUL3,BSN,MPHOSPH9,KIF5C,ARFGEF2,TTLL7,MAD1L1,CTNNB[illegible]

P4,DYNC1I1,HFE,PCLO,DYSF,SND1,CALU,ARFGEF2,FGFR1,HLA-DQA1,SYT1,FURIN,CTTNBP2,CADPS,H

A3,KMT2E,RERE,FOXP1,PSMA5,RSRC1,ESR2,MAPT,ESRRG,WRN,MRM2,F2,TAF1C,ANAPC4,TCF4,CTR9

EZ1,MAPT,WRN,KIZ,PKP4,BBS9,SPAG16,DYNC1I1,KLC1,MYO9B,CUL3,PCLO,HDAC4,BSN,MPHOSPH9,A

5,KMT2E,RERE,FOXP1,PSMA5,RSRC1,ESR2,MAPT,ESRRG,WRN,MRM2,TAF1C,ANAPC4,TCF4,CTR9,SGI

P4,PCLO,DYSF,SND1,CALU,ARFGEF2,FGFR1,HLA-DQA1,SYT1,FURIN,CTTNBP2,CADPS,HLA-DRB5,HLA-

A3,KMT2E,RERE,FOXP1,PSMA5,RSRC1,ESR2,MAPT,ESRRG,WRN,MRM2,F2,TAF1C,ANAPC4,TCF4,CTR9

ERLIN1,MSL2,HLA-B,DGKI,RPA3,AKAP6,KMT2E,GNAI1,MRPL33,RERE,CISD2,FOXP1,HLA-DMA,GRIN2A,F

AKAP6,CISD2,RTN1,VRK2,AGMO,SPPL3,FADS2,DMXL2,CLSTN2,MPHOSPH9,DYSF,EXT1,ATP2A2,ARFG

,RSU1,MAPT,PRKD1,PDE8B,ANAPC4,DYNC1I1,KLC1,TNRC6A,MYO9B,DOCK8,RPTOR,HDAC4,PFAS,NE

AP6,CISD2,RTN1,VRK2,AGMO,SPPL3,FADS2,F2,CLSTN2,COL11A1,EXT1,CALU,ATP2A2,HLA-DQA1,ABCI

LA-B,DGKI,GNAI1,SYNGAP1,HLA-DMA,ENOX1,GRIN2A,FEZ1,CDHR4,MAPT,GPR141,FADS2,PTCH1,PRKI

PL33,GRIN2A,FEZ1,MAPT,WRN,KIZ,PKP4,BBS9,MRM2,SPAG16,DYNC1I1,CTR9,KLC1,TNRC6A,MYO9B,C

PL33,GRIN2A,FEZ1,MAPT,WRN,KIZ,PKP4,BBS9,MRM2,SPAG16,DYNC1I1,CTR9,KLC1,TNRC6A,MYO9B,C

2,GALNT12,ERLIN1,HLA-B,AKAP6,DPP6,ADAM22,CISD2,SYNGAP1,RTN1,HLA-DMA,GRIN2A,CDHR4,VRK

2,GALNT12,ERLIN1,HLA-B,AKAP6,DPP6,ADAM22,CISD2,RTN1,HLA-DMA,GRIN2A,CDHR4,VRK2,AGMO,S

,RERE,CISD2,VRK2,ESR2,TCAIM,MRM2,SPNS1,OLFM4,GPD2,SND1,CKB,GATB,GLT8D1,MAD1L1,TFB1M

AKAP6,CISD2,RTN1,HLA-DMA,VRK2,AGMO,SPPL3,FADS2,DMXL2,BBS9,CLSTN2,SPNS1,MPHOSPH9,DY

2A,MSL2,MSRA,DGKI,RPA3,AKAP6,KMT2E,GNAI1,RERE,FOXP1,PSMA5,VRK2,RSRC1,NCOA5,ESR2,MA

),MYO18A,KMT2A,ERLIN1,MSL2,HLA-B,RPA3,AKAP6,KMT2E,GNAI1,RERE,CISD2,FOXP1,RTN1,HLA-DMA

2,GALNT12,ERLIN1,HLA-B,AKAP6,DPP6,GNAI1,ADAM22,CISD2,SYNGAP1,RTN1,HLA-DMA,GRIN2A,CDH

),MYO18A,KMT2A,ERLIN1,MSL2,HLA-B,RPA3,AKAP6,KMT2E,GNAI1,RERE,CISD2,FOXP1,RTN1,HLA-DMA

PXDNL,ESR2,MDK,DPP4,F2,ITIH3,GABBR1,CNTN4,COL11A1,OLFM4,MMP16,CALU,SFTA2,RAET1E,FGF

PTPRN2,ANXA9,GALNT12,MYO18A,ERLIN1,HLA-B,MSRA,DGKI,AKAP6,ARHGAP15,MRPL33,RERE,CISD2

AGI1,PDE1C,PTPRN2,NPAS3,ANXA9,GALNT12,MYO18A,ERLIN1,HLA-B,MSRA,DGKI,RPA3,AKAP6,ARHG
GPR26,PTPRN2,GALNT12,SEMA3E,ERLIN1,HLA-B,MSRA,DGKI,AKAP6,DPP6,ARHGAP15,GNAI1,ADAM22

LL7,AMIGO1,GRIK1,DBN1,MAGI2,CADM2,CTNNB1,NGEF,SYT1,KCNB1,ANK3,DDN,TWF2,CPEB1,TSHZ3,[illegible]

L7,MAGI2,CTNNB1,NGEF,KCNB1,ANK3,DDN,TWF2,CPEB1,TSHZ3,PTPRO,SHISA9,TENM2,PREX1,RHOA,

C,ARFGEF2,CHRNA3,CACNA1C,TTLL7,AMIGO1,CARMIL1,GRIK1,DBN1,MAGI2,CADM2,CTNND1,CTNNB1,

TNND1,CTNNB1,STAG1,SYT1,ANK3,CPEB1,CADPS,SHISA9,PARD3B,TENM2,SORBS1,RHOA

EDD4L,NR1D2,ANKS1B,ZMIZ1,LDB1,ZNF638,PRPF3,PRKAG2,XRCC6,MEF2C,PALB2,CTNNB1,STAG1,ME

C6A9,CNTNAP2,GABBR1,DYSF,MMP16,ITGA11,ATP2A2,CHRNA3,CACNA1C,RAET1E,FGFR1,HLA-DQA1,

IA5,MAPT,WRN,TSNARE1,EXOC4,PKP4,BBS9,SPAG16,CHRNA5,ANAPC4,TCF4,CACNA2D2,DYNC1I1,HF

AC4,PPM1E,ZFPM2,NEDD4L,NR1D2,ANKS1B,FXR1,ZMIZ1,LDB1,ZNF638,PRPF3,MSH5,PRKAG2,XRCC6,

QA1,GRIK1,BTN2A1,CD55,CACNB2,KCNB1,PTPRF,MICA,PTPRO,SHISA9,HLA-DRB1,NTRK3,SORBS1 [illegible]R1,HLA-DQA1,GRIK1,BTN2A1,CD55,CACNB2,KCNB1,PTPRF,MICA,PTPRO,SHISA9,HLA-DRB1,NTRK3,S[illegible]

[illegible]5B,ST3GAL3,FURIN,PRKG1,GBF1,HLA-DRB5,ZFYVE1,HLA-DRB1,TENM2,GALNT1,SYNE1,GALNT10,HLA-

[illegible]5C,ARFGEF2,CHRNA3,CACNA1C,TTLL7,MAD1L1,DBN1,MAGI2,CALD1,CTNNB1,CPEB1,KDM4A,KIF21B,[illegible]

),SGF29,HDAC4,PPM1E,ZFPM2,COL11A1,NEDD4L,NR1D2,ANKS1B,FXR1,MMP16,CALU,ZMIZ1,LDB1,ZNI

\NKS1B,KIF5C,ARFGEF2,CHRNA3,CACNA1C,TTLL7,SLC30A9,MAD1L1,DBN1,MAGI2,CALD1,FSCN3,CTNI

F29,HDAC4,PPM1E,ZFPM2,THOC7,NEDD4L,NR1D2,ANKS1B,FXR1,ZMIZ1,LDB1,ARID1B,ZNF638,PRPF3,I

),SGF29,HDAC4,PPM1E,ZFPM2,COL11A1,NEDD4L,NR1D2,ANKS1B,FXR1,MMP16,CALU,ZMIZ1,LDB1,ZNI

EZ1,PSMA5,MAPT,WRN,TSNARE1,EXOC4,PKP4,BBS9,SPAG16,CHRNA5,ANAPC4,TCF4,CACNA2D2,DYN

EF2,HLA-DQA1,MGAT5B,MAD1L1,SNAP91,ST3GAL3,SYT1,ABCB9,FURIN,DDN,CADPS,POM121C,GBF1,F

DD4L,CBLB,ARFGEF2,CITED1,CKB,NT5C2,FGFR1,CNOT1,PRKAG2,ETF1,CARMIL1,GMIP,RHOJ,MAD1L1

D1,EXOC4,DPP4,PKP4,BBS9,F2,CHRNA5,CACNA2D2,CLSTN2,HFE,KCNG2,SLC6A9,CNTNAP2,GABBR1,M

CUL3,PCLO,HDAC4,PPM1E,BSN,MPHOSPH9,ANKS1B,FXR1,KIF5C,ARFGEF2,CHRNA3,CACNA1C,TTLL7,L

CUL3,PCLO,HDAC4,PPM1E,BSN,MPHOSPH9,ANKS1B,FXR1,KIF5C,ARFGEF2,CHRNA3,CACNA1C,TTLL7,L

(2,AGMO,SPPL3,GPR141,FADS2,TSNARE1,TMX2,PTCH1,PRKD1,TMEM170B,DPP4,SLC22A23,CHRNA5,C

PPL3,GPR141,FADS2,TSNARE1,TMX2,PTCH1,PRKD1,TMEM170B,DPP4,SLC22A23,CHRNA5,CACNA2D2

SF,GPD2,EXT1,ATP2A2,ARFGEF2,HLA-DQA1,CNOT1,MGAT5B,ST3GAL3,SYT1,ABCB9,FURIN,DDN,CADI

PT,ESRRG,WRN,UTY,ZSCAN9,PRKD1,BBS9,MRM2,SPAG16,ZNF536,TAF1C,NPAS1,ANAPC4,TCF4,CTR9

,GRIN2A,FEZ1,PSMA5,VRK2,AGMO,RSRC1,SPPL3,ESR2,MAPT,ESRRG,WRN,FADS2,KIZ,DMXL2,PKP4,E

R4,VRK2,AGMO,SPPL3,GPR141,FADS2,TSNARE1,TMX2,PTCH1,PRKD1,EXOC4,TMEM170B,DPP4,PKP4,I

,GRIN2A,FEZ1,PSMA5,VRK2,AGMO,RSRC1,SPPL3,ESR2,MAPT,ESRRG,WRN,FADS2,KIZ,DMXL2,PKP4,E

,RTN1,HLA-DMA,PSMA5,VRK2,AGMO,RSU1,SPPL3,ESR2,MAPT,TCAIM,FADS2,TSNARE1,DMXL2,PTCH1

AP15,GNAI1,MRPL33,RERE,CISD2,SYNGAP1,FOXP1,RTN1,HLA-DMA,FEZ1,PSMA5,VRK2,AGMO,PXDNL

2,CISD2,SYNGAP1,SEMA3A,RTN1,HLA-DMA,ENOX1,GRIN2A,FEZ1,CDHR4,VRK2,AGMO,SPPL3,MAPT,GP

,NGEF,SYT1,KCNB1,ANK3,DDN,TWF2,CPEB1,TSHZ3,PSPH,PTPRO,PIP5K1B,SHISA9,TENM2,PREX1,ATF

ED27,ESRRB,MAML3,FOXO3,JMJD1C,RFC2,RNF111,RBL2,POLR2E,TENM2,GATAD2B,THRB,TRIM33,FO

GRIK1,BTN2A1,CADM2,CD55,CTNNB1,CACNB2,KCNB1,ANK3,PTPRF,DDN,CACNA1I,CNNM2,MICA,PTPR

E,KCNG2,CTR9,KLC1,SGF29,MYO9B,CUL3,CNTNAP2,RPTOR,HDAC4,PPM1E,GABBR1,THOC7,GPD2,ITG

MEF2C,PALB2,CTNNB1,STAG1,MED27,ESRRB,MAML3,ZCCHC7,FOXO3,KDM4A,JMJD1C,RFC2,RNF111,F

F638,PRPF3,MSH5,PRKAG2,XRCC6,MEF2C,TFB1M,PALB2,CTNNB1,STAG1,FURIN,MED27,ESRRB,IVD,M

NB1,ANK3,TWF2,CPEB1,KDM4A,PPP2R2B,SYNE1,ELMO1,KIF21B,SORBS1,RHOA,DNAH11,TMOD3,TRIO

MSH5,PRKAG2,XRCC6,MEF2C,ATP23,MAD1L1,PALB2,CTNNB1,STAG1,PCGF3,MED27,ESRRB,POM121C

F638,PRPF3,MSH5,PRKAG2,XRCC6,MEF2C,TFB1M,PALB2,CTNNB1,STAG1,FURIN,MED27,ESRRB,IVD,M

NC1I1,HFE,KCNG2,CTR9,KLC1,SGF29,TNRC6A,MYO9B,CUL3,CNTNAP2,RPTOR,HDAC4,PPM1E,GABBR1

PIP5K1B,HLA-DRB5,PARD3B,HLA-DRB1,GALNT1,SYNE1,GALNT10,HLA-DQB1,CLCN3,DOCK4,CAMKV,CD

,CALD1,CTNND1,CTNNB1,NGEF,PRKG1,DPYD,TANK,CADPS,PSPH,FOXO3,PIP5K1B,PDE4B,PPP2R2B,E

/IPHOSPH9,DYSF,CNTN4,CBLB,SEMA6D,ANKS1B,MMP16,ITGA11,ATP2A2,CHRNA3,CACNA1C,RAET1E,I

-DB1,MSH5,CNOT1,XRCC6,SLC30A9,MAD1L1,DBN1,TFB1M,MAGI2,CALD1,FSCN3,CTNNB1,STAG1,ANK3

-DB1,MSH5,CNOT1,XRCC6,SLC30A9,MAD1L1,DBN1,TFB1M,MAGI2,CALD1,FSCN3,CTNNB1,STAG1,ANK3

;ACNA2D2,CLSTN2,HFE,KCNG2,SLC6A9,CNTNAP2,PKD1L3,DPY19L1,SPNS1,GABBR1,DYSF,CNTN4,SEI

,CLSTN2,HFE,KCNG2,SLC6A9,CNTNAP2,PKD1L3,DPY19L1,SPNS1,GABBR1,DYSF,SEMA6D,MMP16,EXT

PS,POM121C,GBF1,HLA-DRB5,PPP2R2B,HLA-DRB1,GALNT1,SYNE1,GALNT10,HLA-DQB1,IMMP2L,CLCN

,USP4,SGF29,RBM6,CUL3,HDAC4,PPM1E,L3MBTL4,TCF20,ZFPM2,BSN,MPHOSPH9,THOC7,NEDD4L,CE

BBS9,MRM2,F2,SPAG16,TAF1C,ANAPC4,TCF4,DYNC1I1,CLSTN2,CTR9,KLC1,SGF29,MYO9B,CUL3,HDAC

BBS9,SLC22A23,CHRNA5,CACNA2D2,CLSTN2,HFE,KCNG2,SLC6A9,CNTNAP2,PKD1L3,DPY19L1,SPNS1

BBS9,MRM2,F2,SPAG16,TAF1C,ANAPC4,TCF4,DYNC1I1,CLSTN2,CTR9,KLC1,SGF29,MYO9B,CUL3,HDAC

,PRKD1,EXOC4,DPP4,PKP4,MRM2,F2,PDE8B,SPAG16,ANAPC4,DYNC1I1,CLSTN2,HFE,USP4,KLC1,TNR

,RSRC1,RSU1,SPPL3,ESR2,MAPT,TCAIM,FADS2,TSNARE1,KIZ,SSUH2,DMXL2,PTCH1,PRKD1,PEF1,EXO

R141,FADS2,TSNARE1,TMX2,DMXL2,PTCH1,PRKD1,PEF1,EXOC4,TMEM170B,DPP4,PKP4,BBS9,SLC22

[illegible]O,HLA-DRB5,SHISA9,KCNN2,HLA-DRB1,RGS6,NTRK3,HLA-DQB1,CLCN3,ATP2B2,SORBS1,RHOA

[illegible]A11,KIF5C,ARFGEF2,CHRNA3,CACNA1C,LDB1,GATB,ARID1B,RAET1E,HLA-DQA1,PRKAG2,XRCC6,ME[illegible]

[illegible]BL2,POLR2E,ZMIZ2,TENM2,SRPK2,GATAD2B,THRB,BRWD1,TRIM33,SORBS1,FOXO6,MED8,GLI3,ZKS[illegible]

AML3,ZCCHC7,FOXO3,KDM4A,JMJD1C,RFC2,GBF1,RNF111,RBL2,POLR2E,ZMIZ2,TENM2,SRPK2,GATAI

,MAML3,ZCCHC7,FOXO3,KDM4A,JMJD1C,RFC2,RNF111,RBL2,POLR2E,ZMIZ2,TENM2,SRPK2,SYNE1,GA

AML3,ZCCHC7,FOXO3,KDM4A,JMJD1C,RFC2,GBF1,RNF111,RBL2,POLR2E,ZMIZ2,TENM2,SRPK2,GATAI

,THOC7,GPD2,SND1,FXR1,ITGA11,KIF5C,ARFGEF2,CHRNA3,CACNA1C,LDB1,GATB,ARID1B,RAET1E,PI

FGFR1,HLA-DQA1,SLC13A1,AMIGO1,GRIK1,BTN2A1,RHOJ,DBN1,MAGI2,CALD1,RASGRP4,CADM2,CTNN

,MED27,TWF2,CPEB1,ZCCHC7,KDM4A,RFC2,RNF111,KCNN2,RBL2,ZMIZ2,PPP2R2B,SRPK2,SYNE1,ELM

,MED27,TWF2,CPEB1,ZCCHC7,KDM4A,RFC2,RNF111,KCNN2,RBL2,ZMIZ2,PPP2R2B,SRPK2,SYNE1,ELM

MA6D,MMP16,EXT1,ITGA11,ATP2A2,CHRNA3,CACNA1C,RAET1E,FGFR1,HLA-DQA1,GLT8D1,SLC13A1,A

1,ITGA11,ATP2A2,CHRNA3,CACNA1C,RAET1E,FGFR1,HLA-DQA1,GLT8D1,SLC13A1,AMIGO1,SLC30A9,S

LB,SND1,ZNF398,NR1D2,ANKS1B,FXR1,ZNF823,ZMIZ1,DMTF1,CITED1,DDX27,ZSCAN12,LDB1,ARID1B,

4,PPM1E,SPNS1,ZFPM2,MPHOSPH9,DYSF,COL11A1,THOC7,NEDD4L,GPD2,NR1D2,ANKS1B,FXR1,MMF

,GABBR1,DYSF,CNTN4,SEMA6D,MMP16,EXT1,ITGA11,ATP2A2,CHRNA3,CACNA1C,RAET1E,FGFR1,HLA

4,PPM1E,SPNS1,ZFPM2,MPHOSPH9,DYSF,COL11A1,THOC7,NEDD4L,GPD2,NR1D2,ANKS1B,FXR1,MMF

C6A,MYO9B,CUL3,DOCK8,CNTNAP2,RPTOR,PCLO,HDAC4,SPNS1,MPHOSPH9,DYSF,COL11A1,PFAS,OI

C4,PLCL1,DPP4,PKP4,BBS9,MRM2,F2,PDE8B,SPAG16,ANAPC4,DYNC1I1,CLSTN2,HFE,USP4,SBK1,KLC

A23,F2,CHRNA5,CACNA2D2,CLSTN2,HFE,KCNG2,SLC6A9,CNTNAP2,PKD1L3,DPY19L1,PCLO,SPNS1,GA

F2C,ATP23,GRIK1,MAD1L1,SNAP91,CTNNB1,CACNB2,PCGF3,KCNB1,ABCB9,MED27,CACNA1I,MICA,PO

RPF3,HLA-DQA1,CNOT1,PRKAG2,XRCC6,MEF2C,ATP23,GRIK1,MAD1L1,SNAP91,CTNNB1,CACNB2,PC[illegible]

ND1,SNAP91,CD55,CTNNB1,NGEF,ADTRP,NTM,SYT1,CACNB2,KCNB1,ANK3,PTPRF,ABCB9,FURIN,DDN

MO1,RPS6KL1,PPP2R5C,KIF21B,BRWD1,DOCK4,SORBS1,GLI3,ZKSCAN8,RHOA,DNAH11,TMOD3,AFF3,T

MO1,RPS6KL1,PPP2R5C,KIF21B,BRWD1,DOCK4,SORBS1,GLI3,ZKSCAN8,RHOA,DNAH11,TMOD3,AFF3,T

,MIGO1,SLC30A9,SDK1,GRIK1,BTN2A1,MGAT5B,MAD1L1,CADM2,CD55,ST3GAL3,ADTRP,NTM,SYT1,CA(

SDK1,GRIK1,BTN2A1,MGAT5B,MAD1L1,CADM2,CD55,ST3GAL3,ADTRP,SYT1,CACNB2,KCNB1,PTPRF,AE

ZNF638,PRPF3,FGFR1,MSH5,CUEDC2,PRKAG2,XRCC6,SLC30A9,CARMIL1,CREB5,ZSCAN31,MEF2C,AT

P16,EXT1,CALU,ATP2A2,ZMIZ1,KIF5C,ARFGEF2,CHRNA3,CACNA1C,TTLL7,LDB1,ARID1B,ZNF638,PRPF3

-DQA1,GLT8D1,SLC13A1,AMIGO1,SLC30A9,SDK1,GRIK1,BTN2A1,MGAT5B,MAD1L1,CADM2,SNAP91,CD

P16,EXT1,CALU,ATP2A2,ZMIZ1,KIF5C,ARFGEF2,CHRNA3,CACNA1C,TTLL7,LDB1,ARID1B,ZNF638,PRPF3

LFM4,NEDD4L,CBLB,GPD2,SND1,SEMA6D,FXR1,MMP16,EXT1,CALU,ATP2A2,ARFGEF2,CITED1,CKB,CA

C1,FHIT,TNRC6A,MYO9B,CUL3,DOCK8,CNTNAP2,RPTOR,PCLO,TRIM26,HDAC4,PPM1E,SPNS1,GABBR1,

ABBR1,MPHOSPH9,DYSF,CNTN4,CBLB,GPD2,SEMA6D,ANKS1B,MMP16,EXT1,ITGA11,ATP2A2,ARFGEF2

M121C,FOXO3,RFC2,HLA-DRB5,SHISA9,RNF111,RBL2,POLR2E,PPP2R2B,HLA-DRB1,RGS6,SYNE1,HLA-

GF3,KCNB1,ABCB9,MED27,CACNA1I,CPEB1,MICA,POM121C,FOXO3,RFC2,HLA-DRB5,SHISA9,RNF111,R

,PRKG1,CACNA1I,SRR,CNNM2,PPP1R16B,CPEB1,MICA,PTPRO,TMEM219,HLA-DRB5,KEL,SHISA9,KCNN

CNB2,KCNB1,PTPRF,ABCB9,FURIN,CACNA1I,CNNM2,MICA,LRFN5,POM121C,SLC35F2,RFT1,PTPRO,CL

CB9,FURIN,CACNA1I,CNNM2,MICA,LRFN5,POM121C,SLC35F2,RFT1,PTPRO,CLEC17A,TMEM219,HLA-D

P23,ZNF568,ZNF19,DDX3Y,MAD1L1,ZKSCAN3,MAGI2,RASGRP4,PALB2,CTNND1,SOX5,CTNNB1,NEK4,S

,MSH5,HLA-DQA1,CNOT1,PRKAG2,XRCC6,MEF2C,ATP23,MGAT5B,MAD1L1,DBN1,TFB1M,MAGI2,CALD

55,CTNNB1,ST3GAL3,ADTRP,NTM,SYT1,CACNB2,KCNB1,ANK3,PTPRF,ABCB9,FURIN,DDN,CACNA1I,C

,MSH5,HLA-DQA1,CNOT1,PRKAG2,XRCC6,MEF2C,ATP23,MGAT5B,MAD1L1,DBN1,TFB1M,MAGI2,CALD

CNA1C,GATB,NT5C2,FGFR1,HLA-DQA1,GLT8D1,CNOT1,PRKAG2,ETF1,CARMIL1,GMIP,MGAT5B,RHOJ,

,ZFPM2,BSN,MPHOSPH9,DYSF,COL11A1,PFAS,OLFM4,THOC7,NEDD4L,CBLB,GPD2,SND1,SEMA6D,AN

,CHRNA3,CACNA1C,RAET1E,FGFR1,HLA-DQA1,GLT8D1,CNOT1,SLC13A1,XRCC6,AMIGO1,SLC30A9,SD

DQB1,IMMP2L,PPP2R5C,KIF21B,SORBS1,FOXO6,NDUFA2,MED8,GLI3,PIK3R2,DNAH11,HDAC9

BL2,POLR2E,PPP2R2B,HLA-DRB1,RGS6,SYNE1,HLA-DQB1,IMMP2L,RPS6KL1,PPP2R5C,KIF21B,SORBS

N2,BTN3A2,HLA-DRB1,RGS6,PPP1R13B,TENM2,NTRK3,NEGR1,HLA-DQB1,PPP1R16A,ELMO1,WNT3,PC[illegible]

EC17A,TMEM219,HLA-DRB5,KEL,SHISA9,KCNN2,SORCS3,BTN3A2,HLA-DRB1,TENM2,NTRK3,GALNT1,N

RB5,KEL,SHISA9,KCNN2,SORCS3,BTN3A2,HLA-DRB1,TENM2,NTRK3,GALNT1,SYNE1,GALNT10,HLA-DC

TAG1,PCGF3,MED27,DDN,PPP1R16B,ESRRB,POM121C,MAML3,BCL11A,ZCCHC7,TSHZ3,FOXO3,LCOR

1,PALB2,CTNNB1,ST3GAL3,STAG1,SYT1,PCGF3,ANK3,ABCB9,FURIN,MED27,DDN,TWF2,ESRRB,CPEB

NNM2,MICA,LRFN5,POM121C,SLC35F2,RFT1,PTPRO,CLEC17A,TMEM219,HLA-DRB5,KEL,SHISA9,KCNN

1,PALB2,CTNNB1,ST3GAL3,STAG1,SYT1,PCGF3,ABCB9,FURIN,MED27,DDN,ESRRB,CPEB1,IVD,CADPS

MAD1L1,TFB1M,MAGI2,CALD1,CTNND1,SNAP91,CTNNB1,ST3GAL3,NGEF,SYT1,ANK3,ABCB9,FURIN,DD

KS1B,FXR1,MMP16,EXT1,CALU,ATP2A2,ZMIZ1,DMTF1,KIF5C,ARFGEF2,CITED1,CKB,CACNA1C,TTLL7,G

K1,GRIK1,BTN2A1,MGAT5B,RHOJ,MAD1L1,DBN1,MAGI2,CALD1,RASGRP4,CADM2,CTNND1,SNAP91,CD

DH9,CLCN3,SLC39A8,PREX1,ATP2B2,FAM193A,TRAIP,IGSF9B,SORBS1,NEURL1,RHOA,BTN3A1,CAMKV

EGR1,SYNE1,GALNT10,HLA-DQB1,IMMP2L,PCDH9,CLCN3,SLC39A8,ATP2B2,BTN2A2,TENM4,IGSF9B,S

QB1,IMMP2L,PCDH9,CLCN3,SLC39A8,ATP2B2,BTN2A2,TENM4,IGSF9B,SORBS1,BTN3A1,NAALADL2,RN

L,KDM4A,JMJD1C,RFC2,ZNF600,PDE4B,RNF111,RBL2,POLR2E,ZMIZ2,PPP1R13B,TENM2,NFIX,SRPK2,S

,IVD,CADPS,POM121C,MAML3,ZCCHC7,FOXO3,KDM4A,JMJD1C,RFC2,GBF1,HLA-DRB5,RNF111,KCNN

2,SORCS3,BTN3A2,HLA-DRB1,RGS6,TENM2,NTRK3,GALNT1,NEGR1,SYNE1,GALNT10,HLA-DQB1,IMMP

,POM121C,MAML3,ZCCHC7,FOXO3,KDM4A,JMJD1C,RFC2,GBF1,HLA-DRB5,RNF111,RBL2,POLR2E,ZFY

N,PRKG1,CTTNBP2,TWF2,DPYD,CPEB1,TANK,IVD,CADPS,POM121C,TSHZ3,PSPH,FOXO3,GBF1,PIP5K

ATB,NT5C2,ZNF638,PRPF3,FGFR1,HLA-DQA1,GLT8D1,CUEDC2,CNOT1,GUCY1A2,PRKAG2,XRCC6,ET

55,CTNNB1,ST3GAL3,NGEF,ADTRP,NTM,SYT1,CACNB2,KCNB1,ANK3,PTPRF,ABCB9,FURIN,DDN,PRK

;YNE1,GATAD2B,ZNF615,DPP8,NOVA1,IMMP2L,THRB,PPP2R5C,BRWD1,TENM4,KDM3B,TRIM33,SORBS

2,RBL2,POLR2E,ZFYVE1,ZMIZ2,PPP2R2B,HLA-DRB1,TENM2,SRPK2,GALNT1,SYNE1,GALNT10,GATAD2I

2L,WNT3,PCDH9,CLCN3,SLC39A8,ATP2B2,BTN2A2,TENM4,IGSF9B,SORBS1,NDUFA2,RHOA,BTN3A1,N/

VE1,ZMIZ2,PPP2R2B,HLA-DRB1,TENM2,SRPK2,GALNT1,SYNE1,GALNT10,GATAD2B,HLA-DQB1,IMMP2L

(1B,HLA-DRB5,PDE4B,KCNN2,ZFYVE1,ZMIZ2,PPP2R2B,HLA-DRB1,TENM2,MANBA,GALNT1,SYNE1,GALI

=1,CARMIL1,MEF2C,GMIP,DDX3Y,MGAT5B,RHOJ,MAD1L1,DBN1,TFB1M,MAGI2,CALD1,RASGRP4,CADM

31,CACNA1I,SRR,CNNM2,PPP1R16B,CPEB1,CADPS,MICA,LRFN5,POM121C,SLC35F2,FOXO3,RFT1,PTF

:1,BCL11B,FOXO6,CSRNP3,MED8,GLI3,ZKSCAN8,RHOA,AFF3,TRIOBP,HDAC9,ZNF800,TLK1

3,HLA-DQB1,IMMP2L,THRB,CLCN3,SLC39A8,PPA2,PPP2R5C,KIF21B,BRWD1,DOCK4,TRIM33,SORBS1,F

.,THRB,CLCN3,PPA2,PPP2R5C,KIF21B,BRWD1,TRIM33,SORBS1,FOXO6,NDUFA2,MED8,GLI3,ZKSCAN8,

NT10,HLA-DQB1,ELMO1,IMMP2L,WNT3,CLCN3,PPA2,PREX1,ATP2B2,RPS6KL1,DOCK4,TRAIP,SORBS1,I

I2,CTNND1,FSCN3,SNAP91,IMPA2,CTNNB1,ST3GAL3,NGEF,SYT1,ANK3,ABCB9,FURIN,MED27,DDN,PRK
PRO,CLEC17A,TMEM219,GBF1,PIP5K1B,HLA-DRB5,KEL,NKIRAS1,SHISA9,SEMA3F,KCNN2,SORCS3,BTN

FOXO6,AMBRA1,NDUFA2,GLI3,NEURL1,RHOA,PIK3R2,DNAH11,TMOD3,MAIP1,RNF144A,CAMKV,CDKAL

KG1,SRR,CTTNBP2,TWF2,PLCL2,DPYD,CPEB1,TANK,IVD,CADPS,MICA,POM121C,BCL11A,ZCCHC7,TSH

I3A2,PARD3B,PPP2R2B,HLA-DRB1,RGS6,PPP1R13B,TENM2,NTRK3,GALNT1,NEGR1,SYNE1,GALNT10,H

Z3,PSPH,FOXO3,KDM4A,GBF1,PIP5K1B,HLA-DRB5,PDE4B,NKIRAS1,RNF111,KCNN2,RBL2,PMFBP1,ZFY
ILA-DQB1,PPP1R16A,ELMO1,DPP8,IMMP2L,WNT3,PCDH9,CLCN3,SLC39A8,PREX1,ATP2B2,BTN2A2,TEN

NM4,DOCK4,F
