## Supplemental table 3 for "Digestive Dimensions of Autism: A Multiscale Exploration of Gut-Brain Interactions"

| Pathway | Total | Expected | Hits | P.Value | FDR | Genes |
|---|---|---|---|---|---|---|
| Rho GTPase activator activity | 39 | 1.04 | 5 | 0.00362 | 1 | ARHGAP15,MYO9B,GMIP,PREX1,[illegible] |
| Transmembrane receptor protein phosphatase activity | 18 | 0.481 | 3 | 0.0115 | 1 | PTPRN2,PTPRF,PTPRO |
| Steroid hormone receptor activity | 56 | 1.5 | 5 | 0.0166 | 1 | ESR2,ESRRG,NR1D2,ESRRB,THR[illegible] |
| Calmodulin binding | 171 | 4.57 | 10 | 0.0167 | 1 | NRGN,SLC8A3,PDE1C,MYO9B,CA[illegible] |
| Ligand_activated sequence_specific DNA binding RNA polymerase II transcription factor activity | 57 | 1.52 | 5 | 0.0178 | 1 | ESR2,ESRRG,NR1D2,ESRRB,THR[illegible] |
| Protein homodimerization activity | 573 | 15.3 | 24 | 0.0201 | 1 | MTHFD1L,ANXA9,KMT2A,CISD2,F[illegible] |
| Voltage_gated calcium channel activity | 41 | 1.1 | 4 | 0.0234 | 1 | CACNA2D2,CACNA1C,CACNB2,CA[illegible] |
| Calcium ion transmembrane transporter activity | 130 | 3.47 | 8 | 0.0234 | 1 | SLC8A3,GRIN2A,CACNA2D2,ATP2[illegible] |
| SMAD binding | 68 | 1.82 | 5 | 0.0351 | 1 | CITED1,MAGI2,CTNNB1,RNF111,T[illegible] |
| Ras GTPase activator activity | 117 | 3.13 | 7 | 0.0375 | 1 | ARHGAP15,SYNGAP1,MYO9B,GM[illegible] |
| Double_stranded DNA binding | 149 | 3.98 | 8 | 0.0468 | 1 | FOXP1,ZNF638,MSH5,XRCC6,CTN[illegible] |

| | | | | | | |
|---|---|---|---|---|---|---|
| Microtubule motor activity | 75 | 2 | 5 | 0.0501 | 1 | DYNC1I1,KLC1,KIF5C,KIF21B,DNA |
| PDZ domain binding | 76 | 2.03 | 5 | 0.0525 | 1 | SRR,CCDC88C,CLCN3,ATP2B2,DC |
| Structure_specific DNA binding | 242 | 6.47 | 11 | 0.0608 | 1 | RPA3,FOXP1,WRN,ZNF638,MSH5, |
| 3',5'_cyclic_nucleotide phosphodiesterase activity | 34 | 0.909 | 3 | 0.0616 | 1 | PDE1C,PDE8B,PDE4B |
| 3',5'_cyclic_nucleotide phosphodiesterase activity | 34 | 0.909 | 3 | 0.0616 | 1 | PDE1C,PDE8B,PDE4B |
| Oxidoreductase activity, acting on paired donors, with incorporation or reduction of molecular oxygen, 2_oxoglutarate as one donor, and incorporation of one atom each of oxygen into both donors | 58 | 1.55 | 4 | 0.0691 | 1 | KDM5D,UTY,TET2,KDM4A |
| Acetylgalactosaminyltransferase activity | 37 | 0.989 | 3 | 0.0755 | 1 | GALNT12,GALNT1,GALNT10 |

| | | | | | | |
|---|---|---|---|---|---|---|
| Excitatory extracellular ligand_gated ion channel activity | 61 | 1.63 | 4 | 0.08 | 1 | GRIN2A,CHRNA5,CHRNA3,GRIK1 |
| Motor activity | 143 | 3.82 | 7 | 0.0891 | 1 | MYO18A,DYNC1I1,KLC1,MYO9B,K[illegible] |
| Transcription cofactor activity | 552 | 14.8 | 20 | 0.104 | 1 | KMT2E,ESR2,TCF4,HDAC4,TCF20 |
| Acetylcholine_activated cation_selective channel activity | 21 | 0.561 | 2 | 0.107 | 1 | CHRNA5,CHRNA3 |
| Protein phosphatase type 2A regulator activity | 22 | 0.588 | 2 | 0.116 | 1 | PPP2R2B,PPP2R5C |
| Ionotropic glutamate receptor activity | 23 | 0.615 | 2 | 0.125 | 1 | GRIN2A,GRIK1 |
| ADP binding | 23 | 0.615 | 2 | 0.125 | 1 | MYO9B,PRKAG2 |
| Transcription coactivator activity | 312 | 8.34 | 12 | 0.133 | 1 | KMT2E,ESR2,TCF20,ZFPM2,CITE[illegible] |
| Protein phosphatase binding | 74 | 1.98 | 4 | 0.136 | 1 | ANAPC4,CTNND1,CTNNB1,PPP1R[illegible] |
| Protein C_terminus binding | 160 | 4.28 | 7 | 0.137 | 1 | MAGI1,TCF4,ATP2A2,CITED1,XRC[illegible] |
| Cation binding | 4160 | 111 | 121 | 0.144 | 1 | FKBP9,KDM5D,PDE1C,ANXA9,KM[illegible] |
| Phospholipid binding | 514 | 13.7 | 18 | 0.148 | 1 | ANXA9,ARHGAP15,SYNGAP1,PRK[illegible] |
| Deacetylase activity | 52 | 1.39 | 3 | 0.162 | 1 | HDAC4,MACROD2,HDAC9 |
| DNA_dependent ATPase activity | 80 | 2.14 | 4 | 0.166 | 1 | MYO18A,WRN,MSH5,XRCC6 |

| | | | | | | |
|---|---|---|---|---|---|---|
| Calcium channel activity | 111 | 2.97 | 5 | 0.176 | 1 | GRIN2A,CACNA2D2,CACNA1C,CA |
| Steroid binding | 82 | 2.19 | 4 | 0.176 | 1 | ESR2,ESRRG,PTCH1,ESRRB |
| Protein binding transcription factor activity | 600 | 16 | 20 | 0.183 | 1 | KMT2E,ESR2,TCF4,HDAC4,TCF20 |
| N_methyltransferase activity | 84 | 2.24 | 4 | 0.187 | 1 | KMT2D,KMT2A,KMT2E,TFB1M |
| Neurotransmitter receptor activity | 85 | 2.27 | 4 | 0.192 | 1 | ANXA9,CHRNA5,CHRNA3,SORCS |
| Phosphatase binding | 116 | 3.1 | 5 | 0.199 | 1 | ANAPC4,MAGI2,CTNND1,CTNNB1 |
| Protein kinase binding | 376 | 10 | 13 | 0.208 | 1 | FEZ1,MAPT,KIZ,SPAG16,RPTOR, |
| SH3 domain binding | 118 | 3.15 | 5 | 0.208 | 1 | SYNGAP1,MAPT,CTTNBP2,ELMO1 |
| Histone methyltransferase activity | 60 | 1.6 | 3 | 0.216 | 1 | KMT2D,KMT2A,KMT2E |
| Phosphoric diester hydrolase activity | 120 | 3.21 | 5 | 0.218 | 1 | PDE1C,PLCL1,PDE8B,PLCL2,PDE |
| Phosphoric diester hydrolase activity | 120 | 3.21 | 5 | 0.218 | 1 | PDE1C,PLCL1,PDE8B,PLCL2,PDE |
| Delayed rectifier potassium channel activity | 33 | 0.882 | 2 | 0.22 | 1 | KCNG2,KCNB1 |
| Metalloendopeptidase activity | 123 | 3.29 | 5 | 0.233 | 1 | ADAM22,THSD4,MMP16,ATP23,KE |
| S_adenosylmethionine_dependent methyltransferase activity | 123 | 3.29 | 5 | 0.233 | 1 | KMT2D,KMT2A,KMT2E,MRM2,TFB |

| | | | | | | |
|---|---|---|---|---|---|---|
| Small GTPase regulator activity | 319 | 8.53 | 11 | 0.235 | 1 | ARHGAP15,SYNGAP1,MYO9B,ARI |
| ATP_dependent DNA helicase activity | 36 | 0.962 | 2 | 0.25 | 1 | WRN,XRCC6 |
| Histone deacetylase binding | 65 | 1.74 | 3 | 0.251 | 1 | HDAC4,GLI3,HDAC9 |
| Antigen binding | 66 | 1.76 | 3 | 0.259 | 1 | ABCB9,HLA-DRB5,HLA-DRB1 |
| Aminopeptidase activity | 37 | 0.989 | 2 | 0.26 | 1 | DPP4,DPP8 |
| Regulation of DNA_dependent transcription, elongation | 37 | 0.989 | 2 | 0.26 | 1 | CTR9,LDB1 |
| Heparin binding | 130 | 3.47 | 5 | 0.268 | 1 | PTCH1,MDK,FGFR1,PTPRF,TNXB |
| Glutamate receptor activity | 38 | 1.02 | 2 | 0.27 | 1 | GRIN2A,GRIK1 |
| Voltage_gated cation channel activity | 163 | 4.36 | 6 | 0.271 | 1 | CACNA2D2,KCNG2,CACNA1C,CA( |
| Protein dimerization activity | 996 | 26.6 | 30 | 0.273 | 1 | MTHFD1L,NPAS3,ANXA9,KMT2A,C |
| Acetylcholine binding | 12 | 0.321 | 1 | 0.278 | 1 | CHRNA3 |
| Protein deacetylase activity | 39 | 1.04 | 2 | 0.28 | 1 | HDAC4,HDAC9 |
| Positive regulation of transcription, DNA_dependent | 1260 | 33.6 | 37 | 0.293 | 1 | NPAS3,KMT2D,KMT2A,KMT2E,FO) |
| Chromatin binding | 338 | 9.03 | 11 | 0.295 | 1 | RERE,FOXP1,DMTF1,CITED1,LDB |
| Phospholipase C activity | 41 | 1.1 | 2 | 0.3 | 1 | PLCL1,PLCL2 |

| | | | | | | |
|---|---|---|---|---|---|---|
| Extracellular ligand_gated ion channel activity | 104 | 2.78 | 4 | 0.303 | 1 | GRIN2A,CHRNA5,CHRNA3,GRIK1 |
| Metal ion transmembrane transporter activity | 451 | 12.1 | 14 | 0.321 | 1 | SLC8A3,GRIN2A,CACNA2D2,KCN[illegible] |
| Transcription corepressor activity | 208 | 5.56 | 7 | 0.322 | 1 | TCF4,HDAC4,ZFPM2,LDB1,BCL11[illegible] |
| Cation channel activity | 347 | 9.27 | 11 | 0.325 | 1 | GRIN2A,CHRNA5,CACNA2D2,KCN |
| Translation factor activity, nucleic acid binding | 108 | 2.89 | 4 | 0.327 | 1 | EFL1,GATB,EIF1AY,ETF1 |
| Kinase binding | 418 | 11.2 | 13 | 0.327 | 1 | FEZ1,MAPT,KIZ,SPAG16,RPTOR,[illegible] |
| ATPase activity, coupled | 313 | 8.36 | 10 | 0.327 | 1 | MYO18A,WRN,ATP2A2,DDX27,MS |
| Voltage_gated channel activity | 211 | 5.64 | 7 | 0.335 | 1 | CACNA2D2,KCNG2,CACNA1C,CA[illegible] |
| Zinc ion binding | 2010 | 53.8 | 57 | 0.337 | 1 | KDM5D,KMT2D,KMT2A,MSL2,KMT |
| RNA polymerase II distal enhancer sequence_specific DNA binding transcription factor activity | 110 | 2.94 | 4 | 0.339 | 1 | FOXP1,FOXO3,NFIX,FOXO6 |
| Gated channel activity | 389 | 10.4 | 12 | 0.347 | 1 | GRIN2A,CHRNA5,CACNA2D2,KCN |
| ATP_dependent helicase activity | 112 | 2.99 | 4 | 0.351 | 1 | WRN,DDX27,XRCC6,DDX3Y |
| Collagen binding | 47 | 1.26 | 2 | 0.359 | 1 | ITGA11,TNXB |

| | | | | | | |
|---|---|---|---|---|---|---|
| Acetylglucosaminyltransferase activity | 47 | 1.26 | 2 | 0.359 | 1 | EXT1,MGAT5B |
| Lipid binding | 788 | 21.1 | 23 | 0.361 | 1 | ANXA9,ARHGAP15,SYNGAP1,ESR |
| Guanyl_nucleotide exchange factor activity | 218 | 5.83 | 7 | 0.365 | 1 | DOCK8,ARFGEF2,RASGRP4,NGEI |
| Telomeric DNA binding | 17 | 0.454 | 1 | 0.369 | 1 | XRCC6 |
| Magnesium ion binding | 188 | 5.02 | 6 | 0.388 | 1 | WRN,MAST3,SRR,PSPH,SRPK2,PI |
| UDP_glycosyltransferase activity | 153 | 4.09 | 5 | 0.389 | 1 | GALNT12,EXT1,MGAT5B,GALNT1, |
| Identical protein binding | 910 | 24.3 | 26 | 0.391 | 1 | MTHFD1L,ANXA9,KMT2A,CISD2,F |
| Sequence_specific DNA binding | 732 | 19.6 | 21 | 0.4 | 1 | KMT2A,RERE,FOXP1,ESR2,ESRR |
| Taste receptor activity | 19 | 0.508 | 1 | 0.403 | 1 | PKD1L3 |
| Neurotransmitter binding | 19 | 0.508 | 1 | 0.403 | 1 | CHRNA3 |
| Protein methyltransferase activity | 86 | 2.3 | 3 | 0.405 | 1 | KMT2D,KMT2A,KMT2E |
| Protein N_terminus binding | 86 | 2.3 | 3 | 0.405 | 1 | FEZ1,EXOC4,KCNB1 |
| Cation_transporting ATPase activity | 52 | 1.39 | 2 | 0.407 | 1 | ATP2A2,ATP2B2 |
| GTPase activator activity | 267 | 7.14 | 8 | 0.422 | 1 | ARHGAP15,SYNGAP1,MYO9B,GMI |
| DNA helicase activity | 54 | 1.44 | 2 | 0.425 | 1 | WRN,XRCC6 |

| | | | | | | |
|---|---|---|---|---|---|---|
| ATPase activity, coupled to transmembrane movement of ions, phosphorylative mechanism | 54 | 1.44 | 2 | 0.425 | 1 | ATP2A2,ATP2B2 |
| Phosphatidylinositol binding | 160 | 4.28 | 5 | 0.426 | 1 | SNX29,SNAP91,SYT1,TWF2,ZFYVI |
| Protein domain specific binding | 560 | 15 | 16 | 0.428 | 1 | SYNGAP1,MAPT,ESRRG,CTR9,CIT |
| Ras GTPase binding | 125 | 3.34 | 4 | 0.43 | 1 | DGKI,DMXL2,DOCK4,TRIOBP |
| Oxidoreductase activity, acting on paired donors, with incorporation or reduction of molecular oxygen | 197 | 5.26 | 6 | 0.431 | 1 | KDM5D,AGMO,UTY,TET2,FADS2,K |
| ATPase activity | 417 | 11.1 | 12 | 0.438 | 1 | MYO18A,WRN,MYO9B,ATP2A2,DD |
| Transition metal ion binding | 2270 | 60.7 | 62 | 0.451 | 1 | KDM5D,KMT2D,KMT2A,MSL2,KMT |
| Enzyme binding | 1200 | 32 | 33 | 0.456 | 1 | DGKI,KMT2E,FEZ1,ESR2,MAPT,KI |
| Ribonucleoprotein complex binding | 58 | 1.55 | 2 | 0.462 | 1 | EFL1,ETF1 |
| GABA receptor activity | 23 | 0.615 | 1 | 0.464 | 1 | GABBR1 |
| Phosphorus_oxygen lyase activity | 23 | 0.615 | 1 | 0.464 | 1 | GUCY1A2 |
| Cyclase activity | 24 | 0.641 | 1 | 0.478 | 1 | GUCY1A2 |

| | | | | | | |
|---|---|---|---|---|---|---|
| Protein binding, bridging | 135 | 3.61 | 4 | 0.489 | 1 | COL11A1,FSCN3,ANK3,SORBS1 |
| Antiporter activity | 62 | 1.66 | 2 | 0.497 | 1 | SLC8A3,CLCN3 |
| Neutral amino acid transmembrane transporter activity | 26 | 0.695 | 1 | 0.506 | 1 | SLC6A9 |
| Lipoprotein particle binding | 26 | 0.695 | 1 | 0.506 | 1 | MAPT |
| Small GTPase binding | 138 | 3.69 | 4 | 0.506 | 1 | DGKI,DMXL2,DOCK4,TRIOBP |
| Glycosaminoglycan binding | 178 | 4.76 | 5 | 0.519 | 1 | PTCH1,MDK,FGFR1,PTPRF,TNXB |
| GTPase regulator activity | 517 | 13.8 | 14 | 0.519 | 1 | DGKI,ARHGAP15,SYNGAP1,MYO9[illegible] |
| Sialyltransferase activity | 27 | 0.722 | 1 | 0.519 | 1 | ST3GAL3 |
| Monosaccharide binding | 65 | 1.74 | 2 | 0.522 | 1 | CLEC17A,MANBA |
| Calcium ion binding | 673 | 18 | 18 | 0.533 | 1 | FKBP9,ANXA9,CDHR4,PEF1,PLCL[illegible] |
| Thyroid hormone receptor binding | 29 | 0.775 | 1 | 0.544 | 1 | JMJD1C |
| Serine_type peptidase activity | 222 | 5.93 | 6 | 0.547 | 1 | DPP6,DPP4,F2,FURIN,DPP8,IMMP[illegible] |
| Serine hydrolase activity | 224 | 5.99 | 6 | 0.555 | 1 | DPP6,DPP4,F2,FURIN,DPP8,IMMP[illegible] |
| Protein complex binding | 339 | 9.06 | 9 | 0.556 | 1 | GNAI1,ADAM22,WRN,PTCH1,RPT[illegible] |
| Negative regulation of transcription, DNA_dependent | 987 | 26.4 | 26 | 0.56 | 1 | KMT2D,FOXP1,ESR2,UTY,PTCH1,[illegible] |

| | | | | | | |
|---|---|---|---|---|---|---|
| Binding, bridging | 148 | 3.96 | 4 | 0.561 | 1 | COL11A1,FSCN3,ANK3,SORBS1 |
| Actin filament binding | 71 | 1.9 | 2 | 0.57 | 1 | FSCN3,TR IOBP |
| GTPase binding | 150 | 4.01 | 4 | 0.572 | 1 | DGKI,DMXL2,DOCK4,TRIOBP |
| RNA polymeras e II transcriptio n cofactor activity | 73 | 1.95 | 2 | 0.585 | 1 | ZFPM2,ME D8 |
| Cation transmemb rane transporter activity | 694 | 18.5 | 18 | 0.587 | 1 | SLC8A3,GRIN2A,CHRNA5,CACNA2 |
| Ion channel activity | 503 | 13.4 | 13 | 0.589 | 1 | GRIN2A,CHRNA5,CACNA2D2,KCN |
| SH2 domain binding | 33 | 0.882 | 1 | 0.591 | 1 | CTR9 |
| Anion:catio n symporter activity | 33 | 0.882 | 1 | 0.591 | 1 | SLC13A1 |
| Microtubul e binding | 115 | 3.07 | 3 | 0.597 | 1 | MAPT,DYNC1I1,CCDC88C |
| Cytoskelet al protein binding | 738 | 19.7 | 19 | 0.601 | 1 | MAGI1,FEZ1,MAPT,DYNC1I1,MYO |
| Transcripti on factor binding | 509 | 13.6 | 13 | 0.607 | 1 | FOXP1,TCF4,RPTOR,HDAC4,ZFP |
| Hydrolase activity, acting on carbon_nitr ogen (but not peptide) bonds, in linear amides | 76 | 2.03 | 2 | 0.607 | 1 | HDAC4,H DAC9 |
| Ligase activity, forming carbon_nitr ogen bonds | 393 | 10.5 | 10 | 0.608 | 1 | MTHFD1L,ANAPC4,CUL3,PFAS,NE |

| | | | | | | |
|---|---|---|---|---|---|---|
| ATPase activity, coupled to transmembrane movement of ions | 77 | 2.06 | 2 | 0.614 | 1 | ATP2A2,ATP2B2 |
| Peptide binding | 159 | 4.25 | 4 | 0.618 | 1 | ABCB9,FURIN,HLA-DRB5,HLA-DRE |
| Pyrophosphatase activity | 938 | 25.1 | 24 | 0.619 | 1 | MYO18A,GNAI1,WRN,EFL1,DYNC1 |
| Protein tyrosine phosphatase activity | 119 | 3.18 | 3 | 0.62 | 1 | PTPRN2,PTPRF,PTPRO |
| Hydrolase activity, acting on acid anhydrides, catalyzing transmembrane movement of substances | 119 | 3.18 | 3 | 0.62 | 1 | ATP2A2,ABCB9,ATP2B2 |
| Structural constituent of cytoskeleton | 78 | 2.08 | 2 | 0.621 | 1 | MAPT,ANK3 |
| Rho guanyl_nucleotide exchange factor activity | 78 | 2.08 | 2 | 0.621 | 1 | NGEF,PREX1 |
| Substrate_specific channel activity | 515 | 13.8 | 13 | 0.624 | 1 | GRIN2A,CHRNA5,CACNA2D2,KCN |
| Hydrolase activity, acting on acid anhydrides | 944 | 25.2 | 24 | 0.631 | 1 | MYO18A,GNAI1,WRN,EFL1,DYNC1 |

| | | | | | | |
|---|---|---|---|---|---|---|
| Ras guanyl_nucleotide exchange factor activity | 121 | 3.23 | 3 | 0.632 | 1 | RASGRP4,NGEF,PREX1 |
| ATPase activity, coupled to movement of substances | 121 | 3.23 | 3 | 0.632 | 1 | ATP2A2,ABCB9,ATP2B2 |
| Phosphatase regulator activity | 80 | 2.14 | 2 | 0.635 | 1 | PPP2R2B,PPP2R5C |
| Oxidoreductase activity, acting on the CH_CH group of donors | 80 | 2.14 | 2 | 0.635 | 1 | DPYD,IVD |
| Exopeptidase activity | 123 | 3.29 | 3 | 0.643 | 1 | DPP6,DPP4,DPP8 |
| 3'_5' exonuclease activity | 38 | 1.02 | 1 | 0.643 | 1 | WRN |
| Primary active transmembrane transporter activity | 125 | 3.34 | 3 | 0.653 | 1 | ATP2A2,ABCB9,ATP2B2 |
| Growth factor binding | 125 | 3.34 | 3 | 0.653 | 1 | FGFR1,FURIN,NTRK3 |
| Sterol binding | 40 | 1.07 | 1 | 0.662 | 1 | PTCH1 |
| Channel regulator activity | 85 | 2.27 | 2 | 0.667 | 1 | NEDD4L,PRKG1 |
| Actin binding | 373 | 9.97 | 9 | 0.67 | 1 | MYO9B,DBN1,CALD1,FSCN3,TWF[illegible] |
| Protein heterodimerization activity | 374 | 10 | 9 | 0.673 | 1 | FOXP1,PEF1,TCF4,EXT1,MEF2C,S[illegible] |
| Integrin binding | 86 | 2.3 | 2 | 0.674 | 1 | ADAM22,TNXB |

| | | | | | | |
|---|---|---|---|---|---|---|
| Nucleoside _triphosphatase activity | 888 | 23.7 | 22 | 0.676 | 1 | MYO18A,GNAI1,WRN,EFL1,DYNC1 |
| Phosphoric ester hydrolase activity | 496 | 13.3 | 12 | 0.679 | 1 | PDE1C,PTPRN2,PLCL1,PDE8B,PP |
| Transmembrane receptor protein tyrosine kinase activity | 88 | 2.35 | 2 | 0.686 | 1 | FGFR1,NTRK3 |
| Ion binding | 6140 | 164 | 160 | 0.687 | 1 | FKBP9,MTHFD1L,KDM5D,MAGI1,P |
| Helicase activity | 176 | 4.7 | 4 | 0.696 | 1 | WRN,DDX27,XRCC6,DDX3Y |
| RNA polymerase activity | 44 | 1.18 | 1 | 0.697 | 1 | POLR2E |
| Endopeptidase activity | 424 | 11.3 | 10 | 0.701 | 1 | ADAM22,THSD4,PSMA5,SPPL3,DF |
| ATP binding | 1490 | 39.8 | 37 | 0.705 | 1 | MTHFD1L,MAGI1,MYO18A,DGKI,VI |
| Methyltransferase activity | 222 | 5.93 | 5 | 0.711 | 1 | KMT2D,KMT2A,KMT2E,MRM2,TFB |
| Tubulin binding | 180 | 4.81 | 4 | 0.713 | 1 | FEZ1,MAPT,DYNC1I1,CCDC88C |
| Oxidoreductase activity, acting on peroxide as acceptor | 46 | 1.23 | 1 | 0.713 | 1 | PXDNL |
| Inorganic cation transmembrane transporter activity | 550 | 14.7 | 13 | 0.714 | 1 | SLC8A3,GRIN2A,CACNA2D2,KCN( |
| Inorganic cation transmembrane transporter activity | 550 | 14.7 | 13 | 0.714 | 1 | SLC8A3,GRIN2A,CACNA2D2,KCN( |
| Metallopeptidase activity | 224 | 5.99 | 5 | 0.719 | 1 | ADAM22,THSD4,MMP16,ATP23,KE |

| | | | | | | |
|---|---|---|---|---|---|---|
| Protein kinase inhibitor activity | 47 | 1.26 | 1 | 0.721 | 1 | PRKAG2 |
| Ligand_gated channel activity | 182 | 4.86 | 4 | 0.721 | 1 | GRIN2A,CHRNA5,CHRNA3,GRIK1 |
| Enzyme activator activity | 436 | 11.7 | 10 | 0.733 | 1 | ARHGAP15,SYNGAP1,MYO9B,MM |
| Potassium channel activity | 142 | 3.79 | 3 | 0.736 | 1 | KCNG2,KCNB1,KCNN2 |
| Neuropeptide receptor activity | 50 | 1.34 | 1 | 0.743 | 1 | SORCS3 |
| MRNA binding | 99 | 2.65 | 2 | 0.747 | 1 | FXR1,NOVA1 |
| Voltage_gated potassium channel activity | 99 | 2.65 | 2 | 0.747 | 1 | KCNG2,KCNB1 |
| Kinase inhibitor activity | 51 | 1.36 | 1 | 0.749 | 1 | PRKAG2 |
| Transferase activity, transferring one_carbon groups | 233 | 6.23 | 5 | 0.75 | 1 | KMT2D,KMT2A,KMT2E,MRM2,TFB |
| Kinase activator activity | 52 | 1.39 | 1 | 0.756 | 1 | PRKAG2 |
| SH3/SH2 adaptor activity | 52 | 1.39 | 1 | 0.756 | 1 | SORBS1 |
| Adenyl ribonucleotide binding | 1530 | 40.8 | 37 | 0.761 | 1 | MTHFD1L,MAGI1,MYO18A,DGKI,V |
| Transferase activity, transferring hexosyl groups | 237 | 6.33 | 5 | 0.763 | 1 | GALNT12,EXT1,MGAT5B,GALNT1, |
| Ligase activity | 572 | 15.3 | 13 | 0.764 | 1 | MTHFD1L,MSL2,ANAPC4,CUL3,PF |
| Adenyl nucleotide binding | 1530 | 40.9 | 37 | 0.766 | 1 | MTHFD1L,MAGI1,MYO18A,DGKI,V |

| | | | | | | |
|---|---|---|---|---|---|---|
| Solute:sodium symporter activity | 54 | 1.44 | 1 | 0.769 | 1 | SLC6A9 |
| GTPase activity | 241 | 6.44 | 5 | 0.776 | 1 | GNAI1,EFL1,RHOJ,NKIRAS1,RHO[illegible] |
| Serine_type endopeptidase inhibitor activity | 107 | 2.86 | 2 | 0.784 | 1 | ITIH3,FURIN |
| Transferase activity, transferring glycosyl groups | 333 | 8.9 | 7 | 0.791 | 1 | GALNT12,EXT1,GLT8D1,MGAT5B,[illegible] |
| Ubiquitin_protein ligase activity | 291 | 7.78 | 6 | 0.794 | 1 | ANAPC4,CUL3,NEDD4L,CBLB,RN[illegible] |
| Nucleobase_containing compound kinase activity | 59 | 1.58 | 1 | 0.798 | 1 | NT5C2 |
| Small conjugating protein_specific protease activity | 59 | 1.58 | 1 | 0.798 | 1 | USP4 |
| Protein serine/threonine phosphatase activity | 60 | 1.6 | 1 | 0.804 | 1 | PPM1E |
| Phospholipase activity | 112 | 2.99 | 2 | 0.805 | 1 | PLCL1,PLCL2 |
| Serine_type endopeptidase activity | 160 | 4.28 | 3 | 0.805 | 1 | DPP4,F2,FURIN |
| Secondary active transmembrane transporter activity | 212 | 5.67 | 4 | 0.822 | 1 | SLC8A3,SLC6A9,SLC13A1,CLCN3 |

| | | | | | | |
|---|---|---|---|---|---|---|
| Active transmembrane transporter activity | 347 | 9.27 | 7 | 0.824 | 1 | SLC8A3,SLC6A9,ATP2A2,SLC13A1 |
| Translation initiation factor activity | 64 | 1.71 | 1 | 0.824 | 1 | EIF1AY |
| Acid_amino acid ligase activity | 349 | 9.33 | 7 | 0.828 | 1 | ANAPC4,CUL3,NEDD4L,CBLB,TTL |
| Oxidoreductase activity, acting on a sulfur group of donors | 65 | 1.74 | 1 | 0.829 | 1 | MSRA |
| Single_stranded DNA binding | 65 | 1.74 | 1 | 0.829 | 1 | RPA3 |
| Isomerase activity | 170 | 4.54 | 3 | 0.837 | 1 | FKBP9,SRR,PUS7 |
| Steroid hormone receptor binding | 69 | 1.84 | 1 | 0.846 | 1 | CTNNB1 |
| Carbon_oxygen lyase activity | 69 | 1.84 | 1 | 0.846 | 1 | XRCC6 |
| Small conjugating protein ligase activity | 313 | 8.36 | 6 | 0.846 | 1 | ANAPC4,CUL3,NEDD4L,CBLB,RNF |
| Small conjugating protein ligase activity | 313 | 8.36 | 6 | 0.846 | 1 | ANAPC4,CUL3,NEDD4L,CBLB,RNF |
| Transmembrane receptor protein kinase activity | 125 | 3.34 | 2 | 0.851 | 1 | FGFR1,NTRK3 |

| | | | | | | |
|---|---|---|---|---|---|---|
| Inorganic anion transmembrane transporter activity | 127 | 3.39 | 2 | 0.857 | 1 | SLC13A1,CLCN3 |
| Transcription from RNA polymerase II promoter | 1930 | 51.6 | 45 | 0.858 | 1 | KMT2D,KMT2A,FOXP1,ESR2,ESRI |
| Antioxidant activity | 75 | 2 | 1 | 0.87 | 1 | PXDNL |
| Small conjugating protein binding | 75 | 2 | 1 | 0.87 | 1 | RNF111 |
| G_protein coupled receptor binding | 232 | 6.2 | 4 | 0.871 | 1 | GNAI1,USP4,MAGI2,WNT3 |
| Phosphatase activity | 373 | 9.97 | 7 | 0.875 | 1 | PTPRN2,PPM1E,NT5C2,IMPA2,PTI |
| Exonuclease activity | 77 | 2.06 | 1 | 0.876 | 1 | WRN |
| Chloride channel activity | 77 | 2.06 | 1 | 0.876 | 1 | CLCN3 |
| Symporter activity | 134 | 3.58 | 2 | 0.877 | 1 | SLC6A9,SLC13A1 |
| Lyase activity | 186 | 4.97 | 3 | 0.878 | 1 | GUCY1A2,XRCC6,SRR |
| Lipase activity | 136 | 3.63 | 2 | 0.882 | 1 | PLCL1,PLCL2 |
| Coenzyme binding | 188 | 5.02 | 3 | 0.882 | 1 | TYW1B,DPYD,IVD |
| Extracellular matrix structural constituent | 80 | 2.14 | 1 | 0.886 | 1 | COL11A1 |
| Vitamin binding | 80 | 2.14 | 1 | 0.886 | 1 | FTCDNL1 |
| Peptidase activity | 781 | 20.9 | 16 | 0.894 | 1 | DPP6,ADAM22,THSD4,PSMA5,SPF |
| Purine ribonucleotide binding | 1890 | 50.6 | 43 | 0.894 | 1 | MTHFD1L,MAGI1,MYO18A,DGKI,G |
| Phosphoprotein phosphatase activity | 244 | 6.52 | 4 | 0.895 | 1 | PTPRN2,PPM1E,PTPRF,PTPRO |

| | | | | | | |
|---|---|---|---|---|---|---|
| Lipid transporter activity | 84 | 2.24 | 1 | 0.898 | 1 | RFT1 |
| Amino acid transmembrane transporter activity | 84 | 2.24 | 1 | 0.898 | 1 | SLC6A9 |
| Purine nucleotide binding | 1900 | 50.8 | 43 | 0.898 | 1 | MTHFD1L,MAGI1,MYO18A,DGKI,G |
| Drug binding | 85 | 2.27 | 1 | 0.901 | 1 | FKBP9 |
| Kinase regulator activity | 144 | 3.85 | 2 | 0.901 | 1 | PRKAG2,PIK3R2 |
| Anion channel activity | 87 | 2.33 | 1 | 0.906 | 1 | CLCN3 |
| Hydrolase activity, acting on carbon_nitrogen (but not peptide) bonds | 147 | 3.93 | 2 | 0.907 | 1 | HDAC4,HDAC9 |
| DNA binding | 2760 | 73.7 | 64 | 0.912 | 1 | KDM5D,NPAS3,KMT2D,MYO18A,K[illegible] |
| Guanyl nucleotide binding | 402 | 10.7 | 7 | 0.916 | 1 | GNAI1,EFL1,GUCY1A2,RHOJ,PRK[illegible] |
| Nuclease activity | 206 | 5.51 | 3 | 0.916 | 1 | PXDNL,WRN,SND1 |
| Nuclear hormone receptor binding | 153 | 4.09 | 2 | 0.919 | 1 | CTNNB1,JMJD1C |
| RNA splicing, via transesterification reactions | 321 | 8.58 | 5 | 0.934 | 1 | RSRC1,PRPF3,POLR2E,SRPK2,N[illegible] |
| Ion transmembrane transporter activity | 1000 | 26.7 | 20 | 0.934 | 1 | SLC8A3,GRIN2A,CHRNA5,CACNA[illegible] |
| Cofactor binding | 271 | 7.24 | 4 | 0.934 | 1 | TYW1B,SRR,DPYD,IVD |
| GTP binding | 371 | 9.91 | 6 | 0.935 | 1 | GNAI1,EFL1,GUCY1A2,RHOJ,NKIR |

| | | | | | | |
|---|---|---|---|---|---|---|
| Monovalent inorganic cation transmembrane transporter activity | 377 | 10.1 | 6 | 0.941 | 1 | SLC8A3,KCNG2,SLC6A9,SLC13A1, |
| Growth factor activity | 170 | 4.54 | 2 | 0.944 | 1 | MDK,F2 |
| Carbohydrate binding | 227 | 6.07 | 3 | 0.945 | 1 | PKD1L3,CLEC17A,MANBA |
| Carbohydrate binding | 227 | 6.07 | 3 | 0.945 | 1 | PKD1L3,CLEC17A,MANBA |
| Protein serine/threonine kinase activity | 572 | 15.3 | 10 | 0.945 | 1 | VRK2,PRKD1,MAST3,SBK1,ATP23, |
| Receptor signaling protein activity | 110 | 2.94 | 1 | 0.95 | 1 | TRAIP |
| Hydrolase activity, hydrolyzing O_glycosyl compounds | 110 | 2.94 | 1 | 0.95 | 1 | MANBA |
| Hormone receptor binding | 176 | 4.7 | 2 | 0.951 | 1 | CTNNB1,JMJD1C |
| Endonuclease activity | 117 | 3.13 | 1 | 0.959 | 1 | PXDNL |
| Oxidoreductase activity, acting on NAD(P)H | 117 | 3.13 | 1 | 0.959 | 1 | NDUFA2 |
| Nucleotide binding | 2470 | 66 | 54 | 0.96 | 1 | MTHFD1L,MAGI1,MYO18A,DGKI,G |
| Substrate_specific transmembrane transporter activity | 1100 | 29.3 | 21 | 0.962 | 1 | SLC8A3,GRIN2A,CHRNA5,CACNA2 |

| | | | | | | |
|---|---|---|---|---|---|---|
| Enzyme regulator activity | 1140 | 30.5 | 22 | 0.963 | 1 | DGKI,ARHGAP15,SYNGAP1,ITIH3, |
| Protein kinase regulator activity | 123 | 3.29 | 1 | 0.965 | 1 | PRKAG2 |
| Carboxylic acid transmembrane transporter activity | 127 | 3.39 | 1 | 0.968 | 1 | SLC6A9 |
| Thiolester hydrolase activity | 129 | 3.45 | 1 | 0.97 | 1 | USP4 |
| Organic acid transmembrane transporter activity | 130 | 3.47 | 1 | 0.971 | 1 | SLC6A9 |
| Enzyme inhibitor activity | 373 | 9.97 | 5 | 0.973 | 1 | DGKI,ITIH3,PRKAG2,CD55,FURIN |
| Nucleotidyl transferase activity | 135 | 3.61 | 1 | 0.975 | 1 | POLR2E |
| Hydrolase activity, acting on glycosyl bonds | 137 | 3.66 | 1 | 0.976 | 1 | MANBA |
| Peptidase inhibitor activity | 207 | 5.53 | 2 | 0.976 | 1 | ITIH3,FURIN |
| Monooxygenase activity | 141 | 3.77 | 1 | 0.978 | 1 | AGMO |
| Organic anion transmembrane transporter activity | 147 | 3.93 | 1 | 0.982 | 1 | SLC6A9 |
| Peptide receptor activity | 150 | 4.01 | 1 | 0.983 | 1 | SORCS3 |
| Protein tyrosine kinase activity | 222 | 5.93 | 2 | 0.983 | 1 | FGFR1,NTRK3 |

| | | | | | | |
|---|---|---|---|---|---|---|
| Transmembrane transporter activity | 1240 | 33 | 22 | 0.987 | 1 | SLC8A3,GRIN2A,SLC22A23,[illegible] |
| Oxidoreductase activity | 1010 | 27 | 17 | 0.988 | 1 | KDM5D,MSRA,ENOX1,AGMO,[illegible] |
| Substrate_specific transporter activity | 1240 | 33.2 | 22 | 0.988 | 1 | SLC8A3,GRIN2A,CHRNA5,[illegible] |
| Anion transmembrane transporter activity | 304 | 8.12 | 3 | 0.989 | 1 | SLC6A9,SLC13A1,CLCN3 |
| Structural constituent of ribosome | 167 | 4.46 | 1 | 0.989 | 1 | MRPL33 |
| Receptor binding | 1590 | 42.6 | 29 | 0.992 | 1 | HLA-B,AKAP6,GNAI1,ADAM22,[illegible] |
| Oxidoreductase activity, acting on CH_OH group of donors | 182 | 4.86 | 1 | 0.993 | 1 | GPD2 |
| Hydrolase activity, acting on ester bonds | 1010 | 26.9 | 16 | 0.993 | 1 | PDE1C,PTPRN2,PXDNL,WRN,[illegible] |
| Phosphotransferase activity, alcohol group as acceptor | 1110 | 29.6 | 18 | 0.994 | 1 | DGKI,VRK2,PRKD1,MAST3,SBK1,[illegible] |
| Cysteine_type peptidase activity | 187 | 5 | 1 | 0.994 | 1 | USP4 |
| Kinase activity | 1190 | 31.7 | 19 | 0.996 | 1 | DGKI,VRK2,PRKD1,MAST3,SBK1,[illegible] |
| Protein kinase activity | 960 | 25.7 | 14 | 0.997 | 1 | VRK2,PRKD1,MAST3,SBK1,[illegible] |
| RNA binding | 976 | 26.1 | 14 | 0.997 | 1 | EFL1,RBM6,TNRC6A,THOC7,[illegible] |

| | | | | | | |
|---|---|---|---|---|---|---|
| Transferase activity, transferring phosphorus_containing groups | 1370 | 36.5 | 19 | 1 | 1 | DGKI,VRK2,PRKD1,MAST3,SBK1,C |
| Structural molecule activity | 666 | 17.8 | 6 | 1 | 1 | MRPL33,MAPT,COL11A1,MAGI2,C |
| Receptor activity | 1920 | 51.4 | 30 | 1 | 1 | GPR26,PTPRN2,ANXA9,SEMA3E,H |
| Transmembrane signaling receptor activity | 1510 | 40.4 | 20 | 1 | 1 | GPR26,PTPRN2,ANXA9,HLA-B,GR |
| G_protein coupled receptor activity | 984 | 26.3 | 5 | 1 | 1 | GPR26,GPR141,PKD1L3,GABBR1, |
| Transport | 4830 | 129 | 76 | 1 | 1 | SLC8A3,KMT2A,AKAP6,DPP6,RER |

[illegible]OXP1,SPPL3,WRN,DPP4,EXT1,CITED1,LDB1,FGFR1,IMPA2,ABCB9,SRR,DPYD,BCL11A,PSPH,PTPRO,K[illegible]

,ZFPM2,SND1,CITED1,LDB1,ARID1B,SLC30A9,CTNNB1,MED27,ESRRB,MAML3,BCL11A,ZMIZ2,THRB,ME

T2D,KMT2A,MSL2,DGKI,KMT2E,GNAI1,ADAM22,RERE,CISD2,FOXP1,GRIN2A,CDHR4,AGMO,PXDNL,ESR

D1,PLCL1,SNX29,PCLO,DYSF,SNAP91,NGEF,SYT1,TWF2,PLCL2,CADPS,ZFYVE1,ELMO1,PREX1,TRIOB

,ZFPM2,SND1,CITED1,LDB1,ARID1B,SLC30A9,CTNNB1,MED27,ESRRB,MAML3,BCL11A,ZMIZ2,THRB,ME

:ISD2,FOXP1,SPPL3,WRN,PEF1,DPP4,NPAS1,TCF4,EXT1,CITED1,LDB1,FGFR1,MEF2C,SOX5,IMPA2,AB[illegible]

[illegible]P1,ESRRG,PRKD1,MDK,TCF4,CTR9,RPTOR,HDAC4,ZFPM2,ZNF398,ZMIZ1,CITED1,LDB1,ARID1B,XRC[illegible]

G2,SLC6A9,ATP2A2,CACNA1C,SLC13A1,CACNB2,KCNB1,CACNA1I,KCNN2,SLC39A8,ATP2B2

2E,ADAM22,RERE,FOXP1,GRIN2A,ESR2,ESRRG,ZSCAN9,ZNF536,RBM6,TRIM26,HDAC4,L3MBTL4,TCF2

:2,ESRRG,PTCH1,PRKD1,PLCL1,SNX29,PCLO,DYSF,RASGRP4,SNAP91,NGEF,SYT1,TWF2,ESRRB,PLCL

OXP1,SPPL3,WRN,PRKD1,DPP4,EXT1,CITED1,LDB1,FGFR1,IMPA2,SYT1,ABCB9,SRR,DPYD,BCL11A,PS

G,TCF4,RPTOR,HDAC4,NR1D2,LDB1,XRCC6,CREB5,MEF2C,ESRRB,TSHZ3,FOXO3,GATAD2B,THRB,BCI

2E,ADAM22,RERE,FOXP1,GRIN2A,AGMO,ESR2,ESRRG,WRN,FADS2,ZSCAN9,TYW1B,ZNF536,RBM6,TR

Z,DMXL2,DPP4,SPAG16,ANAPC4,CUL3,CNTNAP2,RPTOR,HDAC4,LDB1,PRKAG2,MAGI2,CTNND1,CTNNI

TCF4,CTR9,HDAC4,ZFPM2,NEDD4L,CITED1,LDB1,FGFR1,XRCC6,MEF2C,SOX5,CTNNB1,TSHZ3,FOXO3

2D2,KCNG2,SLC6A9,PKD1L3,ATP2A2,CHRNA3,CACNA1C,SLC13A1,SLC30A9,CACNB2,KCNB1,CACNA1I,

9B,ARFGEF2,CACNA1C,DBN1,CALD1,FSCN3,ANK3,TWF2,KCNN2,CCDC88C,SYNE1,SORBS1,RHOA,TMC

I1,KLC1,FHIT,MYO9B,ATP2A2,KIF5C,DDX27,MSH5,XRCC6,DDX3Y,RHOJ,ABCB9,RFC2,NKIRAS1,LONRF

I1,KLC1,FHIT,MYO9B,ATP2A2,KIF5C,DDX27,MSH5,XRCC6,DDX3Y,RHOJ,ABCB9,RFC2,NKIRAS1,LONRF

I1,KLC1,MYO9B,ATP2A2,KIF5C,DDX27,MSH5,XRCC6,DDX3Y,RHOJ,ABCB9,RFC2,NKIRAS1,LONRF2,AT

DE1C,ANXA9,KMT2D,MYO18A,KMT2A,MSL2,DGKI,KMT2E,GNAI1,ADAM22,RERE,CISD2,FOXP1,GRIN2A,

RK2,WRN,PRKD1,MAST3,SBK1,MYO9B,PFAS,ATP2A2,KIF5C,DDX27,CKB,GATB,FGFR1,MSH5,PRKAG2,

RK2,WRN,PRKD1,MAST3,SBK1,MYO9B,PFAS,ATP2A2,KIF5C,DDX27,CKB,GATB,FGFR1,MSH5,PRKAG2,[illegible]

RK2,WRN,PRKD1,MAST3,SBK1,MYO9B,PFAS,ATP2A2,KIF5C,DDX27,CKB,GATB,FGFR1,MSH5,PRKAG2,[illegible]

RG,PTCH1,PRKD1,TAF1C,TCF4,CTR9,HDAC4,ZFPM2,NEDD4L,NR1D2,ZMIZ1,CITED1,LDB1,FGFR1,XRCC

NAI1,VRK2,WRN,EFL1,PRKD1,MAST3,SBK1,MYO9B,PFAS,ATP2A2,KIF5C,DDX27,CKB,GATB,FGFR1,MSH

NAI1,VRK2,WRN,EFL1,PRKD1,MAST3,SBK1,MYO9B,PFAS,ATP2A2,KIF5C,DDX27,CKB,GATB,FGFR1,MSH

MT2A,RPA3,RERE,FOXP1,ESR2,ESRRG,WRN,ZSCAN9,ZNF536,TAF1C,NPAS1,TCF4,RBM6,RPTOR,TRIM

2D2,KCNG2,SLC6A9,PKD1L3,ATP2A2,CHRNA3,CACNA1C,SLC13A1,SLC30A9,GRIK1,CACNB2,KCNB1,CA

NAI1,ENOX1,VRK2,WRN,EFL1,PRKD1,TYW1B,MAST3,SBK1,RBM6,TNRC6A,MYO9B,PFAS,ATP2A2,KIF5(

2D2,KCNG2,SLC6A9,PKD1L3,ATP2A2,CHRNA3,CACNA1C,SLC13A1,SLC30A9,GRIK1,CACNB2,KCNB1,AB

MYO9B,DOCK8,MMP16,ARFGEF2,PRKAG2,GMIP,RASGRP4,CD55,NGEF,FURIN,GBF1,PPP2R2B,RGS6,S

A5,CACNA2D2,KCNG2,SLC6A9,PKD1L3,ATP2A2,CHRNA3,CACNA1C,SLC13A1,SLC30A9,GRIK1,CACNB2,

2D2,KCNG2,SLC6A9,PKD1L3,ATP2A2,CHRNA3,CACNA1C,SLC13A1,SLC30A9,GRIK1,CACNB2,KCNB1,AB

R2,PTCH1,MDK,DPP4,F2,USP4,CNTNAP2,SEMA6D,ANKS1B,ARFGEF2,SLC30A9,MAGI2,CTNND1,CTNNB

NT5C2,FGFR1,ATP23,NEK4,PRKG1,PIP5K1B,POLR2E,SRPK2,NTRK3,RPS6KL1,PIK3R2,CAMKV,TLK1

CKB,NT5C2,FGFR1,ATP23,NEK4,PRKG1,PIP5K1B,POLR2E,SRPK2,NTRK3,RPS6KL1,PIK3R2,CAMKV,TLK

:KB,NT5C2,FGFR1,ATP23,NEK4,PRKG1,PIP5K1B,POLR2E,SRPK2,NTRK3,RPS6KL1,PIK3R2,CAMKV,TLK[illegible]

[illegible]LA-B,SEMA3A,GRIN2A,ESR2,ESRRG,GPR141,PTCH1,F2,CHRNA5,PKD1L3,GABBR1,SEMA6D,NR1D2,C[illegible]

IN2A,GPR141,PTCH1,CHRNA5,PKD1L3,GABBR1,CHRNA3,FGFR1,HLA-DQA1,GRIK1,PTPRF,PTPRO,SOR

E,HLA-DMA,GRIN2A,FEZ1,RSRC1,MAPT,WRN,TSNARE1,PTCH1,PRKD1,EXOC4,SLC22A23,F2,PDE8B,C[illegible]

2,ESRRG,WRN,UTY,TET2,FADS2,ZSCAN9,PRKD1,PEF1,TYW1B,PLCL1,F2,PDE8B,ZNF536,CACNA2D2,C

C6,SLC30A9,CREB5,MEF2C,ZKSCAN3,SOX5,CTNNB1,ESRRB,MAML3,BCL11A,FOXO3,RNF111,ZMIZ2,NF [illegible]0,ZFPM2,CBLB,ZNF398,NR1D2,MMP16,ZNF823,ZMIZ1,ZNF804A,ZSCAN12,ZNF638,CREB5,ZSCAN31,ZN[illegible]

[illegible]IM26,HDAC4,L3MBTL4,TCF20,ZFPM2,CBLB,ZNF398,NR1D2,MMP16,ZNF823,ZMIZ1,ZNF804A,ZSCAN12,[illegible]

[illegible]1,FURIN,TWF2,PPP1R16B,TANK,FOXO3,KDM4A,THRB,PREX1,DOCK4,SORBS1,FOXO6,GLI3,TRIOBP,[illegible]

CDHR4,VRK2,AGMO,PXDNL,ESR2,ESRRG,WRN,UTY,TET2,FADS2,EFL1,ZSCAN9,PTCH1,PRKD1,PEF1,[illegible]

[illegible]RCC6,DDX3Y,NEK4,ABCB9,PRKG1,SRR,TWF2,RFC2,PIP5K1B,SRPK2,NTRK3,CLCN3,ATP2B2,RPS6KL1

‹RCC6,DDX3Y,NEK4,ABCB9,PRKG1,SRR,TWF2,RFC2,PIP5K1B,SRPK2,NTRK3,CLCN3,ATP2B2,RPS6KL1

‹RCC6,DDX3Y,NEK4,ABCB9,PRKG1,SRR,TWF2,RFC2,PIP5K1B,SRPK2,NTRK3,CLCN3,ATP2B2,RPS6KL1

C6,SLC30A9,CREB5,MEF2C,SOX5,CTNNB1,MED27,ESRRB,MAML3,BCL11A,FOXO3,LCORL,RNF111,POLI

H5,GUCY1A2,PRKAG2,XRCC6,DDX3Y,RHOJ,NEK4,ABCB9,PRKG1,SRR,TWF2,RFC2,PIP5K1B,NKIRAS1,S

H5,GUCY1A2,PRKAG2,XRCC6,DDX3Y,RHOJ,NEK4,ABCB9,PRKG1,SRR,TWF2,RFC2,PIP5K1B,NKIRAS1,S

M26,HDAC4,TCF20,ZFPM2,ZNF398,NR1D2,ZNF823,DMTF1,CITED1,ZSCAN12,LDB1,ARID1B,ZNF638,MSH

C,DDX27,CKB,GATB,NT5C2,ZNF638,FGFR1,MSH5,GUCY1A2,PRKAG2,XRCC6,SLC30A9,DDX3Y,RHOJ,NE

HRNA3,FGFR1,HLA-DQA1,GRIK1,PTPRF,ESRRB,PTPRO,SEMA3F,SORCS3,HLA-DRB1,NTRK3,HLA-DQB1

HRNA5,CACNA2D2,DYNC1I1,HFE,KCNG2,CUL3,SLC6A9,PKD1L3,PCLO,SPNS1,DYSF,THOC7,NEDD4L,CE

CLSTN2,USP4,MAST3,RBM6,NT5DC2,MYO9B,PCLO,TRIM26,HDAC4,PPM1E,L3MBTL4,TCF20,ZFPM2,BSN

F568,ZNF19,ZSWIM6,ZKSCAN3,PCGF3,ESRRB,BCL11A,ZCCHC7,TSHZ3,KDM4A,ZNF600,RNF111,LONRF

:NF638,CREB5,ZSCAN31,ZNF568,ZNF19,ZSWIM6,ZKSCAN3,PCGF3,ESRRB,BCL11A,ZCCHC7,TSHZ3,KD

MDK,TYW1B,PLCL1,F2,PDE8B,ZNF536,CACNA2D2,CLSTN2,USP4,MAST3,SBK1,RBM6,NT5DC2,MYO9B,P

R2E,ZMIZ2,TENM2,NFIX,THRB,BRWD1,TRIM33,BCL11B,FOXO6,CSRNP3,MED8,GLI3,RHOA,HDAC9

5,XRCC6,CREB5,ZSCAN31,MEF2C,ZNF568,ZNF19,DDX3Y,TFB1M,ZKSCAN3,PALB2,SOX5,CTNNB1,ESRF [illegible]K4,ABCB9,PRKG1,SRR,TWF2,DHRS11,DPYD,CPEB1,IVD,RFC2,PIP5K1B,NKIRAS1,SRPK2,NTRK3,CLCN

3LB,CALU,ATP2A2,KIF5C,ARFGEF2,CITED1,CHRNA3,CACNA1C,SLC13A1,PRKAG2,SLC30A9,GRIK1,MAG

I,COL11A1,CBLB,GPD2,ZNF398,NR1D2,MMP16,ZNF823,ITGA11,CALU,ATP2A2,ZMIZ1,ZNF804A,CHRNA3,

F2,ZFYVE1,ZMIZ2,GATAD2B,ZNF615,THRB,TRAIP,TRIM33,BCL11B,GLI3,ZKSCAN8,NEURL1,RNF144A,ZN

M4A,ZNF600,RNF111,LONRF2,ZFYVE1,ZMIZ2,GALNT1,GATAD2B,ZNF615,THRB,TRAIP,TRIM33,BCL11B,

[illegible]CLO,TRIM26,HDAC4,PPM1E,L3MBTL4,TCF20,ZFPM2,BSN,COL11A1,PFAS,CBLB,GPD2,ZNF398,NR1D2,[illegible]

RB,TSHZ3,FOXO3,LCORL,RFC2,ZNF600,RBL2,POLR2E,NFIX,GATAD2B,ZNF615,THRB,TRIM33,BCL11B,F

GI2,SNAP91,CTNNB1,SYT1,CACNB2,KCNB1,ANK3,ABCB9,FURIN,CACNA1I,CNNM2,CADPS,POM121C,SL[illegible]

,CACNA1C,ZSCAN12,NT5C2,ZNF638,CREB5,ZSCAN31,ATP23,ZNF568,ZNF19,GMIP,MGAT5B,ZSWIM6,ZK

MMP16,ZNF823,ITGA11,CALU,ATP2A2,ZMIZ1,ZNF804A,KIF5C,DDX27,CKB,CHRNA3,CACNA1C,ZSCAN12,

C35F2,RFT1,GBF1,HLA-DRB5,KCNN2,BTN3A2,PPP2R2B,HLA-DRB1,ELMO1,IMMP2L,CLCN3,SLC39A8,AT

[illegible]SCAN3,RASGRP4,IMPA2,NEK4,SYT1,PCGF3,FURIN,SRR,ESRRB,PLCL2,DPYD,TANK,CADPS,BCL11A,Z[illegible]

GATB,NT5C2,ZNF638,FGFR1,MSH5,GUCY1A2,FTCDNL1,PRKAG2,XRCC6,CREB5,ZSCAN31,ATP23,ZNF5

CCHC7,TSHZ3,PSPH,CLEC17A,KDM4A,JMJD1C,ZNF600,KEL,PDE4B,RNF111,LONRF2,ZFYVE1,ZMIZ2,M[illegible]

568,ZNF19,GMIP,DDX3Y,MGAT5B,RHOJ,ZSWIM6,ZKSCAN3,RASGRP4,SNAP91,IMPA2,NEK4,SYT1,PCGF

ANBA,SRPK2,GALNT1,GALNT10,GATAD2B,ZNF615,PCDH9,THRB,PPA2,ATP2B2,TRAIP,KDM3B,TRIM33,B

3,PTPRF,ABCB9,FURIN,PRKG1,SRR,TWF2,ESRRB,PLCL2,DPYD,TANK,IVD,CADPS,BCL11A,ZCCHC7,TS

HZ3,PSPH,CLEC17A,KDM4A,JMJD1C,RFC2,ZNF600,PIP5K1B,KEL,PDE4B,NKIRAS1,RNF111,LONRF2,ZF

YVE1,ZMIZ2,MANBA,SRPK2,NTRK3,GALNT1,TNXB,GALNT10,GATAD2B,ZNF615,PCDH9,THRB,CLCN3,PF

PA2,ATP2B2,RPS6KL1,KIF21B,TRAIP,KDM3B,TRIM33,BCL11B,GLI3,ZKSCAN8,NEURL1,RHOA,DNAH11,HI

[illegible]DAC9,RNF144A,CAMKV,ZNF800,CDKAL1,TLK1,SREK1IP1
