## Supplemental table 4 for "Digestive Dimensions of Autism: A Multiscale Exploration of Gut-Brain Interactions"

| Pathway | Total | Expected | Hits | P.Value | FDR | Genes |
|---|---|---|---|---|---|---|
| Antigen processing and presentation of peptide or polysaccharide antigen via MHC class II | 16 | 0.509 | 5 | 0.000104 | 0.0202 | HLA-DMA,HLA-DQA1,HLA-DRB5,HI |
| Antigen processing and presentation | 43 | 1.37 | 6 | 0.00225 | 0.218 | HLA-B,HLA-DMA,HLA-DQA1,HLA-D |
| Anterior/posterior axis specification | 16 | 0.509 | 3 | 0.0132 | 0.851 | LDB1,CTNNB1,WNT3 |
| Dorsal/ventral axis specification | 9 | 0.287 | 2 | 0.0314 | 1 | CTNNB1, WNT3 |
| Nervous system development | 516 | 16.4 | 24 | 0.0405 | 1 | NRGN,SEMA3E,SEMA3A,FEZ1,PRI |
| Chromatin organization | 358 | 11.4 | 17 | 0.0656 | 1 | KDM5D,KMT2D,KMT2A,KMT2E,UT |
| Synaptic vesicle exocytosis | 30 | 0.955 | 3 | 0.069 | 1 | PCLO,SYT1,CADPS |
| Natural killer cell activation | 31 | 0.987 | 3 | 0.0746 | 1 | HFE,RAET1E,MICA |
| Organelle organization | 15 | 0.478 | 2 | 0.0808 | 1 | ATP2A2,KIF5C |
| RRNA metabolic process | 3 | 0.0955 | 1 | 0.0925 | 1 | MAPT |
| Cell adhesion | 629 | 20 | 26 | 0.104 | 1 | MAGI1,ADAM22,FEZ1,CDHR4,DPP |
| Neurotransmitter secretion | 38 | 1.21 | 3 | 0.119 | 1 | PTPRN2,DGKI,SYT1 |

| | | | | | | |
|---|---|---|---|---|---|---|
| Phosphate_containing compound metabolic process | 20 | 0.637 | 2 | 0.132 | 1 | IMPA2,PPA2 |
| Transcription, DNA_templated | 217 | 6.91 | 10 | 0.155 | 1 | ESR2,ESRRG,ZFPM2,NR1D2,LDB1 |
| Pattern specification process | 67 | 2.13 | 4 | 0.164 | 1 | SYNGAP1,PTCH1,RNF111,GLI3 |
| Neutrophil mediated immunity | 8 | 0.255 | 1 | 0.228 | 1 | KMT2E |
| Locomotion | 8 | 0.255 | 1 | 0.228 | 1 | GRIN2A |
| Neuron_neuron synaptic transmission | 9 | 0.287 | 1 | 0.253 | 1 | ARID1B |
| Neuronal action potential propagation | 9 | 0.287 | 1 | 0.253 | 1 | NTRK3 |
| Protein metabolic process | 9 | 0.287 | 1 | 0.253 | 1 | MMP16 |
| Neuromuscular synaptic transmission | 31 | 0.987 | 2 | 0.259 | 1 | CHRNA5,CHRNA3 |
| Macrophage activation | 10 | 0.318 | 1 | 0.276 | 1 | FOXP1 |
| Chemical synaptic transmission | 254 | 8.09 | 10 | 0.291 | 1 | GRIN2A,EXOC4,CHRNA5,BSN,CHI |
| DNA metabolic process | 34 | 1.08 | 2 | 0.295 | 1 | MYO18A, WRN |
| Mammary gland development | 35 | 1.11 | 2 | 0.307 | 1 | PTCH1,GLI3 |
| Cell_matrix adhesion | 91 | 2.9 | 4 | 0.329 | 1 | ITGA11,CTNNB1,SORBS1,RHOA |

| | | | | | | |
|---|---|---|---|---|---|---|
| Mesoderm development | 37 | 1.18 | 2 | 0.33 | 1 | EXT1,PALB2 |
| Heart development | 240 | 7.64 | 9 | 0.356 | 1 | FOXP1,ZFPM2,CACNA1C,MEF2C,[illegible] |
| Unsaturated fatty acid biosynthetic process | 14 | 0.446 | 1 | 0.364 | 1 | FADS2 |
| Ectoderm development | 14 | 0.446 | 1 | 0.364 | 1 | CTNNB1 |
| Cellular process | 14 | 0.446 | 1 | 0.364 | 1 | CTNNB1 |
| Regulation of gene expression, epigenetic | 16 | 0.509 | 1 | 0.404 | 1 | HDAC4 |
| Cellular calcium ion homeostasis | 102 | 3.25 | 4 | 0.409 | 1 | SLC8A3,ATP2A2,KEL,ATP2B2 |
| Respiratory electron transport chain | 17 | 0.541 | 1 | 0.423 | 1 | IMMP2L |
| DNA recombination | 104 | 3.31 | 4 | 0.423 | 1 | RPA3,WRN,XRCC6,PALB2 |
| Cellular glucose homeostasis | 18 | 0.573 | 1 | 0.442 | 1 | PIK3R2 |
| Regulation of translation | 139 | 4.42 | 5 | 0.455 | 1 | TNRC6A,CNOT1,CPEB1,FOXO3,N[illegible] |
| Viral process | 448 | 14.3 | 15 | 0.458 | 1 | HLA-B,VRK2,KLC1,NEDD4L,SND1,[illegible] |
| Sensory perception of sound | 140 | 4.46 | 5 | 0.462 | 1 | COL11A1,FGFR1,THRB,ATP2B2,T[illegible] |
| Protein glycosylation | 171 | 5.44 | 6 | 0.463 | 1 | GALNT12,EXT1,MGAT5B,ST3GAL[illegible] |
| Muscle contraction | 111 | 3.53 | 4 | 0.473 | 1 | DYSF,CALD1,SORBS1,TMOD3 |
| Ion transport | 671 | 21.4 | 22 | 0.475 | 1 | SLC8A3,GRIN2A,SLC22A23,CHRN[illegible] |

| | | | | | | |
|---|---|---|---|---|---|---|
| Gamete generation | 22 | 0.7 | 1 | 0.51 | 1 | WNT3 |
| Nervous system process | 56 | 1.78 | 2 | 0.536 | 1 | CHRNA5,CHRNA3 |
| Response to interferon_ gamma | 24 | 0.764 | 1 | 0.54 | 1 | CITED1 |
| Exocytosis | 122 | 3.88 | 4 | 0.547 | 1 | EXOC4,ARFGEF2,KCNB1,CADPS |
| RNA splicing, via transesterif ication reactions | 25 | 0.796 | 1 | 0.555 | 1 | PRPF3 |
| Chromatin remodeling | 91 | 2.9 | 3 | 0.557 | 1 | RERE,HDAC4,ARID1B |
| Metabolic process | 191 | 6.08 | 6 | 0.571 | 1 | WRN,NT5C2,XRCC6,PRKG1,SRR,I |
| RNA catabolic process | 28 | 0.891 | 1 | 0.596 | 1 | SND1 |
| Endoderm developme nt | 30 | 0.955 | 1 | 0.622 | 1 | EXT1 |
| Muscle organ developme nt | 104 | 3.31 | 3 | 0.648 | 1 | FXR1,ITGA11,MEF2C |
| Fatty acid biosyntheti c process | 69 | 2.2 | 2 | 0.65 | 1 | FADS2,PRKAG2 |
| 7_methylg uanosine mRNA capping | 34 | 1.08 | 1 | 0.668 | 1 | POLR2E |
| Protein methylatio n | 35 | 1.11 | 1 | 0.678 | 1 | ETF1 |
| Mitochondr ial translation | 35 | 1.11 | 1 | 0.678 | 1 | GATB |
| Cellular protein modificatio n process | 110 | 3.5 | 3 | 0.685 | 1 | MSRA,TTLL7,MANBA |
| Skeletal system developme nt | 147 | 4.68 | 4 | 0.693 | 1 | HDAC4,EXT1,FGFR1,CTNNB1 |

| | | | | | | |
|---|---|---|---|---|---|---|
| Glycogen metabolic process | 38 | 1.21 | 1 | 0.708 | 1 | PRKAG2 |
| Blood circulation | 38 | 1.21 | 1 | 0.708 | 1 | IMMP2L |
| Carbohydrate transport | 39 | 1.24 | 1 | 0.717 | 1 | RFT1 |
| DNA replication | 155 | 4.93 | 4 | 0.732 | 1 | RPA3,WRN,RFC2,NFIX |
| Cation transport | 81 | 2.58 | 2 | 0.734 | 1 | PKD1L3,SLC30A9 |
| Cell differentiation | 971 | 30.9 | 28 | 0.737 | 1 | SEMA3E,SEMA3A,PRKD1,MDK,TC |
| Blood coagulation | 193 | 6.14 | 5 | 0.74 | 1 | F2,DOCK8,ZFPM2,CARMIL1,JMJD[illegible] |
| Gluconeogenesis | 44 | 1.4 | 1 | 0.76 | 1 | GPD2 |
| Protein targeting | 44 | 1.4 | 1 | 0.76 | 1 | AKAP6 |
| Rhythmic process | 124 | 3.95 | 3 | 0.76 | 1 | KMT2A,ENOX1,NR1D2 |
| Immune system process | 515 | 16.4 | 14 | 0.766 | 1 | HLA-B,HLA-DMA,PRKD1,TRIM26,R |
| Cellular amino acid biosynthetic process | 45 | 1.43 | 1 | 0.767 | 1 | PSPH |
| Cellular amino acid metabolic process | 47 | 1.5 | 1 | 0.782 | 1 | SRR |
| Cell communication | 47 | 1.5 | 1 | 0.782 | 1 | SLC8A3 |
| RNA metabolic process | 47 | 1.5 | 1 | 0.782 | 1 | POLR2E |
| Immune response | 387 | 12.3 | 10 | 0.793 | 1 | HLA-B,HLA-DMA,HFE,RAET1E,HL[illegible] |
| Cytoskeleton organization | 170 | 5.41 | 4 | 0.795 | 1 | MAST3,PCLO,BRWD1,RHOA |
| Amino acid transport | 52 | 1.66 | 1 | 0.815 | 1 | SLC6A9 |
| RNA splicing | 289 | 9.2 | 7 | 0.818 | 1 | RSRC1,THOC7,ZNF638,PRPF3,SR |

| | | | | | | |
|---|---|---|---|---|---|---|
| Nucleobase_containing compound metabolic process | 55 | 1.75 | 1 | 0.832 | 1 | WRN |
| Sensory perception of pain | 56 | 1.78 | 1 | 0.837 | 1 | GRIN2A |
| MRNA 3'_end processing | 57 | 1.81 | 1 | 0.843 | 1 | THOC7 |
| Phagocytosis | 65 | 2.07 | 1 | 0.879 | 1 | ELMO1 |
| Steroid metabolic process | 121 | 3.85 | 2 | 0.902 | 1 | ERLIN1,DHRS11 |
| Transcription by RNA polymerase II | 626 | 19.9 | 15 | 0.902 | 1 | KMT2A,TAF1C,CTR9,LDB1,CREB5 |
| Protein phosphorylation | 627 | 20 | 15 | 0.903 | 1 | VRK2,RSRC1,PRKD1,MAST3,SBK1 |
| Angiogenesis | 252 | 8.02 | 5 | 0.908 | 1 | SEMA3E,PRKD1,DYSF,FGFR1,SR[illegible] |
| Hemopoiesis | 74 | 2.36 | 1 | 0.909 | 1 | CTNNB1 |
| Protein localization | 76 | 2.42 | 1 | 0.915 | 1 | PARD3B |
| Regulation of transcription by RNA polymerase II | 1250 | 39.7 | 32 | 0.922 | 1 | KDM5D,NPAS3,RERE,ZSCAN9,NP[illegible] |
| Proteolysis | 646 | 20.6 | 15 | 0.924 | 1 | DPP6,ADAM22,THSD4,PSMA5,SP[illegible] |
| Regulation of catalytic activity | 80 | 2.55 | 1 | 0.925 | 1 | PRKAG2 |
| Chromosome segregation | 83 | 2.64 | 1 | 0.932 | 1 | STAG1 |
| Lipid transport | 136 | 4.33 | 2 | 0.934 | 1 | SPNS1,RFT1 |
| Translation | 315 | 10 | 6 | 0.939 | 1 | MRPL33,EFL1,GATB,EIF1AY,ETF1[illegible] |

| | | | | | | |
|---|---|---|---|---|---|---|
| Cholesterol metabolic process | 89 | 2.83 | 1 | 0.944 | 1 | ERLIN1 |
| MRNA splicing, via spliceosome | 236 | 7.51 | 4 | 0.946 | 1 | RSRC1,PRPF3,POLR2E,NOVA1 |
| Mitochondrion organization | 90 | 2.87 | 1 | 0.946 | 1 | TFB1M |
| Circadian rhythm | 90 | 2.87 | 1 | 0.946 | 1 | NTRK3 |
| Regulation of cell cycle | 148 | 4.71 | 2 | 0.952 | 1 | FBXL18,RBL2 |
| Protein folding | 157 | 5 | 2 | 0.962 | 1 | FKBP9,GNAI1 |
| Visual perception | 208 | 6.62 | 3 | 0.964 | 1 | BBS9,COL11A1,CACNB2 |
| Cell proliferation | 386 | 12.3 | 7 | 0.965 | 1 | PRKD1,MRM2,CITED1,RASGRP4,[illegible] |
| Fatty acid metabolic process | 163 | 5.19 | 2 | 0.968 | 1 | FADS2,PRKAG2 |
| Cell cycle | 647 | 20.6 | 13 | 0.975 | 1 | KMT2E,GNAI1,TET2,ANAPC4,DMT[illegible] |
| Vesicle_mediated transport | 274 | 8.72 | 4 | 0.977 | 1 | TSNARE1,CUL3,ARFGEF2,SYT1 |
| MRNA processing | 370 | 11.8 | 6 | 0.98 | 1 | RSRC1,THOC7,PRPF3,CPEB1,SR[illegible] |
| Apoptotic process | 699 | 22.3 | 14 | 0.98 | 1 | KMT2A,SEMA3A,PRKD1,FHIT,FXR[illegible] |
| Cell_cell signaling | 232 | 7.39 | 3 | 0.98 | 1 | ESR2,MAPT,PKP4 |
| DNA repair | 379 | 12.1 | 6 | 0.983 | 1 | RPA3,WRN,MSH5,XRCC6,PALB2,[illegible] |
| Defense response to bacterium | 164 | 5.22 | 1 | 0.995 | 1 | MICA |
| Lipid metabolic process | 538 | 17.1 | 8 | 0.996 | 1 | PTPRN2,ERLIN1,FADS2,PLCL1,PR[illegible] |
| Intracellular protein transport | 353 | 11.2 | 4 | 0.997 | 1 | TSNARE1,KLC1,SGSM2,TLK1 |
| Endocytosis | 184 | 5.86 | 1 | 0.998 | 1 | RHOJ |
| Protein transport | 654 | 20.8 | 9 | 0.999 | 1 | EXOC4,BBS9,ARFGEF2,SNAP91,A[illegible] |

| | | | | | | |
|---|---|---|---|---|---|---|
| Carbohydrate metabolic process | 228 | 7.26 | 1 | 0.999 | 1 | MANBA |
| Negative regulation of apoptotic process | 577 | 18.4 | 7 | 0.999 | 1 | MYO18A,FGFR1,PALB2,CTNNB1,N[illegible] |
| Biological_process | 597 | 19 | 7 | 1 | 1 | TMX2,CALU,AUTS2,MYBPHL,SLC3[illegible] |
| Spermatogenesis | 457 | 14.5 | 3 | 1 | 1 | USP4,SYNE1,IMMP2L |
| Sensory perception of smell | 416 | 13.2 | 1 | 1 | 1 | PDE1C |
| Response to stimulus | 566 | 18 | 1 | 1 | 1 | BBS9 |

KD1,MDK,TCF4,HDAC4,CNTN4,SEMA6D,CHRNA3,TTLL7,ARID1B,AMIGO1,MEF2C,GRIK1,DBN1,MAGI2,C

Y,TET2,SGF29,HDAC4,L3MBTL4,ARID1B,KDM4A,JMJD1C,RBL2,BRWD1,KDM3B,HDAC9,TLK1

4,PKP4,CLSTN2,CNTNAP2,CNTN4,OLFM4,ITGA11,ATP2A2,AMIGO1,SDK1,CADM2,CTNND1,CTNNB1,NT

A5,CACNA2D2,HFE,KCNG2,PKD1L3,ATP2A2,CHRNA3,CACNA1C,SLC13A1,SLC30A9,GRIK1,CACNB2,KC[illegible]

F4,ZFPM2,NEDD4L,SEMA6D,NR1D2,FXR1,CITED1,TTLL7,FGFR1,AMIGO1,MEF2C,DBN1,RASGRP4,CTN[illegible]

AS1,CUL3,ZFPM2,ZNF398,ZNF823,DMTF1,CITED1,ZSCAN12,LDB1,ARID1B,CREB5,ZSCAN31,ZNF19,CTN

INB1,MED27,TSHZ3,FOXO3,LCORL,ZNF600,RBL2,ZNF615,THRB,BRWD1,FOXO6,MED8,ZKSCAN8,RHOA

.,ZNF800
