## Supplemental table 5 for "Digestive Dimensions of Autism: A Multiscale Exploration of Gut-Brain Interactions"

| Pathway | Total | Expected | Hits | P.Value | FDR | Genes |
|---|---|---|---|---|---|---|
| Cell projection | 943 | 20.2 | 47 | 6.19E-08 | 3.28E-06 | SLC8A3,MAGI1,ADAM22,FEZ1,MAI |
| Synapse | 548 | 11.7 | 31 | 9.43E-07 | 2.50E-05 | NRGN,PTPRN2,DGKI,GRIN2A,DM) |
| Dendrite | 453 | 9.7 | 26 | 5.61E-06 | 9.91E-05 | NRGN,SLC8A3,SEMA3A,FEZ1,MAI |
| Postsynaptic membrane | 245 | 5.25 | 17 | 2.29E-05 | 0.000303 | NRGN,GRIN2A,CHRNA5,CLSTN2,( |
| Cell junction | 820 | 17.6 | 36 | 3.84E-05 | 0.000407 | SLC8A3,MAGI1,PTPRN2,GRIN2A,[ |
| Axon | 374 | 8.01 | 20 | 0.000176 | 0.00156 | NRGN,ADAM22,SEMA3A,FEZ1,MA |
| Neuron projection | 411 | 8.8 | 20 | 0.000594 | 0.0045 | GRIN2A,MAPT,WRN,CHRNA5,KLC |
| Neuronal cell body | 403 | 8.63 | 17 | 0.00639 | 0.0414 | NRGN,SLC8A3,DGKI,FEZ1,MAPT,k |
| Golgi apparatus | 1290 | 27.6 | 41 | 0.00704 | 0.0414 | GALNT12,MYO18A,HLA-B,GIGYF2, |
| Plasma membrane | 4750 | 102 | 121 | 0.0152 | 0.0803 | SLC8A3,MAGI1,GPR26,PTPRN2,HI |
| Tubulin complex | 2 | 0.0428 | 1 | 0.0424 | 0.196 | MAPT |
| Protein_containing complex | 656 | 14.1 | 21 | 0.0444 | 0.196 | ERLIN1,DGKI,KMT2E,GNAI1,CISD2 |
| Cytoskeleton | 1310 | 28 | 36 | 0.0721 | 0.294 | MYO18A,KMT2E,GNAI1,FEZ1,MAP |
| Presynaptic membrane | 78 | 1.67 | 4 | 0.0863 | 0.327 | GRIN2A,GABBR1,GRIK1,SYT1 |
| Actin cytoskeleton | 224 | 4.8 | 8 | 0.11 | 0.387 | MSRA,NCOA5,MYO9B,AUTS2,DBN |
| Bicellular tight junction | 123 | 2.63 | 5 | 0.125 | 0.413 | MAGI1,MAGI2,CTNNB1,ANK3,PARI |
| Neuromuscular junction | 66 | 1.41 | 3 | 0.168 | 0.524 | SLC8A3,HDAC4,ANK3 |
| Microtubule | 349 | 7.48 | 10 | 0.217 | 0.64 | SLC8A3,FEZ1,MAPT,DYNC1I1,KLC |
| Protein_DNA complex | 44 | 0.943 | 2 | 0.243 | 0.643 | XRCC6,CTNNB1 |
| Basal part of cell | 15 | 0.321 | 1 | 0.277 | 0.643 | HFE |
| Cytoplasm | 6540 | 140 | 146 | 0.278 | 0.643 | NRGN,MTHFD1L,NREP,SLC8A3,M, |
| Apical part of cell | 89 | 1.91 | 3 | 0.298 | 0.643 | HFE,CTNNB1,SRR |
| Lysosome | 380 | 8.14 | 10 | 0.299 | 0.643 | HLA-DMA,USP4,RPTOR,HLA-DQA' |
| SNARE complex | 51 | 1.09 | 2 | 0.299 | 0.643 | TSNARE1,SYT1 |
| Nucleus | 6480 | 139 | 144 | 0.307 | 0.643 | FBXL18,NRGN,KDM5D,NREP,MAG |
| Intracellular | 1540 | 33 | 36 | 0.315 | 0.643 | GPR26,GNAI1,MRPL33,SYNGAP1,I |

| | | | | | | |
|---|---|---|---|---|---|---|
| Cilium | 354 | 7.58 | 9 | 0.348 | 0.684 | PDE1C,PTCH1,BBS9,SPAG16,CUL |
| Heterotrimeric G_protein complex | 32 | 0.685 | 1 | 0.5 | 0.914 | GNAI1 |
| Gap junction | 32 | 0.685 | 1 | 0.5 | 0.914 | DBN1 |
| Membrane | 8150 | 175 | 173 | 0.587 | 1 | GRAMD1B,MTHFD1L,SLC8A3,MAG |
| Endosome | 644 | 13.8 | 13 | 0.626 | 1 | HLA-DMA,GIGYF2,DYSF,NEDD4L,/ |
| Cellular_component | 456 | 9.77 | 9 | 0.645 | 1 | KDM5D,TMX2,PDE8B,TRIM26,AUT |
| Chromosome | 463 | 9.92 | 9 | 0.663 | 1 | KMT2E,WRN,DYNC1I1,DDX27,XR( |
| Mitochondrion | 1510 | 32.3 | 30 | 0.692 | 1 | MTHFD1L,SLC8A3,MSRA,MRPL33, |
| Endoplasmic reticulum | 1480 | 31.7 | 29 | 0.722 | 1 | FKBP9,SLC8A3,ERLIN1,HLA-B,CIS |
| Clathrin_coated pit | 61 | 1.31 | 1 | 0.734 | 1 | SNAP91 |
| Microvillus | 64 | 1.37 | 1 | 0.751 | 1 | EXOC4 |
| Extracellular matrix | 300 | 6.43 | 5 | 0.773 | 1 | THSD4,COL11A1,MMP16,TNXB,WI |
| Cytosol | 5030 | 108 | 101 | 0.794 | 1 | FBXL18,PDE1C,NPAS3,ANXA9,KM |
| Mitochondrial inner membrane | 430 | 9.21 | 6 | 0.902 | 1 | MRPL33,SPNS1,GPD2,IMMP2L,ND |
| Integral component of membrane | 5470 | 117 | 105 | 0.924 | 1 | GRAMD1B,SLC8A3,GPR26,PTPRN |
| Peroxisome | 119 | 2.55 | 1 | 0.925 | 1 | GBF1 |
| Ribosome | 201 | 4.31 | 2 | 0.932 | 1 | MRPL33,RPS6KL1 |
| Extracellular region | 2280 | 48.9 | 33 | 0.996 | 1 | SEMA3E,SEMA3A,THSD4,ENOX1,I |
| Extracellular space | 1680 | 35.9 | 19 | 1 | 1 | SEMA3E,SEMA3A,ENOX1,PXDNL,I |

PT,KIZ,EXOC4,MDK,DPP4,BBS9,SPAG16,KLC1,CUL3,DOCK8,CNTNAP2,GABBR1,ANKS1B,KIF5C,ARFGEI

KL2,EXOC4,CHRNA5,PCLO,GABBR1,BSN,ANKS1B,KIF5C,ARFGEF2,CHRNA3,CACNA1C,SDK1,GRIK1,MA

PT,CNTNAP2,RPTOR,GABBR1,BSN,FXR1,KIF5C,ARFGEF2,CKB,CHRNA3,CACNA1C,TTLL7,AMIGO1,DBN

GABBR1,ANKS1B,CHRNA3,CACNA1C,GRIK1,KCNB1,ANK3,CPEB1,SHISA9,TENM2,SYNE1,IGSF9B,NEURI

MXL2,DPP4,PKP4,CHRNA5,CNTNAP2,PCLO,GABBR1,BSN,ANKS1B,ARFGEF2,CHRNA3,CACNA1C,SDK1

PT,KLC1,CNTNAP2,BSN,CNTN4,FXR1,ZNF804A,AMIGO1,CADM2,SYT1,KCNB1,ANK3,PTPRO,NTRK3,KIF

1,BSN,KIF5C,CHRNA3,MAGI2,SYT1,KCNB1,ANK3,PTPRF,CPEB1,PSPH,PTPRO,KCNN2,TENM2,TENM4,B

,FEZ1,SPPL3,PTCH1,PRKD1,CLSTN2,TNRC6A,CUL3,CNTNAP2,MPHOSPH9,DYSF,NEDD4L,SEMA6D,EXT

LA-B,DGKI,KMT2E,DPP6,GNAI1,ADAM22,SYNGAP1,ENOX1,GRIN2A,FEZ1,CDHR4,SPPL3,MAPT,GPR141,

2,GIGYF2,VRK2,HDAC4,PPM1E,ATP2A2,LDB1,PRPF3,XRCC6,MEF2C,MAGI2,CTNNB1,TANK,FOXO3,RNF

T,KIZ,PKP4,BBS9,SPAG16,DYNC1I1,KLC1,MYO9B,PCLO,BSN,MPHOSPH9,KIF5C,ARFGEF2,TTLL7,AUTS

AGI1,PTPRN2,MYO18A,MSRA,DGKI,AKAP6,KMT2E,ARHGAP15,GNAI1,FOXP1,GIGYF2,FEZ1,PSMA5,VRK

I1,NPAS3,KMT2D,KMT2A,MSRA,DGKI,RPA3,AKAP6,KMT2E,GNAI1,RERE,FOXP1,PSMA5,VRK2,RSRC1,N

MAPT,WRN,PRKD1,PLCL1,MAST3,MYO9B,DOCK8,TRIM26,NEDD4L,ZNF398,ZNF823,ZNF568,ZNF19,GMI

;I1,GPR26,PTPRN2,GALNT12,MYO18A,ERLIN1,HLA-B,MSRA,AKAP6,KMT2E,DPP6,ARHGAP15,GNAI1,AD/

,CISD2,FEZ1,VRK2,ESR2,MAPT,TCAIM,MRM2,CYP2D7,FHIT,PPM1E,SPNS1,OLFM4,GPD2,GATB,TFB1M,I

D2,RTN1,GRIN2A,GIGYF2,VRK2,AGMO,SPPL3,FADS2,PEF1,CLSTN2,EXT1,CALU,ATP2A2,HLA-DQA1,SL(

T2A,MSRA,DGKI,ARHGAP15,SYNGAP1,GIGYF2,PSMA5,RSU1,MAPT,PRKD1,EXOC4,BBS9,PDE8B,ANAP(

2,GALNT12,ERLIN1,HLA-B,DPP6,ADAM22,CISD2,RTN1,HLA-DMA,GRIN2A,GIGYF2,CDHR4,VRK2,AGMO,S

PSMA5,PXDNL,ESR2,MAPT,MDK,DPP4,F2,ITIH3,GABBR1,CNTN4,COL11A1,OLFM4,MMP16,CALU,SFTA2,

NCOA5,DMXL2,F2,HFE,COL11A1,OLFM4,SEMA6D,CKB,CNOT1,PRKAG2,MICA,SEMA3F,HLA-DRB1,TNXB

F2,CACNA1C,TTLL7,AUTS2,AMIGO1,CARMIL1,RHOJ,DBN1,CADM2,CTNNB1,NEK4,NGEF,KCNB1,ANK3,D

GI2,CADM2,CTNND1,CTNNB1,SYT1,KCNB1,ANK3,CPEB1,CADPS,SHISA9,TENM2,ATP2B2,IGSF9B,NEUR

l,ATP23,GRIK1,DBN1,MAGI2,CADM2,CTNNB1,SYT1,KCNB1,ANK3,CPEB1,CADPS,SHISA9,PARD3B,TENM

T1,CALU,ARFGEF2,SFTA2,HLA-DQA1,GLT8D1,MGAT5B,ST3GAL3,SYT1,ANK3,FURIN,PRKG1,GBF1,HLA-I

PTCH1,PRKD1,EXOC4,TMEM170B,PLCL1,DPP4,PKP4,BBS9,F2,CHRNA5,CACNA2D2,CLSTN2,HFE,KCNG

2,SLC30A9,CARMIL1,MAD1L1,DBN1,CALD1,FSCN3,CTNNB1,ANK3,TWF2,PPP2R2B,SYNE1,KIF21B,SORE

2,PXDNL,RSRC1,MAPT,WRN,KIZ,SSUH2,PRKD1,PEF1,EXOC4,MDK,PLCL1,PKP4,BBS9,CYP2D7,SPAG16

COA5,ESR2,MAPT,ESRRG,WRN,UTY,TET2,SSUH2,ZSCAN9,PTCH1,PRKD1,PKP4,ZNF536,TAF1C,NPAS1

P,RHOJ,DBN1,ZKSCAN3,RASGRP4,CTNNB1,ADTRP,TWF2,PLCL2,JMJD1C,RGS6,NFIX,TNXB,ZNF615,DC

AM22,CISD2,SYNGAP1,RTN1,HLA-DMA,ENOX1,GRIN2A,GIGYF2,FEZ1,CDHR4,VRK2,AGMO,SPPL3,MAP[illegible]

C30A9,ANK3,FURIN,DDN,POM121C,HLA-DRB5,ZFYVE1,HLA-DRB1,TENM2,HLA-DQB1,CDKAL1

C4,DYNC1I1,USP4,KLC1,FHIT,TNRC6A,MYO9B,CUL3,DOCK8,RPTOR,TRIM26,HDAC4,PFAS,THOC7,NED[illegible]

SPPL3,GPR141,FADS2,TSNARE1,KIZ,TMX2,PTCH1,TYW1B,TMEM170B,DPP4,CYP2D7,SLC22A23,CHRN[illegible]

,RAET1E,FGFR1,XRCC6,CD55,ST3GAL3,NTM,FURIN,LUZP2,DHRS11,SEMA3F,GALNT1,TNXB,NEGR1,W[illegible]

[illegible]DN,CTTNBP2,TWF2,PPP1R16B,CPEB1,TSHZ3,SHISA9,TENM2,KIF21B,TENM4,DOCK4,GLI3,NEURL1,RH[illegible]

[illegible]RB5,ZFYVE1,HLA-DRB1,TENM2,GALNT1,SYNE1,GALNT10,HLA-DQB1,CLCN3,PPP2R5C,DOCK4,RNF14[illegible]

[illegible]2,USP4,FHIT,DOCK8,SLC6A9,PKD1L3,GABBR1,DYSF,CNTN4,OLFM4,NEDD4L,CBLB,SEMA6D,ANKS1B,[illegible]

[illegible],DYNC1I1,USP4,SBK1,KLC1,FHIT,TNRC6A,MYO9B,CUL3,DOCK8,RPTOR,TRIM26,HDAC4,PPM1E,GABB[illegible]

[illegible],ANAPC4,TCF4,DYNC1I1,CTR9,USP4,FHIT,SGF29,RBM6,CUL3,TRIM26,HDAC4,PPM1E,L3MBTL4,TCF20,[illegible]

T,GPR141,FADS2,TSNARE1,KIZ,TMX2,DMXL2,PTCH1,PRKD1,PEF1,EXOC4,TYW1B,TMEM170B,DPP4,PK

D4L,CBLB,SND1,ANKS1B,FXR1,DMTF1,ARFGEF2,CITED1,CKB,TTLL7,ARID1B,NT5C2,PRPF3,FGFR1,CU

A5,CACNA2D2,CLSTN2,HFE,KCNG2,SLC6A9,CNTNAP2,PKD1L3,DPY19L1,PCLO,SPNS1,GABBR1,DYSF,S

MMP16,ITGA11,ZNF804A,CHRNA3,CACNA1C,ARID1B,RAET1E,FGFR1,HLA-DQA1,SLC13A1,AMIGO1,SDK

R1,ZFPM2,BSN,MPHOSPH9,PFAS,THOC7,NEDD4L,CBLB,SND1,SEMA6D,ANKS1B,FXR1,ZMIZ1,ZNF804A,

,ZFPM2,BSN,DYSF,THOC7,SND1,ZNF398,NR1D2,ANKS1B,FXR1,ZNF823,ZMIZ1,DMTF1,ZNF804A,CITED1

P4,BBS9,CYP2D7,SLC22A23,CHRNA5,CACNA2D2,CLSTN2,HFE,KCNG2,KLC1,MYO9B,CUL3,DOCK8,SLC

EDC2,CNOT1,PRKAG2,XRCC6,ETF1,CARMIL1,MEF2C,ATP23,GMIP,DDX3Y,RHOJ,MAD1L1,ZKSCAN3,CA

EMA6D,MMP16,EXT1,ITGA11,ATP2A2,CHRNA3,CACNA1C,RAET1E,FGFR1,HLA-DQA1,GLT8D1,SLC13A1

,1,CARMIL1,ATP23,GMIP,GRIK1,BTN2A1,RHOJ,DBN1,MAGI2,CALD1,RASGRP4,CADM2,CTNND1,SNAP91

KIF5C,ARFGEF2,CITED1,CKB,CACNA1C,TTLL7,ARID1B,NT5C2,ZNF638,FGFR1,CUEDC2,CNOT1,GUCY1

,DDX27,CKB,ZSCAN12,LDB1,ARID1B,ZNF638,PRPF3,FGFR1,CUEDC2,CNOT1,XRCC6,AUTS2,SLC30A9,

6A9,CNTNAP2,PKD1L3,DPY19L1,PCLO,SPNS1,GABBR1,MPHOSPH9,DYSF,CNTN4,SND1,SEMA6D,ANKS

LD1,RASGRP4,CTNND1,IMPA2,CTNNB1,NEK4,NGEF,STAG1,ANK3,MED27,PRKG1,SRR,FBXL17,DPYD,C

,AMIGO1,SLC30A9,SDK1,GRIK1,BTN2A1,MGAT5B,CADM2,ST3GAL3,ADTRP,SYT1,PCGF3,KCNB1,PTPRI

,CD55,CTNNB1,ADTRP,NTM,SYT1,CACNB2,KCNB1,ANK3,PTPRF,FURIN,DDN,PRKG1,CACNA1I,SRR,CN

A2,XRCC6,AUTS2,SLC30A9,ETF1,CARMIL1,MEF2C,DDX3Y,RHOJ,MAD1L1,DBN1,ZKSCAN3,MAGI2,CALD

CARMIL1,CREB5,ZSCAN31,MEF2C,ZNF568,ZNF19,DDX3Y,MAD1L1,ZKSCAN3,MAGI2,PALB2,CTNND1,SO

1B,FXR1,MMP16,EXT1,ITGA11,CALU,ATP2A2,ARFGEF2,CHRNA3,CACNA1C,RAET1E,FGFR1,HLA-DQA1

PEB1,TANK,CADPS,ZCCHC7,PSPH,FOXO3,KDM4A,GBF1,PIP5K1B,PDE4B,RNF111,RBL2,POLR2E,PPP2

F,ABCB9,FURIN,CACNA1I,CNNM2,MICA,LRFN5,POM121C,SLC35F2,RFT1,PTPRO,CLEC17A,TMEM219,H

NM2,PPP1R16B,CPEB1,MICA,PTPRO,TMEM219,HLA-DRB5,KEL,SHISA9,KCNN2,BTN3A2,HLA-DRB1,RG

1,RASGRP4,CTNND1,FSCN3,IMPA2,CTNNB1,NEK4,NGEF,SYT1,ANK3,DDN,PRKG1,SRR,CTTNBP2,TWF

X5,CTNNB1,STAG1,PCGF3,MED27,DDN,FBXL17,PPP1R16B,ESRRB,CPEB1,POM121C,MAML3,BCL11A,

,GLT8D1,CNOT1,SLC13A1,XRCC6,AMIGO1,SLC30A9,SDK1,CARMIL1,GRIK1,DDX3Y,BTN2A1,MGAT5B,RI

R2B,RGS6,PPP1R13B,SRPK2,ELMO1,DPP8,PREX1,PPP2R5C,BRWD1,DOCK4,SORBS1,AMBRA1,GLI3,RI

LA-DRB5,KEL,SHISA9,KCNN2,SORCS3,BTN3A2,HLA-DRB1,TENM2,NTRK3,GALNT1,SYNE1,GALNT10,HL

6,PPP1R13B,TENM2,MANBA,NTRK3,NEGR1,HLA-DQB1,PPP1R16A,ELMO1,WNT3,PCDH9,CLCN3,SLC39

2,ESRRB,PLCL2,DPYD,CPEB1,TANK,MICA,BCL11A,PSPH,FOXO3,KDM4A,GBF1,NKIRAS1,RNF111,PMFB

CCHC7,TSHZ3,FOXO3,LCORL,KDM4A,JMJD1C,RFC2,ZNF600,RNF111,RBL2,POLR2E,ZMIZ2,RGS6,PPP1

HOJ,DBN1,MAGI2,CALD1,RASGRP4,CADM2,CTNND1,SNAP91,CD55,CTNNB1,ST3GAL3,NGEF,ADTRP,N

A-DQB1,IMMP2L,PCDH9,CLCN3,SLC39A8,ATP2B2,BTN2A2,PPP2R5C,TENM4,IGSF9B,BTN3A1,NAALADL

A8,PREX1,ATP2B2,BTN2A2,TENM4,DOCK4,IGSF9B,SORBS1,NEURL1,RHOA,BTN3A1,RNF144A,CAMKV

P1,GLCCI1,PARD3B,PPP2R2B,RGS6,PPP1R13B,CCDC88C,SGSM2,SRPK2,NTRK3,SYNE1,ELMO1,DPP8,

R13B,TENM2,NFIX,SRPK2,SYNE1,GATAD2B,ZNF615,NOVA1,THRB,MACROD2,PPP2R5C,BRWD1,TENM

TM,SYT1,CACNB2,PCGF3,KCNB1,ANK3,PTPRF,ABCB9,FURIN,DDN,CACNA1I,CNNM2,PPP1R16B,CPEB1,

WNT3,PPA2,PREX1,ATP2B2,KIF21B,BRWD1,TENM4,DOCK4,TRAIP,SORBS1,FOXO6,AMBRA1,GLI3,NEU

4,TRAIP,KDM3B,TRIM33,SORBS1,BCL11B,FOXO6,CSRNP3,MED8,GLI3,ZKSCAN8,RHOA,PIK3R2,AFF3,P[illegible]

,CADPS,MICA,LRFN5,POM121C,SLC35F2,FOXO3,RFT1,PTPRO,CLEC17A,TMEM219,GBF1,PIP5K1B,HLA-

-DRB5,KEL,PDE4B,NKIRAS1,SHISA9,KCNN2,ZFYVE1,SORCS3,BTN3A2,PARD3B,PPP2R2B,HLA-DRB1,R[illegible]

GS6,TENM2,NTRK3,GALNT1,NEGR1,SYNE1,GALNT10,HLA-DQB1,PPP1R16A,ELMO1,IMMP2L,PCDH9,CL[illegible]

[illegible]N3,SLC39A8
