## Supplemental table 6 for "Digestive Dimensions of Autism: A Multiscale Exploration of Gut-Brain Interactions"

| Pathway | Total | Expected | Hits | P.Value | FDR | Genes |
|---|---|---|---|---|---|---|
| MHC class II receptor activity | 12 | 0.293 | 3 | 0.00269 | 0.333 | HLA-DQA1,HLA-DRB1,HLA-DQB1 |
| Acetylcholine receptor activity | 13 | 0.317 | 3 | 0.00343 | 0.333 | ANXA9,CHRNA5,CHRNA3 |
| Chromatin binding | 413 | 10.1 | 19 | 0.0063 | 0.407 | KDM5D,RERE,FOXP1,NCOA5,WR[illegible] |
| Phosphoric diester hydrolase activity | 55 | 1.34 | 5 | 0.0108 | 0.52 | PDE1C,PLCL1,PDE8B,PLCL2,PDE[illegible] |
| DNA_binding transcription factor activity | 903 | 22 | 33 | 0.0134 | 0.52 | KMT2A,FOXP1,ESR2,ESRRG,ZSC[illegible] |
| Voltage_gated calcium channel activity | 41 | 1 | 4 | 0.0173 | 0.561 | CACNA2D2,CACNA1C,CACNB2,C[illegible] |
| Calmodulin binding | 193 | 4.71 | 10 | 0.0202 | 0.561 | NRGN,SLC8A3,PDE1C,MYO9B,CA[illegible] |
| Nuclear receptor activity | 45 | 1.1 | 4 | 0.0237 | 0.574 | ESR2,NR1D2,ESRRB,THRB |
| Calcium_dependent phospholipid binding | 52 | 1.27 | 4 | 0.0377 | 0.737 | ANXA9,PCLO,DYSF,SYT1 |
| Microtubule motor activity | 76 | 1.85 | 5 | 0.038 | 0.737 | DYNC1I1,KLC1,KIF5C,KIF21B,DNA[illegible] |
| Translation regulator activity | 16 | 0.39 | 2 | 0.0568 | 0.919 | FXR1,CPEB1 |
| Ligand_gated ion channel activity | 36 | 0.878 | 3 | 0.0568 | 0.919 | CHRNA5,CHRNA3,GRIK1 |
| Cation transmembrane transporter activity | 17 | 0.415 | 2 | 0.0634 | 0.946 | PKD1L3,SLC30A9 |
| Potassium channel regulator activity | 40 | 0.976 | 3 | 0.0733 | 1 | DPP6,NEDD4L,AMIGO1 |

| | | | | | | |
|---|---|---|---|---|---|---|
| Guanyl_nucleotide exchange factor activity | 184 | 4.49 | 8 | 0.0818 | 1 | DOCK8,ARFGEF2,RASGRP4,NGE[illegible] |
| Ion channel activity | 202 | 4.93 | 8 | 0.122 | 1 | GRIN2A,CHRNA5,KCNG2,CHRNA[illegible] |
| Motor activity | 112 | 2.73 | 5 | 0.139 | 1 | MYO18A,DYNC1I1,KLC1,MYO9B,D[illegible] |
| Cation channel activity | 28 | 0.683 | 2 | 0.148 | 1 | GRIN2A,PKD1L3 |
| Translation factor activity, RNA binding | 28 | 0.683 | 2 | 0.148 | 1 | GATB,CPEB1 |
| Serine_type peptidase activity | 149 | 3.63 | 6 | 0.157 | 1 | DPP6,DPP4,F2,FURIN,DPP8,IMMP[illegible] |
| Translation release factor activity | 7 | 0.171 | 1 | 0.159 | 1 | ETF1 |
| Voltage_gated ion channel activity | 151 | 3.68 | 6 | 0.164 | 1 | CACNA2D2,KCNG2,CACNA1C,CA[illegible] |
| Guanylate cyclase activity | 8 | 0.195 | 1 | 0.179 | 1 | GUCY1A2 |
| Deacetylase activity | 9 | 0.22 | 1 | 0.199 | 1 | MACROD2 |
| Damaged DNA binding | 64 | 1.56 | 3 | 0.205 | 1 | RPA3,MSH5,XRCC6 |
| Transferase activity, transferring glycosyl groups | 233 | 5.68 | 8 | 0.21 | 1 | GALNT12,DPY19L1,EXT1,GLT8D1,[illegible] |
| DNA_binding transcription factor activity, RNA polymerase II_specific | 1560 | 38 | 43 | 0.216 | 1 | KDM5D,NPAS3,KMT2D,KMT2A,RE[illegible] |
| DNA binding | 2280 | 55.7 | 61 | 0.237 | 1 | KDM5D,NPAS3,KMT2D,MYO18A,K[illegible] |

| | | | | | | |
|---|---|---|---|---|---|---|
| Actin binding | 373 | 9.1 | 11 | 0.303 | 1 | MYO18A,MAPT,MYO9B,DBN1,CAL |
| Transcription coregulator activity | 79 | 1.93 | 3 | 0.303 | 1 | SND1,LDB1,MED8 |
| Ubiquitin_protein transferase activity | 303 | 7.39 | 9 | 0.321 | 1 | FBXL18,ANAPC4,CUL3,NEDD4L,CI |
| Glutamate receptor activity | 16 | 0.39 | 1 | 0.327 | 1 | GRIK1 |
| Nucleotide binding | 1810 | 44.2 | 47 | 0.35 | 1 | MTHFD1L,MAGI1,MYO18A,DGKI,G |
| Transmembrane receptor protein tyrosine kinase activity | 52 | 1.27 | 2 | 0.363 | 1 | FGFR1,NTRK3 |
| Oxidoreductase activity | 590 | 14.4 | 16 | 0.368 | 1 | KDM5D,MSRA,ENOX1,AGMO,PXD |
| Double_stranded DNA binding | 128 | 3.12 | 4 | 0.38 | 1 | MAPT,ZNF638,XRCC6,AFF3 |
| Phosphatase activity | 128 | 3.12 | 4 | 0.38 | 1 | PTPRN2,PTPRF,PSPH,PTPRO |
| DNA helicase activity | 21 | 0.512 | 1 | 0.405 | 1 | WRN |
| Kinase activity | 688 | 16.8 | 18 | 0.414 | 1 | NRGN,DGKI,VRK2,PRKD1,MAST3, |
| Signaling receptor binding | 413 | 10.1 | 11 | 0.426 | 1 | HLA-B,BANK1,DPP4,F2,HFE,BTN2 |
| Endoribonuclease activity | 23 | 0.561 | 1 | 0.434 | 1 | SND1 |
| Calcium ion binding | 703 | 17.1 | 18 | 0.45 | 1 | FKBP9,ANXA9,CDHR4,PEF1,F2,CL |
| Aspartic_type endopeptidase activity | 25 | 0.61 | 1 | 0.461 | 1 | SPPL3 |
| Voltage_gated potassium channel activity | 64 | 1.56 | 2 | 0.465 | 1 | KCNG2,KCNB1 |

| | | | | | | |
|---|---|---|---|---|---|---|
| Ligase activity | 144 | 3.51 | 4 | 0.468 | 1 | MTHFD1L,PFAS,TTLL7,GATB |
| Phosphoprotein phosphatase activity | 145 | 3.54 | 4 | 0.473 | 1 | PTPRN2,PPM1E,PTPRF,PTPRO |
| Lipid transporter activity | 26 | 0.634 | 1 | 0.474 | 1 | RFT1 |
| Antioxidant activity | 26 | 0.634 | 1 | 0.474 | 1 | KDM3B |
| Helicase activity | 146 | 3.56 | 4 | 0.478 | 1 | WRN,DDX27,XRCC6,DDX3Y |
| Translation elongation factor activity | 27 | 0.659 | 1 | 0.487 | 1 | EFL1 |
| Microtubule binding | 234 | 5.71 | 6 | 0.509 | 1 | MAPT,DYNC1I1,DYSF,KIF5C,CCD( |
| Catalytic activity | 524 | 12.8 | 13 | 0.515 | 1 | WRN,TYW1B,FHIT,PPM1E,OLFM4, |
| Hydrolase activity, hydrolyzing O_glycosyl compounds | 31 | 0.756 | 1 | 0.535 | 1 | MANBA |
| Peroxidase activity | 31 | 0.756 | 1 | 0.535 | 1 | PXDNL |
| Lyase activity | 158 | 3.85 | 4 | 0.54 | 1 | TYW1B,GUCY1A2,XRCC6,SRR |
| Sodium channel regulator activity | 32 | 0.78 | 1 | 0.547 | 1 | NEDD4L |
| Peptidase activity | 537 | 13.1 | 13 | 0.551 | 1 | THSD4,PSMA5,SPPL3,DPP4,F2,US |
| SNAP receptor activity | 33 | 0.805 | 1 | 0.558 | 1 | TSNARE1 |
| Calcium channel regulator activity | 34 | 0.829 | 1 | 0.568 | 1 | PRKG1 |
| Metallopeptidase activity | 166 | 4.05 | 4 | 0.58 | 1 | ADAM22,MMP16,ATP23,KEL |
| Nucleic acid binding | 1390 | 33.8 | 33 | 0.585 | 1 | ENOX1,WRN,ZSCAN9,ZNF536,RBI |

| | | | | | | |
|---|---|---|---|---|---|---|
| Extracellular matrix structural constituent | 124 | 3.02 | 3 | 0.586 | 1 | THSD4,COL11A1,TNXB |
| Carbohydrate binding | 212 | 5.17 | 5 | 0.593 | 1 | GALNT12,PKD1L3,CLEC17A,GALN |
| Hydrolase activity | 1680 | 41.1 | 40 | 0.598 | 1 | PDE1C,PTPRN2,THSD4,PSMA5,P |
| Isomerase activity | 128 | 3.12 | 3 | 0.608 | 1 | FKBP9,SRR,PUS7 |
| Transmembrane transporter activity | 130 | 3.17 | 3 | 0.618 | 1 | SLC22A23,ABCB9,SLC35F2 |
| Cytoskeletal protein binding | 88 | 2.15 | 2 | 0.636 | 1 | ANK3,SORBS1 |
| DNA_directed 5'_3' RNA polymerase activity | 41 | 1 | 1 | 0.637 | 1 | POLR2E |
| Nuclease activity | 134 | 3.27 | 3 | 0.638 | 1 | PXDNL,WRN,SND1 |
| Protein kinase activity | 532 | 13 | 12 | 0.65 | 1 | VRK2,PRKD1,MAST3,SBK1,FGFR1 |
| Serine_type endopeptidase inhibitor activity | 93 | 2.27 | 2 | 0.666 | 1 | ITIH3,FURIN |
| Methyltransferase activity | 189 | 4.61 | 4 | 0.681 | 1 | KMT2D,KMT2A,MRM2,TFB1M |
| Single_stranded DNA binding | 96 | 2.34 | 2 | 0.683 | 1 | RPA3,MAPT |
| ATPase activity, coupled to transmembrane movement of substances | 50 | 1.22 | 1 | 0.71 | 1 | ABCB9 |
| Enzyme activator activity | 51 | 1.24 | 1 | 0.717 | 1 | MMP16 |

| | | | | | | |
|---|---|---|---|---|---|---|
| Translation initiation factor activity | 56 | 1.37 | 1 | 0.75 | 1 | EIF1AY |
| Lipid binding | 304 | 7.41 | 6 | 0.755 | 1 | ERLIN1,ESR2,DYSF,CD55,CADPS, |
| Transporter activity | 137 | 3.34 | 2 | 0.851 | 1 | SLC22A23,SLC13A1 |
| Transferase activity | 1810 | 44.2 | 38 | 0.861 | 1 | GALNT12,KMT2D,KMT2A,MSL2,DG |
| GTPase activity | 302 | 7.37 | 5 | 0.864 | 1 | GNAI1,EFL1,RHOJ,NKIRAS1,RHOA |
| MRNA binding | 162 | 3.95 | 2 | 0.909 | 1 | FXR1,NOVA1 |
| Growth factor activity | 165 | 4.02 | 2 | 0.914 | 1 | MDK,F2 |
| Structural constituent of cytoskeleton | 104 | 2.54 | 1 | 0.924 | 1 | ANK3 |
| RNA binding | 1510 | 36.8 | 29 | 0.932 | 1 | MYO18A,CISD2,ENOX1,GIGYF2,N[illegible] |
| Peptidase inhibitor activity | 121 | 2.95 | 1 | 0.95 | 1 | ITIH3 |
| Cysteine_type peptidase activity | 159 | 3.88 | 1 | 0.981 | 1 | USP4 |
| Structural constituent of ribosome | 161 | 3.93 | 1 | 0.982 | 1 | MRPL33 |
| Molecular_function | 726 | 17.7 | 10 | 0.985 | 1 | TMX2,BBS9,ZNF804A,AUTS2,BTN[illegible] |
| Protein binding | 9580 | 234 | 211 | 0.992 | 1 | FBXL18,KDM5D,NREP,MAGI1,ANX |
| G protein_coupled receptor activity | 793 | 19.3 | 3 | 1 | 1 | GPR26,GPR141,GABBR1 |

,UTY,TCF4,HDAC4,CITED1,LDB1,AUTS2,SLC30A9,MEF2C,ZKSCAN3,CTNNB1,STAG1,TSHZ3,KDM4A,GI

AN9,NPAS1,TCF4,L3MBTL4,TCF20,NR1D2,DMTF1,CITED1,ZSCAN12,SLC30A9,CREB5,ZSCAN31,MEF2C,

RE,FOXP1,ESR2,ESRRG,ZSCAN9,ZNF536,NPAS1,TCF4,ZFPM2,ZNF398,NR1D2,ZNF823,DMTF1,ZSCAN1

MT2A,RPA3,RERE,FOXP1,ESR2,MAPT,ESRRG,WRN,TET2,ZSCAN9,ZNF536,TAF1C,NPAS1,TCF4,RBM6,

NAI1,VRK2,WRN,EFL1,PRKD1,TYW1B,MAST3,SBK1,FHIT,MYO9B,PFAS,ATP2A2,KIF5C,DDX27,CKB,TTLL

SBK1,CKB,FGFR1,PRKAG2,MAGI2,NEK4,PRKG1,PIP5K1B,SRPK2,NTRK3,RPS6KL1,CAMKV,TLK1

M6,TNRC6A,ZFPM2,SND1,ZNF398,FXR1,ZNF823,ZNF804A,DDX27,ZSCAN12,ZNF638,CREB5,ZSCAN31,Z [illegible]DNL,SPPL3,WRN,PLCL1,DPP4,F2,PDE8B,USP4,FHIT,NT5DC2,HDAC4,PPM1E,SND1,MMP16,ATP2A2,DD[illegible]

GKI,VRK2,PRKD1,MRM2,MAST3,SBK1,DPY19L1,TRIM26,NEDD4L,CBLB,EXT1,CKB,GATB,FGFR1,GLT8D1

COA5,MAPT,PEF1,RBM6,TNRC6A,TCF20,THOC7,SND1,FXR1,DDX27,EIF1AY,ZNF638,PRPF3,CNOT1,XR[illegible]

A9,SEMA3E,KMT2D,MYO18A,KMT2A,ERLIN1,HLA-B,DGKI,RPA3,AKAP6,KMT2E,GNAI1,ADAM22,RERE,CI

ZNF19,ZKSCAN3,SOX5,CTNNB1,PCGF3,ESRRB,BCL11A,FOXO3,NFIX,THRB,BCL11B,FOXO6,CSRNP3,G

2,ARID1B,SLC30A9,CREB5,ZSCAN31,MEF2C,ZNF568,ZNF19,ZKSCAN3,SOX5,ESRRB,BCL11A,TSHZ3,FC

TRIM26,HDAC4,TCF20,ZFPM2,ZNF398,NR1D2,ZNF823,DMTF1,ZSCAN12,LDB1,ARID1B,ZNF638,MSH5,XF

_7,GATB,NT5C2,FGFR1,MSH5,GUCY1A2,PRKAG2,XRCC6,DDX3Y,RHOJ,NEK4,ABCB9,PRKG1,SRR,DHRS

NF568,ZNF19,DDX3Y,ZKSCAN3,CPEB1,BCL11A,ZCCHC7,TSHZ3,ZNF600,ZNF615,NOVA1,BCL11B,GLI3,Z

DX27,NT5C2,XRCC6,ATP23,DDX3Y,IMPA2,ADTRP,PTPRF,FURIN,PLCL2,PSPH,PTPRO,KEL,PDE4B,MANB

,FTCDNL1,MGAT5B,TFB1M,NEK4,ST3GAL3,PRKG1,PIP5K1B,RNF111,SRPK2,NTRK3,GALNT1,GALNT10,I

SD2,FOXP1,RTN1,HLA-DMA,ENOX1,GRIN2A,GIGYF2,FEZ1,PSMA5,VRK2,RSRC1,RSU1,NCOA5,SPPL3,E

)XO3,LCORL,ZNF600,NFIX,GATAD2B,ZNF615,THRB,BCL11B,FOXO6,CSRNP3,GLI3,ZKSCAN8,ZNF800

RCC6,CREB5,ZSCAN31,MEF2C,ZNF568,ZNF19,DDX3Y,TFB1M,ZKSCAN3,PALB2,SOX5,ESRRB,TSHZ3,FO [illegible]11,DPYD,RFC2,PIP5K1B,NKIRAS1,SRPK2,NTRK3,CLCN3,ATP2B2,RPS6KL1,KIF21B,RHOA,DNAH11,TLK

SR2,MAPT,ESRRG,WRN,TET2,KIZ,ZSCAN9,PTCH1,PRKD1,PEF1,EXOC4,TMEM170B,DPP4,PKP4,BBS9,S

XO3,LCORL,RFC2,ZNF600,RBL2,POLR2E,NFIX,ZNF615,THRB,TRIM33,FOXO6,CSRNP3,GLI3,ZKSCAN8,A

[illegible]LC22A23,F2,CHRNA5,TAF1C,ANAPC4,TCF4,DYNC1I1,HFE,CTR9,USP4,MAST3,KLC1,FHIT,SGF29,RBM6

[illegible],TNRC6A,MYO9B,CUL3,DOCK8,CNTNAP2,PKD1L3,RPTOR,TRIM26,HDAC4,PPM1E,L3MBTL4,SPNS1,TC[illegible]

F20,GABBR1,ZFPM2,DYSF,OLFM4,THOC7,NEDD4L,CBLB,SND1,ZNF398,NR1D2,FXR1,CALU,ATP2A2,KIF

5C,ARFGEF2,CITED1,DDX27,CKB,CHRNA3,CACNA1C,ZSCAN12,LDB1,ARID1B,EIF1AY,RAET1E,NT5C2,P

'RPF3,FGFR1,MSH5,LRRC20,CUEDC2,CNOT1,GUCY1A2,XRCC6,AUTS2,ETF1,CARMIL1,CREB5,MEF2C,(

[illegible]GMIP,MGAT5B,RHOJ,MAD1L1,DBN1,TFB1M,ZKSCAN3,MAGI2,CALD1,PALB2,CTNND1,SNAP91,SOX5,CD[illegible]

55,IMPA2,CTNNB1,STAG1,NTM,SYT1,CACNB2,PCGF3,KCNB1,ANK3,ABCB9,FURIN,MED27,DDN,PRKG1,

CACNA1I,TWF2,PPP1R16B,MYBPHL,DPYD,CPEB1,TANK,IVD,CADPS,MICA,POM121C,ZCCHC7,TSHZ3,F[illegible]

OXO3,LCORL,PTPRO,KDM4A,JMJD1C,TMEM219,RFC2,GBF1,PIP5K1B,KEL,RNF111,KCNN2,RBL2,POLR2

2E,Z
