## Supplemental table 7 for "Digestive Dimensions of Autism: A Multiscale Exploration of Gut-Brain Interactions"

| Pathway | Total | Expected | Hits | P.Value | FDR | Genes |
| --- | --- | --- | --- | --- | --- | --- |
| Type I diabetes mellitus | 43 | 2.15 | 7 | 0.00502 | 1 | PTPRN2,HLA-B,HLA-DMA,HLA-DQ[illegible] |
| Allograft rejection | 38 | 1.9 | 6 | 0.0107 | 1 | HLA-B,HLA-DMA,HLA-DQA1,HLA-[illegible] |
| Graft-versus-host disease | 41 | 2.05 | 6 | 0.0153 | 1 | HLA-B,HLA-DMA,HLA-DQA1,HLA-[illegible] |
| Asthma | 31 | 1.55 | 5 | 0.0177 | 1 | HLA-DMA,HLA-DQA1,HLA-DRB5,H[illegible] |
| Viral myocarditis | 59 | 2.95 | 7 | 0.0269 | 1 | HLA-B,HLA-DMA,HLA-DQA1,CD55,[illegible] |
| Autoimmune thyroid disease | 53 | 2.65 | 6 | 0.0477 | 1 | HLA-B,HLA-DMA,HLA-DQA1,HLA-[illegible] |
| Intestinal immune network for IgA production | 49 | 2.45 | 5 | 0.0965 | 1 | HLA-DMA,HLA-DQA1,HLA-DRB5,H[illegible] |
| Adherens junction | 72 | 3.6 | 6 | 0.15 | 1 | FGFR1,CTNND1,CTNNB1,PTPRF,[illegible] |
| Arrhythmogenic right ventricular cardiomyopathy (ARVC) | 72 | 3.6 | 6 | 0.15 | 1 | CACNA2D2,ITGA11,ATP2A2,CACN[illegible] |
| Antigen processing and presentation | 77 | 3.85 | 6 | 0.187 | 1 | HLA-B,HLA-DMA,HLA-DQA1,HLA-[illegible] |
| Mucin type O-glycan biosynthesis | 31 | 1.55 | 3 | 0.201 | 1 | GALNT12,GALNT1,GALNT10 |
| Inflammatory bowel disease (IBD) | 65 | 3.25 | 5 | 0.224 | 1 | HLA-DMA,HLA-DQA1,HLA-DRB5,H[illegible] |
| Hypertrophic cardiomyopathy (HCM) | 85 | 4.25 | 6 | 0.251 | 1 | CACNA2D2,ITGA11,ATP2A2,CACN[illegible] |
| Staphylococcus aureus infection | 68 | 3.4 | 5 | 0.253 | 1 | HLA-DMA,HLA-DQA1,HLA-DRB5,H[illegible] |

| | | | | | | |
|---|---|---|---|---|---|---|
| Adrenergic signaling in cardiomyocytes | 145 | 7.25 | 9 | 0.301 | 1 | GNAI1,CACNA2D2,ATP2A2,CACNA |
| Cell adhesion molecules (CAMs) | 146 | 7.3 | 9 | 0.308 | 1 | HLA-B,HLA-DMA,CNTNAP2,HLA-D |
| Leishmaniasis | 74 | 3.7 | 5 | 0.311 | 1 | HLA-DMA,HLA-DQA1,HLA-DRB5,H |
| Th1 and Th2 cell differentiation | 92 | 4.6 | 6 | 0.312 | 1 | HLA-DMA,HLA-DQA1,MAML3,HLA- |
| cGMP-PKG signaling pathway | 166 | 8.3 | 10 | 0.318 | 1 | SLC8A3,GNAI1,ATP2A2,CACNA1C |
| Mannose type O-glycan biosynthesis | 23 | 1.15 | 2 | 0.321 | 1 | MGAT5B,ST3GAL3 |
| Mismatch repair | 23 | 1.15 | 2 | 0.321 | 1 | RPA3,RFC2 |
| Hematopoietic cell lineage | 97 | 4.85 | 6 | 0.357 | 1 | HLA-DMA,HLA-DQA1,CD55,HLA-D |
| Circadian entrainment | 97 | 4.85 | 6 | 0.357 | 1 | GNAI1,GRIN2A,CACNA1C,GUCY1 |
| cAMP signaling pathway | 212 | 10.6 | 12 | 0.372 | 1 | GNAI1,GRIN2A,PTCH1,GABBR1,A |
| Longevity regulating pathway - multiple species | 62 | 3.1 | 4 | 0.376 | 1 | RPTOR,PRKAG2,FOXO3,PIK3R2 |
| Basal cell carcinoma | 63 | 3.15 | 4 | 0.388 | 1 | PTCH1,CTNNB1,WNT3,GLI3 |
| AMPK signaling pathway | 120 | 6 | 7 | 0.394 | 1 | RPTOR,PRKAG2,CREB5,FOXO3,P |
| Hedgehog signaling pathway | 47 | 2.35 | 3 | 0.42 | 1 | PTCH1,CUL3,GLI3 |
| Cocaine addiction | 49 | 2.45 | 3 | 0.447 | 1 | GNAI1,GRIN2A,CREB5 |
| Renin secretion | 69 | 3.45 | 4 | 0.456 | 1 | PDE1C,GNAI1,CACNA1C,GUCY1A |

| | | | | | | |
|---|---|---|---|---|---|---|
| Rap1 signaling pathway | 206 | 10.3 | 11 | 0.456 | 1 | MAGI1,GNAI1,GRIN2A,PRKD1,FGF[illegible] |
| Longevity regulating pathway | 89 | 4.45 | 5 | 0.461 | 1 | RPTOR,PRKAG2,CREB5,FOXO3,P[illegible] |
| Purine metabolism | 130 | 6.5 | 7 | 0.477 | 1 | PDE1C,PDE8B,FHIT,PFAS,NT5C2,[illegible] |
| Morphine addiction | 91 | 4.55 | 5 | 0.481 | 1 | PDE1C,GNAI1,PDE8B,GABBR1,PD[illegible] |
| Rheumatoid arthritis | 91 | 4.55 | 5 | 0.481 | 1 | HLA-DMA,HLA-DQA1,HLA-DRB5,H[illegible] |
| Dilated cardiomyopathy | 91 | 4.55 | 5 | 0.481 | 1 | CACNA2D2,ITGA11,ATP2A2,CACN[illegible] |
| Dopaminergic synapse | 131 | 6.55 | 7 | 0.485 | 1 | GNAI1,GRIN2A,KIF5C,CACNA1C,C[illegible] |
| Non-homologous end-joining | 13 | 0.65 | 1 | 0.487 | 1 | XRCC6 |
| Toxoplasmosis | 113 | 5.65 | 6 | 0.501 | 1 | GNAI1,HLA-DMA,HLA-DQA1,HLA-[illegible] |
| Oxytocin signaling pathway | 153 | 7.65 | 8 | 0.501 | 1 | GNAI1,CACNA2D2,CACNA1C,GUC[illegible] |
| Bacterial invasion of epithelial cells | 74 | 3.7 | 4 | 0.511 | 1 | CTNNB1,ELMO1,RHOA,PIK3R2 |
| Glycosaminoglycan biosynthesis - keratan sulfate | 14 | 0.7 | 1 | 0.513 | 1 | ST3GAL3 |
| Regulation of lipolysis in adipocytes | 55 | 2.75 | 3 | 0.524 | 1 | GNAI1,PRKG1,PIK3R2 |
| DNA replication | 36 | 1.8 | 2 | 0.544 | 1 | RPA3,RFC2 |
| Aldosterone synthesis and secretion | 98 | 4.9 | 5 | 0.547 | 1 | PRKD1,CACNA1C,CREB5,CACNA1[illegible] |
| Cardiac muscle contraction | 78 | 3.9 | 4 | 0.553 | 1 | CACNA2D2,ATP2A2,CACNA1C,CA[illegible] |

| | | | | | | |
|---|---|---|---|---|---|---|
| Progesterone-mediated oocyte maturation | 99 | 4.95 | 5 | 0.556 | 1 | GNAI1,ANAPC4,MAD1L1,CPEB1,PI |
| Axon guidance | 181 | 9.05 | 9 | 0.557 | 1 | SEMA3E,GNAI1,SEMA3A,PTCH1,S |
| Aldosterone-regulated sodium reabsorption | 37 | 1.85 | 2 | 0.559 | 1 | NEDD4L,PIK3R2 |
| Endometrial cancer | 58 | 2.9 | 3 | 0.56 | 1 | CTNNB1,FOXO3,PIK3R2 |
| Lysine degradation | 59 | 2.95 | 3 | 0.572 | 1 | KMT2D,KMT2A,KMT2E |
| Long-term depression | 60 | 3 | 3 | 0.584 | 1 | GNAI1,GUCY1A2,PRKG1 |
| Platelet activation | 124 | 6.2 | 6 | 0.593 | 1 | GNAI1,F2,GUCY1A2,PRKG1,RHOA |
| Glycine, serine and threonine metabolism | 40 | 2 | 2 | 0.602 | 1 | SRR,PSPH |
| Taste transduction | 83 | 4.15 | 4 | 0.602 | 1 | PDE1C,PKD1L3,GABBR1,CACNA1[illegible] |
| Other glycan degradation | 18 | 0.9 | 1 | 0.603 | 1 | MANBA |
| Homologous recombination | 41 | 2.05 | 2 | 0.615 | 1 | RPA3,PALB2 |
| Pantothenate and CoA biosynthesis | 19 | 0.95 | 1 | 0.623 | 1 | DPYD |
| Th17 cell differentiation | 107 | 5.35 | 5 | 0.626 | 1 | HLA-DMA,HLA-DQA1,HLA-DRB5,HI |
| Insulin secretion | 86 | 4.3 | 4 | 0.63 | 1 | PCLO,CACNA1C,CREB5,KCNN2 |
| Central carbon metabolism in cancer | 65 | 3.25 | 3 | 0.638 | 1 | FGFR1,NTRK3,PIK3R2 |

| | | | | | | |
|---|---|---|---|---|---|---|
| One carbon pool by folate | 20 | 1 | 1 | 0.642 | 1 | MTHFD1L |
| Phagosome | 152 | 7.6 | 7 | 0.643 | 1 | HLA-B,HLA-DMA,DYNC1I1,HLA-DQ |
| Non-small cell lung cancer | 66 | 3.3 | 3 | 0.649 | 1 | FHIT,FOXO3,PIK3R2 |
| Vasopressin-regulated water reabsorption | 44 | 2.2 | 2 | 0.654 | 1 | DYNC1I1,CREB5 |
| GABAergic synapse | 89 | 4.45 | 4 | 0.657 | 1 | GNAI1,PLCL1,GABBR1,CACNA1C |
| Systemic lupus erythematosus | 133 | 6.65 | 6 | 0.661 | 1 | HLA-DMA,GRIN2A,HLA-DQA1,HLA- |
| HTLV-I infection | 219 | 11 | 10 | 0.663 | 1 | HLA-B,HLA-DMA,ANAPC4,HLA-DQ |
| Leukocyte transendothelial migration | 112 | 5.6 | 5 | 0.666 | 1 | GNAI1,CTNND1,CTNNB1,RHOA,PI |
| Cholinergic synapse | 112 | 5.6 | 5 | 0.666 | 1 | GNAI1,CHRNA3,CACNA1C,CREB5 |
| Amphetamine addiction | 68 | 3.4 | 3 | 0.669 | 1 | GRIN2A,CACNA1C,CREB5 |
| Other types of O-glycan biosynthesis | 22 | 1.1 | 1 | 0.677 | 1 | ST3GAL3 |
| Type II diabetes mellitus | 46 | 2.3 | 2 | 0.678 | 1 | CACNA1C,PIK3R2 |
| Viral carcinogenesis | 201 | 10.1 | 9 | 0.682 | 1 | HLA-B,HDAC4,SND1,CREB5,MAD1 |
| Prolactin signaling pathway | 70 | 3.5 | 3 | 0.688 | 1 | ESR2,FOXO3,PIK3R2 |
| Insulin signaling pathway | 137 | 6.85 | 6 | 0.689 | 1 | RPTOR,CBLB,PRKAG2,PTPRF,SO |
| Nucleotide excision repair | 47 | 2.35 | 2 | 0.689 | 1 | RPA3,RFC2 |
| Protein export | 23 | 1.15 | 1 | 0.693 | 1 | IMMP2L |

| | | | | | | |
|---|---|---|---|---|---|---|
| Thyroid hormone signaling pathway | 116 | 5.8 | 5 | 0.696 | 1 | ATP2A2,CTNNB1,MED27,THRB,PI[illegible] |
| Signaling pathways regulating pluripotency of stem cells | 139 | 6.95 | 6 | 0.702 | 1 | FGFR1,CTNNB1,PCGF3,ESRRB,W[illegible] |
| Glycosaminoglycan biosynthesis - heparan sulfate / heparin | 24 | 1.2 | 1 | 0.709 | 1 | EXT1 |
| Sphingolipid signaling pathway | 119 | 5.95 | 5 | 0.717 | 1 | GNAI1,PPP2R2B,PPP2R5C,RHOA, |
| Prostate cancer | 97 | 4.85 | 4 | 0.722 | 1 | FGFR1,CREB5,CTNNB1,PIK3R2 |
| alpha-Linolenic acid metabolism | 25 | 1.25 | 1 | 0.723 | 1 | FADS2 |
| Phosphatidylinositol signaling system | 99 | 4.95 | 4 | 0.737 | 1 | DGKI,IMPA2,PIP5K1B,PIK3R2 |
| Glycosphingolipid biosynthesis - lacto and neolacto series | 27 | 1.35 | 1 | 0.75 | 1 | ST3GAL3 |
| Biosynthesis of unsaturated fatty acids | 27 | 1.35 | 1 | 0.75 | 1 | FADS2 |
| Fanconi anemia pathway | 54 | 2.7 | 2 | 0.76 | 1 | RPA3,PALB2 |
| Pathogenic Escherichia coli infection | 55 | 2.75 | 2 | 0.769 | 1 | CTNNB1,RHOA |

| | | | | | | |
|---|---|---|---|---|---|---|
| Nicotinate and nicotinamide metabolism | 30 | 1.5 | 1 | 0.786 | 1 | NT5C2 |
| Pyrimidine metabolism | 57 | 2.85 | 2 | 0.786 | 1 | NT5C2,DPYD |
| Insulin resistance | 108 | 5.4 | 4 | 0.796 | 1 | PRKAG2,CREB5,PTPRF,PIK3R2 |
| beta-Alanine metabolism | 31 | 1.55 | 1 | 0.797 | 1 | DPYD |
| RNA polymerase | 31 | 1.55 | 1 | 0.797 | 1 | POLR2E |
| Circadian rhythm | 31 | 1.55 | 1 | 0.797 | 1 | PRKAG2 |
| FoxO signaling pathway | 132 | 6.6 | 5 | 0.797 | 1 | PRKAG2,FOXO3,RBL2,FOXO6,PIK |
| Vascular smooth muscle contraction | 132 | 6.6 | 5 | 0.797 | 1 | CACNA1C,GUCY1A2,CALD1,PRKG |
| Autophagy - other | 32 | 1.6 | 1 | 0.807 | 1 | RPTOR |
| Colorectal cancer | 86 | 4.3 | 3 | 0.812 | 1 | CTNNB1,RHOA,PIK3R2 |
| Apelin signaling pathway | 137 | 6.85 | 5 | 0.823 | 1 | SLC8A3,GNAI1,HDAC4,PRKAG2,M |
| Gap junction | 88 | 4.4 | 3 | 0.824 | 1 | GNAI1,GUCY1A2,PRKG1 |
| Estrogen signaling pathway | 138 | 6.9 | 5 | 0.827 | 1 | GNAI1,ESR2,GABBR1,CREB5,PIK3 |
| Glutamatergic synapse | 114 | 5.7 | 4 | 0.829 | 1 | GNAI1,GRIN2A,CACNA1C,GRIK1 |
| Serotonergic synapse | 115 | 5.75 | 4 | 0.834 | 1 | GNAI1,CYP2D7,CACNA1C,KCNN2 |
| Salivary secretion | 90 | 4.5 | 3 | 0.835 | 1 | GUCY1A2,PRKG1,ATP2B2 |
| Protein digestion and absorption | 90 | 4.5 | 3 | 0.835 | 1 | SLC8A3,DPP4,COL11A1 |

| | | | | | | |
|---|---|---|---|---|---|---|
| Calcium signaling pathway | 188 | 9.4 | 7 | 0.838 | 1 | SLC8A3,PDE1C,GRIN2A,ATP2A2,C |
| mRNA surveillance pathway | 91 | 4.55 | 3 | 0.841 | 1 | ETF1,PPP2R2B,PPP2R5C |
| Mitophagy - animal | 65 | 3.25 | 2 | 0.844 | 1 | FOXO3,AMBRA1 |
| Influenza A | 167 | 8.35 | 6 | 0.849 | 1 | HLA-DMA,HLA-DQA1,HLA-DRB5,HI |
| Thyroid cancer | 37 | 1.85 | 1 | 0.851 | 1 | CTNNB1 |
| Neurotrophin signaling pathway | 119 | 5.95 | 4 | 0.853 | 1 | FOXO3,NTRK3,RHOA,PIK3R2 |
| Long-term potentiation | 67 | 3.35 | 2 | 0.856 | 1 | GRIN2A,CACNA1C |
| Breast cancer | 147 | 7.35 | 5 | 0.866 | 1 | ESR2,FGFR1,CTNNB1,WNT3,PIK3 |
| Phospholipase D signaling pathway | 148 | 7.4 | 5 | 0.87 | 1 | DGKI,F2,PIP5K1B,RHOA,PIK3R2 |
| Ferroptosis | 40 | 2 | 1 | 0.872 | 1 | SLC39A8 |
| Nicotine addiction | 40 | 2 | 1 | 0.872 | 1 | GRIN2A |
| Cell cycle | 124 | 6.2 | 4 | 0.875 | 1 | ANAPC4,MAD1L1,STAG1,RBL2 |
| Endocrine resistance | 98 | 4.9 | 3 | 0.875 | 1 | ESR2,CYP2D7,PIK3R2 |
| Pancreatic secretion | 98 | 4.9 | 3 | 0.875 | 1 | ATP2A2,ATP2B2,RHOA |
| Oocyte meiosis | 125 | 6.25 | 4 | 0.878 | 1 | ANAPC4,MAD1L1,CPEB1,PPP2R5( |
| Choline metabolism in cancer | 99 | 4.95 | 3 | 0.879 | 1 | DGKI,PIP5K1B,PIK3R2 |
| Melanoma | 72 | 3.6 | 2 | 0.882 | 1 | FGFR1,PIK3R2 |
| Epstein-Barr virus infection | 201 | 10.1 | 7 | 0.883 | 1 | HLA-B,HLA-DMA,HLA-DQA1,HLA-D |
| Proteoglycans in cancer | 201 | 10.1 | 7 | 0.883 | 1 | PTCH1,FGFR1,CTNNB1,ANK3,WN |
| T cell receptor signaling pathway | 101 | 5.05 | 3 | 0.888 | 1 | CBLB,RHOA,PIK3R2 |
| Melanogenesis | 101 | 5.05 | 3 | 0.888 | 1 | GNAI1,CTNNB1,WNT3 |

| | | | | | | |
|---|---|---|---|---|---|---|
| Autophagy - animal | 128 | 6.4 | 4 | 0.89 | 1 | RPTOR,ZFYVE1,AMBRA1,PIK3R2 |
| Tuberculosis | 179 | 8.95 | 6 | 0.89 | 1 | HLA-DMA,HLA-DQA1,HLA-DRB5,H[illegible] |
| Inositol phosphate metabolism | 74 | 3.7 | 2 | 0.891 | 1 | IMPA2,PIP5K1B |
| PPAR signaling pathway | 74 | 3.7 | 2 | 0.891 | 1 | FADS2,SORBS1 |
| Chagas disease (American trypanosomiasis) | 103 | 5.15 | 3 | 0.895 | 1 | GNAI1,PPP2R2B,PIK3R2 |
| Carbohydrate digestion and absorption | 44 | 2.2 | 1 | 0.896 | 1 | PIK3R2 |
| Natural killer cell mediated cytotoxicity | 131 | 6.55 | 4 | 0.9 | 1 | HLA-B,RAET1E,MICA,PIK3R2 |
| Pertussis | 76 | 3.8 | 2 | 0.9 | 1 | GNAI1,RHOA |
| ABC transporters | 45 | 2.25 | 1 | 0.901 | 1 | ABCB9 |
| Proteasome | 45 | 2.25 | 1 | 0.901 | 1 | PSMA5 |
| Synaptic vesicle cycle | 78 | 3.9 | 2 | 0.908 | 1 | SLC6A9,SYT1 |
| Transcriptional misregulation in cancer | 186 | 9.3 | 6 | 0.91 | 1 | KMT2A,UTY,LDB1,MEF2C,JMJD1C |
| EGFR tyrosine kinase inhibitor resistance | 79 | 3.95 | 2 | 0.912 | 1 | FOXO3,PIK3R2 |
| RNA degradation | 79 | 3.95 | 2 | 0.912 | 1 | CNOT1,ZCCHC7 |

| | | | | | | |
|---|---|---|---|---|---|---|
| Complement and coagulation cascades | 79 | 3.95 | 2 | 0.912 | 1 | F2,CD55 |
| Valine, leucine and isoleucine degradation | 48 | 2.4 | 1 | 0.915 | 1 | IVD |
| Notch signaling pathway | 48 | 2.4 | 1 | 0.915 | 1 | MAML3 |
| Ubiquitin mediated proteolysis | 137 | 6.85 | 4 | 0.918 | 1 | ANAPC4,CUL3,NEDD4L,CBLB |
| Chemokine signaling pathway | 190 | 9.5 | 6 | 0.92 | 1 | GNAI1,FOXO3,ELMO1,PREX1,RHC |
| ECM-receptor interaction | 82 | 4.1 | 2 | 0.922 | 1 | ITGA11,TNXB |
| Fluid shear stress and atherosclerosis | 139 | 6.95 | 4 | 0.923 | 1 | MEF2C,CTNNB1,RHOA,PIK3R2 |
| Arginine and proline metabolism | 50 | 2.5 | 1 | 0.924 | 1 | CKB |
| N-Glycan biosynthesis | 50 | 2.5 | 1 | 0.924 | 1 | MGAT5B |
| Endocytosis | 244 | 12.2 | 8 | 0.928 | 1 | HLA-B,NEDD4L,CBLB,KIF5C,ARFG |
| Amyotrophic lateral sclerosis (ALS) | 51 | 2.55 | 1 | 0.928 | 1 | GRIN2A |
| ErbB signaling pathway | 85 | 4.25 | 2 | 0.931 | 1 | CBLB,PIK3R2 |
| Tight junction | 170 | 8.5 | 5 | 0.933 | 1 | MAGI1,NEDD4L,PRKAG2,PPP2R2E |
| Salmonella infection | 86 | 4.3 | 2 | 0.934 | 1 | DYNC1I1, KLC1 |
| Fatty acid metabolism | 53 | 2.65 | 1 | 0.935 | 1 | FADS2 |

| | | | | | | |
|---|---|---|---|---|---|---|
| Alzheimer's disease | 171 | 8.55 | 5 | 0.935 | 1 | GRIN2A,MAPT,ATP2A2,CACNA1C, |
| PI3K-Akt signaling pathway | 354 | 17.7 | 12 | 0.946 | 1 | MAGI1,RPTOR,ITGA11,FGFR1,CRI |
| Fc gamma R-mediated phagocytosis | 91 | 4.55 | 2 | 0.947 | 1 | PIP5K1B,PIK3R2 |
| Small cell lung cancer | 93 | 4.65 | 2 | 0.951 | 1 | FHIT,PIK3R2 |
| Alcoholism | 180 | 9 | 5 | 0.951 | 1 | GNAI1,GRIN2A,HDAC4,CREB5,HD/ |
| VEGF signaling pathway | 59 | 2.95 | 1 | 0.952 | 1 | PIK3R2 |
| mTOR signaling pathway | 153 | 7.65 | 4 | 0.952 | 1 | RPTOR,WNT3,RHOA,PIK3R2 |
| Steroid hormone biosynthesis | 60 | 3 | 1 | 0.955 | 1 | DHRS11 |
| Glycerolipid metabolism | 61 | 3.05 | 1 | 0.957 | 1 | DGKI |
| Glycerophospholipid metabolism | 97 | 4.85 | 2 | 0.959 | 1 | DGKI,GPD2 |
| Kaposi's sarcoma-associated herpesvirus infection | 186 | 9.3 | 5 | 0.96 | 1 | HLA-B,CTNNB1,MICA,PREX1,PIK3I |
| Regulation of actin cytoskeleton | 214 | 10.7 | 6 | 0.961 | 1 | F2,ITGA11,FGFR1,PIP5K1B,RHOA, |
| Cytosolic DNA-sensing pathway | 63 | 3.15 | 1 | 0.961 | 1 | POLR2E |
| Cellular senescence | 160 | 8 | 4 | 0.963 | 1 | HLA-B,FOXO3,RBL2,PIK3R2 |
| Shigellosis | 65 | 3.25 | 1 | 0.965 | 1 | ELMO1 |
| Aminoacyl-tRNA biosynthesis | 66 | 3.3 | 1 | 0.967 | 1 | GATB |

| | | | | | | |
|---|---|---|---|---|---|---|
| Acute myeloid leukemia | 66 | 3.3 | 1 | 0.967 | 1 | PIK3R2 |
| MicroRNAs in cancer | 299 | 15 | 9 | 0.968 | 1 | FOXP1,RPTOR,HDAC4,ZFPM2,MM |
| Glucagon signaling pathway | 103 | 5.15 | 2 | 0.968 | 1 | PRKAG2,CREB5 |
| RNA transport | 165 | 8.25 | 4 | 0.969 | 1 | THOC7,FXR1,EIF1AY,POM121C |
| Fc epsilon RI signaling pathway | 68 | 3.4 | 1 | 0.97 | 1 | PIK3R2 |
| Adipocytokine signaling pathway | 69 | 3.45 | 1 | 0.971 | 1 | PRKAG2 |
| Renal cell carcinoma | 69 | 3.45 | 1 | 0.971 | 1 | PIK3R2 |
| RIG-I-like receptor signaling pathway | 70 | 3.5 | 1 | 0.973 | 1 | TANK |
| Focal adhesion | 199 | 9.95 | 5 | 0.974 | 1 | ITGA11,CTNNB1,TNXB,RHOA,PIK3 |
| B cell receptor signaling pathway | 71 | 3.55 | 1 | 0.974 | 1 | PIK3R2 |
| Drug metabolism - cytochrome P450 | 72 | 3.6 | 1 | 0.976 | 1 | CYP2D7 |
| Bile secretion | 72 | 3.6 | 1 | 0.976 | 1 | KCNN2 |
| TNF signaling pathway | 110 | 5.5 | 2 | 0.977 | 1 | CREB5,PIK3R2 |
| Platinum drug resistance | 73 | 3.65 | 1 | 0.977 | 1 | PIK3R2 |
| Thyroid hormone synthesis | 74 | 3.7 | 1 | 0.978 | 1 | CREB5 |
| Ras signaling pathway | 232 | 11.6 | 6 | 0.978 | 1 | SYNGAP1,GRIN2A,FGFR1,RASGR |
| Biosynthesis of amino acids | 75 | 3.75 | 1 | 0.979 | 1 | PSPH |

| | | | | | | |
|---|---|---|---|---|---|---|
| Gastric acid secretion | 75 | 3.75 | 1 | 0.979 | 1 | GNAI1 |
| Pancreatic cancer | 75 | 3.75 | 1 | 0.979 | 1 | PIK3R2 |
| Glioma | 75 | 3.75 | 1 | 0.979 | 1 | PIK3R2 |
| Metabolism of xenobiotics by cytochrome P450 | 76 | 3.8 | 1 | 0.98 | 1 | CYP2D7 |
| Chronic myeloid leukemia | 76 | 3.8 | 1 | 0.98 | 1 | PIK3R2 |
| Retrograde endocannabinoid signaling | 148 | 7.4 | 3 | 0.981 | 1 | GNAI1,CACNA1C,NDUFA2 |
| Non-alcoholic fatty liver disease (NAFLD) | 149 | 7.45 | 3 | 0.982 | 1 | PRKAG2,NDUFA2,PIK3R2 |
| Drug metabolism - other enzymes | 79 | 3.95 | 1 | 0.983 | 1 | DPYD |
| MAPK signaling pathway | 295 | 14.8 | 8 | 0.983 | 1 | MAPT,CACNA2D2,CACNA1C,FGF[illegible] |
| Hippo signaling pathway | 154 | 7.7 | 3 | 0.985 | 1 | CTNNB1,PPP2R2B,WNT3 |
| Hepatitis C | 155 | 7.75 | 3 | 0.986 | 1 | CTNNB1,PPP2R2B,PIK3R2 |
| Lysosome | 123 | 6.15 | 2 | 0.987 | 1 | ABCB9,MANBA |
| Wnt signaling pathway | 158 | 7.9 | 3 | 0.987 | 1 | CTNNB1,WNT3,RHOA |
| Huntington's disease | 193 | 9.65 | 4 | 0.989 | 1 | CREB5,POLR2E,NDUFA2,DNAH11 |
| TGF-beta signaling pathway | 92 | 4.6 | 1 | 0.991 | 1 | RHOA |
| Oxidative phosphorylation | 133 | 6.65 | 2 | 0.992 | 1 | PPA2,NDUFA2 |
| GnRH signaling pathway | 93 | 4.65 | 1 | 0.992 | 1 | CACNA1C |
| Amoebiasis | 96 | 4.8 | 1 | 0.993 | 1 | PIK3R2 |

| | | | | | | |
|---|---|---|---|---|---|---|
| Measles | 138 | 6.9 | 2 | 0.993 | 1 | CBLB,PIK3R2 |
| HIF-1 signaling pathway | 100 | 5 | 1 | 0.994 | 1 | PIK3R2 |
| Inflammatory mediator regulation of TRP channels | 100 | 5 | 1 | 0.994 | 1 | PIK3R2 |
| AGE-RAGE signaling pathway in diabetic complications | 100 | 5 | 1 | 0.994 | 1 | PIK3R2 |
| Parkinson's disease | 142 | 7.1 | 2 | 0.994 | 1 | GNAI1,NDUFA2 |
| Toll-like receptor signaling pathway | 104 | 5.2 | 1 | 0.995 | 1 | PIK3R2 |
| Ribosome biogenesis in eukaryotes | 105 | 5.25 | 1 | 0.996 | 1 | EFL1 |
| Herpes simplex infection | 492 | 24.6 | 13 | 0.997 | 1 | HLA-B,HLA-DMA,ZNF398,ZNF823,[illegible] |
| Carbon metabolism | 116 | 5.8 | 1 | 0.998 | 1 | PSPH |
| Hepatitis B | 163 | 8.15 | 2 | 0.998 | 1 | CREB5,PIK3R2 |
| Neuroactive ligand-receptor interaction | 338 | 16.9 | 7 | 0.998 | 1 | GRIN2A,F2,CHRNA5,GABBR1,CH[illegible] |
| Osteoclast differentiation | 128 | 6.4 | 1 | 0.999 | 1 | PIK3R2 |
| NOD-like receptor signaling pathway | 178 | 8.9 | 2 | 0.999 | 1 | TANK,RHOA |
| Spliceosome | 134 | 6.7 | 1 | 0.999 | 1 | PRPF3 |
| Apoptosis | 136 | 6.8 | 1 | 0.999 | 1 | PIK3R2 |
| Ribosome | 153 | 7.65 | 1 | 1 | 1 | MRPL33 |

| | | | | | | |
|---|---|---|---|---|---|---|
| Jak-STAT signaling pathway | 162 | 8.1 | 1 | 1 | 1 | PIK3R2 |
| Pathways in cancer | 530 | 26.5 | 11 | 1 | 1 | GNAI1,ESR2,PTCH1,F2,FGFR1,RA |
| Olfactory transductio n | 448 | 22.4 | 3 | 1 | 1 | SLC8A3,PDE1C,PRKG1 |
| Metabolic pathways | 1430 | 71.6 | 25 | 1 | 1 | MTHFD1L,PDE1C,GALNT12,DGKI,[illegible] |

FADS2,PDE8B,FHIT,PFAS,EXT1,CKB,GATB,NT5C2,GUCY1A2,MGAT5B,IMPA2,ST3GAL3,SRR,DPYD,IVD,I

PSPH,PIP5K1B,PDE4B,GALNT1,GALNT10,NDUFA2
