## Supplemental table 8 for "Digestive Dimensions of Autism: A Multiscale Exploration of Gut-Brain Interactions"

| Pathway | Total | Expected | Hits | P.Value | FDR | Genes |
|---|---|---|---|---|---|---|
| Reduction of cytosolic Ca++ levels | 10 | 0.58 | 3 | 0.0171 | 1 | SLC8A3,ATP2A2,ATP2B2 |
| Signaling by FGFR1 amplification mutants | 1 | 0.058 | 1 | 0.058 | 1 | FGFR1 |
| Translocation of ZAP-70 to Immunological synapse | 18 | 1.04 | 3 | 0.0826 | 1 | HLA-DQA1,HLA-DRB5,HLA-DRB1 |
| Highly calcium permeable nicotinic acetylcholine receptors | 9 | 0.522 | 2 | 0.0921 | 1 | CHRNA5,CHRNA3 |
| Phosphorylation of CD3 and TCR zeta chains | 20 | 1.16 | 3 | 0.106 | 1 | HLA-DQA1,HLA-DRB5,HLA-DRB1 |
| Removal of aminoterminal propeptides from gamma-carboxylated proteins | 10 | 0.58 | 2 | 0.111 | 1 | F2,FURIN |
| Pyrophosphate hydrolysis | 2 | 0.116 | 1 | 0.113 | 1 | PPA2 |
| Amplification of signal from unattached kinetochores via a MAD2 inhibitory signal | 2 | 0.116 | 1 | 0.113 | 1 | MAD1L1 |

| | | | | | | |
|---|---|---|---|---|---|---|
| Amplification of signal from the kinetochores | 2 | 0.116 | 1 | 0.113 | 1 | MAD1L1 |
| Translocation of BoNT Light chain | 2 | 0.116 | 1 | 0.113 | 1 | SYT1 |
| Activation of Na-permeable Kainate Receptors | 2 | 0.116 | 1 | 0.113 | 1 | GRIK1 |
| Pre-NOTCH Expression and Processing | 22 | 1.28 | 3 | 0.132 | 1 | TNRC6A,ST3GAL3,FURIN |
| Platelet calcium homeostasis | 22 | 1.28 | 3 | 0.132 | 1 | SLC8A3,ATP2A2,ATP2B2 |
| Downregulation of SMAD2/3:SMAD4 transcriptional activity | 23 | 1.33 | 3 | 0.145 | 1 | NEDD4L,RNF111,TRIM33 |
| Depolarization of the Presynaptic Terminal Triggers the Opening of Calcium Channels | 12 | 0.696 | 2 | 0.151 | 1 | CACNA2D2,CACNB2 |
| Presynaptic nicotinic acetylcholine receptors | 12 | 0.696 | 2 | 0.151 | 1 | CHRNA5,CHRNA3 |

| | | | | | | |
|---|---|---|---|---|---|---|
| Gamma-carboxylation, transport, and amino-terminal cleavage of proteins | 12 | 0.696 | 2 | 0.151 | 1 | F2,FURIN |
| Pre-NOTCH Processing in Golgi | 12 | 0.696 | 2 | 0.151 | 1 | ST3GAL3,FURIN |
| Processing of DNA double-strand break ends | 3 | 0.174 | 1 | 0.164 | 1 | RPA3 |
| Cam-PDE 1 activation | 3 | 0.174 | 1 | 0.164 | 1 | PDE1C |
| FGFR1c and Klotho ligand binding and activation | 3 | 0.174 | 1 | 0.164 | 1 | FGFR1 |
| NFG and proNGF binds to p75NTR | 3 | 0.174 | 1 | 0.164 | 1 | SORCS3 |
| Serine biosynthesis | 3 | 0.174 | 1 | 0.164 | 1 | PSPH |
| Highly calcium permeable postsynaptic nicotinic acetylcholine receptors | 13 | 0.753 | 2 | 0.172 | 1 | CHRNA5,CHRNA3 |
| PD-1 signaling | 25 | 1.45 | 3 | 0.174 | 1 | HLA-DQA1,HLA-DRB5,HLA-DRB1 |
| Nuclear Receptor transcription pathway | 53 | 3.07 | 5 | 0.191 | 1 | ESR2,ESRRG,NR1D2,ESRRB,THR |

| | | | | | | |
|---|---|---|---|---|---|---|
| NGF processing | 4 | 0.232 | 1 | 0.212 | 1 | FURIN |
| Axonal growth stimulation | 4 | 0.232 | 1 | 0.212 | 1 | RHOA |
| Repair synthesis of patch ~27-30 bases long by DNA polymerase | 15 | 0.869 | 2 | 0.215 | 1 | RPA3,RFC2 |
| Repair synthesis for gap-filling by DNA polymerase in TC-NER | 15 | 0.869 | 2 | 0.215 | 1 | RPA3,RFC2 |
| Beta-catenin phosphorylation cascade | 15 | 0.869 | 2 | 0.215 | 1 | CTNNB1,PPP2R5C |
| Inhibition of Insulin Secretion by Adrenaline/Noradrenaline | 28 | 1.62 | 3 | 0.219 | 1 | GNAI1,CACNA2D2,CACNB2 |
| Activation of Nicotinic Acetylcholine Receptors | 16 | 0.927 | 2 | 0.237 | 1 | CHRNA5,CHRNA3 |
| Gap-filling DNA repair synthesis and ligation in GG-NER | 16 | 0.927 | 2 | 0.237 | 1 | RPA3,RFC2 |
| Gap-filling DNA repair synthesis and ligation in TC-NER | 16 | 0.927 | 2 | 0.237 | 1 | RPA3,RFC2 |

| | | | | | | |
|---|---|---|---|---|---|---|
| Postsynaptic nicotinic acetylcholine receptors | 16 | 0.927 | 2 | 0.237 | 1 | CHRNA5,CHRNA3 |
| Acetylcholine Binding And Downstream Events | 16 | 0.927 | 2 | 0.237 | 1 | CHRNA5,CHRNA3 |
| Regulation of AMPK activity via LKB1 | 16 | 0.927 | 2 | 0.237 | 1 | RPTOR,PRKAG2 |
| Other semaphorin interactions | 16 | 0.927 | 2 | 0.237 | 1 | SEMA3E,SEMA6D |
| Assembly of the RAD51-ssDNA nucleoprotein complex | 5 | 0.29 | 1 | 0.258 | 1 | RPA3 |
| AMPK inhibits chREBP transcriptional activation activity | 5 | 0.29 | 1 | 0.258 | 1 | PRKAG2 |
| Abacavir metabolism | 5 | 0.29 | 1 | 0.258 | 1 | NT5C2 |
| Sodium-coupled sulphate, di- and tri-carboxylate transporters | 5 | 0.29 | 1 | 0.258 | 1 | SLC13A1 |
| Signaling by NODAL | 17 | 0.985 | 2 | 0.259 | 1 | FURIN,FOXO3 |
| Myogenesis | 31 | 1.8 | 3 | 0.267 | 1 | TCF4,MEF2C,CTNNB1 |

| | | | | | | |
|---|---|---|---|---|---|---|
| Generation of second messenger molecules | 31 | 1.8 | 3 | 0.267 | 1 | HLA-DQA1,HLA-DRB5,HLA-DRB1 |
| CDO in myogenesis | 31 | 1.8 | 3 | 0.267 | 1 | TCF4,MEF2C,CTNNB1 |
| Adherens junctions interactions | 31 | 1.8 | 3 | 0.267 | 1 | CADM2,CTNND1,CTNNB1 |
| Homologous DNA pairing and strand exchange | 6 | 0.348 | 1 | 0.301 | 1 | RPA3 |
| Presynaptic phase of homologous DNA pairing and strand exchange | 6 | 0.348 | 1 | 0.301 | 1 | RPA3 |
| Processing of DNA ends prior to end rejoining | 6 | 0.348 | 1 | 0.301 | 1 | XRCC6 |
| Release of eIF4E | 6 | 0.348 | 1 | 0.301 | 1 | RPTOR |
| Klotho-mediated ligand binding | 6 | 0.348 | 1 | 0.301 | 1 | FGFR1 |
| Downstream TCR signaling | 48 | 2.78 | 4 | 0.302 | 1 | HLA-DQA1,HLA-DRB5,HLA-DRB1,F |
| Energy dependent regulation of mTOR by LKB1-AMPK | 19 | 1.1 | 2 | 0.302 | 1 | RPTOR,PRKAG2 |
| Signaling by NOTCH | 95 | 5.51 | 7 | 0.312 | 1 | TNRC6A,HDAC4,ST3GAL3,FURIN,I |
| Lagging Strand Synthesis | 20 | 1.16 | 2 | 0.324 | 1 | RPA3,RFC2 |

| | | | | | | |
|---|---|---|---|---|---|---|
| Highly sodium permeable acetylcholine nicotinic receptors | 7 | 0.406 | 1 | 0.342 | 1 | CHRNA3 |
| Post-transcriptional Silencing By Small RNAs | 7 | 0.406 | 1 | 0.342 | 1 | TNRC6A |
| Constitutive Signaling by NOTCH1 HD+PEST Domain Mutants | 52 | 3.01 | 4 | 0.356 | 1 | HDAC4,MAML3,NEURL1,HDAC9 |
| Costimulation by the CD28 family | 68 | 3.94 | 5 | 0.359 | 1 | HLA-DQA1,HLA-DRB5,HLA-DRB1,[illegible] |
| Telomere C-strand (Lagging Strand) Synthesis | 22 | 1.28 | 2 | 0.367 | 1 | RPA3,RFC2 |
| Nonhomologous End-joining (NHEJ) | 8 | 0.464 | 1 | 0.38 | 1 | XRCC6 |
| Nef and signal transduction | 8 | 0.464 | 1 | 0.38 | 1 | ELMO1 |
| Sema4D mediated inhibition of cell attachment and migration | 8 | 0.464 | 1 | 0.38 | 1 | RHOA |
| Linoleic acid (LA) metabolism | 8 | 0.464 | 1 | 0.38 | 1 | FADS2 |

| | | | | | | |
|---|---|---|---|---|---|---|
| Signaling by TGF-beta Receptor Complex | 70 | 4.06 | 5 | 0.383 | 1 | NEDD4L,FURIN,RNF111,TRIM33,R |
| Platelet homeostasis | 88 | 5.1 | 6 | 0.402 | 1 | SLC8A3,ATP2A2,GUCY1A2,PRKG1 |
| Double-Strand Break Repair | 24 | 1.39 | 2 | 0.409 | 1 | RPA3,XRCC6 |
| Extension of Telomeres | 24 | 1.39 | 2 | 0.409 | 1 | RPA3,RFC2 |
| Nephrin interactions | 24 | 1.39 | 2 | 0.409 | 1 | MAGI2,PIK3R2 |
| Creatine metabolism | 9 | 0.522 | 1 | 0.416 | 1 | CKB |
| Transport of gamma-carboxylated protein precursors from the endoplasmic reticulum to the Golgi apparatus | 9 | 0.522 | 1 | 0.416 | 1 | F2 |
| S6K1-mediated signalling | 9 | 0.522 | 1 | 0.416 | 1 | RPTOR |
| 2-LTR circle formation | 9 | 0.522 | 1 | 0.416 | 1 | XRCC6 |
| Ca2+ activated K+ channels | 9 | 0.522 | 1 | 0.416 | 1 | KCNN2 |
| Axonal growth inhibition (RHOA activation) | 9 | 0.522 | 1 | 0.416 | 1 | RHOA |
| Import of palmitoyl-CoA into the mitochondrial matrix | 9 | 0.522 | 1 | 0.416 | 1 | PRKAG2 |

| | | | | | | |
|---|---|---|---|---|---|---|
| AKT phosphorylates targets in the nucleus | 9 | 0.522 | 1 | 0.416 | 1 | FOXO3 |
| Synthesis, Secretion, and Inactivation of Glucagon-like Peptide-1 (GLP-1) | 9 | 0.522 | 1 | 0.416 | 1 | DPP4 |
| Sodium/Calcium exchangers | 9 | 0.522 | 1 | 0.416 | 1 | SLC8A3 |
| Synthesis, Secretion, and Inactivation of Glucose-dependent Insulinotropic Polypeptide (GIP) | 9 | 0.522 | 1 | 0.416 | 1 | DPP4 |
| Regulation of cytoskeletal remodeling and cell spreading by IPP complex components | 9 | 0.522 | 1 | 0.416 | 1 | RSU1 |
| Smooth Muscle Contraction | 25 | 1.45 | 2 | 0.43 | 1 | CALD1,SORBS1 |

| | | | | | | |
|---|---|---|---|---|---|---|
| Constitutive Signaling by NOTCH1 PEST Domain Mutants | 59 | 3.42 | 4 | 0.449 | 1 | HDAC4,MAML3,NEURL1,HDAC9 |
| Gamma-carboxylation of protein precursors | 10 | 0.58 | 1 | 0.45 | 1 | F2 |
| Removal of the Flap Intermediate from the C-strand | 10 | 0.58 | 1 | 0.45 | 1 | RPA3 |
| Abacavir transport and metabolism | 10 | 0.58 | 1 | 0.45 | 1 | NT5C2 |
| p75NTR regulates axonogenesis | 10 | 0.58 | 1 | 0.45 | 1 | RHOA |
| COPI Mediated Transport | 10 | 0.58 | 1 | 0.45 | 1 | GBF1 |
| Golgi to ER Retrograde Transport | 10 | 0.58 | 1 | 0.45 | 1 | GBF1 |
| Endosomal/Vacuolar pathway | 10 | 0.58 | 1 | 0.45 | 1 | HLA-B |
| Acetylcholine Neurotransmitter Release Cycle | 10 | 0.58 | 1 | 0.45 | 1 | SYT1 |
| Regulation of Rheb GTPase activity by AMPK | 10 | 0.58 | 1 | 0.45 | 1 | PRKAG2 |

| | | | | | | |
|---|---|---|---|---|---|---|
| Incretin Synthesis, Secretion, and Inactivation | 10 | 0.58 | 1 | 0.45 | 1 | DPP4 |
| Zinc influx into cells by the SLC39 gene family | 10 | 0.58 | 1 | 0.45 | 1 | SLC39A8 |
| Clearance of Nuclear Envelope Membranes from Chromatin | 10 | 0.58 | 1 | 0.45 | 1 | VRK2 |
| Cohesin Loading onto Chromatin | 10 | 0.58 | 1 | 0.45 | 1 | STAG1 |
| PI3K Cascade | 76 | 4.4 | 5 | 0.452 | 1 | RPTOR,FGFR1,PRKAG2,TWF2,PIK |
| Transcriptional activity of SMAD2/SMAD3:SMAD4 heterotrimer | 43 | 2.49 | 3 | 0.458 | 1 | NEDD4L,RNF111,TRIM33 |
| Mitotic Spindle Checkpoint | 27 | 1.56 | 2 | 0.47 | 1 | ANAPC4,MAD1L1 |
| Nitric oxide stimulates guanylate cyclase | 27 | 1.56 | 2 | 0.47 | 1 | GUCY1A2,PRKG1 |
| Purine ribonucleoside monophosphate biosynthesis | 11 | 0.638 | 1 | 0.482 | 1 | PFAS |
| Purine catabolism | 11 | 0.638 | 1 | 0.482 | 1 | NT5C2 |
| Common Pathway | 11 | 0.638 | 1 | 0.482 | 1 | F2 |

| | | | | | | |
|---|---|---|---|---|---|---|
| Processive synthesis on the C-strand of the telomere | 11 | 0.638 | 1 | 0.482 | 1 | RPA3 |
| mTORC1-mediated signalling | 11 | 0.638 | 1 | 0.482 | 1 | RPTOR |
| FGFR1c ligand binding and activation | 11 | 0.638 | 1 | 0.482 | 1 | FGFR1 |
| Serotonin Neurotransmitter Release Cycle | 11 | 0.638 | 1 | 0.482 | 1 | SYT1 |
| Norepinephrine Neurotransmitter Release Cycle | 11 | 0.638 | 1 | 0.482 | 1 | SYT1 |
| Dopamine Neurotransmitter Release Cycle | 11 | 0.638 | 1 | 0.482 | 1 | SYT1 |
| Synthesis of IP2, IP, and Ins in the cytosol | 11 | 0.638 | 1 | 0.482 | 1 | IMPA2 |
| Signaling by activated point mutants of FGFR1 | 11 | 0.638 | 1 | 0.482 | 1 | FGFR1 |
| Establishment of Sister Chromatid Cohesion | 11 | 0.638 | 1 | 0.482 | 1 | STAG1 |
| Regulatory RNA pathways | 28 | 1.62 | 2 | 0.489 | 1 | TNRC6A,POLR2E |
| mTOR signalling | 29 | 1.68 | 2 | 0.508 | 1 | RPTOR,PRKAG2 |

| | | | | | | |
|---|---|---|---|---|---|---|
| PTM: gamma carboxylation, hypusine formation and arylsulfatase activation | 29 | 1.68 | 2 | 0.508 | 1 | F2,FURIN |
| Association of TriC/CCT with target proteins during biosynthesis | 29 | 1.68 | 2 | 0.508 | 1 | FKBP9,ARFGEF2 |
| Pyrimidine catabolism | 12 | 0.696 | 1 | 0.512 | 1 | DPYD |
| Interleukin-7 signaling | 12 | 0.696 | 1 | 0.512 | 1 | PIK3R2 |
| Pre-NOTCH Transcription and Translation | 12 | 0.696 | 1 | 0.512 | 1 | TNRC6A |
| Bile salt and organic anion SLC transporters | 12 | 0.696 | 1 | 0.512 | 1 | SLC13A1 |
| Ionotropic activity of Kainate Receptors | 12 | 0.696 | 1 | 0.512 | 1 | GRIK1 |
| Activation of Ca-permeable Kainate Receptor | 12 | 0.696 | 1 | 0.512 | 1 | GRIK1 |
| Transcription-coupled NER (TC-NER) | 47 | 2.72 | 3 | 0.518 | 1 | RPA3,RFC2,POLR2E |
| Regulation of Insulin Secretion | 82 | 4.75 | 5 | 0.52 | 1 | GNAI1,CACNA2D2,KCNG2,CACNB |
| TCR signaling | 65 | 3.77 | 4 | 0.525 | 1 | HLA-DQA1,HLA-DRB5,HLA-DRB1,[illegible] |

| | | | | | | |
|---|---|---|---|---|---|---|
| PKB-mediated events | 30 | 1.74 | 2 | 0.526 | 1 | RPTOR,PRKAG2 |
| TGF-beta receptor signaling activates SMADs | 30 | 1.74 | 2 | 0.526 | 1 | NEDD4L,FURIN |
| O-linked glycosylation of mucins | 66 | 3.83 | 4 | 0.538 | 1 | GALNT12,ST3GAL3,GALNT1,GALN |
| Adenylate cyclase inhibitory pathway | 13 | 0.753 | 1 | 0.54 | 1 | GNAI1 |
| Caspase-mediated cleavage of cytoskeletal proteins | 13 | 0.753 | 1 | 0.54 | 1 | MAPT |
| Apoptotic cleavage of cell adhesion proteins | 13 | 0.753 | 1 | 0.54 | 1 | CTNNB1 |
| alpha-linolenic acid (ALA) metabolism | 13 | 0.753 | 1 | 0.54 | 1 | FADS2 |
| alpha-linolenic (omega3) and linoleic (omega6) acid metabolism | 13 | 0.753 | 1 | 0.54 | 1 | FADS2 |
| Inhibition of adenylate cyclase pathway | 13 | 0.753 | 1 | 0.54 | 1 | GNAI1 |
| DNA strand elongation | 31 | 1.8 | 2 | 0.544 | 1 | RPA3,RFC2 |
| Activation of Matrix Metalloproteinases | 31 | 1.8 | 2 | 0.544 | 1 | MMP16,FURIN |

| | | | | | | |
|---|---|---|---|---|---|---|
| Signaling by FGFR1 mutants | 31 | 1.8 | 2 | 0.544 | 1 | MYO18A,FGFR1 |
| NOTCH1 Intracellular Domain Regulates Transcription | 50 | 2.9 | 3 | 0.561 | 1 | HDAC4,MAML3,HDAC9 |
| Removal of the Flap Intermediate | 14 | 0.811 | 1 | 0.567 | 1 | RPA3 |
| Leading Strand Synthesis | 14 | 0.811 | 1 | 0.567 | 1 | RFC2 |
| Polymerase switching | 14 | 0.811 | 1 | 0.567 | 1 | RFC2 |
| Mitotic Telophase/Cytokinesis | 14 | 0.811 | 1 | 0.567 | 1 | STAG1 |
| Polymerase switching on the C-strand of the telomere | 14 | 0.811 | 1 | 0.567 | 1 | RFC2 |
| FGFR1 ligand binding and activation | 14 | 0.811 | 1 | 0.567 | 1 | FGFR1 |
| Glutamate Neurotransmitter Release Cycle | 14 | 0.811 | 1 | 0.567 | 1 | SYT1 |
| Constitutive Signaling by NOTCH1 HD Domain Mutants | 14 | 0.811 | 1 | 0.567 | 1 | NEURL1 |

| | | | | | | |
|---|---|---|---|---|---|---|
| SEMA3A-Plexin repulsion signaling by inhibiting Integrin adhesion | 14 | 0.811 | 1 | 0.567 | 1 | SEMA3A |
| Nuclear Envelope Reassembly | 14 | 0.811 | 1 | 0.567 | 1 | VRK2 |
| Initiation of Nuclear Envelope Reformation | 14 | 0.811 | 1 | 0.567 | 1 | VRK2 |
| IRS-mediated signalling | 87 | 5.04 | 5 | 0.574 | 1 | RPTOR,FGFR1,PRKAG2,TWF2,PIK |
| Rho GTPase cycle | 123 | 7.13 | 7 | 0.576 | 1 | ARHGAP15,MYO9B,GMIP,RHOJ,N( |
| Signaling by Rho GTPases | 123 | 7.13 | 7 | 0.576 | 1 | ARHGAP15,MYO9B,GMIP,RHOJ,N( |
| Synthesis of PIPs at the plasma membrane | 33 | 1.91 | 2 | 0.578 | 1 | PIP5K1B,PIK3R2 |
| GPVI-mediated activation cascade | 33 | 1.91 | 2 | 0.578 | 1 | RHOA,PIK3R2 |
| Processive synthesis on the lagging strand | 15 | 0.869 | 1 | 0.592 | 1 | RPA3 |
| NICD traffics to nucleus | 15 | 0.869 | 1 | 0.592 | 1 | MAML3 |
| Notch-HLH transcription pathway | 15 | 0.869 | 1 | 0.592 | 1 | MAML3 |
| Regulation of Complement cascade | 15 | 0.869 | 1 | 0.592 | 1 | CD55 |

| | | | | | | |
|---|---|---|---|---|---|---|
| Purine metabolism | 34 | 1.97 | 2 | 0.595 | 1 | PFAS,NT5C2 |
| Nucleotide Excision Repair | 53 | 3.07 | 3 | 0.601 | 1 | RPA3,RFC2,POLR2E |
| IRS-related events | 90 | 5.22 | 5 | 0.604 | 1 | RPTOR,FGFR1,PRKAG2,TWF2,PIK |
| Semaphorin interactions | 72 | 4.17 | 4 | 0.607 | 1 | SEMA3E,SEMA3A,SEMA6D,RHOA |
| Integration of energy metabolism | 109 | 6.32 | 6 | 0.611 | 1 | GNAI1,CACNA2D2,KCNG2,PRKAG |
| IRS-related events triggered by IGF1R | 91 | 5.27 | 5 | 0.614 | 1 | RPTOR,FGFR1,PRKAG2,TWF2,PIK |
| Homologous recombination repair of replication-independent double-strand breaks | 16 | 0.927 | 1 | 0.616 | 1 | RPA3 |
| Homologous Recombination Repair | 16 | 0.927 | 1 | 0.616 | 1 | RPA3 |
| Class C/3 (Metabotropic glutamate/pheromone receptors) | 16 | 0.927 | 1 | 0.616 | 1 | GABBR1 |
| CRMPs in Sema3A signaling | 16 | 0.927 | 1 | 0.616 | 1 | SEMA3A |
| Sema3A PAK dependent Axon repulsion | 16 | 0.927 | 1 | 0.616 | 1 | SEMA3A |

| | | | | | | |
|---|---|---|---|---|---|---|
| Global Genomic NER (GG-NER) | 36 | 2.09 | 2 | 0.626 | 1 | RPA3,RFC2 |
| Signaling by NOTCH1 t(7;9)(NOTCH1:M1580_K2555) Translocation Mutant | 74 | 4.29 | 4 | 0.629 | 1 | HDAC4,MAML3,NEURL1,HDAC9 |
| Signaling by NOTCH1 in Cancer | 74 | 4.29 | 4 | 0.629 | 1 | HDAC4,MAML3,NEURL1,HDAC9 |
| Signaling by NOTCH1 PEST Domain Mutants in Cancer | 74 | 4.29 | 4 | 0.629 | 1 | HDAC4,MAML3,NEURL1,HDAC9 |
| FBXW7 Mutants and NOTCH1 in Cancer | 74 | 4.29 | 4 | 0.629 | 1 | HDAC4,MAML3,NEURL1,HDAC9 |
| Signaling by NOTCH1 HD Domain Mutants in Cancer | 74 | 4.29 | 4 | 0.629 | 1 | HDAC4,MAML3,NEURL1,HDAC9 |
| Interferon gamma signaling | 74 | 4.29 | 4 | 0.629 | 1 | HLA-B,HLA-DQA1,HLA-DRB5,HLA-[illegible] |
| Signaling by NOTCH1 HD+PEST Domain Mutants in Cancer | 74 | 4.29 | 4 | 0.629 | 1 | HDAC4,MAML3,NEURL1,HDAC9 |
| Signaling by NOTCH1 | 74 | 4.29 | 4 | 0.629 | 1 | HDAC4,MAML3,NEURL1,HDAC9 |
| Trafficking and processing of endosomal TLR | 17 | 0.985 | 1 | 0.638 | 1 | TWF2 |

| | | | | | | |
|---|---|---|---|---|---|---|
| Intrinsic Pathway | 17 | 0.985 | 1 | 0.638 | 1 | F2 |
| Integration of provirus | 17 | 0.985 | 1 | 0.638 | 1 | XRCC6 |
| Synthesis of glycosylphosphatidylinositol (GPI) | 17 | 0.985 | 1 | 0.638 | 1 | SEMA6D |
| TGF-beta receptor signaling in EMT (epithelial to mesenchymal transition) | 17 | 0.985 | 1 | 0.638 | 1 | RHOA |
| Rap1 signalling | 17 | 0.985 | 1 | 0.638 | 1 | PRKG1 |
| TRAF6 mediated IRF7 activation in TLR7/8 or 9 signaling | 17 | 0.985 | 1 | 0.638 | 1 | TWF2 |
| CREB phosphorylation through the activation of CaMKII | 17 | 0.985 | 1 | 0.638 | 1 | GRIN2A |
| Zinc transporters | 17 | 0.985 | 1 | 0.638 | 1 | SLC39A8 |
| Unblocking of NMDA receptor, glutamate binding and activation | 17 | 0.985 | 1 | 0.638 | 1 | GRIN2A |
| Nuclear Envelope Breakdown | 17 | 0.985 | 1 | 0.638 | 1 | VRK2 |

| | | | | | | |
|---|---|---|---|---|---|---|
| Signaling by Type 1 Insulin-like Growth Factor 1 Receptor (IGF1R) | 94 | 5.45 | 5 | 0.643 | 1 | RPTOR,FGFR1,PRKAG2,TWF2,PIK |
| IGF1R signaling cascade | 94 | 5.45 | 5 | 0.643 | 1 | RPTOR,FGFR1,PRKAG2,TWF2,PIK |
| Insulin receptor signalling cascade | 95 | 5.51 | 5 | 0.652 | 1 | RPTOR,FGFR1,PRKAG2,TWF2,PIK |
| Amino acid synthesis and interconversion (transamination) | 18 | 1.04 | 1 | 0.659 | 1 | PSPH |
| Branched-chain amino acid catabolism | 18 | 1.04 | 1 | 0.659 | 1 | IVD |
| RNA Polymerase III Chain Elongation | 18 | 1.04 | 1 | 0.659 | 1 | POLR2E |
| RNA Polymerase III Transcription Termination | 18 | 1.04 | 1 | 0.659 | 1 | POLR2E |
| GABA synthesis, release, reuptake and degradation | 18 | 1.04 | 1 | 0.659 | 1 | SYT1 |
| Signaling by NOTCH2 | 18 | 1.04 | 1 | 0.659 | 1 | NEURL1 |

| | | | | | | |
|---|---|---|---|---|---|---|
| NOTCH2 Activation and Transmission of Signal to the Nucleus | 18 | 1.04 | 1 | 0.659 | 1 | NEURL1 |
| NCAM1 interactions | 39 | 2.26 | 2 | 0.67 | 1 | CACNB2,CACNA1I |
| Botulinum neurotoxicity | 19 | 1.1 | 1 | 0.679 | 1 | SYT1 |
| CD28 dependent PI3K/Akt signaling | 19 | 1.1 | 1 | 0.679 | 1 | PIK3R2 |
| Na+/Cl- dependent neurotransmitter transporters | 19 | 1.1 | 1 | 0.679 | 1 | SLC6A9 |
| Ras activation uopn Ca2+ infux through NMDA receptor | 19 | 1.1 | 1 | 0.679 | 1 | GRIN2A |
| Cell-extracellular matrix interactions | 19 | 1.1 | 1 | 0.679 | 1 | RSU1 |
| Platelet sensitization by LDL | 19 | 1.1 | 1 | 0.679 | 1 | PPP2R5C |
| GABA B receptor activation | 40 | 2.32 | 2 | 0.683 | 1 | GNAI1,GABBR1 |
| Activation of GABAB receptors | 40 | 2.32 | 2 | 0.683 | 1 | GNAI1,GABBR1 |
| Cell-cell junction organization | 60 | 3.48 | 3 | 0.684 | 1 | CADM2,CTNND1,CTNNB1 |

| | | | | | | |
|---|---|---|---|---|---|---|
| MHC class II antigen presentation | 118 | 6.84 | 6 | 0.688 | 1 | HLA-DMA,DYNC1I1,KLC1,HLA-DQ/ |
| Activation of ATR in response to replication stress | 41 | 2.38 | 2 | 0.696 | 1 | RPA3,RFC2 |
| Early Phase of HIV Life Cycle | 20 | 1.16 | 1 | 0.698 | 1 | XRCC6 |
| Tie2 Signaling | 20 | 1.16 | 1 | 0.698 | 1 | PIK3R2 |
| Signaling by FGFR1 fusion mutants | 20 | 1.16 | 1 | 0.698 | 1 | MYO18A |
| Ion transport by P-type ATPases | 42 | 2.43 | 2 | 0.709 | 1 | ATP2A2,ATP2B2 |
| RNA Polymerase I Chain Elongation | 21 | 1.22 | 1 | 0.715 | 1 | TAF1C |
| ERK/MAPK targets | 21 | 1.22 | 1 | 0.715 | 1 | MEF2C |
| Regulation of Insulin-like Growth Factor (IGF) Transport and Uptake by Insulin-like Growth Factor Binding Proteins (IGFBPs) | 21 | 1.22 | 1 | 0.715 | 1 | F2 |
| Mitochondrial tRNA aminoacylation | 21 | 1.22 | 1 | 0.715 | 1 | PPA2 |
| cGMP effects | 21 | 1.22 | 1 | 0.715 | 1 | PRKG1 |

| | | | | | | |
|---|---|---|---|---|---|---|
| Apoptotic cleavage of cellular proteins | 43 | 2.49 | 2 | 0.721 | 1 | MAPT,CTNNB1 |
| Voltage gated Potassium channels | 43 | 2.49 | 2 | 0.721 | 1 | KCNG2,KCNB1 |
| RNA Polymerase I Promoter Escape | 22 | 1.28 | 1 | 0.732 | 1 | TAF1C |
| FGFR ligand binding and activation | 22 | 1.28 | 1 | 0.732 | 1 | FGFR1 |
| Regulation of signaling by CBL | 22 | 1.28 | 1 | 0.732 | 1 | PIK3R2 |
| Degradation of beta-catenin by the destruction complex | 65 | 3.77 | 3 | 0.736 | 1 | PSMA5,CTNNB1,PPP2R5C |
| Signaling by Wnt | 65 | 3.77 | 3 | 0.736 | 1 | PSMA5,CTNNB1,PPP2R5C |
| NCAM signaling for neurite out-growth | 65 | 3.77 | 3 | 0.736 | 1 | FGFR1,CACNB2,CACNA1I |
| PI3K/AKT activation | 106 | 6.14 | 5 | 0.744 | 1 | TNRC6A,FGFR1,FOXO3,RHOA,PIK |
| Signaling by FGFR mutants | 45 | 2.61 | 2 | 0.744 | 1 | MYO18A,FGFR1 |
| RNA Polymerase I Transcription Initiation | 23 | 1.33 | 1 | 0.747 | 1 | TAF1C |
| Dual incision reaction in GG-NER | 23 | 1.33 | 1 | 0.747 | 1 | RPA3 |

| | | | | | | |
|---|---|---|---|---|---|---|
| Formation of incision complex in GG-NER | 23 | 1.33 | 1 | 0.747 | 1 | RPA3 |
| RNA Polymerase I Transcription Termination | 23 | 1.33 | 1 | 0.747 | 1 | TAF1C |
| Abortive elongation of HIV-1 transcript in the absence of Tat | 23 | 1.33 | 1 | 0.747 | 1 | POLR2E |
| ADP signalling through P2Y purinoceptor 12 | 23 | 1.33 | 1 | 0.747 | 1 | GNAI1 |
| PLC beta mediated events | 46 | 2.67 | 2 | 0.755 | 1 | PDE1C,GNAI1 |
| RNA Polymerase I Promoter Clearance | 24 | 1.39 | 1 | 0.762 | 1 | TAF1C |
| Pyrimidine metabolism | 24 | 1.39 | 1 | 0.762 | 1 | DPYD |
| Nuclear Events (kinase and transcription factor activation) | 24 | 1.39 | 1 | 0.762 | 1 | MEF2C |
| Insulin Processing | 24 | 1.39 | 1 | 0.762 | 1 | EXOC4 |
| CTLA4 inhibitory signaling | 24 | 1.39 | 1 | 0.762 | 1 | PPP2R5C |
| G-protein mediated events | 47 | 2.72 | 2 | 0.766 | 1 | PDE1C,GNAI1 |

| | | | | | | |
|---|---|---|---|---|---|---|
| Regulation of Insulin Secretion by Glucagon-like Peptide-1 | 47 | 2.72 | 2 | 0.766 | 1 | KCNG2,KCNB1 |
| Cell junction organization | 89 | 5.16 | 4 | 0.767 | 1 | RSU1,CADM2,CTNND1,CTNNB1 |
| RNA Polymerase III Transcription Initiation From Type 1 Promoter | 25 | 1.45 | 1 | 0.776 | 1 | POLR2E |
| RNA Polymerase III Transcription Initiation From Type 2 Promoter | 25 | 1.45 | 1 | 0.776 | 1 | POLR2E |
| Phosphorylation of the APC/C | 25 | 1.45 | 1 | 0.776 | 1 | ANAPC4 |
| Conversion from APC/C:Cdc20 to APC/C:Cdh1 in late anaphase | 25 | 1.45 | 1 | 0.776 | 1 | ANAPC4 |
| MicroRNA (miRNA) Biogenesis | 25 | 1.45 | 1 | 0.776 | 1 | POLR2E |
| Metal ion SLC transporters | 25 | 1.45 | 1 | 0.776 | 1 | SLC39A8 |

| | | | | | | |
|---|---|---|---|---|---|---|
| Antigen Presentation: Folding, assembly and peptide loading of class I MHC | 25 | 1.45 | 1 | 0.776 | 1 | HLA-B |
| G2/M Checkpoints | 48 | 2.78 | 2 | 0.776 | 1 | RPA3,RFC2 |
| RNA Polymerase I Transcription | 26 | 1.51 | 1 | 0.789 | 1 | TAF1C |
| Inactivation of APC/C via direct inhibition of the APC/C complex | 26 | 1.51 | 1 | 0.789 | 1 | ANAPC4 |
| Inhibition of the proteolytic activity of APC/C required for the onset of anaphase by mitotic spindle checkpoint components | 26 | 1.51 | 1 | 0.789 | 1 | ANAPC4 |
| Post-translational modification: synthesis of GPI-anchored proteins | 26 | 1.51 | 1 | 0.789 | 1 | SEMA6D |
| PI Metabolism | 50 | 2.9 | 2 | 0.795 | 1 | PIP5K1B,PIK3R2 |

| | | | | | | |
|---|---|---|---|---|---|---|
| RNA Polymerase III Transcription Initiation From Type 3 Promoter | 27 | 1.56 | 1 | 0.801 | 1 | POLR2E |
| Activation of G protein gated Potassium channels | 27 | 1.56 | 1 | 0.801 | 1 | GABBR1 |
| G protein gated Potassium channels | 27 | 1.56 | 1 | 0.801 | 1 | GABBR1 |
| Downregulation of TGF-beta receptor signaling | 27 | 1.56 | 1 | 0.801 | 1 | NEDD4L |
| G beta:gamma signalling through PI3Kgamma | 27 | 1.56 | 1 | 0.801 | 1 | RHOA |
| G0 and Early G1 | 27 | 1.56 | 1 | 0.801 | 1 | RBL2 |
| Keratan sulfate biosynthesis | 27 | 1.56 | 1 | 0.801 | 1 | ST3GAL3 |
| Role of phospholipids in phagocytosis | 27 | 1.56 | 1 | 0.801 | 1 | PIK3R2 |
| Inhibition of voltage gated Ca2+ channels via Gbeta/gamma subunits | 27 | 1.56 | 1 | 0.801 | 1 | GABBR1 |

| | | | | | | |
|---|---|---|---|---|---|---|
| Chaperonin-mediated protein folding | 51 | 2.96 | 2 | 0.804 | 1 | FKBP9,ARFGEF2 |
| Transmission across Chemical Synapses | 196 | 11.4 | 9 | 0.81 | 1 | GNAI1,GRIN2A,CHRNA5,CACNA2D |
| RNA Pol II CTD phosphorylation and interaction with CE | 28 | 1.62 | 1 | 0.813 | 1 | POLR2E |
| Effects of PIP2 hydrolysis | 28 | 1.62 | 1 | 0.813 | 1 | DGKI |
| RNA Pol II CTD phosphorylation and interaction with CE | 28 | 1.62 | 1 | 0.813 | 1 | POLR2E |
| SHC-mediated cascade | 28 | 1.62 | 1 | 0.813 | 1 | FGFR1 |
| SMAD2/SMAD3:SMAD4 heterotrimer regulates transcription | 28 | 1.62 | 1 | 0.813 | 1 | RNF111 |
| Interleukin receptor SHC signaling | 28 | 1.62 | 1 | 0.813 | 1 | PIK3R2 |
| Fanconi Anemia pathway | 28 | 1.62 | 1 | 0.813 | 1 | PALB2 |
| Termination of O-glycan biosynthesis | 28 | 1.62 | 1 | 0.813 | 1 | ST3GAL3 |
| TRAF6 mediated IRF7 activation | 28 | 1.62 | 1 | 0.813 | 1 | TANK |
| Muscle contraction | 52 | 3.01 | 2 | 0.813 | 1 | CALD1,SORBS1 |

| | | | | | | |
|---|---|---|---|---|---|---|
| DNA Repair | 117 | 6.78 | 5 | 0.817 | 1 | RPA3,XRCC6,PALB2,RFC2,POLR2 |
| Signaling by Insulin receptor | 117 | 6.78 | 5 | 0.817 | 1 | RPTOR,FGFR1,PRKAG2,TWF2,PIK |
| Formation of Fibrin Clot (Clotting Cascade) | 29 | 1.68 | 1 | 0.824 | 1 | F2 |
| Activated NOTCH1 Transmits Signal to the Nucleus | 29 | 1.68 | 1 | 0.824 | 1 | NEURL1 |
| APC/C:Cdc20 mediated degradation of Cyclin B | 29 | 1.68 | 1 | 0.824 | 1 | ANAPC4 |
| DARPP-32 events | 29 | 1.68 | 1 | 0.824 | 1 | PDE4B |
| G-protein activation | 29 | 1.68 | 1 | 0.824 | 1 | GNAI1 |
| Sema4D induced cell migration and growth-cone collapse | 29 | 1.68 | 1 | 0.824 | 1 | RHOA |
| CREB phosphorylation through the activation of Ras | 29 | 1.68 | 1 | 0.824 | 1 | GRIN2A |
| Interaction between L1 and Ankyrins | 29 | 1.68 | 1 | 0.824 | 1 | ANK3 |
| Degradation of the extracellular matrix | 77 | 4.46 | 3 | 0.832 | 1 | COL11A1,MMP16,FURIN |
| mRNA Capping | 30 | 1.74 | 1 | 0.834 | 1 | POLR2E |
| Calmodulin induced events | 30 | 1.74 | 1 | 0.834 | 1 | PDE1C |

| | | | | | | |
|---|---|---|---|---|---|---|
| CaM pathway | 30 | 1.74 | 1 | 0.834 | 1 | PDE1C |
| CD28 co-stimulation | 30 | 1.74 | 1 | 0.834 | 1 | PIK3R2 |
| Amine compound SLC transporters | 30 | 1.74 | 1 | 0.834 | 1 | SLC6A9 |
| G-protein beta:gamma signalling | 30 | 1.74 | 1 | 0.834 | 1 | RHOA |
| MAPK targets/ Nuclear events mediated by MAP kinases | 30 | 1.74 | 1 | 0.834 | 1 | MEF2C |
| GABA receptor activation | 55 | 3.19 | 2 | 0.837 | 1 | GNAI1,GABBR1 |
| Dual incision reaction in TC-NER | 31 | 1.8 | 1 | 0.844 | 1 | POLR2E |
| Formation of transcription-coupled NER (TC-NER) repair complex | 31 | 1.8 | 1 | 0.844 | 1 | POLR2E |
| APC-Cdc20 mediated degradation of Nek2A | 31 | 1.8 | 1 | 0.844 | 1 | ANAPC4 |
| The role of Nef in HIV-1 replication and disease pathogenesis | 31 | 1.8 | 1 | 0.844 | 1 | ELMO1 |
| HS-GAG biosynthesis | 31 | 1.8 | 1 | 0.844 | 1 | EXT1 |

| | | | | | | |
|---|---|---|---|---|---|---|
| Transcriptional Regulation of White Adipocyte Differentiation | 56 | 3.25 | 2 | 0.844 | 1 | MED27,MED8 |
| Protein folding | 56 | 3.25 | 2 | 0.844 | 1 | FKBP9,ARFGEF2 |
| Neurotransmitter Receptor Binding And Downstream Transmission In The Postsynaptic Cell | 143 | 8.29 | 6 | 0.845 | 1 | GNAI1,GRIN2A,CHRNA5,GABBR1,[illegible] |
| Cell-Cell communication | 143 | 8.29 | 6 | 0.845 | 1 | RSU1,MAGI2,CADM2,CTNND1,CT[illegible] |
| G alpha (12/13) signalling events | 80 | 4.64 | 3 | 0.851 | 1 | NGEF,RHOA,PIK3R2 |
| Apoptotic execution phase | 57 | 3.3 | 2 | 0.851 | 1 | MAPT,CTNNB1 |
| Phospholipase C-mediated cascade | 57 | 3.3 | 2 | 0.851 | 1 | PDE1C,FGFR1 |
| Sphingolipid de novo biosynthesis | 32 | 1.85 | 1 | 0.853 | 1 | PRKD1 |
| Activation of the pre-replicative complex | 32 | 1.85 | 1 | 0.853 | 1 | RPA3 |
| Keratan sulfate/keratin metabolism | 32 | 1.85 | 1 | 0.853 | 1 | ST3GAL3 |
| Ca-dependent events | 32 | 1.85 | 1 | 0.853 | 1 | PDE1C |

| | | | | | | |
|---|---|---|---|---|---|---|
| Pausing and recovery of Tat-mediated HIV-1 elongation | 32 | 1.85 | 1 | 0.853 | 1 | POLR2E |
| Tat-mediated HIV-1 elongation arrest and recovery | 32 | 1.85 | 1 | 0.853 | 1 | POLR2E |
| Biosynthesis of the N-glycan precursor (dolichol lipid-linked oligosaccharide, LLO) and transfer to a nascent protein | 32 | 1.85 | 1 | 0.853 | 1 | RFT1 |
| Antigen Activates B Cell Receptor Leading to Generation of Second Messengers | 32 | 1.85 | 1 | 0.853 | 1 | CBLB |
| PI3K events in ERBB4 signaling | 103 | 5.97 | 4 | 0.856 | 1 | TNRC6A,FGFR1,FOXO3,PIK3R2 |
| PIP3 activates AKT signaling | 103 | 5.97 | 4 | 0.856 | 1 | TNRC6A,FGFR1,FOXO3,PIK3R2 |
| PI-3K cascade | 103 | 5.97 | 4 | 0.856 | 1 | TNRC6A,FGFR1,FOXO3,PIK3R2 |
| Potassium Channels | 103 | 5.97 | 4 | 0.856 | 1 | KCNG2,GABBR1,KCNB1,KCNN2 |
| PI3K/AKT Signaling in Cancer | 103 | 5.97 | 4 | 0.856 | 1 | TNRC6A,FGFR1,FOXO3,PIK3R2 |

| | | | | | | |
|---|---|---|---|---|---|---|
| PI3K events in ERBB2 signaling | 103 | 5.97 | 4 | 0.856 | 1 | TNRC6A,FGFR1,FOXO3,PIK3R2 |
| Metabolism of nucleotides | 81 | 4.69 | 3 | 0.857 | 1 | PFAS,NT5C2,DPYD |
| RNA Polymerase III Transcription | 33 | 1.91 | 1 | 0.861 | 1 | POLR2E |
| RNA Polymerase III Transcription Initiation | 33 | 1.91 | 1 | 0.861 | 1 | POLR2E |
| Elongation arrest and recovery | 33 | 1.91 | 1 | 0.861 | 1 | POLR2E |
| HIV-1 elongation arrest and recovery | 33 | 1.91 | 1 | 0.861 | 1 | POLR2E |
| Pausing and recovery of HIV-1 elongation | 33 | 1.91 | 1 | 0.861 | 1 | POLR2E |
| Complement cascade | 33 | 1.91 | 1 | 0.861 | 1 | CD55 |
| Inwardly rectifying K+ channels | 33 | 1.91 | 1 | 0.861 | 1 | GABBR1 |
| RNA Polymerase III Abortive And Retractive Initiation | 33 | 1.91 | 1 | 0.861 | 1 | POLR2E |
| Signal amplification | 33 | 1.91 | 1 | 0.861 | 1 | GNAI1 |

| | | | | | | |
|---|---|---|---|---|---|---|
| RNA Polymerase I, RNA Polymerase III, and Mitochondrial Transcription | 59 | 3.42 | 2 | 0.864 | 1 | TAF1C,POLR2E |
| Formation of the Early Elongation Complex | 34 | 1.97 | 1 | 0.869 | 1 | POLR2E |
| Formation of the HIV-1 Early Elongation Complex | 34 | 1.97 | 1 | 0.869 | 1 | POLR2E |
| Sema4D in semaphorin signaling | 34 | 1.97 | 1 | 0.869 | 1 | RHOA |
| Activation of Kainate Receptors upon glutamate binding | 34 | 1.97 | 1 | 0.869 | 1 | GRIK1 |
| Thrombin signalling through proteinase activated receptors (PARs) | 34 | 1.97 | 1 | 0.869 | 1 | F2 |
| Peptide hormone metabolism | 60 | 3.48 | 2 | 0.871 | 1 | EXOC4,DPP4 |
| GAB1 signalosome | 106 | 6.14 | 4 | 0.871 | 1 | TNRC6A,FGFR1,FOXO3,PIK3R2 |
| Degradation of collagen | 61 | 3.54 | 2 | 0.877 | 1 | COL11A1,FURIN |
| Neurotransmitter Release Cycle | 35 | 2.03 | 1 | 0.877 | 1 | SYT1 |
| Signal transduction by L1 | 35 | 2.03 | 1 | 0.877 | 1 | FGFR1 |

| | | | | | | |
|---|---|---|---|---|---|---|
| Post NMDA receptor activation events | 35 | 2.03 | 1 | 0.877 | 1 | GRIN2A |
| DAG and IP3 signaling | 35 | 2.03 | 1 | 0.877 | 1 | PDE1C |
| p75 NTR receptor-mediated signalling | 85 | 4.93 | 3 | 0.878 | 1 | NGEF,SORCS3,RHOA |
| Regulation of actin dynamics for phagocytic cup formation | 62 | 3.59 | 2 | 0.882 | 1 | MYO9B,ELMO1 |
| Opioid Signalling | 86 | 4.98 | 3 | 0.883 | 1 | PDE1C,GNAI1,PDE4B |
| Fcgamma receptor (FCGR) dependent phagocytosis | 86 | 4.98 | 3 | 0.883 | 1 | MYO9B,ELMO1,PIK3R2 |
| Mitotic Prophase | 36 | 2.09 | 1 | 0.884 | 1 | VRK2 |
| Platelet Aggregation (Plug Formation) | 36 | 2.09 | 1 | 0.884 | 1 | F2 |
| Cell Cycle Checkpoints | 131 | 7.59 | 5 | 0.884 | 1 | RPA3,PSMA5,ANAPC4,MAD1L1,RF |
| Stimuli-sensing channels | 63 | 3.65 | 2 | 0.888 | 1 | NEDD4L,CLCN3 |
| ER-Phagosome pathway | 63 | 3.65 | 2 | 0.888 | 1 | HLA-B,PSMA5 |
| Class B/2 (Secretin family receptors) | 87 | 5.04 | 3 | 0.888 | 1 | PTCH1,CD55,WNT3 |
| PLC-gamma1 signalling | 37 | 2.14 | 1 | 0.891 | 1 | PDE1C |

| | | | | | | |
|---|---|---|---|---|---|---|
| EGFR interacts with phospholipase C-gamma | 37 | 2.14 | 1 | 0.891 | 1 | PDE1C |
| Factors involved in megakaryocyte development and platelet production | 155 | 8.98 | 6 | 0.893 | 1 | KLC1,DOCK8,ZFPM2,CARMIL1,JM[illegible] |
| Constitutive PI3K/AKT Signaling in Cancer | 89 | 5.16 | 3 | 0.897 | 1 | FGFR1,FOXO3,PIK3R2 |
| PLCG1 events in ERBB2 signaling | 38 | 2.2 | 1 | 0.897 | 1 | PDE1C |
| Post-translational protein modification | 200 | 11.6 | 8 | 0.902 | 1 | GALNT12,F2,SEMA6D,ST3GAL3,F[illegible] |
| FRS2-mediated cascade | 39 | 2.26 | 1 | 0.903 | 1 | FGFR1 |
| Activation of NMDA receptor upon glutamate binding and postsynaptic events | 39 | 2.26 | 1 | 0.903 | 1 | GRIN2A |
| Autodegradation of Cdh1 by Cdh1:APC/C | 68 | 3.94 | 2 | 0.912 | 1 | PSMA5,ANAPC4 |
| Cyclin D associated events in G1 | 41 | 2.38 | 1 | 0.914 | 1 | RBL2 |
| G1 Phase | 41 | 2.38 | 1 | 0.914 | 1 | RBL2 |

| | | | | | | |
|---|---|---|---|---|---|---|
| RNA Polymerase II Promoter Escape | 41 | 2.38 | 1 | 0.914 | 1 | POLR2E |
| RNA Polymerase II Transcription Pre-Initiation And Promoter Opening | 41 | 2.38 | 1 | 0.914 | 1 | POLR2E |
| RNA Polymerase II Transcription Initiation | 41 | 2.38 | 1 | 0.914 | 1 | POLR2E |
| RNA Polymerase II Transcription Initiation And Promoter Clearance | 41 | 2.38 | 1 | 0.914 | 1 | POLR2E |
| Negative regulation of FGFR signaling | 41 | 2.38 | 1 | 0.914 | 1 | FGFR1 |
| HIV-1 Transcription Initiation | 41 | 2.38 | 1 | 0.914 | 1 | POLR2E |
| RNA Polymerase II HIV-1 Promoter Escape | 41 | 2.38 | 1 | 0.914 | 1 | POLR2E |
| Kinesins | 41 | 2.38 | 1 | 0.914 | 1 | KLC1 |
| Meiosis | 117 | 6.78 | 4 | 0.915 | 1 | RPA3,MSH5,STAG1,SYNE1 |
| tRNA Aminoacylation | 42 | 2.43 | 1 | 0.919 | 1 | PPA2 |
| Interleukin-2 signaling | 42 | 2.43 | 1 | 0.919 | 1 | PIK3R2 |
| Synthesis of DNA | 95 | 5.51 | 3 | 0.92 | 1 | RPA3,PSMA5,RFC2 |

| | | | | | | |
|---|---|---|---|---|---|---|
| Transport of glucose and other sugars, bile salts and organic acids, metal ions and amine compounds | 95 | 5.51 | 3 | 0.92 | 1 | SLC6A9,SLC13A1,SLC39A8 |
| Neuronal System | 292 | 16.9 | 12 | 0.924 | 1 | GNAI1,GRIN2A,CHRNA5,CACNA2D |
| APC/C:Cdc20 mediated degradation of Securin | 71 | 4.12 | 2 | 0.924 | 1 | PSMA5,ANAPC4 |
| Translocation of GLUT4 to the Plasma Membrane | 71 | 4.12 | 2 | 0.924 | 1 | EXOC4,PRKAG2 |
| Telomere Maintenance | 72 | 4.17 | 2 | 0.927 | 1 | RPA3,RFC2 |
| S Phase | 122 | 7.07 | 4 | 0.93 | 1 | RPA3,PSMA5,STAG1,RFC2 |
| mRNA Splicing - Minor Pathway | 45 | 2.61 | 1 | 0.933 | 1 | POLR2E |
| Tat-mediated elongation of the HIV-1 transcript | 45 | 2.61 | 1 | 0.933 | 1 | POLR2E |
| Formation of HIV-1 elongation complex containing HIV-1 Tat | 45 | 2.61 | 1 | 0.933 | 1 | POLR2E |
| HIV-1 Transcription Elongation | 45 | 2.61 | 1 | 0.933 | 1 | POLR2E |

| | | | | | | |
|---|---|---|---|---|---|---|
| NRAGE signals death through JNK | 45 | 2.61 | 1 | 0.933 | 1 | NGEF |
| Elastic fibre formation | 45 | 2.61 | 1 | 0.933 | 1 | FURIN |
| Chromosome Maintenance | 124 | 7.19 | 4 | 0.935 | 1 | RPA3,STAG1,RFC2,SYNE1 |
| RNA Polymerase II Transcription Elongation | 46 | 2.67 | 1 | 0.936 | 1 | POLR2E |
| Formation of RNA Pol II elongation complex | 46 | 2.67 | 1 | 0.936 | 1 | POLR2E |
| Formation of HIV-1 elongation complex in the absence of HIV-1 Tat | 46 | 2.67 | 1 | 0.936 | 1 | POLR2E |
| Cdc20:Phospho-APC/C mediated degradation of Cyclin A | 76 | 4.4 | 2 | 0.94 | 1 | PSMA5,ANAPC4 |
| APC/C:Cdh1 mediated degradation of Cdc20 and other APC/C:Cdh1 targeted proteins in late mitosis/early G1 | 76 | 4.4 | 2 | 0.94 | 1 | PSMA5,ANAPC4 |

| | | | | | | |
|---|---|---|---|---|---|---|
| APC/C:Cdc20 mediated degradation of mitotic proteins | 76 | 4.4 | 2 | 0.94 | 1 | PSMA5,ANAPC4 |
| Meiotic Synapsis | 76 | 4.4 | 2 | 0.94 | 1 | STAG1,SYNE1 |
| TRAF6 mediated induction of NFkB and MAP kinases upon TLR7/8 or 9 activation | 76 | 4.4 | 2 | 0.94 | 1 | MEF2C,TWF2 |
| DNA Replication | 102 | 5.91 | 3 | 0.941 | 1 | RPA3,PSMA5,RFC2 |
| Downstream signaling of activated FGFR | 150 | 8.69 | 5 | 0.941 | 1 | PDE1C,TNRC6A,FGFR1,FOXO3,PI |
| Activation of APC/C and APC/C:Cdc20 mediated degradation of mitotic proteins | 77 | 4.46 | 2 | 0.943 | 1 | PSMA5,ANAPC4 |
| Toll Like Receptor 7/8 (TLR7/8) Cascade | 77 | 4.46 | 2 | 0.943 | 1 | MEF2C,TWF2 |
| MyD88 dependent cascade initiated on endosome | 77 | 4.46 | 2 | 0.943 | 1 | MEF2C,TWF2 |

| | | | | | | |
|---|---|---|---|---|---|---|
| Regulation of activated PAK-2p34 by proteasome mediated degradation | 48 | 2.78 | 1 | 0.944 | 1 | PSMA5 |
| Cross-presentation of soluble exogenous antigens (endosomes) | 48 | 2.78 | 1 | 0.944 | 1 | PSMA5 |
| Ion channel transport | 128 | 7.42 | 4 | 0.945 | 1 | NEDD4L,ATP2A2,CLCN3,ATP2B2 |
| Antigen processing-Cross presentation | 78 | 4.52 | 2 | 0.946 | 1 | HLA-B,PSMA5 |
| Ubiquitin-dependent degradation of Cyclin D1 | 49 | 2.84 | 1 | 0.947 | 1 | PSMA5 |
| CDK-mediated phosphorylation and removal of Cdc6 | 49 | 2.84 | 1 | 0.947 | 1 | PSMA5 |
| Ubiquitin-dependent degradation of Cyclin D | 49 | 2.84 | 1 | 0.947 | 1 | PSMA5 |
| Regulation of ornithine decarboxylase (ODC) | 49 | 2.84 | 1 | 0.947 | 1 | PSMA5 |
| Inositol phosphate metabolism | 49 | 2.84 | 1 | 0.947 | 1 | IMPA2 |

| | | | | | | |
|---|---|---|---|---|---|---|
| Toll Like Receptor 9 (TLR9) Cascade | 79 | 4.58 | 2 | 0.949 | 1 | MEF2C,TWF2 |
| Vpu mediated degradation of CD4 | 50 | 2.9 | 1 | 0.95 | 1 | PSMA5 |
| DNA Replication Pre-Initiation | 80 | 4.64 | 2 | 0.951 | 1 | RPA3,PSMA5 |
| M/G1 Transition | 80 | 4.64 | 2 | 0.951 | 1 | RPA3,PSMA5 |
| Immunoregulatory interactions between a Lymphoid and a non-Lymphoid cell | 80 | 4.64 | 2 | 0.951 | 1 | HLA-B,RAET1E |
| Signaling by FGFR in disease | 178 | 10.3 | 6 | 0.951 | 1 | PDE1C,MYO18A,TNRC6A,FGFR1,F |
| Interleukin-3, 5 and GM-CSF signaling | 51 | 2.96 | 1 | 0.953 | 1 | PIK3R2 |
| Generic Transcription Pathway | 434 | 25.2 | 18 | 0.954 | 1 | ESR2,ESRRG,NEDD4L,ZNF398,NR |
| Ubiquitin Mediated Degradation of Phosphorylated Cdc25A | 52 | 3.01 | 1 | 0.956 | 1 | PSMA5 |
| p53-Independent DNA Damage Response | 52 | 3.01 | 1 | 0.956 | 1 | PSMA5 |
| p53-Independent G1/S DNA damage checkpoint | 52 | 3.01 | 1 | 0.956 | 1 | PSMA5 |

| | | | | | | |
|---|---|---|---|---|---|---|
| SCF-beta-TrCP mediated degradation of Emi1 | 53 | 3.07 | 1 | 0.958 | 1 | PSMA5 |
| Autodegradation of the E3 ubiquitin ligase COP1 | 53 | 3.07 | 1 | 0.958 | 1 | PSMA5 |
| Regulation of APC/C activators between G1/S and early anaphase | 84 | 4.87 | 2 | 0.96 | 1 | PSMA5,ANAPC4 |
| Meiotic Recombination | 84 | 4.87 | 2 | 0.96 | 1 | RPA3,MSH5 |
| Stabilization of p53 | 54 | 3.13 | 1 | 0.961 | 1 | PSMA5 |
| Destabilization of mRNA by AUF1 (hnRNP D0) | 54 | 3.13 | 1 | 0.961 | 1 | PSMA5 |
| Assembly of collagen fibrils and other multimeric structures | 54 | 3.13 | 1 | 0.961 | 1 | COL11A1 |
| NGF signalling via TRKA from the plasma membrane | 207 | 12 | 7 | 0.961 | 1 | PDE1C,TNRC6A,FGFR1,MEF2C,F[illegible] |
| Signaling by FGFR | 162 | 9.39 | 5 | 0.963 | 1 | PDE1C,TNRC6A,FGFR1,FOXO3,P[illegible] |
| Vif-mediated degradation of APOBEC3G | 55 | 3.19 | 1 | 0.963 | 1 | PSMA5 |

| | | | | | | |
|---|---|---|---|---|---|---|
| MAP kinase activation in TLR cascade | 55 | 3.19 | 1 | 0.963 | 1 | MEF2C |
| Downstream signal transduction | 163 | 9.45 | 5 | 0.964 | 1 | PDE1C,TNRC6A,FGFR1,FOXO3,PI |
| Heparan sulfate/heparin (HS-GAG) metabolism | 56 | 3.25 | 1 | 0.965 | 1 | EXT1 |
| Signaling by ERBB2 | 164 | 9.51 | 5 | 0.965 | 1 | PDE1C,TNRC6A,FGFR1,FOXO3,PI |
| DAP12 signaling | 164 | 9.51 | 5 | 0.965 | 1 | PDE1C,TNRC6A,FGFR1,FOXO3,PI |
| Signaling by PDGF | 189 | 11 | 6 | 0.967 | 1 | PDE1C,TNRC6A,FGFR1,FURIN,FO |
| CDT1 association with the CDC6:ORC:origin complex | 57 | 3.3 | 1 | 0.967 | 1 | PSMA5 |
| Signaling by SCF-KIT | 142 | 8.23 | 4 | 0.969 | 1 | TNRC6A,FGFR1,FOXO3,PIK3R2 |
| APC/C-mediated degradation of cell cycle proteins | 89 | 5.16 | 2 | 0.969 | 1 | PSMA5,ANAPC4 |
| Regulation of mitotic cell cycle | 89 | 5.16 | 2 | 0.969 | 1 | PSMA5,ANAPC4 |
| SCF(Skp2)-mediated degradation of p27/p21 | 58 | 3.36 | 1 | 0.969 | 1 | PSMA5 |
| p53-Dependent G1/S DNA damage checkpoint | 59 | 3.42 | 1 | 0.971 | 1 | PSMA5 |

| | | | | | | |
|---|---|---|---|---|---|---|
| p53-Dependent G1 DNA Damage Response | 59 | 3.42 | 1 | 0.971 | 1 | PSMA5 |
| Regulation of Apoptosis | 59 | 3.42 | 1 | 0.971 | 1 | PSMA5 |
| Resolution of Sister Chromatid Cohesion | 118 | 6.84 | 3 | 0.971 | 1 | MAD1L1,STAG1,PPP2R5C |
| Collagen biosynthesis and modifying enzymes | 62 | 3.59 | 1 | 0.976 | 1 | COL11A1 |
| G1/S DNA Damage Checkpoints | 62 | 3.59 | 1 | 0.976 | 1 | PSMA5 |
| TRAF6 Mediated Induction of proinflammatory cytokines | 62 | 3.59 | 1 | 0.976 | 1 | MEF2C |
| RNA Polymerase II Pre-transcription Events | 62 | 3.59 | 1 | 0.976 | 1 | POLR2E |
| Cell death signalling via NRAGE, NRIF and NADE | 62 | 3.59 | 1 | 0.976 | 1 | NGEF |
| Downstream Signaling Events Of B Cell Receptor (BCR) | 173 | 10 | 5 | 0.976 | 1 | PSMA5,TNRC6A,FGFR1,FOXO3,PI |
| Mitotic Anaphase | 198 | 11.5 | 6 | 0.977 | 1 | PSMA5,VRK2,ANAPC4,MAD1L1,ST |
| Signalling by NGF | 290 | 16.8 | 10 | 0.977 | 1 | PDE1C,TNRC6A,FGFR1,MEF2C,N( |
| Assembly of the pre-replicative complex | 63 | 3.65 | 1 | 0.977 | 1 | PSMA5 |

| | | | | | | |
|---|---|---|---|---|---|---|
| Mitotic Metaphase and Anaphase | 199 | 11.5 | 6 | 0.977 | 1 | PSMA5,VRK2,ANAPC4,MAD1L1,ST |
| Signaling by the B Cell Receptor (BCR) | 199 | 11.5 | 6 | 0.977 | 1 | PSMA5,TNRC6A,CBLB,FGFR1,FO) |
| Transcription of the HIV genome | 64 | 3.71 | 1 | 0.979 | 1 | POLR2E |
| trans-Golgi Network Vesicle Budding | 64 | 3.71 | 1 | 0.979 | 1 | GBF1 |
| Clathrin derived vesicle budding | 64 | 3.71 | 1 | 0.979 | 1 | GBF1 |
| Signaling by ERBB4 | 152 | 8.81 | 4 | 0.98 | 1 | TNRC6A,FGFR1,FOXO3,PIK3R2 |
| Cyclin E associated events during G1/S transition | 65 | 3.77 | 1 | 0.98 | 1 | PSMA5 |
| Mitotic Prometaphase | 127 | 7.36 | 3 | 0.981 | 1 | MAD1L1,STAG1,PPP2R5C |
| Activation of NF-kappaB in B Cells | 66 | 3.83 | 1 | 0.981 | 1 | PSMA5 |
| Cyclin A:Cdk2-associated events at S phase entry | 66 | 3.83 | 1 | 0.981 | 1 | PSMA5 |
| Signaling by EGFR | 179 | 10.4 | 5 | 0.981 | 1 | PDE1C,TNRC6A,FGFR1,FOXO3,PI |
| Cell surface interactions at the vascular wall | 99 | 5.74 | 2 | 0.981 | 1 | F2,PIK3R2 |

| | | | | | | |
|---|---|---|---|---|---|---|
| RIG-I/MDA5 mediated induction of IFN-alpha/beta pathways | 67 | 3.88 | 1 | 0.982 | 1 | TANK |
| Signaling by EGFR in Cancer | 181 | 10.5 | 5 | 0.982 | 1 | PDE1C,TNRC6A,FGFR1,FOXO3,PI |
| Interferon alpha/beta signaling | 68 | 3.94 | 1 | 0.983 | 1 | HLA-B |
| DAP12 interactions | 182 | 10.5 | 5 | 0.983 | 1 | PDE1C,TNRC6A,FGFR1,FOXO3,PI |
| Switching of origins to a post-replicative state | 69 | 4 | 1 | 0.984 | 1 | PSMA5 |
| Orc1 removal from chromatin | 69 | 4 | 1 | 0.984 | 1 | PSMA5 |
| Sphingolipid metabolism | 70 | 4.06 | 1 | 0.985 | 1 | PRKD1 |
| Separation of Sister Chromatids | 186 | 10.8 | 5 | 0.986 | 1 | PSMA5,ANAPC4,MAD1L1,STAG1,F |
| Removal of licensing factors from origins | 71 | 4.12 | 1 | 0.986 | 1 | PSMA5 |
| Regulation of DNA replication | 71 | 4.12 | 1 | 0.986 | 1 | PSMA5 |
| Developmental Biology | 417 | 24.2 | 15 | 0.986 | 1 | SEMA3E,SEMA3A,TCF4,SEMA6D,F |
| Toll Like Receptor 10 (TLR10) Cascade | 74 | 4.29 | 1 | 0.988 | 1 | MEF2C |
| Toll Like Receptor 5 (TLR5) Cascade | 74 | 4.29 | 1 | 0.988 | 1 | MEF2C |

| | | | | | | |
|---|---|---|---|---|---|---|
| MyD88 cascade initiated on plasma membrane | 74 | 4.29 | 1 | 0.988 | 1 | MEF2C |
| Mitotic G1-G1/S phases | 140 | 8.11 | 3 | 0.99 | 1 | RPA3,PSMA5,RBL2 |
| Platelet activation, signaling and aggregation | 220 | 12.8 | 6 | 0.99 | 1 | DGKI,GNAI1,F2,CALU,RHOA,PIK3F |
| L1CAM interactions | 112 | 6.49 | 2 | 0.991 | 1 | FGFR1,ANK3 |
| G1/S Transition | 113 | 6.55 | 2 | 0.991 | 1 | RPA3,PSMA5 |
| Antigen processing: Ubiquitination & Proteasome degradation | 224 | 13 | 6 | 0.992 | 1 | PSMA5,ANAPC4,CUL3,NEDD4L,CE |
| Interferon Signaling | 173 | 10 | 4 | 0.992 | 1 | HLA-B,HLA-DQA1,HLA-DRB5,HLA-I |
| MyD88:Mal cascade initiated on plasma membrane | 81 | 4.69 | 1 | 0.992 | 1 | MEF2C |
| Toll Like Receptor TLR1:TLR2 Cascade | 81 | 4.69 | 1 | 0.992 | 1 | MEF2C |
| Toll Like Receptor TLR6:TLR2 Cascade | 81 | 4.69 | 1 | 0.992 | 1 | MEF2C |
| Toll Like Receptor 2 (TLR2) Cascade | 81 | 4.69 | 1 | 0.992 | 1 | MEF2C |
| Respiratory electron transport | 82 | 4.75 | 1 | 0.993 | 1 | NDUFA2 |
| Glycosaminoglycan metabolism | 118 | 6.84 | 2 | 0.993 | 1 | EXT1,ST3GAL3 |

| | | | | | | |
|---|---|---|---|---|---|---|
| MPS VI - Maroteaux-Lamy syndrome | 118 | 6.84 | 2 | 0.993 | 1 | EXT1,ST3GAL3 |
| Mucopolysaccharidoses | 118 | 6.84 | 2 | 0.993 | 1 | EXT1,ST3GAL3 |
| MPS IX - Natowicz syndrome | 118 | 6.84 | 2 | 0.993 | 1 | EXT1,ST3GAL3 |
| MPS IIIB - Sanfilippo syndrome B | 118 | 6.84 | 2 | 0.993 | 1 | EXT1,ST3GAL3 |
| MPS I - Hurler syndrome | 118 | 6.84 | 2 | 0.993 | 1 | EXT1,ST3GAL3 |
| MPS II - Hunter syndrome | 118 | 6.84 | 2 | 0.993 | 1 | EXT1,ST3GAL3 |
| MPS VII - Sly syndrome | 118 | 6.84 | 2 | 0.993 | 1 | EXT1,ST3GAL3 |
| MPS IV - Morquio syndrome A | 118 | 6.84 | 2 | 0.993 | 1 | EXT1,ST3GAL3 |
| MPS IIIC - Sanfilippo syndrome C | 118 | 6.84 | 2 | 0.993 | 1 | EXT1,ST3GAL3 |
| MPS IV - Morquio syndrome B | 118 | 6.84 | 2 | 0.993 | 1 | EXT1,ST3GAL3 |
| MPS IIIA - Sanfilippo syndrome A | 118 | 6.84 | 2 | 0.993 | 1 | EXT1,ST3GAL3 |
| MPS IIID - Sanfilippo syndrome D | 118 | 6.84 | 2 | 0.993 | 1 | EXT1,ST3GAL3 |
| Integrin cell surface interactions | 85 | 4.93 | 1 | 0.994 | 1 | ITGA11 |
| Collagen formation | 85 | 4.93 | 1 | 0.994 | 1 | COL11A1 |
| M Phase | 233 | 13.5 | 6 | 0.994 | 1 | PSMA5,VRK2,ANAPC4,MAD1L1,ST |

| | | | | | | |
|---|---|---|---|---|---|---|
| Asparagine N-linked glycosylation | 86 | 4.98 | 1 | 0.994 | 1 | RFT1 |
| Hemostasis | 511 | 29.6 | 18 | 0.994 | 1 | SLC8A3,DGKI,GNAI1,F2,KLC1,DOC |
| TRIF-mediated TLR3/TLR4 signaling | 87 | 5.04 | 1 | 0.995 | 1 | MEF2C |
| Toll-Like Receptors Cascades | 123 | 7.13 | 2 | 0.995 | 1 | MEF2C,TWF2 |
| MyD88-independent cascade | 88 | 5.1 | 1 | 0.995 | 1 | MEF2C |
| Toll Like Receptor 3 (TLR3) Cascade | 88 | 5.1 | 1 | 0.995 | 1 | MEF2C |
| Regulation of mRNA Stability by Proteins that Bind AU-rich Elements | 88 | 5.1 | 1 | 0.995 | 1 | PSMA5 |
| Platelet degranulation | 89 | 5.16 | 1 | 0.995 | 1 | CALU |
| Extracellular matrix organization | 157 | 9.1 | 3 | 0.995 | 1 | COL11A1,MMP16,FURIN |
| Apoptosis | 158 | 9.16 | 3 | 0.996 | 1 | PSMA5,MAPT,CTNNB1 |
| Mitotic M-M/G1 phases | 266 | 15.4 | 7 | 0.996 | 1 | RPA3,PSMA5,VRK2,ANAPC4,MAD[illegible] |
| G alpha (s) signalling events | 127 | 7.36 | 2 | 0.996 | 1 | PDE8B,PDE4B |
| Class I MHC mediated antigen processing & presentation | 267 | 15.5 | 7 | 0.996 | 1 | HLA-B,PSMA5,ANAPC4,CUL3,NED |
| Axon guidance | 292 | 16.9 | 8 | 0.996 | 1 | SEMA3E,SEMA3A,SEMA6D,FGFR1 |

| | | | | | | |
|---|---|---|---|---|---|---|
| HIV Life Cycle | 128 | 7.42 | 2 | 0.996 | 1 | XRCC6,POLR2E |
| Metabolism of amino acids and derivatives | 190 | 11 | 4 | 0.996 | 1 | PSMA5,CKB,IVD,PSPH |
| Response to elevated platelet cytosolic Ca2+ | 94 | 5.45 | 1 | 0.996 | 1 | CALU |
| Transport of inorganic cations/anions and amino acids/oligopeptides | 96 | 5.56 | 1 | 0.997 | 1 | SLC8A3 |
| Phospholipid metabolism | 135 | 7.82 | 2 | 0.997 | 1 | PIP5K1B,PIK3R2 |
| Activated TLR4 signalling | 100 | 5.8 | 1 | 0.998 | 1 | MEF2C |
| Respiratory electron transport, ATP synthesis by chemiosmotic coupling, and heat production by uncoupling proteins. | 101 | 5.85 | 1 | 0.998 | 1 | NDUFA2 |
| Fatty acid, triacylglycerol, and ketone body metabolism | 139 | 8.06 | 2 | 0.998 | 1 | GPD2,PRKAG2 |
| Toll Like Receptor 4 (TLR4) Cascade | 103 | 5.97 | 1 | 0.998 | 1 | MEF2C |

| | | | | | | |
|---|---|---|---|---|---|---|
| Host Interactions of HIV factors | 141 | 8.17 | 2 | 0.998 | 1 | PSMA5,ELMO1 |
| RNA Polymerase II Transcription | 107 | 6.2 | 1 | 0.998 | 1 | POLR2E |
| Late Phase of HIV Life Cycle | 108 | 6.26 | 1 | 0.998 | 1 | POLR2E |
| Transcription | 149 | 8.64 | 2 | 0.999 | 1 | TAF1C,POLR2E |
| HIV Infection | 214 | 12.4 | 4 | 0.999 | 1 | PSMA5,XRCC6,POLR2E,ELMO1 |
| mRNA Splicing | 115 | 6.67 | 1 | 0.999 | 1 | POLR2E |
| mRNA Splicing - Major Pathway | 115 | 6.67 | 1 | 0.999 | 1 | POLR2E |
| G alpha (q) signalling events | 188 | 10.9 | 3 | 0.999 | 1 | DGKI,F2,PIK3R2 |
| Signaling by Interleukins | 116 | 6.72 | 1 | 0.999 | 1 | PIK3R2 |
| Processing of Capped Intron-Containing Pre-mRNA | 119 | 6.9 | 1 | 0.999 | 1 | POLR2E |
| Membrane Trafficking | 203 | 11.8 | 3 | 1 | 1 | EXOC4,PRKAG2,GBF1 |
| Gastrin-CREB signalling pathway via PKC and MAPK | 209 | 12.1 | 3 | 1 | 1 | DGKI,F2,PIK3R2 |
| Adaptive Immune System | 654 | 37.9 | 20 | 1 | 1 | HLA-B,HLA-DMA,PSMA5,ANAPC4,I |
| mRNA Processing | 140 | 8.11 | 1 | 1 | 1 | POLR2E |

| | | | | | | |
|---|---|---|---|---|---|---|
| SLC-mediated transmembrane transport | 251 | 14.5 | 4 | 1 | 1 | SLC8A3,SLC6A9,SLC13A1,SLC39A |
| G alpha (i) signalling events | 184 | 10.7 | 2 | 1 | 1 | GNAI1,GABBR1 |
| The citric acid (TCA) cycle and respiratory electron transport | 145 | 8.4 | 1 | 1 | 1 | NDUFA2 |
| Cytokine Signaling in Immune system | 286 | 16.6 | 5 | 1 | 1 | HLA-B,HLA-DQA1,HLA-DRB5,HLA-[illegible] |
| Cell Cycle, Mitotic | 411 | 23.8 | 9 | 1 | 1 | RPA3,PSMA5,VRK2,ANAPC4,MAD[illegible] |
| Eukaryotic Translation Termination | 178 | 10.3 | 1 | 1 | 1 | ETF1 |
| Nonsense Mediated Decay Independent of the Exon Junction Complex | 184 | 10.7 | 1 | 1 | 1 | ETF1 |
| Peptide ligand-binding receptors | 192 | 11.1 | 1 | 1 | 1 | F2 |
| Innate Immune System | 521 | 30.2 | 11 | 1 | 1 | PDE1C,TNRC6A,MYO9B,FGFR1,M |
| Nonsense Mediated Decay Enhanced by the Exon Junction Complex | 203 | 11.8 | 1 | 1 | 1 | ETF1 |

| | | | | | | |
|---|---|---|---|---|---|---|
| Nonsense-Mediated Decay | 203 | 11.8 | 1 | 1 | 1 | ETF1 |
| Cell Cycle | 508 | 29.4 | 10 | 1 | 1 | RPA3,PSMA5,VRK2,ANAPC4,MAD[illegible] |
| Metabolism of carbohydrates | 258 | 15 | 2 | 1 | 1 | EXT1,ST3GAL3 |
| Transmembrane transport of small molecules | 504 | 29.2 | 8 | 1 | 1 | SLC8A3,SLC6A9,NEDD4L,ATP2A2, |
| Translation | 249 | 14.4 | 1 | 1 | 1 | ETF1 |
| GPCR ligand binding | 415 | 24.1 | 5 | 1 | 1 | PTCH1,F2,GABBR1,CD55,WNT3 |
| Metabolism of mRNA | 317 | 18.4 | 2 | 1 | 1 | PSMA5,ETF1 |
| Metabolism of proteins | 689 | 39.9 | 13 | 1 | 1 | FKBP9,GALNT12,EXOC4,DPP4,F2, |
| Metabolism of RNA | 339 | 19.6 | 2 | 1 | 1 | PSMA5,ETF1 |
| Metabolism of lipids and lipoproteins | 507 | 29.4 | 6 | 1 | 1 | FADS2,PRKD1,GPD2,PRKAG2,PIP[illegible] |
| Class A/1 (Rhodopsin-like receptors) | 312 | 18.1 | 1 | 1 | 1 | F2 |
| Immune System | 1140 | 66 | 27 | 1 | 1 | PDE1C,HLA-B,HLA-DMA,PSMA5,A[illegible] |
| Gene Expression | 1090 | 62.9 | 24 | 1 | 1 | PSMA5,ESR2,ESRRG,TAF1C,TNR[illegible] |
| Disease | 945 | 54.8 | 17 | 1 | 1 | PDE1C,MYO18A,PSMA5,TNRC6A,[illegible] |
| Signaling by GPCR | 931 | 54 | 13 | 1 | 1 | PDE1C,DGKI,GNAI1,PTCH1,F2,PD[illegible] |
| GPCR downstream signaling | 812 | 47.1 | 9 | 1 | 1 | DGKI,GNAI1,F2,PDE8B,GABBR1,N[illegible] |
| Signal Transduction | 1690 | 97.8 | 38 | 1 | 1 | PDE1C,DGKI,ARHGAP15,GNAI1,P[illegible] |
| Metabolism | 1490 | 86.2 | 22 | 1 | 1 | GNAI1,PSMA5,FADS2,PRKD1,CAC |

1D2,ZNF568,ZNF19,ZKSCAN3,MED27,ESRRB,MAML3,ZNF600,RNF111,ZNF615,THRB,TRIM33,MED8,ZK

;K8,ZFPM2,CALU,ATP2A2,GUCY1A2,CARMIL1,PRKG1,JMJD1C,ATP2B2,PPP2R5C,DOCK4,RHOA,PIK3R2

)YNC1I1,KLC1,TNRC6A,CUL3,NEDD4L,CBLB,RAET1E,FGFR1,HLA-DQA1,PRKG1,FOXO3,HLA-DRB5,RNF

NAPC4,DYNC1I1,KLC1,TNRC6A,MYO9B,CUL3,NEDD4L,CBLB,RAET1E,FGFR1,HLA-DQA1,MEF2C,CD55,P

C6A,NEDD4L,ZNF398,NR1D2,ETF1,ZNF568,ZNF19,ZKSCAN3,MED27,ESRRB,MAML3,ZNF600,RNF111,PC

HDAC4,EXT1,FGFR1,XRCC6,ST3GAL3,SYT1,MAML3,FOXO3,POLR2E,ELMO1,NEURL1,PIK3R2,HDAC9

SMA5,PTCH1,F2,PDE8B,TNRC6A,MYO9B,RPTOR,HDAC4,GABBR1,NEDD4L,ITGA11,FGFR1,PRKAG2,MEI

NA2D2,KCNG2,PFAS,GPD2,EXT1,CKB,NT5C2,PRKAG2,IMPA2,ST3GAL3,CACNB2,KCNB1,DPYD,IVD,PSF

[illegible]2C,GMIP,RHOJ,CD55,CTNNB1,ST3GAL3,NGEF,FURIN,TWF2,MAML3,FOXO3,PDE4B,RNF111,SORCS3,[illegible]

WNT3,PPP2R5C,TRIM33,NEURL1,RHOA,PIK3R2,HDAC9
