## Supplemental table 9 for "Digestive Dimensions of Autism: A Multiscale Exploration of Gut-Brain Interactions"

| Pathway | Total | Expected | Hits | P.Value | FDR | Genes | function |
|---|---|---|---|---|---|---|---|
| YNGTTNNNATT_UNKNOWN | 365 | 10 | 26 | 9.25E-06 | 0.00767 | MAGI1,PDE1C,SEMA3A,THSD4,ESRRG,EXOC4,PKP4,BEND4,CNTNAP2,ZFPM2,CBLB,ATP2A2,CITED1,ZNF638,CNOT1,CALD1,CTNND1,SOX5,MAML3,TSHZ3,JMJD1C,NFIX,NTRK3,ELMO1,THRB,DOCK4 | Genes with 3'UTR conta |
| V$P300_01 | 249 | 6.84 | 20 | 1.83E-05 | 0.00767 | KMT2A,RERE,FOXP1,ESRRG,TNRC6A,OLFM4,ANKS1B,DMTF1,GATB,FGFR1,PRKAG2,CACNB2,FURIN,MAML3,BCL11A,SRPK2,ELMO1,WNT3,PREX1,ZNF800 | Genes with 3'UTR conta |
| TAGCTTT,MIR-9 | 236 | 6.48 | 18 | 9.36E-05 | 0.0256 | MAGI1,KLC1,ZFPM2,CNTN4,FXR1,MMP16,ETF1,CREB5,MEF2C,CALD1,KCNN2,SORCS3,NEGR1,THRB,TRIM33,BCL11B,AFF3,SREK1IP1 | Genes having at least or |

| | | | | | | | |
|---|---|---|---|---|---|---|---|
| AACTTT_ UNKNOWN | 1890 | 51.9 | 78 | 0.000123 | 0.0256 | MAGI1,NPAS3,KMT2D,MYO18A,KMT2A,DGKI,GNAI1,SEMA3A,GIGYF2,NCOA5,ESRRG,UTY,KIZ,PTCH1,PLCL1,TCF4,DYNC1I1,CLSTN2,CUL3,CNTNAP2,HDAC4,PPM1E,ZFPM2,CNTN4,CBLB,SEMA6D,ANKS1B,MMP16,EXT1,ZNF804A,CITED1,LDB1,ARID1B,EIF1AY,PRPF3,FGFR1,PRKAG2,AUTS2,ETF1,CREB5,MEF2C,GRIK1,DDX3Y,CALD1,C | Genes having at least or |
| TGAATGT, MIR-181A,MIR-181B,MIR-181C,MIR-181D | 484 | 13.3 | 28 | 0.000173 | 0.0289 | AKAP6,FOXP1,MAPT,CACNA2D2,CUL3,CNTNAP2,CNTN4,CBLB,ATP2A2,ARFGEF2,ETF1,DDX3Y,SOX5,STAG1,CACNB2,PLCL2,BCL11A,GID4,SRPK2,NEGR1,SYNE1,ZNF615,NOVA1,THRB,ATP2B2,BRWD1,MED8,ZNF800 | In humans miRNA-181 i |

| | | | | | | | |
|---|---|---|---|---|---|---|---|
| V$OCT1_03 | 230 | 6.32 | 17 | 0.000212 | 0.0291 | RPA3,SEMA3A,TCF4,BEND4,ANKS1B,CREB5,MEF2C,SOX5,PRKG1,FOXO3,RFT1,KDM4A,PPP2R2B,NFIX,TMOD3,HDAC9,TLK1 | Genes with 3'UTR conta |
| CAGTATT,MIR-200B,MIR-200C,MIR-429 | 469 | 12.9 | 27 | 0.000244 | 0.0291 | MSL2,ESRRG,PLCL1,PPM1E,ZFPM2,CNTN4,SEMA6D,FXR1,CALU,ATP2A2,CACNA1C,CREB5,DDX3Y,NGEF,SYT1,ANK3,SEMA3F,NEGR1,NOVA1,PPP2R5C,BRWD1,TRIM33,BCL11B,AMBRA1,GLI3,RHOA,AFF3 | miRNAs that balance ep |
| AAGCCAT,MIR-135A,MIR-135B | 335 | 9.2 | 21 | 0.000393 | 0.0307 | MSL2,SYNGAP1,PDE8B,BSN,SEMA6D,CREB5,MEF2C,DDX3Y,ANK3,CTTNBP2,BCL11A,NEGR1,THRB,ATP2B2,PPP2R5C,BRWD1,BCL11B,NEURL1,PIK3R2,RNF144A,TLK1 | Both miRNAs are involv |

| | | | | | | | |
|---|---|---|---|---|---|---|---|
| ATGTTAA, MIR-302C | 243 | 6.68 | 17 | 0.000405 | 0.0307 | MAGI1,KMT2E,FOXP1,ZFPM2,FXR1,ZMIZ1,ARID1B,CREB5,DDX3Y,CADM2,SNAP91,BCL11A,GID4,NFIX,WNT3,PCDH9,PPA2 | miR-302 plays an import |
| ATGTACA, MIR-493 | 314 | 8.63 | 20 | 0.000439 | 0.0307 | DGKI,ESRRG,EFL1,SLC22A23,ATP2A2,ZMIZ1,ZNF804A,ARID1B,ETF1,MEF2C,CTNND1,MAML3,GBF1,SORCS3,SRPK2,SYNE1,PREX1,PPP2R5C,AFF3,ZNF800 | miRNA-493 is involved i |
| V$GATA1_03 | 245 | 6.73 | 17 | 0.000445 | 0.0307 | FBXL18,ARHGAP15,FOXP1,GABBR1,ZFPM2,BSN,MMP16,CREB5,MEF2C,SOX5,FURIN,MAML3,KCNN2,TNXB,PPP1R16A,PCDH9,DNAH11 | transcription factor bindi |

| | | | | | | | |
|---|---|---|---|---|---|---|---|
| AAGCAAT, MIR-137 | 223 | 6.13 | 16 | 0.000451 | 0.0307 | NREP,KMT2A,GRIN2A,ESRRG,CUL3,PPM1E,BSN,MMP16,MGAT5B,ST3GAL3,SYT1,FURIN,CADPS,MAML3,NOVA1,TRIM33 | MiR-137 has a critical re |
| GCANCTGNY_V$MYOD_Q6 | 924 | 25.4 | 43 | 0.000478 | 0.0307 | NRGN,SLC8A3,MAGI1,MYO18A,KMT2A,RPA3,KMT2E,ENOX1,GRIN2A,GIGYF2,PKP4,NEDD4L,ATP2A2,ZMIZ1,LDB1,FGFR1,PRKAG2,AUTS2,ETF1,GMIP,CADM2,SOX5,CACNB2,ABCB9,PPP1R16B,CADPS,LRFN5,HLA-DRB5,KCNN2,PARD3B,HLA-DRB1,SRPK2,NTRK3,NEGR1,SYNE1,NOVA1,WNT3,PCDH9,CLCN3,MED8,NEURL1,HDAC9,CA | Genes with 3'UTR conta |

| | | | | | | | |
|---|---|---|---|---|---|---|---|
| V$CART1_01 | 226 | 6.21 | 16 | 0.000522 | 0.0312 | ESRRG,PTCH1,BEND4,TNRC6A,PPM1E,ZFPM2,ANKS1B,CREB5,MEF2C,CALD1,CADM2,SOX5,LRFN5,BCL11A,NFIX,WNT3 | TF binding site of cartila |
| V$PBX1_01 | 252 | 6.92 | 17 | 0.000613 | 0.0341 | KMT2A,MSL2,RPA3,ARHGAP15,FE | |
| TTGTTT_V$FOXO4_01 | 2060 | 56.6 | 80 | 0.000695 | 0.0344 | NREP,MAGI1,NPAS3,MYO18A,KMT2A,MSL2,KMT2E,FOXP1,SEMA3A,RTN1,ESRRG,GPR141,ZSCAN9,PTCH1,PRKD1,EXOC4,TCF4,CACNA2D2,CLSTN2,CTR9,TNRC6A,PKD1L3,ZFPM2,BSN,OLFM4,CBLB,SEMA6D,ANKS1B,FXR1,EXT1,ATP2A2,ZNF804A,CITED1,CKB,LDB1,GATB,ARID1B,NT5C2,CNOT1,GUCY1A2,SLC13A1,PRKAG2,SLC30A9, | TF binding site of FOXO |

| | | | | | | | |
|---|---|---|---|---|---|---|---|
| V$SRY_02 | 255 | 7.01 | 17 | 7.00E-04 | 0.0344 | NREP,MYO18A,ARHGAP15,ESRRG,TCF4,TNRC6A,LDB1,SLC13A1,SOX5,CTNNB1,MAML3,KCNN2,ZFYVE1,PPP2R2B,NFIX,CSRNP3,MED8 | initiation of male sex det |
| GTGGTGA,MIR-197 | 72 | 1.98 | 8 | 0.000779 | 0.0362 | SYNGAP1,SPPL3,USP4,PPM1E,TTLL7,ETF1,CTNND1,GATAD2B | regulates cell proliferatic |
| V$CP2_01 | 260 | 7.14 | 17 | 0.000869 | 0.0382 | SLC8A3,KMT2E,RTN1,PSMA5,CLSTN2,RBM6,DMTF1,CITED1,CALD1,CADM2,RNF111,KCNN2,PPP2R2B,TNXB,WNT3,RHOA,HDAC9 | plays multifaceted role ir |
| V$SOX5_01 | 265 | 7.28 | 17 | 0.00107 | 0.0448 | ARHGAP15,RERE,SEMA3A,ESRRG,PTCH1,TCF4,LDB1,SLC13A1,AUTS2,SOX5,CTNNB1,ANK3,MAML3,BCL11A,ZFYVE1,PPP2R2B,NOVA1 | regulation of embryonic |
| CTGCAGY_UNKNOWN | 765 | 21 | 36 | 0.00114 | 0.0455 | NRGN,MYO18A,KMT2E,GNAI1,FOX | |

| | | | | | | | |
|---|---|---|---|---|---|---|---|
| CCATCCA,MIR-432 | 60 | 1.65 | 7 | 0.00123 | 0.0469 | PPM1E,COL11A1,PCGF3,ABCB9,FURIN,PRKG1,NTRK3 | neurogenesis, neuroplas |
| TGTTTAC,MIR-30A-5P,MIR-30C,MIR-30D,MIR-30B,MIR-30E-5P | 579 | 15.9 | 29 | 0.00137 | 0.0497 | KMT2A,ERLIN1,SEMA3A,GIGYF2,ESRRG,SBK1,TNRC6A,SLC6A9,BSN,NEDD4L,CBLB,CALU,ATP2A2,TTLL7,MAGI2,NEK4,CACNB2,FBXL17,CADPS,PIP5K1B,GLCCI1,SORCS3,GALNT1,TNXB,ATP2B2,BRWD1,PIK3R2,NAALADL2,HDAC9 | The miR-30 family partic |
| GACTGTT,MIR-212,MIR-132 | 161 | 4.42 | 12 | 0.00164 | 0.0554 | NREP,SPPL3,FXR1,MMP16,CALU,( | |
| YTATTTTNR_V$MEF2_02 | 697 | 19.2 | 33 | 0.00166 | 0.0554 | NREP,SLC8A3,MSL2,DGKI,KMT2E, | |
| TTTGCAG,MIR-518A-2 | 210 | 5.77 | 14 | 0.00202 | 0.0651 | MSL2,HDAC4,NEDD4L,ETF1,RASG | |
| SYATTGTG_UNKNOWN | 235 | 6.46 | 15 | 0.00217 | 0.0673 | KMT2A,ADAM22,FOXP1,SEMA3A,F | |
| V$SOX9_B1 | 237 | 6.51 | 15 | 0.00236 | 0.0704 | MYO18A,RERE,BSN,FGFR1,SOX5, | |
| V$CHOP_01 | 238 | 6.54 | 15 | 0.00246 | 0.0708 | NPAS3,PTCH1,ETF1,MEF2C,RHOJ | |
| TCATCTC,MIR-143 | 149 | 4.09 | 11 | 0.00273 | 0.0742 | AKAP6,GIGYF2,ESRRG,PPM1E,BS | |
| V$GATA6_01 | 265 | 7.28 | 16 | 0.00275 | 0.0742 | FOXP1,ESRRG,KIZ,DOCK8,PPM1E | |
| V$S8_01 | 245 | 6.73 | 15 | 0.00324 | 0.0845 | NPAS3,FOXP1,SEMA3A,GRIN2A,E | |
| V$FAC1_01 | 222 | 6.1 | 14 | 0.00336 | 0.0849 | MSRA,FOXP1,SEMA3A,SPPL3,KIZ, | |

| | | | | | | |
|---|---|---|---|---|---|---|
| TCTGATA, MIR-361 | 91 | 2.5 | 8 | 0.00353 | 0.0849 | MAGI1,ZFPM2,SND1,SDK1,NTM,PI |
| ACAACTT, MIR-382 | 72 | 1.98 | 7 | 0.00356 | 0.0849 | MMP16,ARID1B,DDX3Y,SYT1,SOR |
| V$TCF11_01 | 253 | 6.95 | 15 | 0.00436 | 0.101 | NREP,KMT2A,GNAI1,RSRC1,MAPT |
| CAGNYGKNAAA_UNKNOWN | 75 | 2.06 | 7 | 0.00447 | 0.101 | MYO18A,TET2,PRKD1,CACNA2D2, |
| CTTGTAT, MIR-381 | 206 | 5.66 | 13 | 0.00461 | 0.101 | MSL2,ZFPM2,MPHOSPH9,SEMA6D |
| CTGAGCC,MIR-24 | 231 | 6.35 | 14 | 0.00478 | 0.103 | MAGI1,SBK1,BSN,MMP16,RASGRF |
| V$DBP_Q6 | 257 | 7.06 | 15 | 0.00504 | 0.104 | KMT2D,RERE,PTCH1,SLC6A9,ANK |
| TGCTTTG,MIR-330 | 335 | 9.2 | 18 | 0.00532 | 0.104 | SLC8A3,MSL2,GIGYF2,PPM1E,ANK |
| TGGAAA_V$NFAT_Q4_01 | 1900 | 52.1 | 70 | 0.00554 | 0.104 | NRGN,NREP,KMT2D,KMT2A,HLA-E |
| CACTGTG,MIR-128A,MIR-128B | 337 | 9.26 | 18 | 0.00565 | 0.104 | NREP,KMT2A,RERE,PRKD1,CLSTN |
| AACATTC, MIR-409-3P | 142 | 3.9 | 10 | 0.00584 | 0.104 | FOXP1,HDAC4,GPD2,SEMA6D,CR |
| YCATTAA_UNKNOWN | 556 | 15.3 | 26 | 0.00589 | 0.104 | SLC8A3,NPAS3,MSRA,ADAM22,RT |
| TAAWWATAG_V$RSRFC4_Q2 | 165 | 4.53 | 11 | 0.00591 | 0.104 | NREP,SLC8A3,DGKI,FOXP1,PTCH |
| V$BRN2_01 | 237 | 6.51 | 14 | 0.00597 | 0.104 | ESRRG,TCF4,ANKS1B,MMP16,ZMI |
| CACTTTG, MIR-520G,MIR-520H | 237 | 6.51 | 14 | 0.00597 | 0.104 | NPAS3,EFL1,TNRC6A,CUL3,SLC6A |
| V$OCT1_02 | 214 | 5.88 | 13 | 0.00631 | 0.104 | NPAS3,FOXP1,SEMA3A,ESRRG,TE |
| V$CEBPDELTA_Q6 | 240 | 6.59 | 14 | 0.00665 | 0.104 | NREP,RPA3,FOXP1,SEMA3A,PTCH |
| V$NFAT_Q4_01 | 266 | 7.31 | 15 | 0.00686 | 0.104 | NREP,FOXP1,SEMA3A,NEDD4L,EX |
| V$NKX62_Q2 | 241 | 6.62 | 14 | 0.00689 | 0.104 | GNAI1,BEND4,TNRC6A,ZFPM2,ATI |
| V$SMAD4_Q6 | 241 | 6.62 | 14 | 0.00689 | 0.104 | KMT2E,TET2,TCF4,ANKS1B,ATP2A |
| RNGTGGGC_UNKNOWN | 766 | 21 | 33 | 0.00704 | 0.104 | FBXL18,NRGN,SLC8A3,MAGI1,MSI |
| GTCTTCC,MIR-7 | 169 | 4.64 | 11 | 0.00705 | 0.104 | NREP,SLC8A3,ESRRG,SLC6A9,GA |

| | | | | | | |
|---|---|---|---|---|---|---|
| AAGCACA,MIR-218 | 398 | 10.9 | 20 | 0.00706 | 0.104 | KMT2A,MSL2,GNAI1,PKP4,CUL3,T |
| V$MYB_Q5_01 | 267 | 7.34 | 15 | 0.0071 | 0.104 | NRGN,KMT2D,MYO18A,KMT2E,ES |
| YGCGYRCGC_UNKNOWN | 319 | 8.76 | 17 | 0.0072 | 0.104 | NRGN,MAGI1,MSL2,MAPT,SGF29,[illegible] |
| V$PAX4_04 | 218 | 5.99 | 13 | 0.00734 | 0.104 | NREP,FOXP1,TCF4,BEND4,TNRC[illegible] |
| TTANTCA_UNKNOWN | 952 | 26.2 | 39 | 0.00799 | 0.111 | SLC8A3,MAGI1,DGKI,RPA3,KMT2[illegible] |
| YATTNATC_UNKNOWN | 377 | 10.4 | 19 | 0.00828 | 0.113 | NREP,NPAS3,SEMA3A,PKP4,BEN[illegible] |
| V$PAX8_B | 106 | 2.91 | 8 | 0.00879 | 0.116 | NRGN,PTCH1,BEND4,CALD1,TAN[illegible] |
| V$E4BP4_01 | 223 | 6.13 | 13 | 0.00879 | 0.116 | NRGN,NREP,DGKI,ESRRG,BSN,O[illegible] |
| V$RORA2_01 | 151 | 4.15 | 10 | 0.00887 | 0.116 | SEMA3A,BEND4,CITED1,LRRC20,[illegible] |
| TTGCCAA,MIR-182 | 327 | 8.98 | 17 | 0.00911 | 0.117 | MAGI1,KMT2A,MSL2,PRKD1,EXOC |
| V$MYB_Q3 | 250 | 6.87 | 14 | 0.00937 | 0.117 | NRGN,KMT2D,DGKI,KMT2E,FOXP[illegible] |
| TGTYNNNNNRGCARM_UNKNOWN | 86 | 2.36 | 7 | 0.00937 | 0.117 | DGKI,PPM1E,CITED1,TSHZ3,ZFY[illegible] |
| TGTATGA,MIR-485-3P | 153 | 4.2 | 10 | 0.00969 | 0.117 | GNAI1,GIGYF2,HDAC4,ANKS1B,M[illegible] |
| RTTTNNNYTGGM_UNKNOWN | 153 | 4.2 | 10 | 0.00969 | 0.117 | FOXP1,FHIT,SLC6A9,CREB5,CAL[illegible] |
| GTGCCAA,MIR-96 | 303 | 8.33 | 16 | 0.00976 | 0.117 | MAGI1,KMT2A,ERLIN1,CACNA2D2 |
| ATGTCAC,MIR-489 | 88 | 2.42 | 7 | 0.0106 | 0.122 | RERE,RSU1,DDX3Y,BCL11A,NTRK |
| V$CREBP1_01 | 179 | 4.92 | 11 | 0.0106 | 0.122 | NREP,DGKI,BSN,ARID1B,ZNF638,[illegible] |
| GTGTGAG,MIR-342 | 68 | 1.87 | 6 | 0.0108 | 0.122 | KMT2A,EFL1,ARID1B,DDX3Y,CTN[illegible] |
| ACTGCAG,MIR-17-3P | 110 | 3.02 | 8 | 0.0109 | 0.122 | FOXP1,CTR9,RASGRP4,MAML3,T[illegible] |
| V$CDC5_01 | 256 | 7.03 | 14 | 0.0114 | 0.122 | NREP,FOXP1,ESRRG,CHRNA5,ZF |
| AAACCAC,MIR-140 | 111 | 3.05 | 8 | 0.0114 | 0.122 | DPP4,CUL3,HDAC4,CACNA1C,DD[illegible] |
| TGCAAAC,MIR-452 | 111 | 3.05 | 8 | 0.0114 | 0.122 | MMP16,CACNB2,ANK3,FURIN,TSH |
| ATGAAGG,MIR-205 | 157 | 4.31 | 10 | 0.0115 | 0.122 | MAGI1,MSL2,ESRRG,DMXL2,ZNF5 |

| | | | | | | |
|---|---|---|---|---|---|---|
| V$TATA_C | 283 | 7.78 | 15 | 0.0117 | 0.122 | FOXP1,SEMA3A,RTN1,ESR2,ESRF |
| GTACTGT,MIR-101 | 257 | 7.06 | 14 | 0.0117 | 0.122 | MAGI1,KMT2A,SYNGAP1,PTCH1,D |
| CAGTGTT,MIR-141,MIR-200A | 310 | 8.52 | 16 | 0.0119 | 0.122 | FOXP1,ESRRG,CUL3,SLC6A9,HDA |
| CTCTGGA,MIR-520A,MIR-525 | 158 | 4.34 | 10 | 0.012 | 0.122 | KMT2A,BSN,SEMA6D,MSH5,CADM |
| YWATTWNNRGCT_UNKNOWN | 70 | 1.92 | 6 | 0.0124 | 0.123 | PTCH1,CUL3,ZMIZ1,TANK,TSHZ3,I |
| CTACTGT,MIR-199A | 183 | 5.03 | 11 | 0.0124 | 0.123 | DGKI,FOXP1,GIGYF2,DPP4,FXR1,Z |
| V$HNF3ALPHA_Q6 | 208 | 5.72 | 12 | 0.0125 | 0.123 | FOXP1,TET2,MEF2C,CALD1,CADM |
| V$FREAC2_01 | 260 | 7.14 | 14 | 0.0129 | 0.125 | KMT2E,ENOX1,TET2,PTCH1,EXT1 |
| CTTTGCA,MIR-527 | 235 | 6.46 | 13 | 0.0132 | 0.127 | PRKD1,TNRC6A,ZFPM2,SEMA6D,N |
| V$PAX4_02 | 237 | 6.51 | 13 | 0.0141 | 0.134 | ARHGAP15,FOXP1,SEMA3A,TCF4 |
| V$LYF1_01 | 264 | 7.25 | 14 | 0.0146 | 0.134 | NRGN,ANXA9,GIGYF2,TCF4,AUTS |
| V$AFP1_Q6 | 264 | 7.25 | 14 | 0.0146 | 0.134 | FOXP1,PKP4,CNTNAP2,ZFPM2,AN |
| AAGCACT,MIR-520F | 238 | 6.54 | 13 | 0.0146 | 0.134 | KMT2A,KMT2E,DPP4,SLC22A23,SL |
| GAGACTG,MIR-452 | 94 | 2.58 | 7 | 0.0148 | 0.135 | GIGYF2,TCF4,SLC6A9,TCF20,CAL |
| V$HNF3_Q6 | 189 | 5.19 | 11 | 0.0154 | 0.139 | ATP2A2,SLC13A1,MEF2C,CALD1,C |
| WTTGKCTG_UNKNOWN | 516 | 14.2 | 23 | 0.0157 | 0.139 | KMT2E,RERE,FOXP1,SEMA3A,TCF |
| V$AHR_Q5 | 215 | 5.91 | 12 | 0.0159 | 0.139 | NRGN,PSMA5,USP4,PALB2,STAG1 |
| V$OCT1_04 | 241 | 6.62 | 13 | 0.016 | 0.139 | FOXP1,NCOA5,PKP4,ZFPM2,CACN |
| WGTTNNNNNAAA_UNKNOWN | 547 | 15 | 24 | 0.0164 | 0.142 | NREP,KMT2E,ENOX1,PSMA5,ESR |
| V$ER_Q6_01 | 269 | 7.39 | 14 | 0.0169 | 0.144 | GNAI1,TET2,PKP4,SLC6A9,PPM1E |
| V$FOXO4_01 | 243 | 6.68 | 13 | 0.017 | 0.144 | KMT2E,PTCH1,CACNA2D2,EXT1,C |
| V$HIF1_Q5 | 244 | 6.7 | 13 | 0.0176 | 0.147 | KMT2E,TET2,CACNA2D2,USP4,NT |

| | | | | | | |
|---|---|---|---|---|---|---|
| V$RSRFC4_01 | 245 | 6.73 | 13 | 0.0181 | 0.15 | SLC8A3,DGKI,FOXP1,PTCH1,CUL3 |
| CAGGTA_V$AREB6_01 | 792 | 21.8 | 32 | 0.0186 | 0.153 | NRGN,SLC8A3,NPAS3,FOXP1,ENC |
| V$ETS_Q4 | 247 | 6.79 | 13 | 0.0192 | 0.156 | KMT2A,ARHGAP15,FOXP1,SND1,I |
| V$TAL1BETAE47_01 | 248 | 6.81 | 13 | 0.0198 | 0.158 | KMT2E,FOXP1,GIGYF2,ZFPM2,SE |
| V$HMGIY_Q6 | 248 | 6.81 | 13 | 0.0198 | 0.158 | ARHGAP15,FOXP1,CNTN4,NEDD4 |
| V$GATA4_Q3 | 249 | 6.84 | 13 | 0.0204 | 0.159 | SLC8A3,AKAP6,SEMA3A,ESRRG,E |
| TGTTTGY_V$HNF3_Q6 | 738 | 20.3 | 30 | 0.0208 | 0.159 | MAGI1,KMT2A,RERE,FOXP1,SEM |
| CAGCTG_V$AP4_Q5 | 1520 | 41.9 | 55 | 0.0209 | 0.159 | SLC8A3,MAGI1,MYO18A,KMT2A,R |
| GGCACTT,MIR-519E | 124 | 3.41 | 8 | 0.0211 | 0.159 | NEDD4L,DDX3Y,FURIN,RBL2,SYN |
| V$PAX6_01 | 101 | 2.78 | 7 | 0.0212 | 0.159 | TCF4,BEND4,OLFM4,CACNA1C,M |
| V$PAX2_01 | 58 | 1.59 | 5 | 0.0213 | 0.159 | KMT2A,BEND4,ANKS1B,JMJD1C,H |
| V$FOXD3_01 | 199 | 5.47 | 11 | 0.0218 | 0.159 | KMT2E,ATP2A2,CACNA1C,CREB5, |
| V$NKX25_01 | 125 | 3.43 | 8 | 0.022 | 0.159 | TCF4,CREB5,MEF2C,MAML3,BCL1 |
| V$HFH4_01 | 200 | 5.5 | 11 | 0.0225 | 0.159 | NREP,MAGI1,NPAS3,ESRRG,ANK |
| TGTGTGA,MIR-377 | 200 | 5.5 | 11 | 0.0225 | 0.159 | KMT2A,EFL1,PRKD1,CACNA2D2,M |
| V$AP4_Q6 | 226 | 6.21 | 12 | 0.0225 | 0.159 | HDAC4,ZNF398,ZMIZ1,CACNB2,C |
| AATGTGA,MIR-23A,MIR-23B | 419 | 11.5 | 19 | 0.0228 | 0.159 | NRGN,MAGI1,MSL2,PKP4,TNRC6A |
| GTGCCAT,MIR-183 | 175 | 4.81 | 10 | 0.0228 | 0.159 | FOXP1,PKP4,SLC22A23,PPM1E,Z |
| GCATTTG,MIR-105 | 175 | 4.81 | 10 | 0.0228 | 0.159 | KMT2A,ERLIN1,PRKD1,NEDD4L,L |
| YGCANTGCR_UNKNOWN | 126 | 3.46 | 8 | 0.0229 | 0.159 | KMT2E,KIZ,FGFR1,CADM2,LRFN5, |
| TGCACTT,MIR-519C,MIR-519B,MIR-519A | 448 | 12.3 | 20 | 0.023 | 0.159 | RTN1,EFL1,TNRC6A,CUL3,ZFPM2, |

| | | | | | | |
|---|---|---|---|---|---|---|
| GCACTTT,MIR-17-5P,MIR-20A,MIR-106A,MIR-106B,MIR-20B,MIR-519D | 595 | 16.3 | 25 | 0.0234 | 0.16 | NPAS3,EFL1,SLC22A23,TNRC6A,S |
| V$P53_02 | 254 | 6.98 | 13 | 0.0235 | 0.16 | AKAP6,GRIN2A,ESRRG,MDK,GPD |
| MCAATNNNNNGCG_UNKNOWN | 81 | 2.23 | 6 | 0.0239 | 0.161 | RERE,ESRRG,ATP2A2,CNNM2,MA |
| TAATTA_V$CHX10_01 | 810 | 22.3 | 32 | 0.0247 | 0.165 | NREP,RPA3,SEMA3A,ESR2,CLSTN |
| V$RFX1_01 | 256 | 7.03 | 13 | 0.0249 | 0.165 | NRGN,SLC8A3,MSL2,ARHGAP15,E |
| GTGCAAT,MIR-25,MIR-32,MIR-92,MIR-363,MIR-367 | 311 | 8.55 | 15 | 0.0252 | 0.165 | MYO18A,BSN,CNTN4,FXR1,ATP2A |
| TATTATA,MIR-374 | 284 | 7.8 | 14 | 0.0255 | 0.165 | FOXP1,ESRRG,BANK1,TNRC6A,C |
| GTTTGTT,MIR-495 | 257 | 7.06 | 13 | 0.0256 | 0.165 | PTCH1,TNRC6A,SEMA6D,DDX3Y,S |
| GTGTTGA,MIR-505 | 105 | 2.88 | 7 | 0.0256 | 0.165 | MSL2,DDX3Y,NTM,DHRS11,PCDH |
| V$PAX2_02 | 258 | 7.09 | 13 | 0.0263 | 0.168 | RTN1,ESRRG,PTCH1,TCF4,CACN |
| GGCAGCT,MIR-22 | 232 | 6.37 | 12 | 0.0269 | 0.17 | MSL2,DGKI,FOXP1,TET2,DPP4,CU |
| V$ETS1_B | 259 | 7.12 | 13 | 0.027 | 0.17 | KMT2A,ARHGAP15,FOXP1,DYNC1 |
| CCCAGAG,MIR-326 | 155 | 4.26 | 9 | 0.0274 | 0.17 | SYNGAP1,PTCH1,TCF4,PPM1E,SE |
| CAGCAGG,MIR-370 | 155 | 4.26 | 9 | 0.0274 | 0.17 | KMT2A,HDAC4,GABBR1,DDX3Y,S |
| V$POU3F2_02 | 260 | 7.14 | 13 | 0.0278 | 0.171 | FOXP1,SEMA3A,PTCH1,MEF2C,C |
| RYAAAKNNNNNNTTGW_UNKNOWN | 84 | 2.31 | 6 | 0.028 | 0.171 | ENOX1,PKP4,CALD1,SOX5,JMJD1 |
| V$HNF6_Q6 | 234 | 6.43 | 12 | 0.0285 | 0.173 | RERE,SEMA3A,GRIN2A,CLSTN2,S |
| V$CDP_02 | 108 | 2.97 | 7 | 0.0293 | 0.174 | RPA3,TCF4,SLC6A9,MEF2C,SYT1, |
| GGGCATT,MIR-365 | 108 | 2.97 | 7 | 0.0293 | 0.174 | KMT2E,HDAC4,LDB1,CREB5,BCL1 |
| GAGCCTG,MIR-484 | 108 | 2.97 | 7 | 0.0293 | 0.174 | KMT2A,PTPRF,BCL11A,KDM4A,ZF |

| | | | | | | |
|---|---|---|---|---|---|---|
| V$EN1_01 | 109 | 2.99 | 7 | 0.0306 | 0.177 | UTY,TCF4,SLC6A9,CD55,TSHZ3,R |
| V$FOXJ2_01 | 184 | 5.06 | 10 | 0.0309 | 0.177 | NREP,ESRRG,ATP2A2,CREB5,MEI |
| V$OSF2_Q6 | 264 | 7.25 | 13 | 0.0309 | 0.177 | MSRA,KMT2E,WRN,TCF4,SLC6A9, |
| RNCTGNYNRNCTGNY_UNKNOWN | 86 | 2.36 | 6 | 0.0309 | 0.177 | ANXA9,TNRC6A,TCF20,ANKS1B,C |
| V$NKX3A_01 | 237 | 6.51 | 12 | 0.031 | 0.177 | GIGYF2,BEND4,ZFPM2,ATP2A2,CA |
| RTAAACA_V$FREAC2_01 | 919 | 25.3 | 35 | 0.0311 | 0.177 | MSL2,KMT2E,ENOX1,NCOA5,ESRI |
| GAGCTGG,MIR-337 | 159 | 4.37 | 9 | 0.0316 | 0.177 | KMT2A,DGKI,SBK1,ITGA11,RASGF |
| AGCATTA,MIR-155 | 134 | 3.68 | 8 | 0.0316 | 0.177 | DYNC1I1,HDAC4,CNTN4,TSHZ3,LC |
| TGACAGNY_V$MEIS1_01 | 827 | 22.7 | 32 | 0.0318 | 0.177 | NRGN,ESRRG,TET2,KIZ,PTCH1,BE |
| V$OCT1_01 | 266 | 7.31 | 13 | 0.0326 | 0.18 | FOXP1,TCF4,BEND4,CACNA1C,CA |
| TGATTTRY_V$GFI1_01 | 294 | 8.08 | 14 | 0.0329 | 0.181 | ESRRG,CLSTN2,ZFPM2,SND1,CRI |
| V$CEBP_Q2_01 | 267 | 7.34 | 13 | 0.0334 | 0.183 | NREP,FOXP1,BEND4,HDAC4,SEM, |
| V$SREBP1_02 | 88 | 2.42 | 6 | 0.0341 | 0.185 | NRGN,KMT2E,XRCC6,MEF2C,ZFY |
| V$RSRFC4_Q2 | 214 | 5.88 | 11 | 0.0345 | 0.186 | SLC8A3,DGKI,PTCH1,CUL3,ZFPM2 |
| V$ELK1_01 | 269 | 7.39 | 13 | 0.0352 | 0.187 | PSMA5,RBM6,CBLB,DMTF1,PRKA( |
| V$HNF4_01 | 269 | 7.39 | 13 | 0.0352 | 0.187 | PKP4,F2,HDAC4,ARFGEF2,SOX5,N |
| V$GATA3_01 | 242 | 6.65 | 12 | 0.0355 | 0.187 | NRGN,SLC8A3,MYO18A,ANKS1B,M |
| V$RORA1_01 | 242 | 6.65 | 12 | 0.0355 | 0.187 | SLC8A3,TNRC6A,MEF2C,CACNB2, |
| V$HNF1_C | 243 | 6.68 | 12 | 0.0365 | 0.19 | GRIN2A,F2,TCF4,ZFPM2,ANKS1B,I |
| CTATGCA,MIR-153 | 216 | 5.93 | 11 | 0.0366 | 0.19 | MAGI1,AKAP6,PTCH1,ZFPM2,BSN |
| V$HOX13_01 | 46 | 1.26 | 4 | 0.037 | 0.191 | GNAI1,TNRC6A,CALD1,PPP2R2B |
| V$NKX22_01 | 190 | 5.22 | 10 | 0.0372 | 0.191 | FOXP1,GIGYF2,ESRRG,PPM1E,CA |
| V$MEF2_Q6_01 | 244 | 6.7 | 12 | 0.0375 | 0.191 | SLC8A3,DGKI,RERE,FOXP1,HDAC |
| V$CEBP_01 | 272 | 7.47 | 13 | 0.0379 | 0.192 | NREP,KMT2D,FOXP1,SEMA3A,GIG |
| V$HNF1_01 | 245 | 6.73 | 12 | 0.0385 | 0.194 | SEMA3A,ZFPM2,OLFM4,ANKS1B,S |

| | | | | | | |
|---|---|---|---|---|---|---|
| V$LMO2COM_02 | 246 | 6.76 | 12 | 0.0395 | 0.198 | FBXL18,SLC8A3,MYO18A,ENOX1,[illegible] |
| TGCTGAY_UNKNOWN | 538 | 14.8 | 22 | 0.0414 | 0.206 | NREP,PDE1C,WRN,PKP4,TCF4,P[illegible] |
| GACAATC,MIR-219 | 143 | 3.93 | 8 | 0.0438 | 0.216 | SYNGAP1,MAPT,SDK1,FURIN,FBX[illegible] |
| V$HEN1_01 | 196 | 5.39 | 10 | 0.0444 | 0.219 | KMT2A,FOXP1,ESRRG,NT5DC2,P[illegible] |
| RRAGTTGT_UNKNOWN | 251 | 6.9 | 12 | 0.0449 | 0.219 | MAGI1,KMT2D,RERE,UTY,TCF4,C[illegible] |
| V$HEN1_02 | 198 | 5.44 | 10 | 0.047 | 0.225 | KMT2A,ESRRG,TCF4,CACNB2,AB[illegible] |
| V$HNF1_Q6 | 253 | 6.95 | 12 | 0.0472 | 0.225 | SEMA3A,F2,ZFPM2,OLFM4,SFTA2 |
| TTTGCAC,MIR-19A,MIR-19B | 516 | 14.2 | 21 | 0.0475 | 0.225 | FOXP1,RTN1,GRIN2A,TNRC6A,HD[illegible] |
| V$LEF1_Q2 | 226 | 6.21 | 11 | 0.048 | 0.225 | RERE,ESRRG,SLC6A9,COL11A1,C[illegible] |
| CACCAGC,MIR-138 | 226 | 6.21 | 11 | 0.048 | 0.225 | HDAC4,ZMIZ1,MGAT5B,ST3GAL3,[illegible] |
| AGCGCAG,MIR-191 | 13 | 0.357 | 2 | 0.0481 | 0.225 | MAGI1,ETF1 |
| V$OCT1_05 | 254 | 6.98 | 12 | 0.0483 | 0.225 | ESRRG,PPM1E,ATP2A2,CADM2,C[illegible] |
| AAAYWAACM_V$HFH4_01 | 254 | 6.98 | 12 | 0.0483 | 0.225 | NREP,SEMA3A,NCOA5,TCF4,ATP[illegible] |
| ATACTGT,MIR-144 | 199 | 5.47 | 10 | 0.0484 | 0.225 | ESRRG,SEMA6D,SNAP91,STAG1,[illegible] |
| V$MYB_Q6 | 255 | 7.01 | 12 | 0.0495 | 0.227 | KMT2D,MYO18A,KMT2A,KMT2E,G[illegible] |
| V$NGFIC_01 | 255 | 7.01 | 12 | 0.0495 | 0.227 | NRGN,PRPF3,GUCY1A2,ETF1,ME[illegible] |
| CAGCTTT,MIR-320 | 256 | 7.03 | 12 | 0.0507 | 0.232 | ESRRG,HDAC4,SEMA6D,MMP16,C[illegible] |
| AGCACTT,MIR-93,MIR-302A,MIR-302B,MIR-302C,MIR-302D,MIR-372,MIR-373,MIR-520E,MIR-520A,MIR-526B,MIR-520B,MIR-520C,MIR-520D | 343 | 9.42 | 15 | 0.0521 | 0.234 | NPAS3,KMT2A,SLC22A23,SLC6A9,[illegible] |

| | | | | | | |
|---|---|---|---|---|---|---|
| AAAGACA,MIR-511 | 202 | 5.55 | 10 | 0.0526 | 0.234 | ESRRG,TNRC6A,SEMA6D,DDX3Y, |
| ACTGTAG,MIR-139 | 123 | 3.38 | 7 | 0.053 | 0.234 | FOXP1,ESRRG,CACNA2D2,AUTS2 |
| V$HIF1_Q3 | 230 | 6.32 | 11 | 0.0532 | 0.234 | KMT2E,GRIN2A,TET2,PPM1E,MEF |
| V$IRF1_Q6 | 258 | 7.09 | 12 | 0.0532 | 0.234 | ESRRG,BANK1,TNRC6A,CALU,CA[illegible] |
| TTGGAGA,MIR-515-5P,MIR-519E | 149 | 4.09 | 8 | 0.0534 | 0.234 | MTHFD1L,KMT2A,CLSTN2,GABBR |
| CTTTGTA,MIR-524 | 433 | 11.9 | 18 | 0.0537 | 0.234 | KMT2E,GNAI1,FOXP1,GIGYF2,CA[illegible] |
| GCTGAGT,MIR-512-5P | 52 | 1.43 | 4 | 0.0541 | 0.234 | ZNF536,CTNNB1,NFIX,SRPK2 |
| CATGTAA,MIR-496 | 176 | 4.84 | 9 | 0.0542 | 0.234 | MSL2,ZNF536,CNTNAP2,SEMA6D, |
| V$ZID_01 | 259 | 7.12 | 12 | 0.0545 | 0.234 | ANXA9,FOXP1,THSD4,ESRRG,TC[illegible] |
| V$AR_Q6 | 259 | 7.12 | 12 | 0.0545 | 0.234 | KMT2D,RPA3,PTCH1,TNRC6A,CIT |
| GGCNNMSMYNTTG_UNKNOWN | 75 | 2.06 | 5 | 0.0552 | 0.234 | KMT2A,TET2,CUEDC2,STAG1,KC[illegible] |
| ACACTCC,MIR-122A | 75 | 2.06 | 5 | 0.0552 | 0.234 | TNRC6A,CPEB1,NEGR1,SLC39A8, |
| AAGGGAT,MIR-188 | 75 | 2.06 | 5 | 0.0552 | 0.234 | MEF2C,NTM,RNF111,PCDH9,BCL1 |
| V$AP2REP_01 | 178 | 4.89 | 9 | 0.0574 | 0.242 | KMT2A,GRIN2A,MAPT,FHIT,NT5D[illegible] |
| V$GFI1_01 | 262 | 7.2 | 12 | 0.0584 | 0.244 | TET2,TCF4,ZFPM2,CREB5,MEF2C |
| V$PAX4_01 | 262 | 7.2 | 12 | 0.0584 | 0.244 | MAGI1,GRIN2A,GIGYF2,MAPT,PT[illegible] |
| V$TCF11MAFG_01 | 207 | 5.69 | 10 | 0.06 | 0.25 | NRGN,FOXP1,PSMA5,PTCH1,DYN |
| ATCTTGC,MIR-31 | 77 | 2.12 | 5 | 0.0605 | 0.251 | ABCB9,CADPS,SEMA3F,GATAD2B |
| V$NKX61_01 | 236 | 6.48 | 11 | 0.0617 | 0.254 | RPA3,ESRRG,TCF4,CUL3,PPM1E,[illegible] |
| TTAYRTAA_V$E4BP4_01 | 265 | 7.28 | 12 | 0.0625 | 0.256 | NREP,DGKI,KMT2E,BSN,ARID1B,Z |
| V$FOXJ2_02 | 237 | 6.51 | 11 | 0.0631 | 0.257 | FOXP1,SEMA3A,FEZ1,ESRRG,TC[illegible] |
| V$GNCF_01 | 78 | 2.14 | 5 | 0.0633 | 0.257 | TNRC6A,MEF2C,BCL11A,ZFYVE1,[illegible] |
| V$PR_Q2 | 266 | 7.31 | 12 | 0.064 | 0.258 | RERE,SEMA3A,SSUH2,BEND4,CA[illegible] |
| V$PPARA_02 | 129 | 3.54 | 7 | 0.0652 | 0.262 | NRGN,SEMA3A,ESRRG,ANKS1B,F |
| V$SMAD3_Q6 | 239 | 6.57 | 11 | 0.0662 | 0.265 | ANXA9,FOXP1,ESRRG,ITIH3,SLC6 |

| | | | | | | |
|---|---|---|---|---|---|---|
| YAATNRNNNYNATT_UNKNOWN | 104 | 2.86 | 6 | 0.0668 | 0.266 | MAGI1,NPAS3,RPA3,CACNA1C,CR |
| ACACTAC,MIR-142-3P | 130 | 3.57 | 7 | 0.0673 | 0.267 | KMT2A,RERE,UTY,ATP2A2,PCGF3 |
| ATATGCA,MIR-448 | 212 | 5.82 | 10 | 0.0681 | 0.269 | GNAI1,DPP4,CNTN4,CACNA1C,AU |
| KCCGNSWTTT_UNKNOWN | 105 | 2.88 | 6 | 0.0693 | 0.27 | KMT2E,DMTF1,BCL11A,FOXO3,MI |
| CATTGTYY_V$SOX9_B1 | 358 | 9.84 | 15 | 0.0698 | 0.27 | MYO18A,TCF4,SLC6A9,NT5C2,FGI |
| V$STAT5A_04 | 213 | 5.85 | 10 | 0.0698 | 0.27 | PTCH1,SLC22A23,ZFPM2,ANKS1B |
| V$FOX_Q2 | 213 | 5.85 | 10 | 0.0698 | 0.27 | NREP,ESRRG,TET2,EXT1,ATP2A2 |
| V$YY1_02 | 242 | 6.65 | 11 | 0.071 | 0.272 | KMT2E,ENOX1,GIGYF2,TET2,TCF4 |
| V$EVI1_03 | 57 | 1.57 | 4 | 0.0711 | 0.272 | ESRRG,ZFPM2,CREB5,SRPK2 |
| TCCCCAC,MIR-491 | 57 | 1.57 | 4 | 0.0711 | 0.272 | SYNGAP1,SEMA6D,TRIM33,TRIOE |
| V$GATA1_01 | 244 | 6.7 | 11 | 0.0742 | 0.281 | SLC8A3,RERE,GRIN2A,XRCC6,AN |
| V$PTF1BETA_Q6 | 244 | 6.7 | 11 | 0.0742 | 0.281 | SLC8A3,MSL2,KMT2E,CITED1,CAC |
| V$GATA1_04 | 245 | 6.73 | 11 | 0.0759 | 0.285 | FBXL18,SLC8A3,ENOX1,BSN,ANKS |
| V$FOXO1_01 | 245 | 6.73 | 11 | 0.0759 | 0.285 | KMT2E,ESRRG,SEMA6D,EXT1,CAI |
| V$YY1_01 | 246 | 6.76 | 11 | 0.0776 | 0.288 | KMT2E,TCF4,RHOJ,CALD1,STAG1 |
| V$CIZ_01 | 246 | 6.76 | 11 | 0.0776 | 0.288 | SLC8A3,SEMA3A,PKP4,TCF4,ZFPI |
| GGCNRNWCTTYS_UNKNOWN | 83 | 2.28 | 5 | 0.0781 | 0.289 | NREP,ENOX1,CBLB,SND1,ELMO1 |
| AGTTCTC,MIR-146A,MIR-146B | 59 | 1.62 | 4 | 0.0786 | 0.29 | MMP16,SYT1,NOVA1,THRB |
| V$ISRE_01 | 247 | 6.79 | 11 | 0.0794 | 0.291 | AKAP6,PSMA5,PTCH1,CACNA2D2 |
| GGGAGGRR_V$MAZ_Q6 | 2270 | 62.5 | 73 | 0.0822 | 0.299 | ANXA9,KMT2A,MSL2,KMT2E,ARH( |
| ACATATC,MIR-190 | 60 | 1.65 | 4 | 0.0825 | 0.299 | TCF4,TNRC6A,BCL11A,PCDH9 |
| V$SRF_Q5_01 | 220 | 6.04 | 10 | 0.0825 | 0.299 | RERE,FOXP1,RSU1,PTCH1,LDB1,I |
| V$MAZ_Q6 | 193 | 5.3 | 9 | 0.0853 | 0.305 | RERE,MDK,F2,LRRC20,CTNND1,P |

| | | | | | | |
|---|---|---|---|---|---|---|
| CACTGCC,MIR-34A,MIR-34C,MIR-449 | 280 | 7.69 | 12 | 0.086 | 0.305 | MSL2,AKAP6,FOXP1,MAPT,PKP4,S |
| V$HMEF2_Q6 | 138 | 3.79 | 7 | 0.0862 | 0.305 | ZFPM2,SLC13A1,PPP1R16B,PPP2I |
| AGCGCTT,MIR-518F,MIR-518E,MIR-518A | 18 | 0.495 | 2 | 0.0863 | 0.305 | AUTS2,CPEB1 |
| TGCACGA,MIR-517A,MIR-517C | 18 | 0.495 | 2 | 0.0863 | 0.305 | BSN,DBN1 |
| GGTGAAG,MIR-412 | 61 | 1.68 | 4 | 0.0865 | 0.305 | PKP4,CUL3,BSN,FOXO3 |
| V$TAL1ALPHAE47_01 | 252 | 6.92 | 11 | 0.0884 | 0.307 | KMT2E,FOXP1,SND1,SEMA6D,MEI |
| V$NRF1_Q6 | 252 | 6.92 | 11 | 0.0884 | 0.307 | NRGN,MAGI1,MAPT,SGF29,SND1,. |
| V$FREAC7_01 | 195 | 5.36 | 9 | 0.0896 | 0.307 | FEZ1,PTCH1,SEMA6D,TSHZ3,PPP |
| V$PAX4_03 | 253 | 6.95 | 11 | 0.0902 | 0.307 | NRGN,NREP,NEDD4L,CITED1,FGF |
| V$CP2_02 | 253 | 6.95 | 11 | 0.0902 | 0.307 | RTN1,MDK,CITED1,CALD1,CTNND |
| V$SRY_01 | 224 | 6.15 | 10 | 0.0903 | 0.307 | KMT2A,ESRRG,ANKS1B,ATP2A2,L |
| V$LHX3_01 | 224 | 6.15 | 10 | 0.0903 | 0.307 | NREP,TNRC6A,OLFM4,PRKG1,TSI |
| V$LBP1_Q6 | 224 | 6.15 | 10 | 0.0903 | 0.307 | KMT2A,ZMIZ1,SOX5,CACNB2,ABC |
| AAAGGGA,MIR-204,MIR-211 | 224 | 6.15 | 10 | 0.0903 | 0.307 | KMT2A,RERE,ESRRG,DMTF1,CRE |
| ATGTAGC,MIR-221,MIR-222 | 140 | 3.85 | 7 | 0.0914 | 0.309 | KMT2A,MSL2,SBK1,KLC1,ZFPM2,S |
| V$AREB6_02 | 254 | 6.98 | 11 | 0.0921 | 0.311 | MYO18A,SEMA3A,RTN1,SPPL3,SL |
| V$FOXO4_02 | 255 | 7.01 | 11 | 0.0941 | 0.316 | NPAS3,KMT2A,ESR2,PTCH1,TNRC |
| GTTNYYNNGGTNA_UNKNOWN | 88 | 2.42 | 5 | 0.0947 | 0.317 | SLC8A3,WRN,RGS6,PPP1R16A,CS |
| V$SP1_Q6 | 256 | 7.03 | 11 | 0.096 | 0.319 | GRIN2A,TCF4,CTR9,ATP2A2,DMTI |
| V$GATA_Q6 | 198 | 5.44 | 9 | 0.0962 | 0.319 | NREP,ESRRG,PPM1E,BSN,ANKS1 |

| | | | | | | |
|---|---|---|---|---|---|---|
| GGCCAGT,MIR-193A,MIR-193B | 89 | 2.45 | 5 | 0.0982 | 0.323 | KMT2A,MMP16,PTPRF,NOVA1,PPI |
| AGGCACT,MIR-515-3P | 89 | 2.45 | 5 | 0.0982 | 0.323 | NEDD4L,SEMA6D,SYT1,SYNE1,GA |
| V$MEF2_02 | 228 | 6.26 | 10 | 0.0986 | 0.323 | NREP,SLC8A3,DGKI,FOXP1,ESRR |
| V$SP1_Q4_01 | 258 | 7.09 | 11 | 0.1 | 0.325 | GRIN2A,TCF4,CTR9,ATP2A2,DMTI |
| ATAAGCT,MIR-21 | 116 | 3.19 | 6 | 0.1 | 0.325 | SOX5,RNF111,GID4,GLCCI1,GATA |
| ACTTTAT,MIR-142-5P | 288 | 7.91 | 12 | 0.101 | 0.325 | RTN1,SLC22A23,ZFPM2,SOX5,STA |
| V$CEBP_C | 200 | 5.5 | 9 | 0.101 | 0.325 | RERE,FOXP1,ETF1,CREB5,RASGI |
| V$AMEF2_Q6 | 259 | 7.12 | 11 | 0.102 | 0.325 | SEMA3A,RTN1,ESRRG,BEND4,CR |
| V$MAF_Q6 | 259 | 7.12 | 11 | 0.102 | 0.325 | FBXL18,NPAS3,KMT2A,ESRRG,PP |
| V$EVI1_05 | 172 | 4.73 | 8 | 0.102 | 0.325 | RTN1,ENOX1,ESRRG,KIZ,ZFPM2,( |
| V$MEF2_01 | 144 | 3.96 | 7 | 0.102 | 0.325 | ESRRG,ZFPM2,MEF2C,GRIK1,ANK |
| TTANWNANTGGM_UNKNOWN | 65 | 1.79 | 4 | 0.103 | 0.327 | FOXP1,ESRRG,ANKS1B,PPP2R5C |
| ACTGTGA,MIR-27A,MIR-27B | 474 | 13 | 18 | 0.104 | 0.327 | NREP,KMT2A,NCOA5,PPM1E,SEM |
| V$IPF1_Q4 | 260 | 7.14 | 11 | 0.104 | 0.327 | NPAS3,GNAI1,BEND4,SEMA6D,ME |
| V$NKX25_02 | 262 | 7.2 | 11 | 0.108 | 0.339 | ESRRG,PPM1E,CNTN4,CACNA1C, |
| CAATGCA,MIR-33 | 92 | 2.53 | 5 | 0.109 | 0.34 | CNTN4,MMP16,CACNA1C,BCL11A |
| CTTTGT_V$LEF1_Q2 | 1970 | 54.2 | 63 | 0.11 | 0.341 | NREP,MAGI1,KMT2D,KMT2E,RERI |
| V$PU1_Q6 | 234 | 6.43 | 10 | 0.112 | 0.346 | KMT2D,KMT2E,CKB,MSH5,CREB5, |
| V$LMO2COM_01 | 264 | 7.25 | 11 | 0.112 | 0.347 | KMT2E,MAPT,DMTF1,KCNB1,CNN |
| CTTTGA_V$LEF1_Q2 | 1230 | 33.9 | 41 | 0.114 | 0.351 | KMT2E,RERE,SEMA3A,RTN1,GIGY |
| V$CDPCR3HD_01 | 236 | 6.48 | 10 | 0.116 | 0.355 | KMT2A,TET2,CLSTN2,EXT1,MGAT |
| V$PIT1_Q6 | 236 | 6.48 | 10 | 0.116 | 0.355 | SEMA3A,FEZ1,ESRRG,BEND4,ANI |

| | | | | | | |
|---|---|---|---|---|---|---|
| V$TCF1P_Q6 | 266 | 7.31 | 11 | 0.117 | 0.355 | KMT2E,SEMA3A,NCOA5,ESRRG,T |
| V$IK2_01 | 267 | 7.34 | 11 | 0.119 | 0.36 | NRGN,PTCH1,HDAC4,ZFPM2,CALI |
| V$EVI1_04 | 238 | 6.54 | 10 | 0.121 | 0.362 | RERE,KIZ,PTCH1,BEND4,EXT1,CA |
| V$CACCCBINDINGFACTOR_Q6 | 268 | 7.36 | 11 | 0.121 | 0.362 | MAPT,PTCH1,SLC6A9,LDB1,CADM |
| V$EVI1_06 | 22 | 0.604 | 2 | 0.121 | 0.362 | TCF4,CREB5 |
| GTCAACC,MIR-380-5P | 22 | 0.604 | 2 | 0.121 | 0.362 | KMT2A,RNF111 |
| V$CRX_Q4 | 269 | 7.39 | 11 | 0.123 | 0.365 | RPA3,ESRRG,MSH5,CREB5,DPYD |
| V$TITF1_Q3 | 239 | 6.57 | 10 | 0.124 | 0.365 | FOXP1,TCF4,TNRC6A,MEF2C,SOX |
| TCCATTKW_UNKNOWN | 239 | 6.57 | 10 | 0.124 | 0.365 | MYO18A,BSN,FGFR1,GMIP,CALD1 |
| GCAAGAC,MIR-431 | 45 | 1.24 | 3 | 0.126 | 0.369 | ESRRG,AMIGO1,PTPRF |
| V$FOXO1_02 | 240 | 6.59 | 10 | 0.126 | 0.369 | KMT2E,UTY,TET2,PTCH1,MDK,TN |
| GTATGAT,MIR-154,MIR-487 | 70 | 1.92 | 4 | 0.126 | 0.369 | ANKS1B,MAGI2,TSHZ3,RHOA |
| TCTCTCC,MIR-185 | 124 | 3.41 | 6 | 0.127 | 0.369 | MAGI1,SYNGAP1,BSN,LRFN5,NTR |
| TGGTGCT,MIR-29A,MIR-29B,MIR-29C | 521 | 14.3 | 19 | 0.128 | 0.372 | NREP,KMT2A,HDAC4,PPM1E,COL |
| GTCGATC,MIR-369-5P | 5 | 0.137 | 1 | 0.13 | 0.374 | RNF111 |
| GGCGGCA,MIR-371 | 5 | 0.137 | 1 | 0.13 | 0.374 | SYNGAP1 |
| CCTGTGA,MIR-513 | 125 | 3.43 | 6 | 0.13 | 0.374 | NPAS3,CNTNAP2,PPM1E,FGFR1,( |
| ACCTGTTG_UNKNOWN | 154 | 4.23 | 7 | 0.132 | 0.378 | ANXA9,MMP16,BCL11A,FOXO3,PII |
| V$HFH1_01 | 243 | 6.68 | 10 | 0.133 | 0.38 | SEMA6D,ATP2A2,CREB5,BCL11A,. |
| TGCACTG,MIR-148A,MIR-152,MIR-148B | 304 | 8.35 | 12 | 0.134 | 0.38 | KMT2A,ERLIN1,ESRRG,TNRC6A,Z |

| | | | | | | |
|---|---|---|---|---|---|---|
| YTAATTAA_V$LHX3_01 | 184 | 5.06 | 8 | 0.135 | 0.382 | NREP,SEMA3A,TNRC6A,PPM1E,LI |
| V$HTF_01 | 72 | 1.98 | 4 | 0.136 | 0.383 | SND1,TSHZ3,GBF1,BCL11B |
| V$GATA1_02 | 244 | 6.7 | 10 | 0.136 | 0.383 | SLC8A3,ESRRG,CACNA1C,CREB5 |
| V$FOXO3_01 | 245 | 6.73 | 10 | 0.139 | 0.385 | SEMA3A,UTY,PTCH1,TNRC6A,JMJ |
| V$SREBP1_Q6 | 245 | 6.73 | 10 | 0.139 | 0.385 | NRGN,ANXA9,KMT2A,KMT2E,MAM |
| V$YY1_Q6 | 245 | 6.73 | 10 | 0.139 | 0.385 | KMT2E,ENOX1,GIGYF2,MAPT,TET |
| V$SP1_Q2_01 | 245 | 6.73 | 10 | 0.139 | 0.385 | GRIN2A,CTR9,ATP2A2,LDB1,CADI |
| V$NFAT_Q6 | 246 | 6.76 | 10 | 0.141 | 0.389 | KMT2A,FOXP1,ENOX1,CNOT1,CRI |
| V$LFA1_Q6 | 246 | 6.76 | 10 | 0.141 | 0.389 | NRGN,NPAS3,GRIN2A,CKB,SOX5, |
| V$HAND1E47_01 | 277 | 7.61 | 11 | 0.142 | 0.391 | WRN,CREB5,MEF2C,CALD1,SOX5 |
| V$AP3_Q6 | 247 | 6.79 | 10 | 0.144 | 0.394 | NREP,PPM1E,ZFPM2,CALD1,CTNI |
| V$ICSBP_Q6 | 248 | 6.81 | 10 | 0.146 | 0.4 | NPAS3,BANK1,PKP4,HDAC4,SEMA |
| ATAGGAA,MIR-202 | 102 | 2.8 | 5 | 0.149 | 0.406 | KMT2A,MSL2,SNAP91,BCL11A,TRI |
| ATTACAT,MIR-380-3P | 102 | 2.8 | 5 | 0.149 | 0.406 | UTY,ZFPM2,MEF2C,GID4,ZNF800 |
| AAAYRNCTG_UNKNOWN | 374 | 10.3 | 14 | 0.151 | 0.407 | MYO18A,KMT2A,RTN1,ESRRG,DY |
| RGAGGAARY_V$PU1_Q6 | 502 | 13.8 | 18 | 0.151 | 0.407 | RBM6,ITGA11,DMTF1,CKB,MSH5,I |
| V$E47_02 | 250 | 6.87 | 10 | 0.152 | 0.407 | MYO18A,KMT2E,MMP16,MEF2C,JI |
| TAATGTG,MIR-323 | 160 | 4.4 | 7 | 0.152 | 0.407 | PKP4,TNRC6A,SEMA6D,NFIX,PCD |
| CGGTGTG,MIR-220 | 6 | 0.165 | 1 | 0.154 | 0.41 | THRB |
| V$FREAC3_01 | 251 | 6.9 | 10 | 0.154 | 0.41 | ESRRG,ARID1B,CALD1,MAML3,BC |
| V$T3R_Q6 | 251 | 6.9 | 10 | 0.154 | 0.41 | SLC8A3,RERE,BEND4,CACNA2D2, |
| V$SRF_Q4 | 221 | 6.07 | 9 | 0.156 | 0.411 | RERE,FOXP1,RSU1,KIZ,PTCH1,SI |
| V$EVI1_02 | 132 | 3.63 | 6 | 0.156 | 0.411 | ENOX1,ESRRG,ZFPM2,CREB5,SR |
| RYCACNNRNNRNCAG_UNKNOWN | 76 | 2.09 | 4 | 0.156 | 0.411 | SEMA3A,MDK,MEF2C,CSRNP3 |
| V$GATA1_05 | 283 | 7.78 | 11 | 0.157 | 0.413 | TNRC6A,ZFPM2,CREB5,SOX5,ESI |

| | | | | | | |
|---|---|---|---|---|---|---|
| TAATAAT, MIR-126 | 222 | 6.1 | 9 | 0.159 | 0.414 | ESRRG,HDAC4,SEMA6D,MMP16,G |
| ATGTTTC, MIR-494 | 162 | 4.45 | 7 | 0.159 | 0.414 | KMT2A,BEND4,ANKS1B,CACNA1C |
| V$E47_01 | 253 | 6.95 | 10 | 0.16 | 0.415 | KMT2E,FOXP1,NCOA5,CACNA2D2 |
| AAANWWTGC_UNKNOWN | 193 | 5.3 | 8 | 0.163 | 0.421 | FOXP1,MEF2C,SOX5,ANK3,MAML |
| CTTTAAR_UNKNOWN | 972 | 26.7 | 32 | 0.164 | 0.424 | NRGN,NREP,MAGI1,HLA-B,GNAI1, |
| V$AP4_Q6_01 | 255 | 7.01 | 10 | 0.165 | 0.426 | MYO18A,RPA3,GIGYF2,RSRC1,ZM |
| V$IK3_01 | 225 | 6.18 | 9 | 0.168 | 0.43 | SEMA3A,RTN1,PKP4,SLC6A9,GAB |
| V$TAL1BETAITF2_01 | 256 | 7.03 | 10 | 0.168 | 0.43 | FOXP1,ZFPM2,SEMA6D,CADM2,S |
| CAGTCAC,MIR-134 | 52 | 1.43 | 3 | 0.171 | 0.433 | ZFPM2,ETF1,AFF3 |
| GTAAACC,MIR-299-5P | 52 | 1.43 | 3 | 0.171 | 0.433 | HDAC4,CALU,ETF1 |
| YTTCCNNNGGAMR_UNKNOWN | 52 | 1.43 | 3 | 0.171 | 0.433 | MYO18A,ESRRG,FURIN |
| TGAGATT, MIR-216 | 107 | 2.94 | 5 | 0.172 | 0.433 | MMP16,ETF1,SRR,GATAD2B,BCL1 |
| AATGGAG,MIR-136 | 80 | 2.2 | 4 | 0.178 | 0.447 | MSL2,ESRRG,ETF1,PPA2 |
| CATRRAGC_UNKNOWN | 138 | 3.79 | 6 | 0.18 | 0.451 | COL11A1,ANK3,DHRS11,DPP8,AT |
| TCCAGAT,MIR-516-5P | 109 | 2.99 | 5 | 0.181 | 0.452 | MMP16,ATP2A2,DBN1,GID4,TLK1 |
| RYTTCCTG_V$ETS2_B | 1080 | 29.8 | 35 | 0.181 | 0.452 | KMT2D,MYO18A,KMT2A,ARHGAP |
| V$EGR2_01 | 199 | 5.47 | 8 | 0.183 | 0.453 | NRGN,MSL2,RERE,USP4,ATP2A2, |
| V$NF1_Q6 | 261 | 7.17 | 10 | 0.183 | 0.453 | KMT2E,RERE,CACNA1C,MEF2C,S |
| V$HP1SITEFACTOR_Q6 | 230 | 6.32 | 9 | 0.183 | 0.453 | NPAS3,ESRRG,TET2,DMTF1,CAD |
| GTGGGTGK_UNKNOWN | 293 | 8.05 | 11 | 0.184 | 0.454 | NRGN,KMT2A,MAPT,CTNND1,PPF |
| GTAAGAT,MIR-200A | 54 | 1.48 | 3 | 0.185 | 0.455 | KMT2E,FURIN,NOVA1 |
| WCTCNATGGY_UNKNOWN | 82 | 2.25 | 4 | 0.189 | 0.46 | PKP4,TCF4,JMJD1C,BCL11B |

| | | | | | | |
|---|---|---|---|---|---|---|
| TTCCGTT,MIR-191 | 29 | 0.797 | 2 | 0.189 | 0.46 | FOXP1,ATP2B2 |
| GTAGGCA,MIR-189 | 29 | 0.797 | 2 | 0.189 | 0.46 | SRPK2,PPP2R5C |
| V$MZF1_02 | 232 | 6.37 | 9 | 0.19 | 0.46 | KMT2E,FOXP1,PTCH1,ANKS1B,PT |
| CTACCTC,LET-7A,LET-7B,LET-7C,LET-7D,LET-7E,LET-7F,MIR-98,LET-7G,LET-7I | 391 | 10.7 | 14 | 0.19 | 0.46 | AKAP6,KMT2E,ATP2A2,PCGF3,AB |
| V$STAT5A_02 | 141 | 3.87 | 6 | 0.192 | 0.464 | FOXP1,PTCH1,CACNB2,PPP1R16I |
| YATGNWAAT_V$OCT_C | 360 | 9.89 | 13 | 0.193 | 0.465 | FOXP1,ENOX1,TCF4,CUL3,CADM2 |
| TAAYNRNNTCC_UNKNOWN | 172 | 4.73 | 7 | 0.195 | 0.469 | ESRRG,ZFPM2,ATP2A2,FURIN,ES |
| AACTGGA,MIR-145 | 234 | 6.43 | 9 | 0.196 | 0.469 | ERLIN1,SEMA3A,BEND4,BSN,NED |
| V$HFH8_01 | 203 | 5.58 | 8 | 0.196 | 0.469 | ESRRG,CREB5,ZFYVE1,PPP2R2B, |
| ACGCACA,MIR-210 | 8 | 0.22 | 1 | 0.2 | 0.476 | SYNGAP1 |
| V$HOXA4_Q2 | 267 | 7.34 | 10 | 0.201 | 0.477 | RPA3,RSRC1,HDAC4,PPM1E,CREI |
| V$MZF1_01 | 236 | 6.48 | 9 | 0.202 | 0.478 | KMT2E,GABBR1,ST3GAL3,BCL11A |
| GTGCCTT,MIR-506 | 727 | 20 | 24 | 0.203 | 0.478 | KMT2A,AKAP6,GNAI1,PRKD1,MDK |
| V$MSX1_01 | 174 | 4.78 | 7 | 0.203 | 0.478 | KMT2A,FOXP1,NT5DC2,SND1,KDM |
| V$MYCMAX_02 | 268 | 7.36 | 10 | 0.204 | 0.478 | KMT2A,PTCH1,TCF4,BEND4,FXR1 |
| TACTTGA,MIR-26A,MIR-26B | 300 | 8.24 | 11 | 0.204 | 0.478 | ERLIN1,AKAP6,RTN1,SLC22A23,TI |
| V$MEF2_03 | 238 | 6.54 | 9 | 0.209 | 0.481 | SLC8A3,DGKI,FOXP1,PTCH1,LRR( |
| V$POU1F1_Q6 | 238 | 6.54 | 9 | 0.209 | 0.481 | ESRRG,PKP4,OLFM4,ZMIZ1,CREB |
| V$MEIS1BHOXA9_01 | 145 | 3.98 | 6 | 0.209 | 0.481 | ESRRG,UTY,MDK,ANKS1B,GBF1,I |
| V$DR3_Q4 | 145 | 3.98 | 6 | 0.209 | 0.481 | NREP,PTCH1,NT5DC2,ATP2A2,PR |

| | | | | | | |
|---|---|---|---|---|---|---|
| CAGCCTC,MIR-485-5P | 145 | 3.98 | 6 | 0.209 | 0.481 | ZMIZ1,CADM2,SOX5,NGEF,ABCB9 |
| ATGCTGG,MIR-338 | 115 | 3.16 | 5 | 0.21 | 0.481 | MSL2,SEMA6D,NOVA1,MACROD2, |
| ATCATGA,MIR-433 | 115 | 3.16 | 5 | 0.21 | 0.481 | MSL2,ZMIZ1,ABCB9,GALNT1,BRW |
| V$RP58_01 | 207 | 5.69 | 8 | 0.211 | 0.481 | NRGN,FOXP1,GIGYF2,SND1,SEMA |
| GCTCTTG,MIR-335 | 86 | 2.36 | 4 | 0.211 | 0.481 | CALU,ETF1,PCDH9,SREK1IP1 |
| RYTGCNNRGNAAC_V$MIF1_01 | 86 | 2.36 | 4 | 0.211 | 0.481 | NRGN,NCOA5,PTCH1,GBF1 |
| V$MEIS1_01 | 239 | 6.57 | 9 | 0.212 | 0.483 | MAPT,ESRRG,BEND4,DYNC1I1,CC |
| SMTTTTGT_UNKNOWN | 402 | 11 | 14 | 0.218 | 0.493 | FOXP1,TCF4,ANKS1B,ZNF638,CN( |
| TGCTGCT,MIR-15A,MIR-16,MIR-15B,MIR-195,MIR-424,MIR-497 | 601 | 16.5 | 20 | 0.218 | 0.493 | KMT2A,ESRRG,TCAIM,PTCH1,DYN |
| CCANNAGRKGGC_UNKNOWN | 117 | 3.21 | 5 | 0.219 | 0.493 | ESR2,MDK,COL11A1,STAG1,CADF |
| V$MEIS1AHOXA9_01 | 117 | 3.21 | 5 | 0.219 | 0.493 | ESRRG,BEND4,ANKS1B,CREB5,N( |
| GCTNWTTGK_UNKNOWN | 306 | 8.41 | 11 | 0.222 | 0.498 | NREP,RERE,F2,SND1,ANKS1B,ATI |
| V$TCF4_Q5 | 242 | 6.65 | 9 | 0.223 | 0.498 | RERE,ANKS1B,SOX5,NFIX,ELMO1 |
| GGCNKCCATNK_UNKNOWN | 118 | 3.24 | 5 | 0.224 | 0.5 | KMT2E,NCOA5,TET2,STAG1,CSRN |
| AGTCAGC,MIR-345 | 60 | 1.65 | 3 | 0.228 | 0.506 | PRPF3,CACNB2,RNF111 |
| YKACATTT_UNKNOWN | 276 | 7.58 | 10 | 0.229 | 0.508 | ADAM22,DYNC1I1,MEF2C,ZNF568 |
| V$CEBPA_01 | 244 | 6.7 | 9 | 0.23 | 0.508 | NREP,RPA3,KMT2E,ANKS1B,ITGA |
| V$TBP_01 | 245 | 6.73 | 9 | 0.233 | 0.514 | ESRRG,CUL3,ANKS1B,CKB,SOX5, |
| KMCATNNWGGA_UNKNOWN | 90 | 2.47 | 4 | 0.235 | 0.515 | PTCH1,STAG1,GBF1,SREK1IP1 |

| | | | | | | |
|---|---|---|---|---|---|---|
| GTGTCAA,MIR-514 | 61 | 1.68 | 3 | 0.235 | 0.515 | KMT2A,ZNF536,BRWD1 |
| TCANNTGAY_V$SREBP1_01 | 475 | 13.1 | 16 | 0.235 | 0.515 | ANXA9,FOXP1,GIGYF2,SPPL3,SPN |
| YTAAYNGCT_UNKNOWN | 151 | 4.15 | 6 | 0.236 | 0.515 | ZFPM2,CITED1,PPP1R16B,PPP2R |
| GCAAAAA,MIR-129 | 183 | 5.03 | 7 | 0.239 | 0.52 | ESRRG,EFL1,SBK1,ANKS1B,CTNN |
| V$ZIC2_01 | 247 | 6.79 | 9 | 0.24 | 0.521 | FOXP1,UTY,PTCH1,PKP4,CKB,CA |
| TTCNRGNNNNTTC_V$HSF_Q6 | 152 | 4.18 | 6 | 0.24 | 0.521 | FOXP1,RTN1,RBM6,ZMIZ1,TNXB,Z |
| V$ROAZ_01 | 10 | 0.275 | 1 | 0.243 | 0.525 | PPP1R16A |
| CAGGTCC,MIR-492 | 63 | 1.73 | 3 | 0.25 | 0.534 | DBN1,AMBRA1,CSRNP3 |
| CCACACA,MIR-147 | 63 | 1.73 | 3 | 0.25 | 0.534 | CREB5,SEMA3F,TRIOBP |
| GCGNNANTTCC_UNKNOWN | 123 | 3.38 | 5 | 0.25 | 0.534 | NRGN,CACNA2D2,CBLB,SND1,GA |
| AGGAAGC,MIR-516-3P | 123 | 3.38 | 5 | 0.25 | 0.534 | SDK1,CREB5,SYT1,PPA2,BCL11B |
| WTGAAAT_UNKNOWN | 616 | 16.9 | 20 | 0.251 | 0.536 | SLC8A3,RERE,SEMA3A,GIGYF2,M |
| GTTATAT,MIR-410 | 93 | 2.56 | 4 | 0.253 | 0.537 | MSL2,RERE,CREB5,ATP2B2 |
| V$CEBP_Q3 | 251 | 6.9 | 9 | 0.254 | 0.538 | DOCK8,PPM1E,ANKS1B,CALD1,C |
| V$AP2_Q3 | 251 | 6.9 | 9 | 0.254 | 0.538 | MAPT,PPM1E,XRCC6,ETF1,FOXO |
| ACTGCCT,MIR-34B | 219 | 6.02 | 8 | 0.255 | 0.539 | MAGI1,ERLIN1,FOXP1,CACNA2D2 |
| V$AREB6_04 | 252 | 6.92 | 9 | 0.258 | 0.541 | KMT2A,ESRRG,WRN,PKP4,TCF4, |
| V$MTF1_Q4 | 252 | 6.92 | 9 | 0.258 | 0.541 | SEMA3A,CTNND1,CNNM2,CADPS, |
| V$HNF3B_01 | 221 | 6.07 | 8 | 0.263 | 0.55 | UTY,ZFPM2,CALD1,TSHZ3,NFIX,P |
| YRTCANNRCGC_UNKNOWN | 65 | 1.79 | 3 | 0.265 | 0.55 | XRCC6,MAML3,CDKAL1 |
| AGCTCCT,MIR-28 | 95 | 2.61 | 4 | 0.265 | 0.55 | MSL2,AMIGO1,NTM,PTPRF |
| V$CDX2_Q5 | 254 | 6.98 | 9 | 0.265 | 0.55 | NREP,FOXP1,ZFPM2,CACNA1C,S |
| V$GRE_C | 126 | 3.46 | 5 | 0.265 | 0.55 | SEMA3A,MEF2C,TANK,TMOD3,CA |

| | | | | | | |
|---|---|---|---|---|---|---|
| CATTTCA,MIR-203 | 287 | 7.89 | 10 | 0.266 | 0.55 | MAGI1,ZMIZ1,MEF2C,PRKG1,GLC |
| V$PITX2_Q2 | 255 | 7.01 | 9 | 0.269 | 0.552 | KMT2E,SLC13A1,CREB5,BCL11A,J |
| V$TEF_Q6 | 255 | 7.01 | 9 | 0.269 | 0.552 | DGKI,KMT2E,FOXP1,CREB5,CADM |
| V$SMAD_Q6 | 255 | 7.01 | 9 | 0.269 | 0.552 | NRGN,SLC8A3,KMT2E,PTCH1,DYN |
| YAATNANRNNNCAG_UNKNOWN | 66 | 1.81 | 3 | 0.272 | 0.557 | CALD1,CD55,TMOD3 |
| V$P53_DECAMER_Q2 | 256 | 7.03 | 9 | 0.272 | 0.557 | MSL2,AKAP6,GRIN2A,SLC6A9,SEM |
| V$PBX1_02 | 128 | 3.52 | 5 | 0.276 | 0.563 | RPA3,ZFPM2,MEF2C,MGAT5B,TLK |
| GGCAGTG,MIR-324-3P | 97 | 2.67 | 4 | 0.277 | 0.563 | FXR1,DMTF1,CKB,ST3GAL3 |
| V$CEBPB_02 | 258 | 7.09 | 9 | 0.28 | 0.565 | RPA3,FOXP1,ITGA11,CITED1,ARID |
| V$AP2_Q6 | 258 | 7.09 | 9 | 0.28 | 0.565 | SPPL3,PPM1E,BSN,LDB1,GUCY1A |
| V$STAT6_02 | 258 | 7.09 | 9 | 0.28 | 0.565 | CBLB,MEF2C,RHOJ,CTNNB1,TANK |
| TAGAACC,MIR-182 | 38 | 1.04 | 2 | 0.281 | 0.565 | GALNT1,ZNF800 |
| CAGGTG_V$E12_Q6 | 2480 | 68.3 | 73 | 0.281 | 0.565 | NRGN,NREP,MAGI1,PDE1C,ANXA |
| TTCYRGAA_UNKNOWN | 326 | 8.96 | 11 | 0.286 | 0.57 | NRGN,SEMA3A,RTN1,MAPT,CNTN |
| GCCATNTTG_V$YY1_Q6 | 427 | 11.7 | 14 | 0.287 | 0.57 | FBXL18,KMT2E,FOXP1,ENOX1,GI |
| V$OCT1_07 | 162 | 4.45 | 6 | 0.287 | 0.57 | ESRRG,LRFN5,NFIX,BCL11B,HDA |
| V$ALPHACP1_01 | 260 | 7.14 | 9 | 0.287 | 0.57 | NRGN,NT5DC2,ATP2A2,CALD1,ES |
| V$STAT_Q6 | 260 | 7.14 | 9 | 0.287 | 0.57 | RTN1,TET2,BSN,CALD1,ANK3,PPF |
| GTGCAAA,MIR-507 | 131 | 3.6 | 5 | 0.292 | 0.578 | ESRRG,SEMA6D,BCL11A,PCDH9, |
| GGGTGGRR_V$PAX4_03 | 1290 | 35.6 | 39 | 0.294 | 0.579 | NREP,MYO18A,KMT2A,KMT2E,MA |
| V$CEBPB_01 | 262 | 7.2 | 9 | 0.295 | 0.579 | NREP,FOXP1,PTCH1,FHIT,ARID1E |
| V$E12_Q6 | 262 | 7.2 | 9 | 0.295 | 0.579 | NEDD4L,PRKAG2,KCNB1,CNNM2, |
| CTCTATG,MIR-368 | 40 | 1.1 | 2 | 0.301 | 0.588 | MEF2C,PARD3B |
| V$ARP1_01 | 165 | 4.53 | 6 | 0.301 | 0.588 | ESRRG,PPM1E,NFIX,SORBS1,MEI |

| | | | | | | |
|---|---|---|---|---|---|---|
| V$NFY_Q6_01 | 264 | 7.25 | 9 | 0.302 | 0.588 | NT5DC2,ATP2A2,CADM2,KCNN2,E |
| YNTTTNNNANGCARM_UNKNOWN | 70 | 1.92 | 3 | 0.302 | 0.588 | GNAI1,CACNA1C,CSRNP3 |
| V$XBP1_01 | 133 | 3.65 | 5 | 0.303 | 0.588 | TCF4,SND1,CITED1,CADPS,BCL11 |
| TCCGTCC,MIR-184 | 13 | 0.357 | 1 | 0.304 | 0.59 | SLC6A9 |
| V$STAT5A_03 | 265 | 7.28 | 9 | 0.306 | 0.591 | RPA3,KIZ,TCF4,NEDD4L,ATP2A2,( |
| V$HEB_Q6 | 265 | 7.28 | 9 | 0.306 | 0.591 | MYO18A,MDK,ZNF398,ZMIZ1,SOX! |
| V$OCT1_06 | 266 | 7.31 | 9 | 0.31 | 0.595 | GALNT12,FOXP1,PKP4,BEND4,CR |
| AAGTCCA,MIR-422B,MIR-422A | 71 | 1.95 | 3 | 0.31 | 0.595 | NEK4,SRR,NTRK3 |
| ACATTCC,MIR-1,MIR-206 | 300 | 8.24 | 10 | 0.311 | 0.597 | FOXP1,HDAC4,BSN,GPD2,SEMA6I |
| V$HSF1_01 | 267 | 7.34 | 9 | 0.313 | 0.598 | RTN1,WRN,CACNA2D2,SGF29,GR |
| V$OCT1_B | 267 | 7.34 | 9 | 0.313 | 0.598 | FOXP1,BEND4,CACNA1C,MGAT5E |
| ACTWSNACTNY_UNKNOWN | 103 | 2.83 | 4 | 0.314 | 0.598 | KMT2D,FHIT,BCL11A,TMOD3 |
| V$CEBP_Q2 | 234 | 6.43 | 8 | 0.315 | 0.599 | FOXP1,TCF4,BEND4,HDAC4,ITGA |
| V$PR_02 | 136 | 3.74 | 5 | 0.319 | 0.603 | RERE,SEMA3A,SOX5,ELMO1,DOC |
| AGGGCAG,MIR-18A | 136 | 3.74 | 5 | 0.319 | 0.603 | MAPT,TET2,FURIN,ZFYVE1,CAMK |
| TTTGTAG,MIR-520D | 336 | 9.23 | 11 | 0.319 | 0.603 | MSL2,CNTNAP2,HDAC4,PPM1E,AT |
| V$GATA2_01 | 104 | 2.86 | 4 | 0.32 | 0.603 | SLC8A3,GRIN2A,MAML3,NEURL1 |
| V$EGR1_01 | 269 | 7.39 | 9 | 0.321 | 0.603 | NRGN,MSL2,PTCH1,PRPF3,MEF2( |
| V$GR_Q6 | 271 | 7.45 | 9 | 0.329 | 0.615 | RERE,PKP4,CNTN4,AUTS2,PPP2R |
| V$GR_01 | 204 | 5.61 | 7 | 0.329 | 0.615 | RERE,SEMA3A,KIZ,JMJD1C,ELMO |
| V$NCX_01 | 171 | 4.7 | 6 | 0.33 | 0.615 | KMT2E,CACNA2D2,CNOT1,PPP1R |
| GGGGCCC,MIR-296 | 74 | 2.03 | 3 | 0.332 | 0.615 | SYNGAP1,FOXP1,DBN1 |
| V$OLF1_01 | 272 | 7.47 | 9 | 0.333 | 0.615 | KMT2E,ESRRG,PTCH1,SLC6A9,SD |
| V$MMEF2_Q6 | 272 | 7.47 | 9 | 0.333 | 0.615 | DGKI,SEMA3A,RTN1,FXR1,GRIK1, |
| ATTCTTT,MIR-186 | 272 | 7.47 | 9 | 0.333 | 0.615 | CNTNAP2,TCF20,DDX3Y,CADM2,S |

| | | | | | | |
|---|---|---|---|---|---|---|
| V$AP4_Q5 | 273 | 7.5 | 9 | 0.336 | 0.62 | SLC8A3,HDAC4,SEMA6D,ZNF398,[illegible] |
| V$COREBINDINGFACTOR_Q6 | 273 | 7.5 | 9 | 0.336 | 0.62 | RPA3,ARHGAP15,ATP2A2,ETF1,S[illegible] |
| V$POU6F1_01 | 240 | 6.59 | 8 | 0.34 | 0.623 | RERE,FOXP1,GRIN2A,ZFPM2,CAC[illegible] |
| GAANYNYGACNY_UNKNOWN | 75 | 2.06 | 3 | 0.34 | 0.623 | BEND4,CARMIL1,ESRRB |
| V$AHRARNT_02 | 15 | 0.412 | 1 | 0.342 | 0.625 | SORCS3 |
| GTATTAT,MIR-369-3P | 207 | 5.69 | 7 | 0.343 | 0.625 | KMT2A,MSL2,PTCH1,ZFPM2,ST3G[illegible] |
| AGTCTTA,MIR-499 | 76 | 2.09 | 3 | 0.348 | 0.63 | ESRRG,SOX5,NOVA1 |
| V$CACBINDINGPROTEIN_Q6 | 242 | 6.65 | 8 | 0.348 | 0.63 | NRGN,NEDD4L,IVD,LRFN5,BCL11[illegible] |
| V$IRF_Q6 | 242 | 6.65 | 8 | 0.348 | 0.63 | KMT2A,SEMA3A,PSMA5,SEMA6D,[illegible] |
| V$ER_Q6 | 276 | 7.58 | 9 | 0.348 | 0.63 | KMT2E,ESRRG,CNNM2,ESRRB,M[illegible] |
| V$SRF_C | 211 | 5.8 | 7 | 0.36 | 0.651 | FOXP1,RSU1,PTCH1,RHOJ,CALD1[illegible] |
| V$PPARG_01 | 46 | 1.26 | 2 | 0.362 | 0.652 | NOVA1,CSRNP3 |
| CCAWYNNGAAR_UNKNOWN | 145 | 3.98 | 5 | 0.368 | 0.661 | PPM1E,HLA-DQA1,CTNND1,RBL2,[illegible] |
| V$FXR_IR1_Q6 | 112 | 3.08 | 4 | 0.37 | 0.664 | TNRC6A,ZMIZ1,SOX5,PPP2R2B |
| CRGAARNNNNCGA_UNKNOWN | 47 | 1.29 | 2 | 0.372 | 0.664 | BCL11B,PIK3R2 |
| TAGGTCA,MIR-192,MIR-215 | 47 | 1.29 | 2 | 0.372 | 0.664 | PKP4,ZNF536 |
| V$STAT1_03 | 248 | 6.81 | 8 | 0.373 | 0.665 | NRGN,ADAM22,OLFM4,ZNF568,G[illegible] |
| GGTGTGT,MIR-329 | 113 | 3.1 | 4 | 0.377 | 0.668 | ESRRG,SLC22A23,MEF2C,THRB |
| V$CMYB_01 | 249 | 6.84 | 8 | 0.377 | 0.668 | NRGN,KMT2D,NCOA5,RBM6,PPM[illegible] |
| TGACCTTG_V$SF1_Q6 | 249 | 6.84 | 8 | 0.377 | 0.668 | CKB,SDK1,MEF2C,CACNB2,PPP1[illegible] |
| V$STAT3_02 | 147 | 4.04 | 5 | 0.379 | 0.668 | KMT2A,NCOA5,TNRC6A,CALU,KC[illegible] |
| CYTAGCAAY_UNKNOWN | 147 | 4.04 | 5 | 0.379 | 0.668 | NRGN,KMT2D,PTCH1,TNRC6A,RG[illegible] |
| V$HSF2_01 | 250 | 6.87 | 8 | 0.381 | 0.67 | HLA-B,WRN,CACNA2D2,SGF29,M[illegible] |

| | | | | | | |
|---|---|---|---|---|---|---|
| V$MYOGNF1_01 | 48 | 1.32 | 2 | 0.382 | 0.67 | KMT2E,BCL11A |
| CGTSACG_V$PAX3_B | 148 | 4.07 | 5 | 0.384 | 0.674 | MTHFD1L,GIGYF2,BCL11A,RFT1,[illegible] |
| V$CREB_Q3 | 252 | 6.92 | 8 | 0.39 | 0.681 | FEZ1,ESRRG,PKP4,ANKS1B,BCL1[illegible] |
| GGGNRMNNYCAT_UNKNOWN | 82 | 2.25 | 3 | 0.393 | 0.685 | ETF1,ST3GAL3,ELMO1 |
| V$STAT_01 | 253 | 6.95 | 8 | 0.394 | 0.686 | MYO18A,TET2,GABBR1,CKB,NT5C[illegible] |
| V$FREAC4_01 | 150 | 4.12 | 5 | 0.395 | 0.686 | DGKI,TET2,CALD1,TMOD3,NAALA[illegible] |
| RYTGCNWTGGNR_UNKNOWN | 116 | 3.19 | 4 | 0.395 | 0.686 | SLC8A3,PPM1E,CACNB2,KCNN2 |
| V$HLF_01 | 254 | 6.98 | 8 | 0.398 | 0.687 | NREP,FOXP1,BSN,SEMA6D,ARID1[illegible] |
| V$E4F1_Q6 | 289 | 7.94 | 9 | 0.399 | 0.687 | PTCH1,PKP4,ANKS1B,ATP2A2,CK[illegible] |
| CTCCAAG,MIR-432 | 83 | 2.28 | 3 | 0.4 | 0.687 | RASGRP4,FURIN,TSHZ3 |
| V$MAZR_01 | 220 | 6.04 | 7 | 0.401 | 0.687 | KMT2E,MAPT,ETF1,GRIK1,PTPRF,[illegible] |
| TCTGATC,MIR-383 | 50 | 1.37 | 2 | 0.401 | 0.687 | MSL2,SOX5 |
| GGGATGC,MIR-324-5P | 50 | 1.37 | 2 | 0.401 | 0.687 | NTRK3,CAMKV |
| CTAWWWATA_V$RSRFC4_Q2 | 361 | 9.92 | 11 | 0.406 | 0.694 | SLC8A3,DGKI,RPA3,FOXP1,PTCH[illegible] |
| CTGYNNCTYTAA_UNKNOWN | 84 | 2.31 | 3 | 0.407 | 0.694 | NRGN,BCL11A,KCNN2 |
| TCTGGAC,MIR-198 | 84 | 2.31 | 3 | 0.407 | 0.694 | BSN,CREB5,BTN2A1 |
| V$MYOD_Q6_01 | 257 | 7.06 | 8 | 0.411 | 0.697 | CLSTN2,CTR9,ITGA11,JMJD1C,SE[illegible] |
| CGGAARNGGCNG_UNKNOWN | 51 | 1.4 | 2 | 0.411 | 0.697 | CTR9,ATP2A2 |
| STTTCRNTTT_V$IRF_Q6 | 188 | 5.17 | 6 | 0.414 | 0.699 | PSMA5,HDAC4,SOX5,BCL11A,NFI[illegible] |
| V$CETS1P54_01 | 258 | 7.09 | 8 | 0.415 | 0.699 | CTR9,CALU,DMTF1,PRKAG2,MEF[illegible] |
| V$AREB6_03 | 258 | 7.09 | 8 | 0.415 | 0.699 | PDE1C,MMP16,CTNND1,PPP1R16[illegible] |

| | | | | | | |
|---|---|---|---|---|---|---|
| V$ERR1_Q2 | 259 | 7.12 | 8 | 0.419 | 0.705 | TNRC6A,PPM1E,TTLL7,MEF2C,CA |
| V$EGR3_01 | 86 | 2.36 | 3 | 0.422 | 0.708 | USP4,SORCS3,HDAC9 |
| GATAAGR_V$GATA_C | 295 | 8.11 | 9 | 0.422 | 0.708 | MYO18A,PPM1E,GABBR1,ANKS1B |
| V$VDR_Q3 | 225 | 6.18 | 7 | 0.424 | 0.708 | NRGN,PTCH1,FSCN3,KCNB1,FUR |
| MGGAAGTG_V$GABP_B | 757 | 20.8 | 22 | 0.424 | 0.708 | ARHGAP15,FOXP1,CTR9,TNRC6A |
| V$TEF1_Q6 | 226 | 6.21 | 7 | 0.428 | 0.711 | MDK,SLC6A9,ZMIZ1,ANK3,PTPRF, |
| V$HFH3_01 | 191 | 5.25 | 6 | 0.428 | 0.711 | ATP2A2,MEF2C,CADM2,NOVA1,SL |
| CCAWWNAAGG_V$SRF_Q4 | 87 | 2.39 | 3 | 0.43 | 0.711 | PTCH1,TCF4,CALD1 |
| TTGCACT,MIR-130A,MIR-301,MIR-130B | 403 | 11.1 | 12 | 0.43 | 0.711 | KMT2A,RTN1,TNRC6A,CUL3,ZFPM |
| V$TST1_01 | 262 | 7.2 | 8 | 0.432 | 0.713 | KMT2E,RTN1,ESRRG,CACNA2D2,( |
| V$HNF4_Q6 | 263 | 7.23 | 8 | 0.436 | 0.717 | KMT2A,MMP16,XRCC6,CACNB2,T! |
| ACAGGGT,MIR-10A,MIR-10B | 123 | 3.38 | 4 | 0.439 | 0.717 | ESRRG,NFIX,GALNT1,BRWD1 |
| RACTNNRTTTNC_UNKNOWN | 123 | 3.38 | 4 | 0.439 | 0.717 | ADAM22,SGF29,TNRC6A,ZNF568 |
| GTGACTT,MIR-224 | 158 | 4.34 | 5 | 0.439 | 0.717 | RERE,SPPL3,CREB5,DDX3Y,AFF3 |
| CAGCACT,MIR-512-3P | 158 | 4.34 | 5 | 0.439 | 0.717 | KMT2A,CADM2,SGSM2,ATP2B2,ZN |
| CCCACAT,MIR-299-3P | 54 | 1.48 | 2 | 0.439 | 0.717 | TCF4,BCL11A |
| V$STAT6_01 | 264 | 7.25 | 8 | 0.44 | 0.717 | KIZ,TCF4,NEDD4L,ATP2A2,CADM2 |
| V$GATA_C | 266 | 7.31 | 8 | 0.448 | 0.726 | ESRRG,PPM1E,GABBR1,ANKS1B,( |
| V$STAT4_01 | 266 | 7.31 | 8 | 0.448 | 0.726 | MSL2,PKP4,CARMIL1,MEF2C,RHO |
| TGCCTTA,MIR-124A | 552 | 15.2 | 16 | 0.45 | 0.726 | RERE,PRKD1,CACNA2D2,HDAC4,! |
| ACCATTT,MIR-522 | 160 | 4.4 | 5 | 0.45 | 0.726 | MSL2,FOXP1,PPM1E,STAG1,PRK( |
| V$SPZ1_01 | 231 | 6.35 | 7 | 0.45 | 0.726 | LDB1,CADM2,BCL11A,KCNN2,NTR |

| | | | | | | |
|---|---|---|---|---|---|---|
| V$PAX3_B | 90 | 2.47 | 3 | 0.451 | 0.726 | SEMA3A,CNNM2,BCL11B |
| TTGGGAG,MIR-150 | 90 | 2.47 | 3 | 0.451 | 0.726 | FOXP1,PKP4,ETF1 |
| RGAANNTTC_V$HSF1_01 | 446 | 12.3 | 13 | 0.454 | 0.728 | HLA-B,SEMA3A,RTN1,CACNA2D2, |
| V$CDPCR3_01 | 56 | 1.54 | 2 | 0.458 | 0.733 | NPAS1,HDAC4 |
| V$STAT3_01 | 22 | 0.604 | 1 | 0.458 | 0.733 | ELMO1 |
| V$TEL2_Q6 | 233 | 6.4 | 7 | 0.459 | 0.733 | ARHGAP15,CTR9,TNRC6A,DMTF1 |
| CCTNTMAGA_UNKNOWN | 127 | 3.49 | 4 | 0.463 | 0.737 | SND1,ADTRP,NTRK3,BCL11B |
| ACTAYRNNNCCCR_UNKNOWN | 449 | 12.3 | 13 | 0.464 | 0.737 | KMT2D,ADAM22,WRN,SGF29,SND |
| V$CDP_01 | 92 | 2.53 | 3 | 0.466 | 0.737 | RERE,ESRRG,NFIX |
| V$LXR_DR4_Q3 | 92 | 2.53 | 3 | 0.466 | 0.737 | FOXP1,HDAC4,MEF2C |
| V$AP2_Q6_01 | 272 | 7.47 | 8 | 0.473 | 0.748 | KMT2E,RBM6,PPM1E,LDB1,DBN1, |
| GTACAGG,MIR-486 | 58 | 1.59 | 2 | 0.476 | 0.748 | MAML3,AFF3 |
| ACTGAAA,MIR-30A-3P,MIR-30E-3P | 201 | 5.52 | 6 | 0.477 | 0.748 | FOXP1,FXR1,MEF2C,GALNT1,NEG |
| GGGACCA,MIR-133A,MIR-133B | 201 | 5.52 | 6 | 0.477 | 0.748 | SLC8A3,NCOA5,DDX3Y,CTTNBP2, |
| V$OCT1_Q5_01 | 273 | 7.5 | 8 | 0.477 | 0.748 | FOXP1,ATP2A2,MGAT5B,CADM2,F |
| V$CDPCR1_01 | 130 | 3.57 | 4 | 0.481 | 0.752 | KMT2A,EXT1,MGAT5B,NOVA1 |
| V$ETS2_B | 274 | 7.53 | 8 | 0.481 | 0.752 | DMTF1,CKB,FURIN,FOXO3,ELMO1 |
| V$AR_03 | 59 | 1.62 | 2 | 0.485 | 0.754 | NRGN,RERE |
| GTCAGGA,MIR-378 | 59 | 1.62 | 2 | 0.485 | 0.754 | NFIX,ATP2B2 |
| V$EGR_Q6 | 275 | 7.56 | 8 | 0.485 | 0.754 | PTCH1,CALU,PRPF3,KCNB1,PPP1 |
| AACYNNNNTTCCS_UNKNOWN | 95 | 2.61 | 3 | 0.487 | 0.754 | ADAM22,SGF29,GATB |
| CGCTGCT,MIR-503 | 24 | 0.659 | 1 | 0.488 | 0.754 | BSN |

| | | | | | | |
|---|---|---|---|---|---|---|
| AGTCTAG,MIR-151 | 24 | 0.659 | 1 | 0.488 | 0.754 | NTM |
| CCTGCTG,MIR-214 | 240 | 6.59 | 7 | 0.49 | 0.756 | MYO18A,MAPT,MRM2,NEDD4L,AN |
| AACTGAC,MIR-223 | 96 | 2.64 | 3 | 0.494 | 0.76 | MMP16,MEF2C,FOXO3 |
| V$RFX1_02 | 278 | 7.64 | 8 | 0.498 | 0.765 | SLC8A3,SPAG16,HDAC4,PPM1E,Z |
| V$GCM_Q2 | 242 | 6.65 | 7 | 0.499 | 0.766 | MAGI1,EXT1,CADM2,KCNB1,CNNM |
| V$GR_Q6_01 | 279 | 7.67 | 8 | 0.502 | 0.767 | KMT2D,RSRC1,TET2,CALD1,ANK3 |
| V$MEF2_04 | 25 | 0.687 | 1 | 0.502 | 0.767 | DGKI |
| ACCAAAG,MIR-9 | 499 | 13.7 | 14 | 0.507 | 0.773 | FOXP1,CNTN4,SEMA6D,MMP16,AI |
| V$ELF1_Q6 | 244 | 6.7 | 7 | 0.508 | 0.773 | DMTF1,MSH5,GMIP,CTNND1,ELMO |
| ATAACCT,MIR-154 | 62 | 1.7 | 2 | 0.511 | 0.775 | PPM1E,CADM2 |
| RYTAAWNNNTGAY_UNKNOWN | 62 | 1.7 | 2 | 0.511 | 0.775 | ZFPM2,CREB5 |
| V$NFMUE1_Q6 | 245 | 6.73 | 7 | 0.512 | 0.775 | GIGYF2,TCF4,EXT1,STAG1,GBF1,I |
| GTTRYCATRR_UNKNOWN | 172 | 4.73 | 5 | 0.513 | 0.775 | HDAC4,PPM1E,LDB1,ELMO1,CAMI |
| AGGTGCA,MIR-500 | 99 | 2.72 | 3 | 0.514 | 0.776 | ESRRG,DDX3Y,TSHZ3 |
| V$TAXCREB_02 | 26 | 0.714 | 1 | 0.516 | 0.777 | BCL11A |
| V$FOXM1_01 | 246 | 6.76 | 7 | 0.516 | 0.777 | SEMA3A,CREB5,MEF2C,SOX5,PCI |
| ACAWNRNSRCGG_UNKNOWN | 63 | 1.73 | 2 | 0.52 | 0.778 | CKB,ZNF638 |
| V$TGIF_01 | 247 | 6.79 | 7 | 0.521 | 0.778 | NRGN,DYNC1I1,CLSTN2,DYSF,AN |
| V$E2F1_Q3_01 | 247 | 6.79 | 7 | 0.521 | 0.778 | NRGN,KMT2E,GRIN2A,EIF1AY,STA |
| V$POU3F2_01 | 100 | 2.75 | 3 | 0.521 | 0.778 | NOVA1,BCL11B,HDAC9 |
| V$AHR_01 | 64 | 1.76 | 2 | 0.528 | 0.786 | BCL11A,NTRK3 |
| CTAGGAA,MIR-384 | 64 | 1.76 | 2 | 0.528 | 0.786 | SYNGAP1,ESRRG |
| TCCCRNNRTGC_UNKNOWN | 213 | 5.85 | 6 | 0.533 | 0.792 | KMT2D,EXOC4,RBM6,BSN,CSRNP |

| | | | | | | |
|---|---|---|---|---|---|---|
| ATGGYGGA_UNKNOWN | 102 | 2.8 | 3 | 0.535 | 0.792 | CACNA2D2,EXT1,BCL11B |
| YRCCAKNNGNCGC_UNKNOWN | 65 | 1.79 | 2 | 0.537 | 0.794 | GRIN2A,PTPRF |
| V$MYCMAX_03 | 252 | 6.92 | 7 | 0.542 | 0.801 | GIGYF2,SPPL3,UTY,PTCH1,SPNS1 |
| CAAGGAT,MIR-362 | 66 | 1.81 | 2 | 0.545 | 0.802 | CNTN4,BCL11A |
| AGGAGTG,MIR-483 | 66 | 1.81 | 2 | 0.545 | 0.802 | NTRK3,AMBRA1 |
| V$HNF4_01_B | 253 | 6.95 | 7 | 0.546 | 0.803 | ARFGEF2,CNOT1,BCL11A,JMJD1C |
| ATACCTC,MIR-202 | 179 | 4.92 | 5 | 0.548 | 0.804 | CALU,CALD1,SYT1,GALNT1,SREK |
| AACWWCAANK_UNKNOWN | 142 | 3.9 | 4 | 0.55 | 0.805 | FGFR1,SOX5,DOCK4,AFF3 |
| V$NFY_01 | 254 | 6.98 | 7 | 0.55 | 0.805 | KMT2E,ATP2A2,LRRC20,ELMO1,W |
| GACAGGG,MIR-339 | 67 | 1.84 | 2 | 0.553 | 0.807 | SYNGAP1,NOVA1 |
| V$ZIC3_01 | 255 | 7.01 | 7 | 0.555 | 0.808 | FOXP1,ESRRG,PTCH1,CTNND1,P |
| V$MIF1_01 | 181 | 4.97 | 5 | 0.558 | 0.812 | NRGN,NCOA5,PTCH1,CBLB,SOX5 |
| TGACATY_UNKNOWN | 665 | 18.3 | 18 | 0.561 | 0.814 | PDE1C,NPAS3,TET2,OLFM4,ANKS |
| V$SF1_Q6 | 257 | 7.06 | 7 | 0.563 | 0.816 | TNRC6A,PPM1E,CKB,TTLL7,CACN |
| GAGCCAG,MIR-149 | 145 | 3.98 | 4 | 0.567 | 0.817 | EXT1,ATP2A2,NTRK3,TNXB |
| RAAGNYNNCTTY_UNKNOWN | 145 | 3.98 | 4 | 0.567 | 0.817 | ESRRG,DYSF,OLFM4,BCL11A |
| V$SREBP_Q3 | 258 | 7.09 | 7 | 0.567 | 0.817 | NRGN,KMT2E,PPM1E,ANKS1B,BC |
| GTTAAAG,MIR-302B | 69 | 1.9 | 2 | 0.569 | 0.818 | WNT3,PCDH9 |
| ATGCTGC,MIR-103,MIR-107 | 221 | 6.07 | 6 | 0.57 | 0.818 | ZFPM2,FURIN,BCL11A,PDE4B,NO |
| V$ATF_01 | 259 | 7.12 | 7 | 0.571 | 0.819 | PFAS,SND1,ETF1,CALD1,KCNN2,P |
| GTTGNYNNRGNAAC_UNKNOWN | 108 | 2.97 | 3 | 0.573 | 0.821 | SPAG16,ZFYVE1,ELMO1 |
| TTCYNRGAA_V$STAT5B_01 | 335 | 9.2 | 9 | 0.575 | 0.822 | MYO18A,TET2,SEMA6D,NT5C2,AN |

| | | | | | | |
|---|---|---|---|---|---|---|
| CACGTTT,MIR-302A | 31 | 0.852 | 1 | 0.579 | 0.824 | NREP |
| V$ZIC1_01 | 261 | 7.17 | 7 | 0.579 | 0.824 | MSL2,GIGYF2,PTCH1,DMTF1,CTN |
| RACCACAR_V$AML_Q6 | 261 | 7.17 | 7 | 0.579 | 0.824 | MSRA,RPA3,CITED1,FGFR1,ETF1, |
| TGACCTY_V$ERR1_Q2 | 1040 | 28.7 | 28 | 0.58 | 0.824 | NREP,ANXA9,KMT2A,DGKI,KMT2E |
| V$AP4_01 | 262 | 7.2 | 7 | 0.583 | 0.824 | FOXP1,COL11A1,CACNB2,BCL11A |
| V$MAX_01 | 262 | 7.2 | 7 | 0.583 | 0.824 | KMT2A,TCF4,BEND4,SPNS1,GUCY |
| V$NFY_Q6 | 262 | 7.2 | 7 | 0.583 | 0.824 | HLA-DMA,PTCH1,CBLB,HLA-DRB5 |
| GCAAGGA,MIR-502 | 71 | 1.95 | 2 | 0.585 | 0.824 | BCL11A,MACROD2 |
| V$CHX10_01 | 225 | 6.18 | 6 | 0.587 | 0.824 | PPM1E,SND1,LRFN5,PPP1R16A,P |
| TTTTGAG,MIR-373 | 225 | 6.18 | 6 | 0.587 | 0.824 | GNAI1,PKP4,TNRC6A,PPM1E,PPP |
| TGANNYRGCA_V$TCF11MAFG_01 | 301 | 8.27 | 8 | 0.588 | 0.824 | SLC8A3,DYNC1I1,ANKS1B,TTLL7, |
| GCCNNNWTAAR_UNKNOWN | 149 | 4.09 | 4 | 0.589 | 0.824 | MAGI1,DGKI,PTCH1,SOX5 |
| V$AP1_Q6_01 | 264 | 7.25 | 7 | 0.592 | 0.826 | ESRRG,TNRC6A,GABBR1,ANK3,S |
| GTCNYYATGR_UNKNOWN | 111 | 3.05 | 3 | 0.592 | 0.826 | PKP4,PPM1E,JMJD1C |
| V$MYOD_01 | 265 | 7.28 | 7 | 0.596 | 0.828 | CACNA2D2,CACNB2,KCNB1,CNN |
| V$OCT_Q6 | 265 | 7.28 | 7 | 0.596 | 0.828 | FOXP1,ATP2A2,MGAT5B,CADM2,F |
| V$AML_Q6 | 266 | 7.31 | 7 | 0.6 | 0.833 | GALNT12,PTCH1,ATP2A2,ETF1,C |
| GGTAACC,MIR-409-5P | 33 | 0.907 | 1 | 0.602 | 0.834 | TSHZ3 |
| V$MYCMAX_B | 268 | 7.36 | 7 | 0.607 | 0.841 | PRKAG2,SDK1,ETF1,LRFN5,BCL1 |
| GCTTGAA,MIR-498 | 114 | 3.13 | 3 | 0.61 | 0.841 | PTCH1,CADM2,PDE4B |
| V$VDR_Q6 | 269 | 7.39 | 7 | 0.611 | 0.841 | GRIN2A,ESRRG,PTCH1,SOX5,PPF |
| V$ATF1_Q6 | 231 | 6.35 | 6 | 0.613 | 0.841 | FOXP1,ESRRG,PFAS,CALD1,NOV |
| V$PAX5_01 | 154 | 4.23 | 4 | 0.615 | 0.841 | CALD1,MAML3,ZNF800,TLK1 |

| | | | | | | |
|---|---|---|---|---|---|---|
| RGTTAMWNATT_V$HNF1_01 | 75 | 2.06 | 2 | 0.615 | 0.841 | KMT2A,SFTA2 |
| V$OCT1_Q6 | 270 | 7.42 | 7 | 0.615 | 0.841 | ESRRG,TCF4,ATP2A2,CADM2,SO[illegible] |
| V$COMP1_01 | 115 | 3.16 | 3 | 0.616 | 0.841 | EXT1,CREB5,CALD1 |
| ATGCAGT,MIR-217 | 115 | 3.16 | 3 | 0.616 | 0.841 | BCL11A,GATAD2B,NOVA1 |
| V$NFKB_Q6_01 | 232 | 6.37 | 6 | 0.617 | 0.841 | DGKI,RASGRP4,SOX5,PPP1R13B, |
| V$NMYC_01 | 271 | 7.45 | 7 | 0.619 | 0.843 | GRIN2A,TET2,BEND4,ZCCHC7,FO |
| TGGNNNNNNKCCAR_UNKNOWN | 424 | 11.6 | 11 | 0.62 | 0.843 | KMT2E,RERE,RTN1,ZFPM2,ATP2A |
| GCACCTT,MIR-18A,MIR-18B | 117 | 3.21 | 3 | 0.628 | 0.852 | TSHZ3,CSRNP3,TRIOBP |
| V$LXR_Q3 | 77 | 2.12 | 2 | 0.629 | 0.852 | SND1,FAM193A |
| GGARNTKYCCA_UNKNOWN | 78 | 2.14 | 2 | 0.636 | 0.86 | SLC6A9,NTRK3 |
| V$SP1_01 | 237 | 6.51 | 6 | 0.638 | 0.86 | GRIN2A,GMIP,CADM2,SRR,GBF1,[illegible] |
| V$NFKAPPAB65_01 | 237 | 6.51 | 6 | 0.638 | 0.86 | WRN,RASGRP4,SOX5,KCNN2,PP[illegible] |
| V$ZF5_B | 239 | 6.57 | 6 | 0.646 | 0.869 | NCOA5,ZNF398,ETF1,PPA2,TRIM3 |
| AGGGCCA,MIR-328 | 80 | 2.2 | 2 | 0.65 | 0.872 | DYNC1I1,ST3GAL3 |
| TAANNYSGCG_UNKNOWN | 80 | 2.2 | 2 | 0.65 | 0.872 | CNOT1,RHOA |
| V$SRF_Q6 | 241 | 6.62 | 6 | 0.654 | 0.874 | RERE,FOXP1,RSU1,PTCH1,RHOJ, |
| V$PAX8_01 | 38 | 1.04 | 1 | 0.654 | 0.874 | JMJD1C |
| V$TFIII_Q6 | 205 | 5.63 | 5 | 0.668 | 0.891 | ESRRG,LDB1,BCL11A,NFIX,PCDH[illegible] |
| V$MYOD_Q6 | 245 | 6.73 | 6 | 0.669 | 0.891 | MAGI1,KMT2E,PKP4,NEDD4L,PRK |
| V$NFY_C | 245 | 6.73 | 6 | 0.669 | 0.891 | ESRRG,CBLB,CNNM2,ELMO1,WN[illegible] |
| V$IRF2_01 | 125 | 3.43 | 3 | 0.672 | 0.892 | HDAC4,BCL11A,SORBS1 |
| V$AR_Q2 | 125 | 3.43 | 3 | 0.672 | 0.892 | RERE,TSHZ3,WNT3 |
| TCTATGA,MIR-376A,MIR-376B | 84 | 2.31 | 2 | 0.676 | 0.894 | MACROD2,BRWD1 |
| V$NERF_Q2 | 247 | 6.79 | 6 | 0.677 | 0.894 | FURIN,GBF1,NTRK3,DPP8,SORBS |

| | | | | | | |
|---|---|---|---|---|---|---|
| AAAGGAT,MIR-501 | 126 | 3.46 | 3 | 0.677 | 0.894 | MEF2C,SRR,NFIX |
| CAGNWMCNNNGAC_UNKNOWN | 85 | 2.34 | 2 | 0.682 | 0.898 | MGAT5B,JMJD1C |
| CAGGGTC,MIR-504 | 85 | 2.34 | 2 | 0.682 | 0.898 | NTM,NFIX |
| V$IRF1_01 | 250 | 6.87 | 6 | 0.688 | 0.904 | HDAC4,LDB1,CACNB2,BCL11A,NF |
| V$STAT5A_01 | 251 | 6.9 | 6 | 0.692 | 0.908 | SEMA6D,ATP2A2,NT5C2,ANK3,WN |
| V$STAT1_02 | 252 | 6.92 | 6 | 0.695 | 0.911 | NRGN,ADAM22,ZNF568,NTRK3,SY |
| V$HMX1_01 | 43 | 1.18 | 1 | 0.699 | 0.913 | CACNB2 |
| ACAACCT,MIR-453 | 43 | 1.18 | 1 | 0.699 | 0.913 | BCL11B |
| TNCATNTCCYR_UNKNOWN | 131 | 3.6 | 3 | 0.702 | 0.915 | ADAM22,ZNF568,WNT3 |
| V$WHN_B | 254 | 6.98 | 6 | 0.702 | 0.915 | GRIN2A,ESRRG,HDAC4,SND1,ANI |
| V$CREL_01 | 256 | 7.03 | 6 | 0.71 | 0.921 | WRN,RASGRP4,SOX5,KCNN2,PPF |
| V$PXR_Q2 | 256 | 7.03 | 6 | 0.71 | 0.921 | ARHGAP15,ITIH3,CREB5,NTM,JMJ |
| TTTNNANAGCYR_UNKNOWN | 133 | 3.65 | 3 | 0.712 | 0.923 | FOXP1,TCF4,PPP2R2B |
| CCTGAGT,MIR-510 | 45 | 1.24 | 1 | 0.715 | 0.925 | NOVA1 |
| WGGAATGY_V$TEF1_Q6 | 378 | 10.4 | 9 | 0.716 | 0.925 | GALNT12,KMT2E,PTCH1,MDK,PKF |
| GGGNNTTTCC_V$NFKB_Q6_01 | 134 | 3.68 | 3 | 0.717 | 0.925 | NRGN,SOX5,NTRK3 |
| CTCNANGTGNY_UNKNOWN | 91 | 2.5 | 2 | 0.718 | 0.925 | CKB,LRFN5 |
| V$PPAR_DR1_Q2 | 259 | 7.12 | 6 | 0.72 | 0.926 | HDAC4,ARFGEF2,LRFN5,GBF1,CS |
| TCTAGAG,MIR-517 | 46 | 1.26 | 1 | 0.723 | 0.928 | CUL3 |
| V$DR4_Q2 | 260 | 7.14 | 6 | 0.723 | 0.928 | NREP,FOXP1,HDAC4,SOX5,NOVA |
| V$CREB_Q2_01 | 220 | 6.04 | 5 | 0.727 | 0.929 | PFAS,XRCC6,CALD1,CACNB2,CLC |
| V$HNF4_DR1_Q3 | 261 | 7.17 | 6 | 0.727 | 0.929 | TCAIM,SOX5,JMJD1C,GBF1,WNT3 |
| V$TTF1_Q6 | 262 | 7.2 | 6 | 0.73 | 0.932 | RSRC1,TCF4,ZNF804A,CITED1,NC |

| | | | | | | |
|---|---|---|---|---|---|---|
| V$COUP_01 | 263 | 7.23 | 6 | 0.733 | 0.932 | ESRRG,ARFGEF2,JMJD1C,GBF1,( |
| V$NFKB_C | 263 | 7.23 | 6 | 0.733 | 0.932 | WRN,SOX5,BCL11A,PPP1R13B,RH |
| V$BACH1_01 | 263 | 7.23 | 6 | 0.733 | 0.932 | GNAI1,GIGYF2,TNRC6A,DYSF,SRF |
| V$AHRARNT_01 | 140 | 3.85 | 3 | 0.744 | 0.943 | RBM6,BCL11A,RGS6 |
| GGATTA_V$PITX2_Q2 | 587 | 16.1 | 14 | 0.745 | 0.943 | SEMA3A,ENOX1,SPAG16,TCF4,US |
| V$AP1FJ_Q2 | 268 | 7.36 | 6 | 0.75 | 0.943 | GNAI1,SLC6A9,GABBR1,JMJD1C,F |
| V$EFC_Q6 | 268 | 7.36 | 6 | 0.75 | 0.943 | NRGN,NPAS3,NCOA5,PTCH1,GBF |
| V$NF1_Q6_01 | 268 | 7.36 | 6 | 0.75 | 0.943 | KMT2E,CACNA1C,RHOJ,TANK,FAI |
| V$NRSF_01 | 97 | 2.67 | 2 | 0.75 | 0.943 | MGAT5B,ATP2B2 |
| CCAGGGG,MIR-331 | 97 | 2.67 | 2 | 0.75 | 0.943 | GABBR1,FURIN |
| V$SRF_01 | 50 | 1.37 | 1 | 0.752 | 0.943 | CALD1 |
| WWTAAGGC_UNKNOWN | 142 | 3.9 | 3 | 0.753 | 0.943 | KDM4A,SORBS1,HDAC9 |
| YGTCCTTGR_UNKNOWN | 98 | 2.69 | 2 | 0.755 | 0.945 | SEMA6D,ATP2A2 |
| V$AREB6_01 | 271 | 7.45 | 6 | 0.759 | 0.946 | RTN1,ENOX1,CREB5,CADM2,AFF3 |
| V$HNF4ALPHA_Q6 | 271 | 7.45 | 6 | 0.759 | 0.946 | ZNF804A,ARFGEF2,CADM2,GBF1, |
| ATCMNTCCGY_UNKNOWN | 51 | 1.4 | 1 | 0.759 | 0.946 | MAML3 |
| V$NFE2_01 | 272 | 7.47 | 6 | 0.762 | 0.948 | ESRRG,FHIT,TNRC6A,TTLL7,TNXE |
| ACTACCT,MIR-196A,MIR-196B | 145 | 3.98 | 3 | 0.765 | 0.95 | ABCB9,PPP1R16B,LCORL |
| V$PR_01 | 148 | 4.07 | 3 | 0.777 | 0.962 | JMJD1C,ELMO1,PCDH9 |
| TCCAGAG,MIR-518C | 148 | 4.07 | 3 | 0.777 | 0.962 | ATP2A2,NFIX,PCDH9 |
| ACAWYAAAG_UNKNOWN | 103 | 2.83 | 2 | 0.779 | 0.962 | DOCK4,CSRNP3 |
| V$IK1_01 | 278 | 7.64 | 6 | 0.78 | 0.962 | FOXP1,WRN,PTCH1,ZFPM2,MMP1 |
| CCCNNGGGAR_V$OLF1_01 | 320 | 8.79 | 7 | 0.781 | 0.962 | ANXA9,KMT2A,LRRC20,GRIK1,CA( |
| GATTGGY_V$NFY_Q6_01 | 1160 | 31.9 | 28 | 0.792 | 0.975 | NRGN,SLC8A3,KMT2E,HLA-DMA,F |

| | | | | | | |
|---|---|---|---|---|---|---|
| GGAANCGGAANY_UNKNOWN | 106 | 2.91 | 2 | 0.793 | 0.975 | PRPF3,MED8 |
| V$AP2ALPHA_01 | 241 | 6.62 | 5 | 0.796 | 0.977 | PPM1E,CITED1,LDB1,SOX5,KDM3[illegible] |
| CTCTAGA,MIR-526C,MIR-518F,MIR-526A | 58 | 1.59 | 1 | 0.802 | 0.983 | NREP |
| CCAATNNSNNNGCG_UNKNOWN | 59 | 1.62 | 1 | 0.807 | 0.988 | MEF2C |
| V$AR_01 | 157 | 4.31 | 3 | 0.81 | 0.99 | TNRC6A,CBLB,ZNF800 |
| V$STAT5B_01 | 247 | 6.79 | 5 | 0.813 | 0.991 | BEND4,NT5C2,ANK3,DOCK4,TMO[illegible] |
| MYAATNNNNNNNGGC_UNKNOWN | 111 | 3.05 | 2 | 0.813 | 0.991 | SOX5,DOCK4 |
| V$ELK1_02 | 248 | 6.81 | 5 | 0.816 | 0.991 | CALU,PRKAG2,PALB2,DPP8,MED[illegible] |
| V$ATF3_Q6 | 248 | 6.81 | 5 | 0.816 | 0.991 | KMT2E,PKP4,PFAS,CREB5,KCNN[illegible] |
| TATAAA_V$TATA_01 | 1300 | 35.6 | 31 | 0.818 | 0.992 | RPA3,FOXP1,SEMA3A,ESR2,ESR[illegible] |
| V$AP2GAMMA_01 | 250 | 6.87 | 5 | 0.821 | 0.995 | PPM1E,LDB1,GMIP,SOX5,KDM3B |
| TATCTGG,MIR-488 | 62 | 1.7 | 1 | 0.823 | 0.996 | SYNGAP1 |
| V$RREB1_01 | 207 | 5.69 | 4 | 0.825 | 0.996 | RERE,WRN,AUTS2,NTRK3 |
| RNTCANNRNNYNATTW_UNKNOWN | 63 | 1.73 | 1 | 0.828 | 0.997 | SRPK2 |
| TTGCWCAAY_V$CEBPB_02 | 63 | 1.73 | 1 | 0.828 | 0.997 | MSH5 |
| WCAANNNYCAG_UNKNOWN | 254 | 6.98 | 5 | 0.831 | 1 | KMT2D,SLC6A9,FGFR1,PPP2R2B,[illegible] |
| V$FXR_Q3 | 116 | 3.19 | 2 | 0.832 | 1 | ITIH3,NTRK3 |
| V$MYCMAX_01 | 255 | 7.01 | 5 | 0.834 | 1 | KMT2A,TET2,BEND4,FOXO3,CAM[illegible] |
| V$ETF_Q6 | 117 | 3.21 | 2 | 0.836 | 1 | RERE,TRIM33 |
| GATGKMRGCG_UNKNOWN | 65 | 1.79 | 1 | 0.837 | 1 | WNT3 |

| | | | | | | |
|---|---|---|---|---|---|---|
| V$CEBPGAMMA_Q6 | 257 | 7.06 | 5 | 0.839 | 1 | ANKS1B,CREB5,MEF2C,JMJD1C,N |
| V$DR1_Q3 | 257 | 7.06 | 5 | 0.839 | 1 | ARFGEF2,SOX5,GBF1,CSRNP3,MI |
| V$ATF4_Q2 | 258 | 7.09 | 5 | 0.841 | 1 | BSN,KCNN2,TNXB,PCDH9,SORBS |
| AAGWWRNYGGC_UNKNOWN | 119 | 3.27 | 2 | 0.843 | 1 | PPM1E,MED27 |
| V$PAX_Q6 | 259 | 7.12 | 5 | 0.844 | 1 | SEMA3A,LDB1,NTM,NOVA1,DOCK |
| TGANTCA_V$AP1_C | 1120 | 30.8 | 26 | 0.844 | 1 | GNAI1,RERE,RTN1,GIGYF2,PSMA |
| CCAGGTT,MIR-490 | 67 | 1.84 | 1 | 0.846 | 1 | ETF1 |
| V$ARNT_01 | 261 | 7.17 | 5 | 0.848 | 1 | KMT2A,SPPL3,BEND4,ZCCHC7,BC |
| V$LEF1_Q6 | 261 | 7.17 | 5 | 0.848 | 1 | SOX5,TNXB,WNT3,SORBS1,AFF3 |
| AGCYRWTTC_UNKNOWN | 122 | 3.35 | 2 | 0.853 | 1 | KMT2A,KDM3B |
| V$CREB_Q2 | 263 | 7.23 | 5 | 0.853 | 1 | PKP4,PFAS,ETF1,CACNB2,CLCN3 |
| V$AML1_01 | 263 | 7.23 | 5 | 0.853 | 1 | SLC6A9,ATP2A2,SOX5,ANK3,JMJD |
| V$AML1_Q6 | 263 | 7.23 | 5 | 0.853 | 1 | SLC6A9,ATP2A2,SOX5,ANK3,JMJD |
| GCGSCMNTTT_UNKNOWN | 69 | 1.9 | 1 | 0.854 | 1 | CNOT1 |
| V$ATF6_01 | 123 | 3.38 | 2 | 0.856 | 1 | STAG1,GBF1 |
| V$E2F_Q2 | 176 | 4.84 | 3 | 0.866 | 1 | NEGR1,TRIM33,MED8 |
| CTGRYYYNATT_UNKNOWN | 72 | 1.98 | 1 | 0.866 | 1 | CACNB2 |
| V$AP1_Q4 | 271 | 7.45 | 5 | 0.87 | 1 | RPA3,GNAI1,ESRRB,SRPK2,PCDH |
| V$USF_02 | 272 | 7.47 | 5 | 0.872 | 1 | SEMA3A,GIGYF2,SPPL3,SPNS1,CI |
| V$AP1_Q2_01 | 275 | 7.56 | 5 | 0.878 | 1 | TCF4,DYSF,ANK3,TNXB,PCDH9 |
| GGGYGTGNY_UNKNOWN | 664 | 18.2 | 14 | 0.879 | 1 | KMT2E,RTN1,GIGYF2,SPPL3,MDK |
| CCGNMNNTNACG_UNKNOWN | 77 | 2.12 | 1 | 0.884 | 1 | KCNN2 |
| TGCGCANK_UNKNOWN | 545 | 15 | 11 | 0.888 | 1 | KMT2A,MSL2,KMT2E,RSRC1,NCO |

| | | | | | | |
|---|---|---|---|---|---|---|
| CTCAGGG,MIR-125B,MIR-125A | 329 | 9.04 | 6 | 0.893 | 1 | MYO18A,GRIN2A,BSN,PTPRF,ZFY |
| V$TAXCREB_01 | 137 | 3.76 | 2 | 0.894 | 1 | MAGI1,KDM3B |
| YYCATTCAWW_UNKNOWN | 191 | 5.25 | 3 | 0.9 | 1 | CKB,ANK3,PDE4B |
| CCAWNWWNNNGGC_UNKNOWN | 84 | 2.31 | 1 | 0.904 | 1 | STAG1 |
| V$E2A_Q2 | 243 | 6.68 | 4 | 0.905 | 1 | ATP2A2,FBXL17,PPP2R2B,ZNF80( |
| V$COUP_DR1_Q6 | 247 | 6.79 | 4 | 0.911 | 1 | ARFGEF2,GBF1,CSRNP3,MED8 |
| RRCCGTTA_UNKNOWN | 87 | 2.39 | 1 | 0.912 | 1 | FXR1 |
| V$ARNT_02 | 248 | 6.81 | 4 | 0.913 | 1 | GIGYF2,SPPL3,SPNS1,PTPRF |
| V$HSF_Q6 | 201 | 5.52 | 3 | 0.917 | 1 | RTN1,ANK3,TNXB |
| V$TFIIA_Q6 | 251 | 6.9 | 4 | 0.918 | 1 | ANKS1B,ATP2A2,JMJD1C,SRPK2 |
| V$ER_Q6_02 | 252 | 6.92 | 4 | 0.919 | 1 | SLC8A3,KMT2E,PPM1E,DOCK4 |
| GTGACGY_V$E4F1_Q6 | 658 | 18.1 | 13 | 0.919 | 1 | MTHFD1L,RERE,GIGYF2,PKP4,SG |
| V$NFKB_Q6 | 254 | 6.98 | 4 | 0.922 | 1 | WRN,SOX5,TANK,PPP1R13B |
| V$TATA_01 | 255 | 7.01 | 4 | 0.923 | 1 | ESRRG,CKB,SOX5,RHOA |
| V$PEA3_Q6 | 255 | 7.01 | 4 | 0.923 | 1 | NRGN,NT5C2,ANK3,TRIM33 |
| V$MYOGENIN_Q6 | 255 | 7.01 | 4 | 0.923 | 1 | NEDD4L,SLC35F2,PARD3B,NEGR[illegible] |
| V$NRF2_Q4 | 255 | 7.01 | 4 | 0.923 | 1 | ESRRG,CACNB2,PPP2R2B,TNXB |
| V$USF_01 | 256 | 7.03 | 4 | 0.925 | 1 | GIGYF2,ESR2,UTY,ZCCHC7 |
| RCGCANGCGY_V$NRF1_Q6 | 918 | 25.2 | 19 | 0.925 | 1 | NRGN,RSU1,MAPT,SGF29,NT5DC[illegible] |
| V$CREBP1CJUN_01 | 259 | 7.12 | 4 | 0.929 | 1 | PKP4,PFAS,STAG1,CLCN3 |
| V$USF_Q6 | 259 | 7.12 | 4 | 0.929 | 1 | GIGYF2,TET2,SPNS1,XRCC6 |
| V$GABP_B | 259 | 7.12 | 4 | 0.929 | 1 | PRKAG2,GALNT10,DPP8,RHOA |
| V$AP1_Q4_01 | 261 | 7.17 | 4 | 0.931 | 1 | GNAI1,ESRRG,SRPK2,TNXB |

| | | | | | | |
|---|---|---|---|---|---|---|
| ACACTGG,MIR-199A,MIR-199B | 157 | 4.31 | 2 | 0.933 | 1 | DDX3Y,ATP2B2 |
| V$AP1_Q2 | 265 | 7.28 | 4 | 0.936 | 1 | GNAI1,JMJD1C,SRPK2,PCDH9 |
| TMTCGCGANR_UNKNOWN | 160 | 4.4 | 2 | 0.937 | 1 | SPAG16,ANAPC4 |
| V$OCT_C | 268 | 7.36 | 4 | 0.94 | 1 | CADM2,GALNT1,CSRNP3,HDAC9 |
| CCCNNNNNNAAGWT_UNKNOWN | 102 | 2.8 | 1 | 0.942 | 1 | CNOT1 |
| CACGTG_V$MYC_Q2 | 1030 | 28.4 | 21 | 0.945 | 1 | KMT2A,GRIN2A,GIGYF2,SPPL3,ES |
| SCGGAAGY_V$ELK1_02 | 1200 | 32.9 | 25 | 0.945 | 1 | MRPL33,PSMA5,RSU1,ANAPC4,CT |
| V$SREBP1_01 | 168 | 4.62 | 2 | 0.948 | 1 | GIGYF2,SPPL3 |
| V$E2F_02 | 235 | 6.46 | 3 | 0.959 | 1 | MAPT,MSH5,STAG1 |
| V$E2F1DP1_01 | 235 | 6.46 | 3 | 0.959 | 1 | MAPT,MSH5,STAG1 |
| V$E2F1DP2_01 | 235 | 6.46 | 3 | 0.959 | 1 | MAPT,MSH5,STAG1 |
| V$E2F4DP2_01 | 235 | 6.46 | 3 | 0.959 | 1 | MAPT,MSH5,STAG1 |
| V$ZF5_01 | 238 | 6.54 | 3 | 0.961 | 1 | LRFN5,SORCS3,NEURL1 |
| V$E2F4DP1_01 | 239 | 6.57 | 3 | 0.962 | 1 | MAPT,MSH5,STAG1 |
| GGAMTNNNNNTCCY_UNKNOWN | 117 | 3.21 | 1 | 0.962 | 1 | FHIT |
| V$MYC_Q2 | 185 | 5.08 | 2 | 0.965 | 1 | KMT2A,PTPRF |
| V$SP1_Q6_01 | 243 | 6.68 | 3 | 0.965 | 1 | TCF4,ATP2A2,LDB1 |
| V$E2F1_Q4 | 244 | 6.7 | 3 | 0.966 | 1 | SGF29,GRIK1,STAG1 |
| ARGGGTTAA_UNKNOWN | 121 | 3.32 | 1 | 0.966 | 1 | RGS6 |
| V$ATF_B | 187 | 5.14 | 2 | 0.966 | 1 | PKP4,PFAS |
| V$E2F_03 | 245 | 6.73 | 3 | 0.967 | 1 | NRGN,AUTS2,STAG1 |
| V$SP3_Q3 | 245 | 6.73 | 3 | 0.967 | 1 | FOXP1,GRIN2A,CADM2 |
| V$USF2_Q6 | 251 | 6.9 | 3 | 0.971 | 1 | KMT2A,GUCY1A2,PTPRF |
| V$IRF7_01 | 252 | 6.92 | 3 | 0.971 | 1 | CALD1,SOX5,NFIX |

| | | | | | | |
|---|---|---|---|---|---|---|
| V$AP1_01 | 267 | 7.34 | 3 | 0.979 | 1 | TNRC6A,NEDD4L,TNXB |
| V$CREB_Q4 | 268 | 7.36 | 3 | 0.98 | 1 | PKP4,PFAS,CLCN3 |
| V$NRF2_01 | 270 | 7.42 | 3 | 0.98 | 1 | PALB2,DPP8,MED8 |
| TGCCAAR_V$NF1_Q6 | 722 | 19.8 | 12 | 0.981 | 1 | MAPT,ESRRG,TCAIM,PTCH1,TCF4 |
| V$BACH2_01 | 271 | 7.45 | 3 | 0.981 | 1 | TNRC6A,DYSF,TNXB |
| V$CREB_Q4_01 | 211 | 5.8 | 2 | 0.981 | 1 | PKP4,PFAS |
| GGGCGGR_V$SP1_Q6 | 2940 | 80.8 | 65 | 0.982 | 1 | NRGN,FKBP9,MTHFD1L,MAGI1,MY |
| V$AP1_C | 275 | 7.56 | 3 | 0.982 | 1 | ESRRG,ANK3,PCDH9 |
| V$USF_C | 279 | 7.67 | 3 | 0.984 | 1 | GRIN2A,BEND4,CAMKV |
| V$E2F1_Q4_01 | 228 | 6.26 | 2 | 0.988 | 1 | STAG1,NEGR1 |
| TGACGTCA_V$ATF3_Q6 | 231 | 6.35 | 2 | 0.988 | 1 | PKP4,PFAS |
| V$E2F1_Q6 | 232 | 6.37 | 2 | 0.989 | 1 | MSH5,STAG1 |
| V$E2F_Q4 | 234 | 6.43 | 2 | 0.989 | 1 | STAG1,WNT3 |
| V$E2F_Q3_01 | 235 | 6.46 | 2 | 0.989 | 1 | STAG1,NEGR1 |
| V$E2F_Q4_01 | 237 | 6.51 | 2 | 0.99 | 1 | STAG1,NEGR1 |
| V$E2F1_Q6_01 | 238 | 6.54 | 2 | 0.99 | 1 | CADM2,STAG1 |
| V$USF_Q6_01 | 239 | 6.57 | 2 | 0.99 | 1 | TET2,SPNS1 |
| V$E2F_Q6_01 | 240 | 6.59 | 2 | 0.991 | 1 | STAG1,SORBS1 |
| SGCGSSAAA_V$E2F1DP2_01 | 168 | 4.62 | 1 | 0.991 | 1 | MSH5 |
| V$E2F1_Q3 | 244 | 6.7 | 2 | 0.992 | 1 | MAML3,FOXO3 |
| TGAYRTCA_V$ATF3_Q6 | 538 | 14.8 | 7 | 0.993 | 1 | PKP4,PFAS,RHOJ,CTNND1,MAML3 |
| V$CREB_02 | 254 | 6.98 | 2 | 0.993 | 1 | MAGI1,MED27 |
| V$AP1_Q6 | 259 | 7.12 | 2 | 0.994 | 1 | GNAI1,PCDH9 |
| V$CREB_01 | 262 | 7.2 | 2 | 0.995 | 1 | PKP4,PFAS |
| TGASTMAGC_V$NFE2_01 | 195 | 5.36 | 1 | 0.996 | 1 | TNXB |

| | | | | | | |
|---|---|---|---|---|---|---|
| V$E2F_Q3 | 227 | 6.24 | 1 | 0.998 | 1 | FOXO3 |
| V$E2F1DP1RB_01 | 231 | 6.35 | 1 | 0.998 | 1 | MAPT |
| V$E2F_Q6 | 232 | 6.37 | 1 | 0.999 | 1 | STAG1 |
| V$NFKAPPAB_01 | 251 | 6.9 | 1 | 0.999 | 1 | SOX5 |
| V$CREBP1_Q2 | 254 | 6.98 | 1 | 0.999 | 1 | BCL11A |

ining motif NNNGGGAGTNNNNS which matches annotation for PCAF: p300/CBP-associated factor - transcri

ne occurence of the motif TAGCTTT in their 3' untranslated region. The motif represents putative target (that is ıe occurrence of the highly conserved motif M17 AACTTT in the regions spanning 4 kb centered on their trans ıining motif NNNRTAATNANNN which matches annotation for POU2F1: POU domain, class 2, transcription fa[illegible]

ed in inflammatory diseases, miRNA-135a regulates sterss reactions - including regulation of HT1A-5.

tant role in diverse biological processes, including the pluripotency of human embryonic stem cells (hESCs), se

ng site of GATA1 - TF expresses in hematopoetic cells and plays an important role in their development and fu egulatory role in brain function, with several reports displaying an association with proliferation and differentiation

ining motif GCANCTGNY which matches annotation for MYOD1: myogenic factor 3 - regulates muscle cell diff ge paired-class homeoprotein 1 - Most probably regulates the expression of genes involved in the developmen

:Z1,BEND4,SLC6A9,ZFPM2,NEDD4L,ZMIZ1,SLC13A1,CALD1,SYT1,DPYD,CSRNP3,TMOD3,HDAC9

)4 - regulates many cellular pathways, including oxidative stress signaling, longevity, insulin signaling, cell cycle ı chemoresistance, epithelial-mesenchymal transition (EMT), allergic response, inflammation, and AD

ℵP1,GRIN2A,MAPT,ESRRG,PTCH1,DMTF1,CITED1,FGFR1,CADM2,CTNND1,STAG1,CACNB2,ANK3,LRFN sticity, synaptic function, regulation of genes of development and function of adipocytes, regulation of inflamma

cipates in autophagy, apoptosis, oxidative stress, and inflammation through post-transcriptional regulation of tar

,FOXP1,ESRRG,DYNC1I1,CTR9,CUL3,DOCK8,HDAC4,PPM1E,ZFPM2,ANKS1B,FXR1,CARMIL1,CREB5,GI

KS1B,FGFR1,ETF1,MEF2C,PRKG1,FBXL17,RNF111,SRPK2,GATAD2B,MACROD2,ATP2B2,BRWD1,BCL11

3,RPA3,RERE,FOXP1,SEMA3A,ENOX1,GIGYF2,ESRRG,PTCH1,PLCL1,PKP4,TCF4,ITIH3,CUL3,CNTNAP2

N2,PPM1E,TCF20,SEMA6D,ARID1B,SYT1,ABCB9,FURIN,BCL11A,RGS6,NTRK3,ELMO1,ZKSCAN8,RNF14

N1,ENOX1,ESRRG,PTCH1,USP4,PPM1E,MSH5,ETF1,CREB5,MGAT5B,RHOJ,SOX5,CD55,NTM,FURIN,P

L2,RPA3,RERE,FOXP1,GRIN2A,GIGYF2,NCOA5,TET2,PTCH1,PKP4,DYNC1I1,USP4,NT5DC2,PPM1E,GAB

CF20,ZMIZ1,GUCY1A2,DBN1,MAGI2,CADM2,SOX5,PRKG1,CACNA1I,LCORL,ZMIZ2,NFIX,ELMO1,KIF21B,

PPM1E,MMP16,ATP2A2,PRKAG2,PALB2,SRR,LRFN5,SORCS3,IMMP2L,TRIM33,NEURL1,RHOA

E,GNAI1,FOXP1,SEMA3A,RTN1,GRIN2A,ESRRG,KIZ,PTCH1,SPAG16,BEND4,NT5DC2,ZFPM2,NEDD4L,ITG

D4,ANKS1B,ATP2A2,SLC13A1,CARMIL1,CREB5,CALD1,SYT1,MED27,PPP2R2B,NFIX,ELMO1,NOVA1,DOC

C4,PPM1E,GABBR1,FXR1,MMP16,ATP2A2,MEF2C,CADM2,PCGF3,ANK3,BCL11A,FOXO3,ELMO1

,SLC6A9,EXT1,CACNA1C,LDB1,CTNND1,SOX5,ST3GAL3,JMJD1C,GID4,NOVA1,BRWD1,HDAC9

F4,ANKS1B,CALU,ARID1B,CNOT1,CALD1,CADM2,CTNND1,ESRRB,TANK,MAML3,JMJD1C,GBF1,PPP2R2

RG,TET2,TCF4,EXT1,LDB1,FGFR1,CREB5,CALD1,PTPRF,LRFN5,MAML3,JMJD1C,PPP2R2B,NOVA1,WN

)X1,ESR2,ESRRG,WRN,PTCH1,FHIT,HDAC4,NEDD4L,CREB5,CADM2,SOX5,CACNB2,PTPRF,MED27,LRI

\3A,RTN1,WRN,PTCH1,TCF4,CNTNAP2,ATP2A2,CITED1,LDB1,ARID1B,CREB5,MEF2C,GRIK1,CALD1,CA

PA3,FOXP1,GIGYF2,RSRC1,SPPL3,ESRRG,MDK,TCF4,BEND4,CLSTN2,HDAC4,PPM1E,DYSF,CNTN4,NE

\,CUL3,SEMA6D,AUTS2,MEF2C,BCL11A,TSHZ3,KDM4A,PDE4B,PPP1R13B,CLCN3,BRWD1,BCL11B,AMBI

,CREB5,DDX3Y,CALD1,SOX5,SYT1,GBF1,NKIRAS1,KCNN2,NFIX,GATAD2B,MACROD2,PREX1,BRWD1,C

;LC6A9,HDAC4,ZFPM2,NEDD4L,DMTF1,CREB5,CALD1,CADM2,FURIN,TSHZ3,PTPRO,GBF1,NKIRAS1,RB

N2,TNRC6A,PPM1E,CNTN4,OLFM4,CBLB,ANKS1B,ATP2A2,ZMIZ1,SLC13A1,CARMIL1,CREB5,MEF2C,CAL

RG,TET2,KIZ,PTCH1,MDK,TCF4,DYNC1I1,SEMA6D,MMP16,EXT1,CREB5,CALD1,SOX5,NGEF,CACNB2,C

END4,DYNC1I1,CLSTN2,TNRC6A,DYSF,COL11A1,SEMA6D,ANKS1B,FXR1,DMTF1,ZNF638,CREB5,GRIK1

PM1E,GABBR1,ZFPM2,SND1,LDB1,CARMIL1,CTNND1,ANK3,PPP1R16B,ESRRB,JMJD1C,PPP2R2B,HLA-D

AC4,ZFPM2,BSN,MPHOSPH9,CACNA1C,DDX3Y,DBN1,MAGI2,SOX5,SYT1,TSHZ3,RNF111,WNT3,ATP2B

CNA2D2,COL11A1,CALU,ZMIZ1,MEF2C,DBN1,PCGF3,PPP1R16B,JMJD1C,RNF111,NFIX,GATAD2B,PCDH

GAP15,ADAM22,RERE,HLA-DMA,SPPL3,MAPT,ESRRG,WRN,SSUH2,MDK,PKP4,TCF4,BEND4,TNRC6A,SI

IA6D,STAG1,SYT1,CACNB2,ABCB9,JMJD1C,RNF111,RGS6,ELMO1,NOVA1,THRB,CLCN3,RNF144A,ZNF8

E,FOXP1,SEMA3A,RTN1,PSMA5,RSRC1,NCOA5,SPPL3,MAPT,ESRRG,UTY,TET2,MDK,F2,CTR9,RBM6,TN

YF2,ESRRG,KIZ,PTCH1,SLC22A23,TCF4,CACNA2D2,CLSTN2,SLC6A9,COL11A1,NEDD4L,SEMA6D,ANKS

11A1,CALU,CREB5,RASGRP4,CTNND1,PCGF3,ANK3,ABCB9,BCL11A,GID4,PPP1R13B,NFIX,SYNE1,BRW

ETF1,CREB5,RHOJ,CTNND1,SOX5,MIRLET7BHG,GID4,ZFYVE1,ELMO1,DPP8,WNT3,PCDH9,TMOD3

RERE,SEMA3A,ESRRG,PLCL1,TCF4,PPM1E,CNTN4,SND1,CITED1,ETF1,CREB5,MEF2C,SOX5,CTNNB1,

I5,FOXP1,KIZ,BEND4,HFE,CNTN4,CBLB,SND1,DMTF1,ARFGEF2,NT5C2,CNOT1,PRKAG2,ETF1,MEF2C,F [illegible],HDAC4,SEMA6D,ANKS1B,CALU,DDX3Y,ZKSCAN3,CTNND1,FURIN,CPEB1,BCL11A,PDE4B,NFIX,NEGR1

NC1I1,RBM6,PPM1E,GABBR1,SEMA6D,DMTF1,MSH5,DDX3Y,FURIN,RNF111,SYNE1,PCDH9,ATP2B2,PPF

DK,PLCL1,PKP4,TCF4,BEND4,OLFM4,CARMIL1,MEF2C,CADM2,SOX5,MAML3,TSHZ3,SRPK2,SORBS1,B[illegible]

9,MYO18A,DGKI,RPA3,SEMA3A,RTN1,GRIN2A,GIGYF2,RSRC1,NCOA5,SPPL3,MAPT,ESRRG,KIZ,PRKD1,

PT,WRN,PTCH1,TCF4,ITIH3,ZFPM2,NEDD4L,ANKS1B,CITED1,LDB1,EIF1AY,FGFR1,AUTS2,GMIP,GRIK1,

,DOCK8,CNTN4,ITGA11,DMTF1,CNOT1,PRKAG2,ETF1,PALB2,FURIN,GALNT10,ELMO1,DPP8,PREX1,CS

SEMA6D,ITGA11,GLT8D1,ETF1,CACNB2,CADPS,NFIX,NEGR1,THRB,MACROD2,BRWD1,RNF144A

[illegible]1B,MEF2C,CALD1,CTNND1,ADTRP,LRFN5,BCL11A,TSHZ3,KCNN2,RGS6,ELMO1,WNT3,PPP2R5C,SORB[illegible]

:,ESRRG,F2,SLC6A9,PPM1E,CNTN4,CALU,CKB,SDK1,MEF2C,SOX5,CACNB2,ANK3,PPP1R16B,ESRRB,D

PTCH1,NT5DC2,CNTN4,CALU,LRRC20,CNOT1,ETF1,CREB5,CADM2,STAG1,ANK3,CNNM2,TANK,IVD,KDM [illegible]RG,UTY,TCF4,TNRC6A,CUL3,CNTN4,SEMA6D,ANKS1B,CITED1,CKB,CREB5,CALD1,SOX5,ADTRP,ANK3,[illegible]

5,ESRRG,GPR141,TNRC6A,CNTNAP2,DYSF,ATP2A2,TTLL7,CREB5,MEF2C,TFB1M,ANK3,FURIN,DPYD,P

2,BSN,SND1,ZNF638,PRPF3,CNOT1,PALB2,STAG1,SRR,ZCCHC7,RNF111,SORCS3,MACROD2,RHOA,TM

R2,UTY,TET2,PTCH1,TCF4,BEND4,SPNS1,FXR1,GUCY1A2,ZCCHC7,FOXO3,SEMA3F,PPP2R2B,TNXB,B

R9,TNRC6A,SND1,CALU,ATP2A2,ZSCAN12,PRKAG2,GMIP,PALB2,STAG1,ZCCHC7,RFC2,SEMA3F,GAL

YO18A,KMT2E,RTN1,GRIN2A,PSMA5,RSU1,NCOA5,SPPL3,MAPT,UTY,TET2,PTCH1,TCF4,DYNC1I1,CLST

, seed match) of human mature miRNA hsa-miR-9* (v7.1 miRBase). miR-9 is highly expressed in human brain cription starting sites [-2kb,+2kb]. The motif does not match any known transcription factor binding site.

ctor 1 - Transcription factor that binds to the octamer motif (5'-ATTTGCAT-3') and activates the promoters of th

ferentiation by inducing cell cycle arrest, a prerequisite for myogenic initiation. The protein is also involved in mu t of mesenchyme-derived craniofacial structures. Early on in development, it plays a role in forebrain mesenchy

I5,BCL11A,TSHZ3,FOXO3,JMJD1C,PIP5K1B,KCNN2,PPP2R2B,NFIX,TNXB,ELMO1,NOVA1,PCDH9,PPP2R

RIK1,TFB1M,CTNND1,PPP1R16B,ESRRB,MAML3,BCL11A,JMJD1C,NFIX,SYNE1,PCDH9,TRIM33,CSRNP3,

,HDAC4,BSN,NEDD4L,SND1,SEMA6D,EXT1,CALU,ATP2A2,ZMIZ1,CACNA1C,TTLL7,ARID1B,NT5C2,MSH[illegible]

[illegible]BR1,CBLB,EXT1,CALU,DMTF1,CKB,CALD1,CADM2,KCNB1,PPP1R16B,MAML3,PPP2R2B,KDM3B,CSRNP[illegible]

GA11,ZMIZ1,GATB,PRPF3,FGFR1,CREB5,DBN1,SOX5,ESRRB,DPYD,TSHZ3,PTPRO,JMJD1C,PPP2R2B,N

FN5,MAML3,BCL11A,PDE4B,SEMA3F,PPP2R2B,NFIX,SRPK2,CLCN3,RPS6KL1,FAM193A,BCL11B,CSRNP

DM2,SOX5,ADTRP,BCL11A,TSHZ3,SORCS3,PPP2R2B,SRPK2,MACROD2,BCL11B,TMOD3,AFF3

DD4L,SEMA6D,ZNF398,FXR1,ZMIZ1,DMTF1,TTLL7,ETF1,MEF2C,CADM2,SOX5,CACNB2,ABCB9,CNNM2, [illegible]D1,CADM2,CTNND1,SOX5,LRFN5,PTPRO,PPP2R2B,PPP1R13B,NFIX,SRPK2,PPP1R16A,ELMO1,NOVA1[illegible]

ADPS,MAML3,BCL11A,FOXO3,ZFYVE1,PPP2R2B,NFIX,SRPK2,NTRK3,ELMO1,IMMP2L,PCDH9,BCL11B,C

,RHOJ,CALD1,SOX5,ESRRB,TSHZ3,FOXO3,GBF1,PPP2R2B,RGS6,SRPK2,ELMO1,WNT3,PPP2R5C,DOC

_C6A9,CNTNAP2,HDAC4,PPM1E,GABBR1,ZFPM2,CNTN4,COL11A1,CBLB,ITGA11,ATP2A2,CACNA1C,LD[illegible]

ıRC6A,ZFPM2,CNTN4,COL11A1,OLFM4,NEDD4L,SEMA6D,MMP16,ZMIZ1,CKB,ZSCAN12,LDB1,LRRC20,C

1B,XRCC6,CREB5,MEF2C,CTNND1,SOX5,ANK3,DHRS11,DPYD,CADPS,MAML3,TSHZ3,PTPRO,GBF1,PPI

JMJD1C,PPP2R2B,NFIX,SRPK2,IMMP2L,PCDH9,THRB,MACROD2,RPS6KL1,SORBS1,BCL11B,AFF3,NAA

RHOJ,TFB1M,CALD1,FURIN,SRR,PPP1R16B,ZCCHC7,FOXO3,SEMA3F,MIRLET7BHG,WNT3,CLCN3,PPP2

,MDK,PKP4,CACNA2D2,MPHOSPH9,CNTN4,NEDD4L,MMP16,EXT1,ITGA11,CKB,LDB1,GATB,PRKAG2,AU

,CADM2,CTNND1,CD55,STAG1,PTPRF,DHRS11,LRFN5,JMJD1C,PIP5K1B,KEL,ZFYVE1,PPP2R2B,NFIX,TI

N2,CTR9,FHIT,ZFPM2,EXT1,CALU,DMTF1,LDB1,ZNF638,PRPF3,FGFR1,MSH5,CNOT1,XRCC6,SDK1,ME

ıe genes for some small nuclear RNAs (snRNA) and of genes such as those for histone H2B and immunoglobu

yme survival. May also induce epithelial to mesenchymal transition (EMT) through the expression of SNAI1.

5,PRKAG2,AUTS2,CREB5,MEF2C,GMIP,DDX3Y,DBN1,CALD1,RASGRP4,CADM2,CTNND1,SOX5,IMPA2,S

PPP1R16B,ESRRB,CADPS,LRFN5,JMJD1C,RNF111,KCNN2,ZFYVE1,RGS6,MANBA,NFIX,NOVA1,PCDH9,

B1,GATB,FGFR1,LRRC20,CNOT1,AUTS2,ETF1,CALD1,CADM2,CTNND1,SOX5,STAG1,CACNB2,PTPRF,F[illegible]

:REB5,MEF2C,GMIP,BTN2A1,CALD1,SOX5,CTNNB1,STAG1,ANK3,MED27,PPP1R16B,ESRRB,BCL11A,FO

[illegible]TS2,SDK1,CARMIL1,MEF2C,DDX3Y,MGAT5B,CADM2,CTNND1,STAG1,SYT1,CACNB2,KCNB1,ANK3,CN[illegible]

F2C,GRIK1,MGAT5B,CADM2,SOX5,CD55,STAG1,ABCB9,SRR,CADPS,LRFN5,BCL11A,ZCCHC7,FOXO3,G

TAG1,CACNB2,ESRRB,MAML3,BCL11A,PTPRO,JMJD1C,PDE4B,KCNN2,PPP1R13B,NFIX,SRPK2,TNXB,E

URIN,CNNM2,PPP1R16B,CADPS,LRFN5,BCL11A,ZCCHC7,FOXO3,KDM4A,KEL,NKIRAS1,RNF111,ZFYVE[illegible]

[illegible]XO3,PTPRO,JMJD1C,SEMA3F,ZFYVE1,PPP2R2B,NFIX,WNT3,PCDH9,SLC39A8,DOCK4,KDM3B,BCL11B,

IM2,PPP1R16B,LRFN5,SLC35F2,BCL11A,PTPRO,JMJD1C,PIP5K1B,PDE4B,SEMA3F,KCNN2,PARD3B,PPF

BF1,KCNN2,PPP2R2B,MANBA,NTRK3,SYNE1,DPP8,IMMP2L,WNT3,MACROD2,DOCK4,TRIM33,BCL11B,N

1,PPP2R2B,NFIX,NTRK3,TNXB,SYNE1,WNT3,PCDH9,THRB,RPS6KL1,DOCK4,TRIM33,SORBS1,BCL11B,[illegible]

P2R2B,RGS6,PPP1R13B,NFIX,NTRK3,NEGR1,SYNE1,ELMO1,WNT3,KDM3B,SORBS1,BCL11B,NEURL1,AI

RHOA,CAMKV,TLK1

=F3,HDAC9,ZNF800
