## Supplemental table 1 for "Digestive Dimensions of Autism: A Multiscale Exploration of Gut-Brain Interactions"

| Pathway | Total | Expected | Hits | P.Value | FDR | Genes |
|---|---|---|---|---|---|---|
| Regulation of neurogenesis | 444 | 12 | 32 | 4.78E-07 | 0.000392 | NREP,SYNGAP1,SEMA3A,FEZ1,M/ |
| Regulation of axonogenesis | 104 | 2.82 | 11 | 0.000117 | 0.0478 | SYNGAP1,SEMA3A,MAPT,AMIGO1 |
| Regulation of cell morphogenesis | 325 | 8.8 | 21 | 0.000216 | 0.0591 | SEMA3E,ARHGAP15,SYNGAP1,SE |
| Developmental growth | 290 | 7.85 | 17 | 0.00242 | 0.464 | KMT2D,FOXP1,SEMA3A,MAPT,ZM |
| Cell morphogenesis involved in differentiation | 827 | 22.4 | 36 | 0.00336 | 0.464 | NREP,SEMA3E,SYNGAP1,FOXP1,S |
| Axonogenesis | 578 | 15.7 | 27 | 0.00425 | 0.464 | NREP,SEMA3E,SYNGAP1,FOXP1,S |
| Generation of neurons | 1300 | 35.3 | 51 | 0.00475 | 0.464 | NREP,SEMA3E,SYNGAP1,FOXP1,S |
| Neuron projection development | 816 | 22.1 | 35 | 0.0048 | 0.464 | NREP,SEMA3E,SYNGAP1,FOXP1,S |
| Axon guidance | 394 | 10.7 | 20 | 0.00545 | 0.464 | SEMA3E,FOXP1,SEMA3A,FEZ1,CN |
| Regulation of cellular component size | 192 | 5.2 | 12 | 0.00615 | 0.464 | SEMA3A,MAPT,RPTOR,TWF2,BCL |
| Neurogenesis | 1390 | 37.7 | 53 | 0.00672 | 0.464 | NREP,SEMA3E,ADAM22,SYNGAP1 |
| Transcription initiation from RNA polymerase II promoter | 219 | 5.93 | 13 | 0.00679 | 0.464 | ESR2,ESRRG,TCF4,NEDD4L,NR1D |
| Central nervous system development | 784 | 21.2 | 33 | 0.00801 | 0.486 | SLC8A3,ADAM22,FOXP1,SEMA3A, |

| | | | | | | |
|---|---|---|---|---|---|---|
| Neuron differentiation | 1190 | 32.1 | 46 | 0.00841 | 0.486 | NREP,SEMA3E,SYNGAP1,FOXP1,[illegible] |
| Neuron development | 945 | 25.6 | 38 | 0.0094 | 0.486 | NREP,SEMA3E,SYNGAP1,FOXP1,[illegible] |
| Regulation of cell differentiation | 1290 | 34.9 | 49 | 0.00948 | 0.486 | NREP,HLA-B,KMT2E,SYNGAP1,FC[illegible] |
| DNA_dependent transcription, initiation | 257 | 6.96 | 14 | 0.0105 | 0.505 | ESR2,ESRRG,TAF1C,TCF4,NEDD[illegible] |
| Brain development | 559 | 15.1 | 24 | 0.0178 | 0.812 | SLC8A3,SEMA3A,ESR2,PTCH1,MD[illegible] |
| Cell development | 1840 | 49.9 | 64 | 0.0211 | 0.903 | NREP,SEMA3E,KMT2D,ADAM22,S[illegible] |
| Positive regulation of cell differentiation | 571 | 15.5 | 24 | 0.0223 | 0.903 | KMT2E,FOXP1,HLA-DMA,FEZ1,MA[illegible] |
| Regulation of GTPase activity | 257 | 6.96 | 13 | 0.0231 | 0.903 | PTPRN2,DGKI,ARHGAP15,SYNGA[illegible] |
| Regulation of anatomical structure morphogenesis | 702 | 19 | 28 | 0.0264 | 0.982 | SEMA3E,ARHGAP15,SYNGAP1,FC[illegible] |
| Regulation of Ras protein signal transduction | 302 | 8.18 | 14 | 0.036 | 1 | DGKI,ARHGAP15,SYNGAP1,PKP4,[illegible] |
| Detection of biotic stimulus | 28 | 0.758 | 3 | 0.0392 | 1 | HLA-B,HLA-DRB5,HLA-DRB1 |
| Locomotory behavior | 147 | 3.98 | 8 | 0.0467 | 1 | NPAS3,ADAM22,GIGYF2,MAPT,CH[illegible] |
| Regulation of Ras GTPase activity | 176 | 4.77 | 9 | 0.0504 | 1 | DGKI,ARHGAP15,SYNGAP1,PKP4,[illegible] |

| | | | | | | |
|---|---|---|---|---|---|---|
| NLS_bearing substrate import into nucleus | 14 | 0.379 | 2 | 0.0537 | 1 | RERE,CBLB |
| Regulation of Rho GTPase activity | 101 | 2.74 | 6 | 0.0564 | 1 | ARHGAP15,PKP4,MYO9B,GMIP,PF |
| Behavior | 597 | 16.2 | 23 | 0.0571 | 1 | NPAS3,ADAM22,SYNGAP1,SEMA3 |
| Negative regulation of cell differentiation | 540 | 14.6 | 21 | 0.0619 | 1 | SYNGAP1,SEMA3A,PTCH1,F2,ZNF |
| Cell recognition | 80 | 2.17 | 5 | 0.0658 | 1 | SEMA3A,CNTNAP2,CNTN4,AMIGO |
| Ras protein signal transduction | 274 | 7.42 | 12 | 0.07 | 1 | DGKI,SYNGAP1,RSU1,MYO9B,CUL |
| Transmission of nerve impulse | 871 | 23.6 | 31 | 0.0725 | 1 | DPP6,GNAI1,ADAM22,SYNGAP1,G |
| Nervous system development | 2190 | 59.4 | 70 | 0.0765 | 1 | NRGN,NREP,SLC8A3,SEMA3E,AD |
| Negative regulation of cellular component organization | 370 | 10 | 15 | 0.079 | 1 | SYNGAP1,SEMA3A,PRKD1,DPP4, |
| Regulation of small GTPase mediated signal transduction | 444 | 12 | 17 | 0.0963 | 1 | DGKI,ARHGAP15,SYNGAP1,PKP4, |
| Chromatin modification | 512 | 13.9 | 19 | 0.103 | 1 | KDM5D,KMT2D,KMT2A,MSL2,KMT |

| | | | | | | |
|---|---|---|---|---|---|---|
| Regulation of Rho protein signal transduction | 177 | 4.79 | 8 | 0.109 | 1 | ARHGAP15,PKP4,MYO9B,CUL3,G[illegible] |
| Female gamete generation | 94 | 2.55 | 5 | 0.112 | 1 | KMT2D,ZMIZ1,CTNNB1,FOXO3,IM[illegible] |
| Regulation of catabolic process | 595 | 16.1 | 21 | 0.131 | 1 | PTPRN2,DGKI,ARHGAP15,CISD2,[illegible] |
| Excretion | 72 | 1.95 | 4 | 0.131 | 1 | MDK,NEDD4L,CHRNA3,CACNA1C |
| Cell_cell junction organization | 186 | 5.04 | 8 | 0.133 | 1 | PKP4,NEDD4L,CADM2,CTNND1,C[illegible] |
| Regulation of T cell activation | 279 | 7.56 | 11 | 0.138 | 1 | HLA-DMA,DPP4,CBLB,HLA-DQA1,[illegible] |
| Positive regulation of transcription from RNA polymerase II promoter | 800 | 21.7 | 27 | 0.14 | 1 | KMT2D,KMT2A,FOXP1,PRKD1,TC[illegible] |
| Histone modification | 312 | 8.45 | 12 | 0.142 | 1 | KDM5D,KMT2D,KMT2A,MSL2,KMT[illegible] |
| Growth | 839 | 22.7 | 28 | 0.148 | 1 | KMT2D,FOXP1,SEMA3A,GIGYF2,E[illegible] |
| Negative regulation of developmental process | 674 | 18.3 | 23 | 0.151 | 1 | SEMA3E,SYNGAP1,SEMA3A,PTC[illegible] |
| Regulation of cell shape | 106 | 2.87 | 5 | 0.16 | 1 | SEMA3E,ARHGAP15,F2,RHOJ,BR[illegible] |
| Covalent chromatin modification | 320 | 8.67 | 12 | 0.161 | 1 | KDM5D,KMT2D,KMT2A,MSL2,KMT[illegible] |
| Cell_cell junction assembly | 78 | 2.11 | 4 | 0.161 | 1 | PKP4,NEDD4L,PTPRO,RHOA |

| | | | | | | |
|---|---|---|---|---|---|---|
| Regulation of transforming growth factor beta receptor signaling pathway | 107 | 2.9 | 5 | 0.165 | 1 | NREP,NEDD4L,CITED1,RNF111,TF |
| Protein O_linked glycosylation | 79 | 2.14 | 4 | 0.166 | 1 | GALNT12,ST3GAL3,GALNT1,GALN |
| Skeletal muscle tissue development | 197 | 5.34 | 8 | 0.167 | 1 | FOXP1,CACNA2D2,HDAC4,MEF2C |
| Neurotransmitter secretion | 108 | 2.93 | 5 | 0.169 | 1 | TSNARE1,PCLO,CHRNA3,SYT1,CA |
| Detection of external stimulus | 108 | 2.93 | 5 | 0.169 | 1 | HLA-B,COL11A1,HLA-DRB5,HLA-D |
| Cytoplasm organization | 7 | 0.19 | 1 | 0.175 | 1 | ZMIZ1 |
| Cytokine secretion | 110 | 2.98 | 5 | 0.178 | 1 | HLA-DRB5,BTN3A2,HLA-DRB1,BTN |
| B cell differentiation | 82 | 2.22 | 4 | 0.182 | 1 | FOXP1,HDAC4,BCL11A,HDAC9 |
| Positive regulation of T cell activation | 202 | 5.47 | 8 | 0.183 | 1 | HLA-DMA,DPP4,HLA-DQA1,HLA-DI |
| Heterophilic cell_cell adhesion | 29 | 0.785 | 2 | 0.185 | 1 | AMIGO1,TENM2 |
| Actin filament bundle assembly | 83 | 2.25 | 4 | 0.188 | 1 | CUL3,PPM1E,SORBS1,RHOA |
| Synaptic transmission | 765 | 20.7 | 25 | 0.191 | 1 | GNAI1,SYNGAP1,GRIN2A,TSNARE |
| Positive regulation of hydrolase activity | 497 | 13.5 | 17 | 0.193 | 1 | PDE1C,ARHGAP15,SYNGAP1,WRI |
| T cell activation | 432 | 11.7 | 15 | 0.196 | 1 | HLA-DMA,DPP4,CBLB,HLA-DQA1,( |

| | | | | | | |
|---|---|---|---|---|---|---|
| Regulation of myeloid cell differentiation | 145 | 3.93 | 6 | 0.2 | 1 | KMT2E,CTR9,LDB1,MEF2C,CTNNB[illegible] |
| Gland development | 303 | 8.21 | 11 | 0.201 | 1 | SEMA3A,ESR2,PTCH1,MDK,FGFR[illegible] |
| Lymphocyte activation | 601 | 16.3 | 20 | 0.201 | 1 | FOXP1,HLA-DMA,BANK1,DPP4,HD[illegible] |
| Gliogenesis | 176 | 4.77 | 7 | 0.201 | 1 | ADAM22,F2,SOX5,CTNNB1,NTRK3[illegible] |
| Ribonucleotide metabolic process | 805 | 21.8 | 26 | 0.201 | 1 | PTPRN2,DGKI,ARHGAP15,GNAI1,[illegible] |
| Regulation of protein polymerization | 115 | 3.11 | 5 | 0.202 | 1 | MAPT,TWF2,PREX1,RHOA,TRIOB[illegible] |
| Actin cytoskeleton organization | 502 | 13.6 | 17 | 0.204 | 1 | SEMA3E,MYO18A,FOXP1,CUL3,P[illegible] |
| Maintenance of protein location | 116 | 3.14 | 5 | 0.206 | 1 | DBN1,CTNNB1,ANK3,TWF2,SYNE[illegible] |
| T cell proliferation | 147 | 3.98 | 6 | 0.209 | 1 | CBLB,CTNNB1,HLA-DRB5,HLA-DR[illegible] |
| Regulation of membrane potential | 306 | 8.29 | 11 | 0.209 | 1 | DPP6,ADAM22,GRIN2A,NEDD4L,A[illegible] |
| Cell maturation | 117 | 3.17 | 5 | 0.211 | 1 | ESR2,CNTNAP2,FGFR1,CTNNB1,[illegible] |
| Regulation of heart contraction | 148 | 4.01 | 6 | 0.213 | 1 | SEMA3A,KCNG2,HDAC4,ATP2A2,[illegible] |
| Actin filament_based process | 577 | 15.6 | 19 | 0.221 | 1 | SEMA3E,MYO18A,FOXP1,UTY,MY[illegible] |
| Negative regulation of myeloid cell differentiation | 60 | 1.63 | 3 | 0.222 | 1 | CTR9,LDB1,CTNNB1 |

| | | | | | | |
|---|---|---|---|---|---|---|
| Regulation of neuron apoptotic process | 150 | 4.06 | 6 | 0.222 | 1 | CITED1,MEF2C,FURIN,PPP2R2B,S |
| Positive regulation of developmental process | 817 | 22.1 | 26 | 0.223 | 1 | KMT2E,FOXP1,HLA-DMA,FEZ1,MA |
| Regulation of T cell proliferation | 120 | 3.25 | 5 | 0.226 | 1 | CBLB,CTNNB1,HLA-DRB5,HLA-DR |
| Rho protein signal transduction | 121 | 3.28 | 5 | 0.231 | 1 | MYO9B,CUL3,RHOJ,NGEF,RHOA |
| Purine ribonucleotide metabolic process | 787 | 21.3 | 25 | 0.231 | 1 | PTPRN2,DGKI,ARHGAP15,GNAI1,S |
| Interferon_gamma production | 91 | 2.46 | 4 | 0.233 | 1 | HLA-DRB5,BTN3A2,HLA-DRB1,BTN |
| Cellular carbohydrate catabolic process | 62 | 1.68 | 3 | 0.236 | 1 | HDAC4,GPD2,PRKAG2 |
| Actin filament organization | 250 | 6.77 | 9 | 0.238 | 1 | CUL3,PPM1E,CARMIL1,DBN1,TWF |
| Positive regulation of lymphocyte activation | 250 | 6.77 | 9 | 0.238 | 1 | HLA-DMA,DPP4,HLA-DQA1,MEF2C |

| | | | | | | |
|---|---|---|---|---|---|---|
| Adaptive immune response based on somatic recombination of immune receptors built from immunoglobulin superfamily domains | 218 | 5.9 | 8 | 0.24 | 1 | HLA-DMA,MEF2C,CD55,ABCB9,HL |
| Regulation of gene expression, epigenetic | 155 | 4.2 | 6 | 0.244 | 1 | KMT2D,KMT2E,UTY,TET2,TNRC6A |
| Cell fate commitment | 254 | 6.88 | 9 | 0.252 | 1 | CTR9,FGFR1,MEF2C,SOX5,CTNNI |
| Striated muscle tissue development | 354 | 9.59 | 12 | 0.253 | 1 | FOXP1,CACNA2D2,HDAC4,ZFPM2 |
| Cellular monovalent inorganic cation homeostasis | 36 | 0.975 | 2 | 0.255 | 1 | NEDD4L,CLCN3 |
| Muscle organ development | 389 | 10.5 | 13 | 0.258 | 1 | FOXP1,CACNA2D2,HDAC4,ZFPM2 |
| Positive regulation of cellular component organization | 560 | 15.2 | 18 | 0.26 | 1 | FEZ1,MAPT,PRKD1,CTR9,PPM1E,I |
| Glycoprotein catabolic process | 11 | 0.298 | 1 | 0.261 | 1 | MANBA |
| Cell_matrix adhesion | 159 | 4.31 | 6 | 0.262 | 1 | SEMA3E,ITGA11,CTNNB1,TNXB,S |
| Negative regulation of cell adhesion | 96 | 2.6 | 4 | 0.263 | 1 | SEMA3E,ADAM22,PRKG1,PTPRO |

| | | | | | | |
|---|---|---|---|---|---|---|
| Small GTPase mediated signal transduction | 596 | 16.1 | 19 | 0.264 | 1 | DGKI,ARHGAP15,SYNGAP1,RSU1, |
| Insulin receptor signaling pathway | 192 | 5.2 | 7 | 0.265 | 1 | RPTOR,FGFR1,PRKAG2,FOXO3,S |
| Generation of a signal involved in cell_cell signaling | 393 | 10.6 | 13 | 0.269 | 1 | TSNARE1,PDE8B,CACNA2D2,KCN |
| Synapse organization | 161 | 4.36 | 6 | 0.271 | 1 | SEMA3E,CACNA2D2,CTNNB1,CAC |
| Regulation of lymphocyte activation | 360 | 9.75 | 12 | 0.272 | 1 | HLA-DMA,DPP4,CBLB,HLA-DQA1,I |
| Androgen receptor signaling pathway | 67 | 1.81 | 3 | 0.272 | 1 | ESR2,CTNNB1,RHOA |
| Regulation of hydrolase activity | 843 | 22.8 | 26 | 0.273 | 1 | PDE1C,PTPRN2,DGKI,ARHGAP15, |
| Purine nucleotide metabolic process | 844 | 22.9 | 26 | 0.275 | 1 | PTPRN2,DGKI,ARHGAP15,GNAI1,S |
| Dephosphorylation | 295 | 7.99 | 10 | 0.279 | 1 | PTPRN2,PPM1E,MEF2C,MAGI2,IM |
| Leukocyte activation | 707 | 19.1 | 22 | 0.28 | 1 | KMT2E,FOXP1,HLA-DMA,BANK1,D |
| Gene silencing | 99 | 2.68 | 4 | 0.281 | 1 | KMT2D,UTY,TNRC6A,SND1 |
| Post_translational protein modification | 196 | 5.31 | 7 | 0.282 | 1 | GALNT12,F2,ST3GAL3,FURIN,RFT |
| Potassium ion transport | 131 | 3.55 | 5 | 0.282 | 1 | DPP6,KCNG2,NEDD4L,ANK3,KCNI |

| | | | | | | |
|---|---|---|---|---|---|---|
| Regulation of developmental process | 1880 | 50.9 | 55 | 0.286 | 1 | NREP,SEMA3E,HLA-B,KMT2E,ARH |
| Pattern specification process | 503 | 13.6 | 16 | 0.29 | 1 | SEMA3E,SYNGAP1,FOXP1,PTCH1 |
| Maintenance of protein location in cell | 101 | 2.74 | 4 | 0.293 | 1 | DBN1,CTNNB1,ANK3,SYNE1 |
| Negative regulation of translation | 70 | 1.9 | 3 | 0.295 | 1 | GIGYF2,TNRC6A,FXR1 |
| Regulation of cell growth | 302 | 8.18 | 10 | 0.303 | 1 | SEMA3A,ESR2,MAPT,F2,TWF2,BC |
| Regulation of action potential | 169 | 4.58 | 6 | 0.309 | 1 | DPP6,ADAM22,AMIGO1,CTNNB1,A |
| Nucleotide_excision repair | 104 | 2.82 | 4 | 0.311 | 1 | RPA3,SLC30A9,RFC2,POLR2E |
| Regulation of growth | 548 | 14.8 | 17 | 0.317 | 1 | FOXP1,SEMA3A,ESR2,MAPT,WRN |
| Positive regulation of transcription, DNA_dependent | 1260 | 34.1 | 37 | 0.323 | 1 | NPAS3,KMT2D,KMT2A,KMT2E,FOX |
| Positive regulation of transcription, DNA_dependent | 1260 | 34.1 | 37 | 0.323 | 1 | NPAS3,KMT2D,KMT2A,KMT2E,FOX |
| Negative regulation of transcription from RNA polymerase II promoter | 552 | 15 | 17 | 0.328 | 1 | ESR2,PTCH1,TCF4,CTR9,HDAC4,Z |

| | | | | | | |
|---|---|---|---|---|---|---|
| Actin filament polymerization | 107 | 2.9 | 4 | 0.33 | 1 | TWF2,PREX1,RHOA,TRIOBP |
| Regulation of actin polymerization or depolymerization | 107 | 2.9 | 4 | 0.33 | 1 | TWF2,PREX1,RHOA,TRIOBP |
| Adaptive immune response | 241 | 6.53 | 8 | 0.33 | 1 | HLA-DMA,MEF2C,CD55,ABCB9,HL |
| Cell_substrate adhesion | 241 | 6.53 | 8 | 0.33 | 1 | SEMA3E,ITGA11,CTNNB1,PTPRO, |
| Protein processing | 140 | 3.79 | 5 | 0.33 | 1 | PTCH1,CD55,FURIN,IMMP2L,GLI3 |
| Response to drug | 344 | 9.32 | 11 | 0.33 | 1 | KMT2A,GRIN2A,MDK,HDAC4,PPM[illegible] |
| Myoblast differentiation | 44 | 1.19 | 2 | 0.335 | 1 | MEF2C,CTNNB1 |
| Regulation of actin filament length | 108 | 2.93 | 4 | 0.336 | 1 | TWF2,PREX1,RHOA,TRIOBP |
| Developmental maturation | 177 | 4.79 | 6 | 0.347 | 1 | ESR2,CNTNAP2,FGFR1,CTNNB1,[illegible] |
| Positive regulation of cytoskeleton organization | 110 | 2.98 | 4 | 0.348 | 1 | MAPT,PPM1E,NTRK3,RHOA |
| DNA recombination | 247 | 6.69 | 8 | 0.354 | 1 | RPA3,FOXP1,WRN,MSH5,XRCC6,[illegible] |
| Regulation of hormone secretion | 213 | 5.77 | 7 | 0.356 | 1 | PDE8B,CACNA2D2,KCNG2,CACN[illegible] |
| Microtubule_based movement | 179 | 4.85 | 6 | 0.357 | 1 | MAPT,DYNC1I1,KLC1,KIF5C,KIF21 |
| Maintenance of location in cell | 112 | 3.03 | 4 | 0.36 | 1 | DBN1,CTNNB1,ANK3,SYNE1 |

| | | | | | | |
|---|---|---|---|---|---|---|
| Post_Golgi vesicle_mediated transport | 79 | 2.14 | 3 | 0.362 | 1 | ARFGEF2,ANK3,GBF1 |
| Chromatin organization | 637 | 17.3 | 19 | 0.366 | 1 | KDM5D,KMT2D,KMT2A,MSL2,KMT |
| Intracellular steroid hormone receptor signaling pathway | 113 | 3.06 | 4 | 0.367 | 1 | KMT2D,ESR2,CTNNB1,RHOA |
| Detection of stimulus | 216 | 5.85 | 7 | 0.369 | 1 | HLA-B,PKD1L3,COL11A1,SYT1,HL |
| Cell activation | 960 | 26 | 28 | 0.369 | 1 | DGKI,KMT2E,GNAI1,FOXP1,HLA-D |
| Phosphatidylinositol_mediated signaling | 148 | 4.01 | 5 | 0.373 | 1 | F2,FGFR1,FOXO3,RHOA,PIK3R2 |
| Vesicle localization | 82 | 2.22 | 3 | 0.384 | 1 | DYNC1I1,CUL3,GBF1 |
| Transforming growth factor beta receptor signaling pathway | 221 | 5.99 | 7 | 0.391 | 1 | NREP,NEDD4L,CITED1,FURIN,RN |
| Positive regulation of RNA metabolic process | 1330 | 36.1 | 38 | 0.394 | 1 | NPAS3,KMT2D,KMT2A,KMT2E,FO |
| DNA_dependent transcription, elongation | 118 | 3.2 | 4 | 0.397 | 1 | TAF1C,CTR9,LDB1,POLR2E |
| TRNA processing | 84 | 2.28 | 3 | 0.399 | 1 | TYW1B,PUS7,CDKAL1 |
| Protein tetramerization | 84 | 2.28 | 3 | 0.399 | 1 | SRR,HLA-DRB5,HLA-DRB1 |
| Protein maturation | 153 | 4.14 | 5 | 0.4 | 1 | PTCH1,CD55,FURIN,IMMP2L,GLI3 |
| Regulation of cell adhesion | 294 | 7.96 | 9 | 0.402 | 1 | SEMA3E,ADAM22,DPP4,PKP4,LDB |

| | | | | | | |
|---|---|---|---|---|---|---|
| Lymphocyte differentiation | 260 | 7.04 | 8 | 0.408 | 1 | FOXP1,HLA-DMA,HDAC4,CTNNB1 |
| Cytokinesis | 120 | 3.25 | 4 | 0.41 | 1 | PKP4,CUL3,ANK3,RHOA |
| Glutamate receptor signaling pathway | 52 | 1.41 | 2 | 0.413 | 1 | GRIN2A,GRIK1 |
| Hormone secretion | 262 | 7.1 | 8 | 0.416 | 1 | PDE8B,CACNA2D2,KCNG2,PCLO,[illegible] |
| Regulation of neurotransmitter levels | 157 | 4.25 | 5 | 0.421 | 1 | TSNARE1,PCLO,CHRNA3,SYT1,C[illegible] |
| Humoral immune response | 157 | 4.25 | 5 | 0.421 | 1 | MEF2C,CD55,HLA-DRB5,HLA-DRB[illegible] |
| Regulation of chromosome organization | 122 | 3.3 | 4 | 0.422 | 1 | PRKD1,CTR9,CTNNB1,TLK1 |
| Regulation of translation | 228 | 6.18 | 7 | 0.422 | 1 | GIGYF2,TNRC6A,FXR1,ETF1,CPE[illegible] |
| Calcium ion transport | 300 | 8.13 | 9 | 0.425 | 1 | SLC8A3,F2,ATP2A2,CACNA1C,CT[illegible] |
| Cell projection assembly | 229 | 6.2 | 7 | 0.427 | 1 | SPAG16,CARMIL1,TWF2,PPP1R16[illegible] |
| Chromosome organization | 878 | 23.8 | 25 | 0.427 | 1 | KDM5D,KMT2D,KMT2A,MSL2,RPA[illegible] |
| Regulation of cytokine secretion | 88 | 2.38 | 3 | 0.428 | 1 | HLA-DRB5,HLA-DRB1,BTN2A2 |
| B cell activation | 194 | 5.25 | 6 | 0.429 | 1 | FOXP1,BANK1,HDAC4,MEF2C,BC[illegible] |
| Smooth muscle contraction | 89 | 2.41 | 3 | 0.435 | 1 | CHRNA3,CACNA1C,TSHZ3 |
| Homeostasis of number of cells | 196 | 5.31 | 6 | 0.439 | 1 | KMT2E,GIGYF2,LDB1,MEF2C,FOX[illegible] |

| | | | | | | |
|---|---|---|---|---|---|---|
| Establishment of organelle localization | 125 | 3.39 | 4 | 0.44 | 1 | FEZ1,MAPT,CUL3,GBF1 |
| Negative regulation of growth | 197 | 5.34 | 6 | 0.444 | 1 | SEMA3A,ESR2,PTCH1,BCL11A,SE |
| Metal ion transport | 631 | 17.1 | 18 | 0.445 | 1 | SLC8A3,DPP6,F2,HFE,KCNG2,NED |
| Mesoderm development | 126 | 3.41 | 4 | 0.446 | 1 | EXT1,FGFR1,PALB2,WNT3 |
| Cellular response to nutrient levels | 126 | 3.41 | 4 | 0.446 | 1 | WRN,HFE,TNRC6A,RPTOR |
| Regulation of cytoskeleton organization | 306 | 8.29 | 9 | 0.449 | 1 | SEMA3E,MAPT,PPM1E,CTNNB1,T\ |
| Intracellular receptor mediated signaling pathway | 270 | 7.31 | 8 | 0.449 | 1 | KMT2D,KMT2E,ESR2,ZNF536,NR1 |
| Activation of phospholipase C activity | 56 | 1.52 | 2 | 0.45 | 1 | PDE1C,RASGRP4 |
| Negative regulation of angiogenesis | 56 | 1.52 | 2 | 0.45 | 1 | SEMA3E,FOXO6 |
| Regulation of synapse structure and activity | 56 | 1.52 | 2 | 0.45 | 1 | SYNGAP1,MEF2C |
| Energy reserve metabolic process | 199 | 5.39 | 6 | 0.454 | 1 | CACNA2D2,KCNG2,CACNA1C,PRK |
| Ovulation cycle | 92 | 2.49 | 3 | 0.456 | 1 | ESR2,FOXO3,IMMP2L |
| Cyclic nucleotide metabolic process | 164 | 4.44 | 5 | 0.458 | 1 | PDE8B,GABBR1,GUCY1A2,PDE4B |

| | | | | | | |
|---|---|---|---|---|---|---|
| Blood coagulation | 564 | 15.3 | 16 | 0.461 | 1 | SLC8A3,DGKI,GNAI1,F2,KLC1,DO( |
| Cellular component disassembly | 310 | 8.4 | 9 | 0.464 | 1 | MAPT,DPP4,KLC1,MMP16,ETF1,C |
| Ectoderm development | 23 | 0.623 | 1 | 0.469 | 1 | CTNNB1 |
| Learning or memory | 166 | 4.5 | 5 | 0.469 | 1 | SYNGAP1,GRIN2A,MDK,CACNA1C |
| Hemopoiesis | 640 | 17.3 | 18 | 0.469 | 1 | KMT2A,HLA-B,KMT2E,FOXP1,HLA- |
| Production of molecular mediator of immune response | 130 | 3.52 | 4 | 0.47 | 1 | FOXP1,HLA-DRB5,HLA-DRB1,HLA- |
| Coagulation | 568 | 15.4 | 16 | 0.473 | 1 | SLC8A3,DGKI,GNAI1,F2,KLC1,DO( |
| Protein polymerization | 167 | 4.52 | 5 | 0.474 | 1 | MAPT,TWF2,PREX1,RHOA,TRIOBI |
| Establishment of vesicle localization | 59 | 1.6 | 2 | 0.477 | 1 | CUL3,GBF1 |
| Hemostasis | 570 | 15.4 | 16 | 0.478 | 1 | SLC8A3,DGKI,GNAI1,F2,KLC1,DO( |
| Cell_cell adhesion | 461 | 12.5 | 13 | 0.481 | 1 | ANXA9,CDHR4,DPP4,CLSTN2,CNT |
| Detection of stimulus involved in sensory perception | 96 | 2.6 | 3 | 0.484 | 1 | PKD1L3,COL11A1,ATP2B2 |
| Regulation of angiogenesis | 169 | 4.58 | 5 | 0.485 | 1 | SEMA3E,PRKD1,CTNNB1,FOXO6,I |
| Neuron apoptotic process | 169 | 4.58 | 5 | 0.485 | 1 | CITED1,MEF2C,PPP2R2B,SRPK2,I |
| Nucleus organization | 60 | 1.63 | 2 | 0.486 | 1 | SRPK2,SYNE1 |

| | | | | | | |
|---|---|---|---|---|---|---|
| Negative regulation of immune system process | 207 | 5.61 | 6 | 0.491 | 1 | CBLB,CD55,HLA-DRB5,HLA-DRB1, |
| Immune system development | 722 | 19.6 | 20 | 0.492 | 1 | KMT2A,HLA-B,KMT2E,FOXP1,HLA- |
| Protein oligomerization | 357 | 9.67 | 10 | 0.502 | 1 | KCNG2,MAGI2,CTNNB1,SYT1,KCN |
| Actin polymerization or depolymerization | 136 | 3.68 | 4 | 0.505 | 1 | TWF2,PREX1,RHOA,TRIOBP |
| TRNA metabolic process | 173 | 4.69 | 5 | 0.505 | 1 | TYW1B,GATB,PPA2,PUS7,CDKAL1 |
| Transition metal ion transport | 100 | 2.71 | 3 | 0.511 | 1 | HFE,SLC30A9,SLC39A8 |
| Regulation of homeostatic process | 324 | 8.78 | 9 | 0.517 | 1 | F2,CACNA1C,LDB1,CTNNB1,ANK3 |
| Transcription from RNA polymerase III promoter | 64 | 1.73 | 2 | 0.521 | 1 | RPTOR,POLR2E |
| Homophilic cell adhesion | 139 | 3.76 | 4 | 0.522 | 1 | CDHR4,CLSTN2,AMIGO1,PCDH9 |
| Double_strand break repair | 139 | 3.76 | 4 | 0.522 | 1 | RPA3,XRCC6,ATP23,PALB2 |
| Muscle cell differentiation | 363 | 9.83 | 10 | 0.524 | 1 | FOXP1,CACNA2D2,HDAC4,MEF2C |
| Notch signaling pathway | 177 | 4.79 | 5 | 0.526 | 1 | TNRC6A,FURIN,MAML3,NEURL1,H |
| Nucleotide metabolic process | 1040 | 28.3 | 28 | 0.549 | 1 | PTPRN2,DGKI,ARHGAP15,GNAI1,S |
| CAMP_mediated signaling | 29 | 0.785 | 1 | 0.549 | 1 | PCLO |

| | | | | | | |
|---|---|---|---|---|---|---|
| S phase of mitotic cell cycle | 144 | 3.9 | 4 | 0.55 | 1 | RPA3,GIGYF2,PSMA5,RFC2 |
| Myeloid cell differentiation | 296 | 8.02 | 8 | 0.553 | 1 | KMT2E,TET2,CTR9,LDB1,MEF2C,F |
| Proteoglycan biosynthetic process | 68 | 1.84 | 2 | 0.553 | 1 | EXT1,CTNNB1 |
| Calcium_mediated signaling | 107 | 2.9 | 3 | 0.557 | 1 | HDAC4,CACNA1C,TENM2 |
| Regulation of cellular pH | 30 | 0.813 | 1 | 0.562 | 1 | CLCN3 |
| Striated muscle contraction | 108 | 2.93 | 3 | 0.564 | 1 | UTY,HDAC4,CACNA1C |
| T cell differentiation | 186 | 5.04 | 5 | 0.57 | 1 | HLA-DMA,CTNNB1,BCL11A,BCL11 |
| Epithelial cell differentiation | 339 | 9.18 | 9 | 0.572 | 1 | ESR2,PTCH1,CITED1,MEF2C,MAG |
| Hematopoietic or lymphoid organ development | 679 | 18.4 | 18 | 0.572 | 1 | KMT2A,HLA-B,KMT2E,FOXP1,HLA- |
| Regulation of body fluid levels | 680 | 18.4 | 18 | 0.575 | 1 | SLC8A3,DGKI,GNAI1,F2,KLC1,DOC |
| Protein targeting to mitochondrion | 71 | 1.92 | 2 | 0.577 | 1 | PPP2R2B,IMMP2L |
| Mitotic cell cycle checkpoint | 149 | 4.04 | 4 | 0.577 | 1 | PSMA5,ANAPC4,MAD1L1,PPP2R5( |
| Cellular response to extracellular stimulus | 149 | 4.04 | 4 | 0.577 | 1 | WRN,HFE,TNRC6A,RPTOR |
| Protein homooligomerization | 226 | 6.12 | 6 | 0.577 | 1 | KCNG2,SYT1,KCNB1,SRR,BCL11A |

| | | | | | | |
|---|---|---|---|---|---|---|
| Chromatin remodeling | 111 | 3.01 | 3 | 0.582 | 1 | RERE,HDAC4,ARID1B |
| Organelle localization | 189 | 5.12 | 5 | 0.584 | 1 | FEZ1,MAPT,DYNC1I1,CUL3,GBF1 |
| Response to external stimulus | 1510 | 40.9 | 40 | 0.585 | 1 | SEMA3E,HLA-B,FOXP1,SEMA3A,G |
| Response to steroid hormone stimulus | 267 | 7.23 | 7 | 0.588 | 1 | KMT2D,MDK,CITED1,CTNNB1,PSF |
| Negative regulation of transcription, DNA_dependent | 987 | 26.7 | 26 | 0.589 | 1 | KMT2D,FOXP1,ESR2,UTY,PTCH1,[illegible] |
| Negative regulation of transcription, DNA_dependent | 987 | 26.7 | 26 | 0.589 | 1 | KMT2D,FOXP1,ESR2,UTY,PTCH1,[illegible] |
| Neurological system process | 1480 | 40 | 39 | 0.593 | 1 | DPP6,GNAI1,ADAM22,SYNGAP1,G |
| Sperm motility | 33 | 0.894 | 1 | 0.596 | 1 | NEURL1 |
| Maintenance of location | 192 | 5.2 | 5 | 0.598 | 1 | DBN1,CTNNB1,ANK3,TWF2,SYNE[illegible] |
| S phase | 153 | 4.14 | 4 | 0.598 | 1 | RPA3,GIGYF2,PSMA5,RFC2 |
| Viral reproduction | 803 | 21.7 | 21 | 0.599 | 1 | HLA-B,PSMA5,VRK2,ZSCAN9,KLC[illegible] |
| Body fluid secretion | 74 | 2 | 2 | 0.6 | 1 | ATP2B2,NEURL1 |
| Exocytosis | 309 | 8.37 | 8 | 0.602 | 1 | TSNARE1,EXOC4,PCLO,CALU,ARI |
| Cellular polysaccharide metabolic process | 115 | 3.11 | 3 | 0.606 | 1 | EXT1,PRKAG2,SORBS1 |
| Interaction with host | 426 | 11.5 | 11 | 0.608 | 1 | HLA-B,VRK2,KLC1,THOC7,NEDD4I |
| Embryonic morphogenesis | 581 | 15.7 | 15 | 0.613 | 1 | PTCH1,CTR9,CUL3,COL11A1,EXT[illegible] |

| | | | | | | |
|---|---|---|---|---|---|---|
| Cell cycle arrest | 428 | 11.6 | 11 | 0.614 | 1 | RPA3,KMT2E,PSMA5,ANAPC4,CUL |
| Amine transport | 76 | 2.06 | 2 | 0.614 | 1 | CHRNA3,GRIK1 |
| Cytoskeleton_dependent intracellular transport | 76 | 2.06 | 2 | 0.614 | 1 | MAPT,DYNC1I1 |
| Positive regulation of nucleobase_containing compound metabolic process | 1490 | 40.4 | 39 | 0.616 | 1 | NPAS3,KMT2D,KMT2A,KMT2E,FOX |
| Protein secretion | 196 | 5.31 | 5 | 0.616 | 1 | HLA-DRB5,BTN3A2,HLA-DRB1,BTN |
| Proteoglycan metabolic process | 117 | 3.17 | 3 | 0.618 | 1 | COL11A1,EXT1,CTNNB1 |
| DNA damage response, signal transduction by p53 class mediator | 117 | 3.17 | 3 | 0.618 | 1 | PSMA5,FOXO3,PPP2R5C |
| Cell division | 507 | 13.7 | 13 | 0.62 | 1 | GNAI1,PTCH1,MDK,PKP4,ANAPC4 |
| Regulation of cell_cell adhesion | 77 | 2.09 | 2 | 0.621 | 1 | DPP4,PRKG1 |
| Cellular membrane organization | 471 | 12.8 | 12 | 0.627 | 1 | SYNGAP1,PTCH1,CUL3,CNTNAP2 |
| Cellular component disassembly involved in execution phase of apoptosis | 78 | 2.11 | 2 | 0.628 | 1 | MAPT,CTNNB1 |

| | | | | | | |
|---|---|---|---|---|---|---|
| Actin filament_based movement | 78 | 2.11 | 2 | 0.628 | 1 | UTY,MYO9B |
| One_carbon metabolic process | 36 | 0.975 | 1 | 0.628 | 1 | MTHFD1L |
| Cyclic_nucleotide_mediated signaling | 36 | 0.975 | 1 | 0.628 | 1 | PCLO |
| Embryo development | 1080 | 29.4 | 28 | 0.633 | 1 | KMT2D,KMT2A,FOXP1,UTY,PTCH[illegible] |
| Positive regulation of epithelial cell proliferation | 120 | 3.25 | 3 | 0.635 | 1 | FOXP1,PRKD1,CTNNB1 |
| DNA_dependent DNA replication | 121 | 3.28 | 3 | 0.641 | 1 | RPA3,WRN,RFC2 |
| Negative regulation of cytoskeleton organization | 80 | 2.17 | 2 | 0.642 | 1 | TWF2,TRIOBP |
| Monovalent inorganic cation homeostasis | 80 | 2.17 | 2 | 0.642 | 1 | NEDD4L,CLCN3 |
| Protein complex assembly | 861 | 23.3 | 22 | 0.644 | 1 | MAGI1,KMT2A,HLA-DMA,MAPT,HF[illegible] |
| Extracellular structure organization | 242 | 6.55 | 6 | 0.644 | 1 | THSD4,DPP4,COL11A1,MMP16,FU[illegible] |
| Activation of immune response | 399 | 10.8 | 10 | 0.644 | 1 | CBLB,HLA-DQA1,MEF2C,CD55,MI[illegible] |
| Spermatid development | 81 | 2.19 | 2 | 0.649 | 1 | PPP2R2B,NEURL1 |

| | | | | | | |
|---|---|---|---|---|---|---|
| Morphogenesis of an epithelium | 440 | 11.9 | 11 | 0.65 | 1 | SEMA3E,SEMA3A,ESR2,PTCH1,CI |
| Cytoskeleton organization | 980 | 26.5 | 25 | 0.653 | 1 | SEMA3E,MYO18A,FOXP1,MAPT,KI |
| Spliceosomal complex assembly | 39 | 1.06 | 1 | 0.658 | 1 | SRPK2 |
| Leukocyte differentiation | 404 | 10.9 | 10 | 0.659 | 1 | HLA-B,KMT2E,FOXP1,HLA-DMA,HI |
| Golgi vesicle transport | 206 | 5.58 | 5 | 0.66 | 1 | PRKD1,CUL3,ARFGEF2,ANK3,GBF |
| DNA modification | 83 | 2.25 | 2 | 0.662 | 1 | KMT2E,TET2 |
| Second_messenger_mediated signaling | 166 | 4.5 | 4 | 0.662 | 1 | PCLO,HDAC4,CACNA1C,TENM2 |
| Negative regulation of RNA metabolic process | 1020 | 27.7 | 26 | 0.664 | 1 | KMT2D,FOXP1,ESR2,UTY,PTCH1,[illegible] |
| Mitotic spindle organization | 40 | 1.08 | 1 | 0.667 | 1 | RHOA |
| Polysaccharide metabolic process | 126 | 3.41 | 3 | 0.668 | 1 | EXT1,PRKAG2,SORBS1 |
| Endothelial cell migration | 126 | 3.41 | 3 | 0.668 | 1 | PRKD1,DPP4,HDAC9 |
| Heart development | 487 | 13.2 | 12 | 0.673 | 1 | FOXP1,UTY,PTCH1,ZFPM2,COL11 |
| Positive regulation of immune response | 487 | 13.2 | 12 | 0.673 | 1 | HLA-DMA,CBLB,HLA-DQA1,MEF2C |

| | | | | | | |
|---|---|---|---|---|---|---|
| Regulation of cellular component organization | 1520 | 41.3 | 39 | 0.673 | 1 | SEMA3E,ARHGAP15,SYNGAP1,SE |
| Regulation of muscle contraction | 127 | 3.44 | 3 | 0.673 | 1 | HDAC4,CHRNA3,CACNA1C |
| Positive regulation of cell adhesion | 127 | 3.44 | 3 | 0.673 | 1 | LDB1,RHOA,TRIOBP |
| Tissue morphogenesis | 566 | 15.3 | 14 | 0.675 | 1 | SEMA3E,SEMA3A,ESR2,PTCH1,ZF |
| Reciprocal meiotic recombination | 41 | 1.11 | 1 | 0.676 | 1 | MSH5 |
| Regulation of endocytosis | 128 | 3.47 | 3 | 0.678 | 1 | PRKD1,NEDD4L,MAGI2 |
| Spermatid differentiation | 86 | 2.33 | 2 | 0.681 | 1 | PPP2R2B,NEURL1 |
| Regulation of G_protein coupled receptor protein signaling pathway | 129 | 3.49 | 3 | 0.683 | 1 | F2,RASGRP4,RGS6 |
| Signal transduction in response to DNA damage | 129 | 3.49 | 3 | 0.683 | 1 | PSMA5,FOXO3,PPP2R5C |
| Positive regulation of transport | 570 | 15.4 | 14 | 0.685 | 1 | KMT2A,PTCH1,F2,NEDD4L,GRIK1, |
| Regulation of multicellular organismal process | 2480 | 67.1 | 64 | 0.686 | 1 | NREP,SEMA3E,HLA-B,KMT2E,SYN |

| | | | | | | |
|---|---|---|---|---|---|---|
| Protein dephosphorylation | 172 | 4.66 | 4 | 0.689 | 1 | PTPRN2,PPM1E,PTPRF,PTPRO |
| Cellular homeostasis | 883 | 23.9 | 22 | 0.69 | 1 | SLC8A3,DPP6,ADAM22,GRIN2A,TI |
| Secondary metabolic process | 44 | 1.19 | 1 | 0.702 | 1 | CITED1 |
| Focal adhesion assembly | 44 | 1.19 | 1 | 0.702 | 1 | SORBS1 |
| Cation transport | 851 | 23 | 21 | 0.703 | 1 | SLC8A3,DPP6,F2,HFE,KCNG2,PKD |
| Cation transport | 851 | 23 | 21 | 0.703 | 1 | SLC8A3,DPP6,F2,HFE,KCNG2,PKD |
| Neuropeptide signaling pathway | 90 | 2.44 | 2 | 0.705 | 1 | PKD1L3,SORCS3 |
| Cellular carbohydrate metabolic process | 259 | 7.02 | 6 | 0.707 | 1 | HDAC4,GPD2,EXT1,PRKAG2,IMPA |
| Negative regulation of protein metabolic process | 540 | 14.6 | 13 | 0.708 | 1 | GRIN2A,GIGYF2,PSMA5,F2,ANAP( |
| Adenylate cyclase_inhibiting G_protein coupled receptor signaling pathway | 45 | 1.22 | 1 | 0.71 | 1 | GNAI1 |
| Base_excision repair | 45 | 1.22 | 1 | 0.71 | 1 | WRN |
| Detection of chemical stimulus | 91 | 2.46 | 2 | 0.711 | 1 | PKD1L3,SYT1 |
| Wound healing | 700 | 19 | 17 | 0.714 | 1 | SLC8A3,DGKI,GNAI1,F2,KLC1,DO( |
| Negative regulation of cellular protein metabolic process | 463 | 12.5 | 11 | 0.714 | 1 | GIGYF2,PSMA5,F2,ANAPC4,USP4, |

| | | | | | | |
|---|---|---|---|---|---|---|
| Ion homeostasis | 818 | 22.2 | 20 | 0.715 | 1 | SLC8A3,DPP6,ADAM22,GRIN2A,F2 |
| Cellular protein complex assembly | 343 | 9.29 | 8 | 0.715 | 1 | MAPT,CHRNA3,SNAP91,TWF2,CA |
| Spindle organization | 92 | 2.49 | 2 | 0.716 | 1 | KIZ,RHOA |
| Detection of abiotic stimulus | 92 | 2.49 | 2 | 0.716 | 1 | COL11A1,ATP2B2 |
| Response to bacterium | 385 | 10.4 | 9 | 0.72 | 1 | HLA-B,CTR9,CITED1,MEF2C,SRR, |
| Negative regulation of cell proliferation | 585 | 15.8 | 14 | 0.721 | 1 | ESR2,PTCH1,HDAC4,CBLB,MAGI2 |
| Transmembrane receptor protein tyrosine kinase signaling pathway | 782 | 21.2 | 19 | 0.721 | 1 | PDE1C,GIGYF2,PRKD1,RPTOR,C |
| G1 phase of mitotic cell cycle | 47 | 1.27 | 1 | 0.725 | 1 | RBL2 |
| Centrosome cycle | 47 | 1.27 | 1 | 0.725 | 1 | CTNNB1 |
| Cellular component assembly | 1790 | 48.5 | 45 | 0.727 | 1 | MAGI1,KMT2A,RPA3,HLA-DMA,TH |
| Positive regulation of angiogenesis | 94 | 2.55 | 2 | 0.727 | 1 | PRKD1,HDAC9 |
| Regulation of organelle organization | 589 | 16 | 14 | 0.73 | 1 | SEMA3E,MAPT,PRKD1,ANAPC4,C |
| Glucan metabolic process | 95 | 2.57 | 2 | 0.733 | 1 | PRKAG2,SORBS1 |
| Cellular nitrogen compound catabolic process | 986 | 26.7 | 24 | 0.738 | 1 | PTPRN2,DGKI,RPA3,ARHGAP15,G |

| | | | | | | |
|---|---|---|---|---|---|---|
| Response to hormone stimulus | 751 | 20.3 | 18 | 0.738 | 1 | KMT2D,GNAI1,MDK,RPTOR,CITED |
| Regulation of pH | 49 | 1.33 | 1 | 0.74 | 1 | CLCN3 |
| Sensory perception of taste | 49 | 1.33 | 1 | 0.74 | 1 | PKD1L3 |
| G1 phase | 49 | 1.33 | 1 | 0.74 | 1 | RBL2 |
| Negative regulation of cellular biosyntheti c process | 1220 | 33.1 | 30 | 0.742 | 1 | KMT2D,FOXP1,GIGYF2,ESR2,UTY |
| Regionaliz ation | 353 | 9.56 | 8 | 0.744 | 1 | PTCH1,LDB1,FGFR1,MEF2C,PALB |
| Pyrimidine nucleotide metabolic process | 50 | 1.35 | 1 | 0.747 | 1 | DPYD |
| Amino acid transport | 143 | 3.87 | 3 | 0.748 | 1 | PRKAG2,GRIK1,SYT1 |
| Response to carbohydra te stimulus | 143 | 3.87 | 3 | 0.748 | 1 | HLA-DRB5,HLA-DRB1,RHOA |
| Regulation of secretion | 517 | 14 | 12 | 0.748 | 1 | PDE8B,CACNA2D2,KCNG2,PCLO,[illegible] |
| Regulation of mitosis | 99 | 2.68 | 2 | 0.753 | 1 | ANAPC4,MAD1L1 |
| Cell_cell signaling | 1310 | 35.4 | 32 | 0.753 | 1 | GNAI1,SYNGAP1,GRIN2A,ESR2,T[illegible] |
| Intrinsic apoptotic signaling pathway in response to DNA damage | 51 | 1.38 | 1 | 0.754 | 1 | PPP2R5C |
| Negative regulation of cell cycle | 520 | 14.1 | 12 | 0.755 | 1 | RPA3,KMT2E,PSMA5,ANAPC4,CU[illegible] |
| Regulation of protein secretion | 145 | 3.93 | 3 | 0.757 | 1 | HLA-DRB5,HLA-DRB1,BTN2A2 |

| | | | | | | |
|---|---|---|---|---|---|---|
| Macromolecular complex disassembly | 189 | 5.12 | 4 | 0.757 | 1 | KLC1,ETF1,TWF2,TRIOBP |
| Superoxide metabolic process | 52 | 1.41 | 1 | 0.761 | 1 | PREX1 |
| Carbohydrate transport | 147 | 3.98 | 3 | 0.765 | 1 | PRKAG2,RFT1,SORBS1 |
| Adenylate cyclase_activating G_protein coupled receptor signaling pathway | 53 | 1.44 | 1 | 0.767 | 1 | GPR26 |
| Protein autophosphorylation | 192 | 5.2 | 4 | 0.768 | 1 | VRK2,PRKD1,FGFR1,NTRK3 |
| Anatomical structure formation involved in morphogenesis | 2090 | 56.6 | 52 | 0.77 | 1 | NREP,SEMA3E,SYNGAP1,FOXP1,[illegible] |
| Negative regulation of biosynthetic process | 1240 | 33.6 | 30 | 0.773 | 1 | KMT2D,FOXP1,GIGYF2,ESR2,UTY |
| Positive regulation of I_kappaB kinase/NF_kappaB cascade | 150 | 4.06 | 3 | 0.777 | 1 | PRKD1,CTNNB1,RHOA |
| Cell cycle checkpoint | 281 | 7.61 | 6 | 0.777 | 1 | RPA3,PSMA5,ANAPC4,MAD1L1,R[illegible] |
| Respiratory gaseous exchange | 55 | 1.49 | 1 | 0.78 | 1 | TSHZ3 |
| Fatty acid biosynthetic process | 151 | 4.09 | 3 | 0.78 | 1 | AGMO,FADS2,PRKAG2 |

| | | | | | | |
|---|---|---|---|---|---|---|
| Negative regulation of nucleobase_containing compound metabolic process | 1130 | 30.6 | 27 | 0.782 | 1 | KMT2D,FOXP1,ESR2,UTY,PTCH1,[illegible] |
| Negative regulation of multicellular organismal process | 410 | 11.1 | 9 | 0.785 | 1 | PTCH1,F2,ATP2A2,MEF2C,GRIK1,[illegible] |
| Aging | 197 | 5.34 | 4 | 0.785 | 1 | CISD2,WRN,ZMIZ1,SRR |
| Positive regulation of translation | 56 | 1.52 | 1 | 0.786 | 1 | RHOA |
| Regulation of transcription from RNA polymerase II promoter | 1610 | 43.6 | 39 | 0.793 | 1 | KMT2D,KMT2A,FOXP1,ESR2,PTC[illegible] |
| Rhythmic process | 200 | 5.42 | 4 | 0.795 | 1 | ENOX1,ESR2,FOXO3,IMMP2L |
| Sulfur compound biosynthetic process | 155 | 4.2 | 3 | 0.795 | 1 | EXT1,CTNNB1,ST3GAL3 |
| Glycoprotein metabolic process | 458 | 12.4 | 10 | 0.799 | 1 | GALNT12,COL11A1,EXT1,MGAT5[illegible] |
| Development of primary sexual characteristics | 246 | 6.66 | 5 | 0.8 | 1 | ESR2,ZFPM2,CTNNB1,FOXO3,IM[illegible] |
| I_kappaB kinase/NF_kappaB cascade | 246 | 6.66 | 5 | 0.8 | 1 | PRKD1,CTNNB1,TANK,NKIRAS1,R[illegible] |
| Regulation of transport | 1220 | 33.2 | 29 | 0.803 | 1 | KMT2A,DPP6,MAPT,PTCH1,PRKD[illegible] |

| | | | | | | |
|---|---|---|---|---|---|---|
| Microtubule polymerization or depolymerization | 59 | 1.6 | 1 | 0.803 | 1 | MAPT |
| Secretion | 947 | 25.7 | 22 | 0.803 | 1 | TSNARE1,EXOC4,PDE8B,CACNA2 |
| Carbohydrate biosynthetic process | 203 | 5.5 | 4 | 0.804 | 1 | GPD2,EXT1,IMPA2,SORBS1 |
| Mitochondrial transport | 159 | 4.31 | 3 | 0.809 | 1 | PRKAG2,PPP2R2B,IMMP2L |
| Secretion by cell | 831 | 22.5 | 19 | 0.81 | 1 | TSNARE1,EXOC4,PDE8B,CACNA2 |
| Cellular protein complex disassembly | 160 | 4.33 | 3 | 0.812 | 1 | ETF1,TWF2,TRIOBP |
| Positive regulation of cell cycle | 113 | 3.06 | 2 | 0.815 | 1 | FGFR1,SRPK2 |
| ER to Golgi vesicle_mediated transport | 62 | 1.68 | 1 | 0.818 | 1 | CUL3 |
| Regulation of protein stability | 114 | 3.09 | 2 | 0.819 | 1 | USP4,CCDC88C |
| Negative regulation of hydrolase activity | 253 | 6.85 | 5 | 0.819 | 1 | PTPRN2,ITIH3,GMIP,ZFYVE1,CSRI |
| Bone remodeling | 63 | 1.71 | 1 | 0.823 | 1 | CTNNB1 |
| Macromolecular complex assembly | 1120 | 30.4 | 26 | 0.825 | 1 | MAGI1,KMT2A,RPA3,HLA-DMA,MA |
| Regulation of blood pressure | 164 | 4.44 | 3 | 0.825 | 1 | NEDD4L,CACNA1C,PTPRO |
| Viral reproductive process | 597 | 16.2 | 13 | 0.827 | 1 | HLA-B,VRK2,KLC1,THOC7,NEDD4I |
| Inorganic anion transport | 64 | 1.73 | 1 | 0.828 | 1 | SLC13A1 |

| | | | | | | |
|---|---|---|---|---|---|---|
| Transmembrane receptor protein serine/threonine kinase signaling pathway | 388 | 10.5 | 8 | 0.83 | 1 | NREP,NEDD4L,CITED1,MAGI2,FUI |
| Glucose catabolic process | 117 | 3.17 | 2 | 0.83 | 1 | HDAC4,PRKAG2 |
| Keratinocyte differentiation | 117 | 3.17 | 2 | 0.83 | 1 | CTNND1,BCL11B |
| Microtubule_based process | 516 | 14 | 11 | 0.831 | 1 | MAPT,KIZ,DYNC1I1,KLC1,KIF5C,C |
| Acute inflammatory response | 118 | 3.2 | 2 | 0.833 | 1 | F2,CD55 |
| Cellular cation homeostasis | 433 | 11.7 | 9 | 0.834 | 1 | SLC8A3,F2,HFE,NEDD4L,ATP2A2,( |
| Protein complex disassembly | 167 | 4.52 | 3 | 0.834 | 1 | ETF1,TWF2,TRIOBP |
| Protein N_linked glycosylation | 119 | 3.22 | 2 | 0.837 | 1 | MGAT5B, RFT1 |
| Phospholipase C_activating G_protein coupled receptor signaling pathway | 66 | 1.79 | 1 | 0.837 | 1 | F2 |
| Nuclear transport | 392 | 10.6 | 8 | 0.838 | 1 | RERE,RSRC1,WRN,THOC7,CBLB,( |
| Glycoprotein biosynthetic process | 393 | 10.6 | 8 | 0.84 | 1 | GALNT12,EXT1,MGAT5B,CTNNB1, |
| Phosphatidylinositol biosynthetic process | 120 | 3.25 | 2 | 0.84 | 1 | PIP5K1B,PIK3R2 |

| | | | | | | |
|---|---|---|---|---|---|---|
| Adenylate cyclase_modulating G_protein coupled receptor signaling pathway | 120 | 3.25 | 2 | 0.84 | 1 | GPR26,GNAI1 |
| Cell migration | 1050 | 28.5 | 24 | 0.84 | 1 | MYO18A,SEMA3A,ESR2,MAPT,PR[illegible] |
| Chemical homeostasis | 1050 | 28.5 | 24 | 0.84 | 1 | SLC8A3,DPP6,ADAM22,GRIN2A,P[illegible] |
| Response to heat | 67 | 1.81 | 1 | 0.842 | 1 | MICA |
| Viral genome replication | 67 | 1.81 | 1 | 0.842 | 1 | SRPK2 |
| Pigment biosynthetic process | 67 | 1.81 | 1 | 0.842 | 1 | CITED1 |
| Negative regulation of sequence_specific DNA binding transcription factor activity | 121 | 3.28 | 2 | 0.844 | 1 | PTCH1,HDAC4 |
| Nucleotide biosynthetic process | 352 | 9.53 | 7 | 0.845 | 1 | GABBR1,PFAS,ATP2A2,GUCY1A2, |
| Regulation of viral reproduction | 122 | 3.3 | 2 | 0.847 | 1 | POLR2E,SRPK2 |
| Ribonucleoprotein complex assembly | 124 | 3.36 | 2 | 0.853 | 1 | EFL1,SRPK2 |
| Chromosome segregation | 174 | 4.71 | 3 | 0.854 | 1 | MSH5,CTNNB1,STAG1 |
| Centrosome organization | 70 | 1.9 | 1 | 0.854 | 1 | CTNNB1 |
| Peptidyl_amino acid modification | 739 | 20 | 16 | 0.855 | 1 | FKBP9,KMT2A,MSL2,PRKD1,F2,C[illegible] |

| | | | | | | |
|---|---|---|---|---|---|---|
| Negative regulation of cellular metabolic process | 1660 | 45.1 | 39 | 0.855 | 1 | KMT2D,FOXP1,GIGYF2,PSMA5,ES |
| Negative regulation of metabolic process | 1820 | 49.4 | 43 | 0.856 | 1 | KMT2D,FOXP1,GRIN2A,GIGYF2,PS |
| Sodium ion transport | 125 | 3.39 | 2 | 0.856 | 1 | NEDD4L,ANK3 |
| Nucleobase_containing compound transport | 175 | 4.74 | 3 | 0.857 | 1 | THOC7,KIF5C,POM121C |
| Cellular modified amino acid biosynthetic process | 71 | 1.92 | 1 | 0.858 | 1 | MTHFD1L |
| Tissue remodeling | 126 | 3.41 | 2 | 0.859 | 1 | MEF2C,CTNNB1 |
| Protein polyubiquitination | 177 | 4.79 | 3 | 0.862 | 1 | PSMA5,ANAPC4,RNF111 |
| DNA catabolic process | 72 | 1.95 | 1 | 0.862 | 1 | RPA3 |
| Epidermis development | 319 | 8.64 | 6 | 0.867 | 1 | PTCH1,ATP2A2,LDB1,CTNND1,CT |
| Response to extracellular stimulus | 320 | 8.67 | 6 | 0.869 | 1 | WRN,HFE,TNRC6A,RPTOR,RASGI |
| Protein import into nucleus | 228 | 6.18 | 4 | 0.869 | 1 | RERE,CBLB,GLI3,RHOA |
| Microtubule organizing center organization | 74 | 2 | 1 | 0.87 | 1 | CTNNB1 |
| Response to toxin | 130 | 3.52 | 2 | 0.871 | 1 | RSRC1,MEF2C |
| Response to light stimulus | 229 | 6.2 | 4 | 0.872 | 1 | SYNGAP1,GRIN2A,WRN,CACNA1( |

| | | | | | | |
|---|---|---|---|---|---|---|
| Phosphatidylinositol metabolic process | 181 | 4.9 | 3 | 0.872 | 1 | IMPA2,PIP5K1B,PIK3R2 |
| Glutamine family amino acid metabolic process | 75 | 2.03 | 1 | 0.873 | 1 | PFAS |
| Nuclear import | 232 | 6.28 | 4 | 0.878 | 1 | RERE,CBLB,GLI3,RHOA |
| Response to oxidative stress | 279 | 7.56 | 5 | 0.878 | 1 | MSRA,VRK2,PXDNL,WRN,PRKD1 |
| Skeletal system development | 459 | 12.4 | 9 | 0.879 | 1 | HDAC4,COL11A1,EXT1,FGFR1,ME |
| Carbohydrate catabolic process | 184 | 4.98 | 3 | 0.879 | 1 | HDAC4,GPD2,PRKAG2 |
| Negative regulation of cell migration | 133 | 3.6 | 2 | 0.879 | 1 | SEMA3A,MAGI2 |
| Negative regulation of intracellular transport | 77 | 2.09 | 1 | 0.88 | 1 | MAPT |
| Regulation of JUN kinase activity | 77 | 2.09 | 1 | 0.88 | 1 | TNXB |
| Transcription from RNA polymerase II promoter | 1930 | 52.2 | 45 | 0.88 | 1 | KMT2D,KMT2A,FOXP1,ESR2,ESR[illegible] |
| Regulation of sequence_specific DNA binding transcription factor activity | 372 | 10.1 | 7 | 0.881 | 1 | FOXP1,ESR2,PTCH1,PRKD1,HDA[illegible] |

| | | | | | | |
|---|---|---|---|---|---|---|
| Regulation of response to external stimulus | 461 | 12.5 | 9 | 0.882 | 1 | SEMA3A,PRKD1,F2,CD55,PRKG1,[illegible] |
| Regulation of cell cycle | 886 | 24 | 19 | 0.883 | 1 | RPA3,KMT2E,PSMA5,PTCH1,PKP4 |
| Tube developme nt | 506 | 13.7 | 10 | 0.884 | 1 | SEMA3E,FOXP1,PTCH1,ZFPM2,C[illegible] |
| Negative regulation of secretion | 135 | 3.66 | 2 | 0.884 | 1 | PDE8B,BT N2A2 |
| Intrinsic apoptotic signaling pathway | 135 | 3.66 | 2 | 0.884 | 1 | CUL3,PPP2R5C |
| Cytokine_ mediated signaling pathway | 374 | 10.1 | 7 | 0.884 | 1 | HLA-B,VRK2,CTR9,HLA-DQA1,HLA |
| Enzyme linked receptor protein signaling pathway | 1180 | 31.8 | 26 | 0.885 | 1 | NREP,PDE1C,GIGYF2,PRKD1,RPT |
| Regulation of endothelial cell proliferatio n | 79 | 2.14 | 1 | 0.886 | 1 | PRKD1 |
| Positive regulation of binding | 79 | 2.14 | 1 | 0.886 | 1 | MEF2C |
| Regulation of protein import into nucleus | 137 | 3.71 | 2 | 0.889 | 1 | GLI3,RHO A |
| Sphingolipi d biosyntheti c process | 80 | 2.17 | 1 | 0.89 | 1 | PRKD1 |
| Homeostat ic process | 1510 | 40.8 | 34 | 0.892 | 1 | SLC8A3,RPA3,KMT2E,DPP6,ADAM |
| Peripheral nervous system developme nt | 81 | 2.19 | 1 | 0.893 | 1 | ADAM22 |

| | | | | | | |
|---|---|---|---|---|---|---|
| Cation homeostasis | 514 | 13.9 | 10 | 0.894 | 1 | SLC8A3,F2,HFE,NEDD4L,ATP2A2,[illegible] |
| Regulation of intracellular transport | 290 | 7.85 | 5 | 0.897 | 1 | MAPT,F2,CACNA1C,GLI3,RHOA |
| Microtubule cytoskeleton organization | 337 | 9.13 | 6 | 0.898 | 1 | MAPT,KIZ,CTNNB1,CCDC88C,NEU[illegible] |
| Apoptotic mitochondrial changes | 83 | 2.25 | 1 | 0.898 | 1 | PPP2R2B |
| Positive regulation of catalytic activity | 1070 | 29 | 23 | 0.9 | 1 | PDE1C,ARHGAP15,SYNGAP1,PSM[illegible] |
| Receptor_mediated endocytosis | 142 | 3.85 | 2 | 0.901 | 1 | HFE,MAGI2 |
| Pigment metabolic process | 84 | 2.28 | 1 | 0.901 | 1 | CITED1 |
| DNA damage checkpoint | 143 | 3.87 | 2 | 0.903 | 1 | PSMA5,PPP2R5C |
| Protein_DNA complex assembly | 143 | 3.87 | 2 | 0.903 | 1 | RPA3,TCF4 |
| Regulation of lipid metabolic process | 246 | 6.66 | 4 | 0.904 | 1 | PRKAG2,RBL2,SORBS1,BCL11B |
| Positive regulation of immune system process | 739 | 20 | 15 | 0.905 | 1 | FOXP1,HLA-DMA,DPP4,CBLB,HLA[illegible] |
| Nucleocytoplasmic transport | 388 | 10.5 | 7 | 0.905 | 1 | RERE,RSRC1,THOC7,CBLB,CITED[illegible] |
| Response to nutrient levels | 295 | 7.99 | 5 | 0.905 | 1 | WRN,HFE,TNRC6A,RPTOR,PSPH |

| | | | | | | |
|---|---|---|---|---|---|---|
| RNA export from nucleus | 86 | 2.33 | 1 | 0.906 | 1 | THOC7 |
| Response to endogenous stimulus | 1360 | 36.9 | 30 | 0.907 | 1 | NREP,SLC8A3,PDE1C,KMT2D,GN[illegible] |
| Positive regulation of sequence_specific DNA binding transcription factor activity | 198 | 5.36 | 3 | 0.907 | 1 | ESR2,PRKD1,HDAC4 |
| Regulation of immune effector process | 249 | 6.74 | 4 | 0.909 | 1 | FOXP1,CD55,ABCB9,ELMO1 |
| Tube morphogenesis | 347 | 9.4 | 6 | 0.912 | 1 | SEMA3E,PTCH1,CITED1,MEF2C,C[illegible] |
| Sensory organ development | 485 | 13.1 | 9 | 0.913 | 1 | HLA-DMA,COL11A1,FGFR1,CTNN[illegible] |
| Feeding behavior | 89 | 2.41 | 1 | 0.914 | 1 | GIGYF2 |
| Immune effector process | 576 | 15.6 | 11 | 0.915 | 1 | KMT2E,FOXP1,HLA-DMA,CD55,AB[illegible] |
| Leukocyte chemotaxis | 150 | 4.06 | 2 | 0.917 | 1 | PTPRO,PDE4B |
| Meiosis I | 91 | 2.46 | 1 | 0.918 | 1 | MSH5 |
| DNA integrity checkpoint | 152 | 4.12 | 2 | 0.921 | 1 | PSMA5,PPP2R5C |
| Endothelial cell proliferation | 92 | 2.49 | 1 | 0.921 | 1 | PRKD1 |
| Organelle organization | 2820 | 76.3 | 66 | 0.921 | 1 | KDM5D,SEMA3E,KMT2D,MYO18A,[illegible] |
| M phase of mitotic cell cycle | 447 | 12.1 | 8 | 0.921 | 1 | MDK,ANAPC4,CUL3,MPHOSPH9,[illegible] |

| | | | | | | |
|---|---|---|---|---|---|---|
| Myeloid leukocyte differentiation | 153 | 4.14 | 2 | 0.922 | 1 | KMT2E,CTNNB1 |
| Membrane fusion | 93 | 2.52 | 1 | 0.923 | 1 | DYSF |
| G1/S transition of mitotic cell cycle | 209 | 5.66 | 3 | 0.925 | 1 | RPA3,PSMA5,CUL3 |
| Regulation of I_kappaB kinase/NF_kappaB cascade | 210 | 5.69 | 3 | 0.927 | 1 | PRKD1,CTNNB1,RHOA |
| Gamete generation | 543 | 14.7 | 10 | 0.927 | 1 | KDM5D,KMT2D,SPAG16,ZMIZ1,CD |
| Positive regulation of cell migration | 263 | 7.12 | 4 | 0.929 | 1 | PRKD1,NTRK3,RHOA,HDAC9 |
| Epithelial to mesenchymal transition | 96 | 2.6 | 1 | 0.929 | 1 | CTNNB1 |
| Protein targeting to membrane | 158 | 4.28 | 2 | 0.931 | 1 | PTCH1,ANK3 |
| Innate immune response | 638 | 17.3 | 12 | 0.932 | 1 | HLA-B,PRKD1,CITED1,HLA-DQA1,I |
| Regulation of immune response | 727 | 19.7 | 14 | 0.933 | 1 | HLA-B,HLA-DMA,CBLB,RAET1E,HL |
| System process | 2170 | 58.8 | 49 | 0.934 | 1 | DPP6,GNAI1,ADAM22,SYNGAP1,S |
| Negative regulation of transport | 318 | 8.61 | 5 | 0.935 | 1 | MAPT,PRKD1,PDE8B,NEDD4L,BTN |
| Protein catabolic process | 644 | 17.4 | 12 | 0.937 | 1 | ERLIN1,GRIN2A,PSMA5,ANAPC4,U |
| Regulation of biological quality | 3400 | 92.1 | 80 | 0.938 | 1 | SLC8A3,SEMA3E,DGKI,RPA3,KMT |

| | | | | | | |
|---|---|---|---|---|---|---|
| RNA splicing, via transesterification reactions | 321 | 8.69 | 5 | 0.938 | 1 | RSRC1,PRPF3,POLR2E,SRPK2,NC |
| G_protein coupled receptor signaling pathway, coupled to cyclic nucleotide second messenger | 164 | 4.44 | 2 | 0.939 | 1 | GPR26,GNAI1 |
| Protein import | 272 | 7.37 | 4 | 0.94 | 1 | RERE,CBLB,GLI3,RHOA |
| Monocarboxylic acid transport | 103 | 2.79 | 1 | 0.941 | 1 | PRKAG2 |
| Ribosome biogenesis | 166 | 4.5 | 2 | 0.942 | 1 | EFL1,MRM2 |
| Vasculature development | 652 | 17.7 | 12 | 0.943 | 1 | SEMA3E,PRKD1,ZFPM2,ZMIZ1,CIT |
| Epidermal growth factor receptor signaling pathway | 167 | 4.52 | 2 | 0.943 | 1 | PDE1C,FOXO3 |
| Positive regulation of cell proliferation | 786 | 21.3 | 15 | 0.944 | 1 | KMT2D,FOXP1,PRKD1,DPP4,F2,CU |
| Cellular protein catabolic process | 518 | 14 | 9 | 0.945 | 1 | ERLIN1,PSMA5,ANAPC4,USP4,CU |
| Angiogenesis | 426 | 11.5 | 7 | 0.946 | 1 | SEMA3E,PRKD1,FGFR1,CTNNB1,S |
| RNA 3'_end processing | 106 | 2.87 | 1 | 0.946 | 1 | CNOT1 |
| Negative regulation of signal transduction | 790 | 21.4 | 15 | 0.946 | 1 | SYNGAP1,ESR2,PTCH1,ZNF536,C |

| | | | | | | |
|---|---|---|---|---|---|---|
| Negative regulation of catalytic activity | 568 | 15.4 | 10 | 0.948 | 1 | PTPRN2,PSMA5,ANAPC4,ITIH3,PP |
| Negative regulation of response to stimulus | 967 | 26.2 | 19 | 0.948 | 1 | SYNGAP1,SEMA3A,ESR2,PTCH1,Z |
| Peptide metabolic process | 108 | 2.93 | 1 | 0.949 | 1 | FURIN |
| Regulation of nucleocytoplasmic transport | 172 | 4.66 | 2 | 0.949 | 1 | GLI3,RHOA |
| RNA splicing | 478 | 12.9 | 8 | 0.95 | 1 | RSRC1,THOC7,ZNF638,PRPF3,PO |
| RRNA processing | 109 | 2.95 | 1 | 0.95 | 1 | MRM2 |
| Phagocytosis | 109 | 2.95 | 1 | 0.95 | 1 | ELMO1 |
| Tissue development | 1680 | 45.6 | 36 | 0.951 | 1 | SEMA3E,FOXP1,SEMA3A,ESR2,PT |
| Response to wounding | 1310 | 35.6 | 27 | 0.952 | 1 | NREP,SLC8A3,DGKI,GNAI1,GRIN2 |
| Interphase of mitotic cell cycle | 435 | 11.8 | 7 | 0.953 | 1 | RPA3,GIGYF2,PSMA5,ANAPC4,CU |
| Response to temperature stimulus | 111 | 3.01 | 1 | 0.953 | 1 | MICA |
| ER_nucleus signaling pathway | 111 | 3.01 | 1 | 0.953 | 1 | ATP2A2 |
| Protein modification by small protein conjugation | 713 | 19.3 | 13 | 0.953 | 1 | PSMA5,ANAPC4,CTR9,USP4,CUL3 |
| Male gonad development | 112 | 3.03 | 1 | 0.954 | 1 | ZFPM2 |
| Response to UV | 112 | 3.03 | 1 | 0.954 | 1 | WRN |

| | | | | | | |
|---|---|---|---|---|---|---|
| Response to ionizing radiation | 112 | 3.03 | 1 | 0.954 | 1 | XRCC6 |
| Energy derivation by oxidation of organic compounds | 437 | 11.8 | 7 | 0.954 | 1 | CACNA2D2,KCNG2,CACNA1C,PRI |
| Sensory perception | 532 | 14.4 | 9 | 0.955 | 1 | GRIN2A,BBS9,PKD1L3,COL11A1,F |
| Regulation of immune system process | 1190 | 32.3 | 24 | 0.955 | 1 | HLA-B,KMT2E,FOXP1,HLA-DMA,DI |
| Glucose metabolic process | 290 | 7.85 | 4 | 0.957 | 1 | HDAC4,GPD2,PRKAG2,SORBS1 |
| Positive regulation of NF_kappa B transcription factor activity | 115 | 3.11 | 1 | 0.958 | 1 | PRKD1 |
| DNA repair | 538 | 14.6 | 9 | 0.958 | 1 | RPA3,WRN,MSH5,XRCC6,SLC30A |
| Interphase | 443 | 12 | 7 | 0.959 | 1 | RPA3,GIGYF2,PSMA5,ANAPC4,CU |
| Negative regulation of phosphate metabolic process | 293 | 7.94 | 4 | 0.959 | 1 | PPM1E,GABBR1,PRKAG2,NTRK3 |
| Response to radiation | 345 | 9.34 | 5 | 0.959 | 1 | SYNGAP1,GRIN2A,WRN,CACNA1 |
| DNA replication | 346 | 9.37 | 5 | 0.96 | 1 | RPA3,WRN,FHIT,RFC2,NFIX |
| Viral infectious cycle | 241 | 6.53 | 3 | 0.961 | 1 | FURIN,POLR2E,SRPK2 |
| RRNA metabolic process | 118 | 3.2 | 1 | 0.961 | 1 | MRM2 |

| | | | | | | |
|---|---|---|---|---|---|---|
| Positive regulation of cysteine_type endopeptidase activity involved in apoptotic process | 118 | 3.2 | 1 | 0.961 | 1 | RHOA |
| Protein targeting | 545 | 14.8 | 9 | 0.962 | 1 | AKAP6,RERE,PTCH1,CBLB,ANK3,[illegible] |
| Sensory perception of chemical stimulus | 120 | 3.25 | 1 | 0.963 | 1 | PKD1L3 |
| Regulation of mitotic cell cycle | 351 | 9.51 | 5 | 0.963 | 1 | PSMA5,PTCH1,ANAPC4,MAD1L1,[illegible] |
| JAK_STAT cascade | 121 | 3.28 | 1 | 0.964 | 1 | CTR9 |
| Monovalent inorganic cation transport | 353 | 9.56 | 5 | 0.965 | 1 | DPP6,KCNG2,NEDD4L,ANK3,KCN[illegible] |
| Organic acid transport | 246 | 6.66 | 3 | 0.965 | 1 | PRKAG2,GRIK1,SYT1 |
| Negative regulation of phosphorylation | 246 | 6.66 | 3 | 0.965 | 1 | PPM1E,PRKAG2,NTRK3 |
| Carboxylic acid transport | 246 | 6.66 | 3 | 0.965 | 1 | PRKAG2,GRIK1,SYT1 |
| Positive regulation of secretion | 246 | 6.66 | 3 | 0.965 | 1 | GRIK1,HLA-DRB5,HLA-DRB1 |
| MRNA processing | 551 | 14.9 | 9 | 0.966 | 1 | RSRC1,THOC7,PRPF3,CNOT1,CP[illegible] |
| Regulation of binding | 189 | 5.12 | 2 | 0.966 | 1 | HDAC4,MEF2C |
| Negative regulation of transferase activity | 189 | 5.12 | 2 | 0.966 | 1 | PPM1E,PRKAG2 |

| | | | | | | |
|---|---|---|---|---|---|---|
| Regulation of cell migration | 456 | 12.4 | 7 | 0.966 | 1 | SEMA3A,PRKD1,ZSWIM6,MAGI2,N |
| Generation of precursor metabolites and energy | 603 | 16.3 | 10 | 0.968 | 1 | ENOX1,FADS2,CACNA2D2,KCNG2 |
| Regulation of catalytic activity | 1730 | 46.9 | 36 | 0.968 | 1 | PDE1C,PTPRN2,DGKI,ARHGAP15, |
| Organ morphogenesis | 966 | 26.2 | 18 | 0.968 | 1 | FOXP1,ESR2,PTCH1,ZFPM2,COL1 |
| Fatty acid oxidation | 125 | 3.39 | 1 | 0.968 | 1 | PRKAG2 |
| Triglyceride metabolic process | 126 | 3.41 | 1 | 0.969 | 1 | TNXB |
| Leukocyte migration | 309 | 8.37 | 4 | 0.97 | 1 | F2,PTPRO,PDE4B,PIK3R2 |
| Protein ubiquitination | 658 | 17.8 | 11 | 0.971 | 1 | PSMA5,ANAPC4,CTR9,USP4,CUL3 |
| Positive regulation of MAP kinase activity | 197 | 5.34 | 2 | 0.972 | 1 | FGFR1,NTRK3 |
| Stress_activated protein kinase signaling cascade | 257 | 6.96 | 3 | 0.972 | 1 | MEF2C,TNXB,RHOA |
| Sulfur compound metabolic process | 313 | 8.48 | 4 | 0.972 | 1 | MSRA,EXT1,CTNNB1,ST3GAL3 |
| Membrane lipid biosynthetic process | 130 | 3.52 | 1 | 0.972 | 1 | PRKD1 |
| Activation of MAPK activity | 131 | 3.55 | 1 | 0.973 | 1 | NTRK3 |
| Mitosis | 420 | 11.4 | 6 | 0.973 | 1 | ANAPC4,CUL3,MAD1L1,NEK4,STA |
| Coenzyme biosynthetic process | 133 | 3.6 | 1 | 0.975 | 1 | MTHFD1L |

| | | | | | | |
|---|---|---|---|---|---|---|
| Hormone metabolic process | 203 | 5.5 | 2 | 0.975 | 1 | HFE,FURIN |
| Cellular amino acid metabolic process | 670 | 18.1 | 11 | 0.976 | 1 | MTHFD1L,MSRA,PSMA5,PFAS,CK |
| Cellular amino acid metabolic process | 670 | 18.1 | 11 | 0.976 | 1 | MTHFD1L,MSRA,PSMA5,PFAS,CK |
| Regulation of cell proliferation | 1430 | 38.7 | 28 | 0.977 | 1 | KMT2D,FOXP1,ESR2,PTCH1,PRK |
| Lipid transport | 268 | 7.26 | 3 | 0.978 | 1 | PTCH1,SPNS1,PRKAG2 |
| Response to DNA damage stimulus | 862 | 23.3 | 15 | 0.978 | 1 | RPA3,PSMA5,WRN,MSH5,XRCC6, |
| Ribonucleoprotein complex biogenesis | 269 | 7.29 | 3 | 0.978 | 1 | EFL1,MRM2,SRPK2 |
| Nuclear export | 139 | 3.76 | 1 | 0.978 | 1 | THOC7 |
| Negative regulation of apoptotic process | 679 | 18.4 | 11 | 0.979 | 1 | MYO18A,SYNGAP1,CITED1,FGFR |
| Negative regulation of apoptotic process | 679 | 18.4 | 11 | 0.979 | 1 | MYO18A,SYNGAP1,CITED1,FGFR |
| Regulation of MAP kinase activity | 272 | 7.37 | 3 | 0.98 | 1 | FGFR1,NTRK3,TNXB |
| Proteolysis | 1100 | 29.7 | 20 | 0.98 | 1 | ERLIN1,DPP6,ADAM22,PSMA5,DP |
| Anatomical structure morphogenesis | 2820 | 76.2 | 61 | 0.981 | 1 | NREP,SEMA3E,ARHGAP15,SYNG |
| Defense response to virus | 215 | 5.82 | 2 | 0.982 | 1 | MICA,ELMO1 |

| | | | | | | |
|---|---|---|---|---|---|---|
| Negative regulation of programmed cell death | 691 | 18.7 | 11 | 0.982 | 1 | MYO18A,SYNGAP1,CITED1,FGFR1 |
| Sexual reproduction | 645 | 17.5 | 10 | 0.983 | 1 | KDM5D,KMT2D,SPAG16,ZMIZ1,CD |
| Positive regulation of multicellular organismal process | 649 | 17.6 | 10 | 0.984 | 1 | F2,CACNA2D2,ATP2A2,ARFGEF2,I |
| Regulation of JNK cascade | 150 | 4.06 | 1 | 0.984 | 1 | TNXB |
| G2/M transition of mitotic cell cycle | 150 | 4.06 | 1 | 0.984 | 1 | ANAPC4 |
| RNA processing | 890 | 24.1 | 15 | 0.985 | 1 | RSRC1,TYW1B,MRM2,RBM6,THO( |
| Chromatin assembly or disassembly | 152 | 4.12 | 1 | 0.985 | 1 | TLK1 |
| Defense response to bacterium | 152 | 4.12 | 1 | 0.985 | 1 | MICA |
| Ion transport | 1340 | 36.4 | 25 | 0.986 | 1 | SLC8A3,DPP6,SLC22A23,F2,HFE,K |
| Cell proliferation | 1900 | 51.5 | 38 | 0.986 | 1 | KMT2D,FOXP1,ESR2,PTCH1,PRKD |
| Membrane lipid metabolic process | 233 | 6.31 | 2 | 0.988 | 1 | AGMO,PRKD1 |
| M phase | 671 | 18.2 | 10 | 0.988 | 1 | KIZ,MDK,ANAPC4,CUL3,MPHOSPH |
| Female pregnancy | 162 | 4.39 | 1 | 0.989 | 1 | HFE |
| Cytokine production | 576 | 15.6 | 8 | 0.989 | 1 | HLA-B,ARFGEF2,FURIN,HLA-DRB5 |
| Cellular amino acid catabolic process | 166 | 4.5 | 1 | 0.99 | 1 | IVD |

| | | | | | | |
|---|---|---|---|---|---|---|
| Protein folding | 241 | 6.53 | 2 | 0.99 | 1 | FKBP9,HLA-DMA |
| Cellular modified amino acid metabolic process | 241 | 6.53 | 2 | 0.99 | 1 | MTHFD1L, CKB |
| Mitochondrion organization | 306 | 8.29 | 3 | 0.99 | 1 | GATB,PPP2R2B,IMMP2L |
| Vesicle_mediated transport | 1110 | 30.1 | 19 | 0.991 | 1 | TSNARE1,PRKD1,EXOC4,HFE,CUI |
| Response to hypoxia | 245 | 6.64 | 2 | 0.991 | 1 | DPP4,RHOA |
| Sphingolipid metabolic process | 173 | 4.69 | 1 | 0.992 | 1 | PRKD1 |
| Regulation of molecular function | 2250 | 60.9 | 45 | 0.992 | 1 | PDE1C,PTPRN2,KMT2A,DGKI,ARH |
| Anion transport | 374 | 10.1 | 4 | 0.992 | 1 | SLC13A1,PRKAG2,GRIK1,SYT1 |
| Reproductive process | 1740 | 47.2 | 33 | 0.992 | 1 | KDM5D,NPAS3,KMT2D,HLA-B,SYN |
| Translation | 698 | 18.9 | 10 | 0.993 | 1 | MRPL33,GIGYF2,TNRC6A,FXR1,G |
| Multi_organism process | 1710 | 46.2 | 32 | 0.993 | 1 | NPAS3,HLA-B,PSMA5,VRK2,ZSCAI |
| Cellular macromolecule catabolic process | 849 | 23 | 13 | 0.993 | 1 | ERLIN1,RPA3,PSMA5,ANAPC4,USI |
| Glycerophospholipid biosynthetic process | 255 | 6.91 | 2 | 0.993 | 1 | PIP5K1B,PIK3R2 |
| RNA catabolic process | 256 | 6.93 | 2 | 0.993 | 1 | CNOT1,ETF1 |
| Steroid biosynthetic process | 183 | 4.96 | 1 | 0.994 | 1 | PRKAG2 |
| Meiotic cell cycle | 183 | 4.96 | 1 | 0.994 | 1 | MSH5 |
| Response to virus | 325 | 8.8 | 3 | 0.994 | 1 | MEF2C,MICA,ELMO1 |

| | | | | | | |
|---|---|---|---|---|---|---|
| Activation of protein kinase activity | 260 | 7.04 | 2 | 0.994 | 1 | CHRNA3,NTRK3 |
| Positive regulation of metabolic process | 2690 | 72.9 | 55 | 0.994 | 1 | NPAS3,KMT2D,KMT2A,KMT2E,FO |
| Glycerophospholipid metabolic process | 327 | 8.86 | 3 | 0.994 | 1 | IMPA2,PIP5K1B,PIK3R2 |
| Apoptotic signaling pathway | 261 | 7.07 | 2 | 0.994 | 1 | CUL3,PPP2R5C |
| Cofactor biosynthetic process | 185 | 5.01 | 1 | 0.994 | 1 | MTHFD1L |
| Response to other organism | 716 | 19.4 | 10 | 0.994 | 1 | HLA-B,CTR9,CITED1,MEF2C,SRR, |
| Amine metabolic process | 268 | 7.26 | 2 | 0.995 | 1 | MSRA,PSMA5 |
| Fatty acid metabolic process | 397 | 10.8 | 4 | 0.995 | 1 | AGMO,FADS2,PRKAG2,TNXB |
| Regulation of cytokine production | 513 | 13.9 | 6 | 0.995 | 1 | HLA-B,ARFGEF2,FURIN,HLA-DRB |
| Inflammatory response | 569 | 15.4 | 7 | 0.995 | 1 | PRKD1,F2,HDAC4,CD55,HLA-DRB |
| Response to abiotic stimulus | 876 | 23.7 | 13 | 0.995 | 1 | SYNGAP1,GRIN2A,WRN,DPP4,PK |
| Positive regulation of defense response | 273 | 7.39 | 2 | 0.995 | 1 | MEF2C,MICA |
| Cellular catabolic process | 2140 | 58.1 | 41 | 0.996 | 1 | PTPRN2,ERLIN1,DGKI,RPA3,ARH |
| Positive regulation of cellular metabolic process | 2530 | 68.4 | 50 | 0.996 | 1 | NPAS3,KMT2D,KMT2A,KMT2E,FO |
| Protein localization | 1850 | 50 | 34 | 0.996 | 1 | AKAP6,RERE,SYNGAP1,HLA-DMA |

| | | | | | | |
|---|---|---|---|---|---|---|
| Positive regulation of protein metabolic process | 1080 | 29.3 | 17 | 0.996 | 1 | PSMA5,PRKD1,F2,ANAPC4,CTR9,F |
| Establishment of protein localization | 1460 | 39.4 | 25 | 0.996 | 1 | AKAP6,RERE,HLA-DMA,UTY,TSNA |
| Reproduction | 1860 | 50.2 | 34 | 0.996 | 1 | KDM5D,NPAS3,KMT2D,HLA-B,SYN |
| Intracellular protein transport | 793 | 21.5 | 11 | 0.996 | 1 | AKAP6,RERE,TSNARE1,PTCH1,CE |
| JNK cascade | 204 | 5.53 | 1 | 0.996 | 1 | TNXB |
| Phospholipid biosynthetic process | 285 | 7.72 | 2 | 0.997 | 1 | PIP5K1B,PIK3R2 |
| Intracellular transport | 1510 | 40.9 | 26 | 0.997 | 1 | SLC8A3,AKAP6,RERE,RSRC1,MAF |
| Response to biotic stimulus | 749 | 20.3 | 10 | 0.997 | 1 | HLA-B,CTR9,CITED1,MEF2C,SRR,I |
| Cellular localization | 2650 | 71.9 | 52 | 0.997 | 1 | SLC8A3,AKAP6,RERE,SYNGAP1,F |
| Positive regulation of protein modification process | 867 | 23.5 | 12 | 0.998 | 1 | PSMA5,PRKD1,F2,ANAPC4,CTR9,F |
| MRNA metabolic process | 817 | 22.1 | 11 | 0.998 | 1 | PSMA5,RSRC1,THOC7,PRPF3,CN( |
| Mitotic cell cycle | 968 | 26.2 | 14 | 0.998 | 1 | RPA3,GIGYF2,PSMA5,PTCH1,MDK |
| Positive regulation of cellular protein metabolic process | 968 | 26.2 | 14 | 0.998 | 1 | PSMA5,PRKD1,F2,ANAPC4,CTR9,F |
| Positive regulation of MAPK cascade | 368 | 9.97 | 3 | 0.998 | 1 | FGFR1,CTNNB1,NTRK3 |
| Regulation of MAPK cascade | 559 | 15.1 | 6 | 0.998 | 1 | SYNGAP1,VRK2,FGFR1,CTNNB1,N |

| | | | | | | |
|---|---|---|---|---|---|---|
| G_protein coupled receptor signaling pathway | 779 | 21.1 | 10 | 0.998 | 1 | GPR26,DGKI,GNAI1,GPR141,F2,PK |
| Cell cycle process | 1420 | 38.3 | 23 | 0.998 | 1 | RPA3,KMT2E,GIGYF2,PSMA5,KIZ,I |
| Peptidyl_tyrosine phosphorylation | 228 | 6.18 | 1 | 0.998 | 1 | FGFR1 |
| Peptidyl_tyrosine modification | 230 | 6.23 | 1 | 0.998 | 1 | FGFR1 |
| Monocarboxylic acid metabolic process | 567 | 15.4 | 6 | 0.998 | 1 | MTHFD1L,AGMO,FADS2,PRKAG2, |
| Positive regulation of transferase activity | 510 | 13.8 | 5 | 0.998 | 1 | RPTOR,CHRNA3,FGFR1,PRKAG2, |
| Establishment of localization in cell | 2320 | 62.8 | 43 | 0.998 | 1 | SLC8A3,AKAP6,RERE,FEZ1,RSRC |
| Intracellular signal transduction | 2450 | 66.3 | 46 | 0.998 | 1 | DGKI,ARHGAP15,SYNGAP1,PSMA |
| Alcohol metabolic process | 388 | 10.5 | 3 | 0.999 | 1 | GPD2,PRKAG2,IMPA2 |
| Cellular respiration | 236 | 6.39 | 1 | 0.999 | 1 | NDUFA2 |
| Phospholipid metabolic process | 463 | 12.5 | 4 | 0.999 | 1 | PRKD1,IMPA2,PIP5K1B,PIK3R2 |
| Protein transport | 1400 | 37.9 | 22 | 0.999 | 1 | AKAP6,RERE,HLA-DMA,TSNARE1, |
| Cell cycle | 1860 | 50.4 | 32 | 0.999 | 1 | RPA3,KMT2E,GNAI1,GIGYF2,PSMA |
| Macromolecule catabolic process | 1070 | 29 | 15 | 0.999 | 1 | ERLIN1,RPA3,GRIN2A,PSMA5,ANA |
| Macromolecule catabolic process | 1070 | 29 | 15 | 0.999 | 1 | ERLIN1,RPA3,GRIN2A,PSMA5,ANA |
| Cell cycle phase | 1070 | 29.1 | 15 | 0.999 | 1 | RPA3,GIGYF2,PSMA5,KIZ,MDK,AN |

| | | | | | | |
|---|---|---|---|---|---|---|
| Positive regulation of response to stimulus | 1550 | 42.1 | 25 | 0.999 | 1 | KMT2D,KMT2A,DGKI,HLA-DMA,PR |
| Response to organic substance | 2500 | 67.7 | 46 | 0.999 | 1 | NREP,SLC8A3,PDE1C,KMT2D,HLA |
| MAPK cascade | 665 | 18 | 7 | 0.999 | 1 | SYNGAP1,VRK2,FGFR1,MEF2C,CT |
| Cellular response to stress | 1620 | 44 | 26 | 0.999 | 1 | NREP,RPA3,PSMA5,VRK2,PXDNL, |
| Regulation of apoptotic process | 1540 | 41.6 | 24 | 0.999 | 1 | MYO18A,SYNGAP1,GRIN2A,PSMA |
| Coenzyme metabolic process | 266 | 7.2 | 1 | 0.999 | 1 | MTHFD1L |
| Positive regulation of cytokine production | 268 | 7.26 | 1 | 0.999 | 1 | ARFGEF2 |
| Positive regulation of protein phosphorylation | 681 | 18.4 | 7 | 0.999 | 1 | PRKD1,F2,RPTOR,CHRNA3,FGFR |
| Regulation of protein metabolic process | 1820 | 49.4 | 30 | 0.999 | 1 | GRIN2A,GIGYF2,PSMA5,PRKD1,F2 |
| Regulation of kinase activity | 743 | 20.1 | 8 | 1 | 1 | RPTOR,PPM1E,CHRNA3,FGFR1,P |
| Regulation of programmed cell death | 1550 | 42.1 | 24 | 1 | 1 | MYO18A,SYNGAP1,GRIN2A,PSMA |
| Regulation of cellular protein metabolic process | 1560 | 42.2 | 24 | 1 | 1 | GIGYF2,PSMA5,PRKD1,F2,ANAPC |
| Regulation of transcription, DNA_dependent | 3770 | 102 | 75 | 1 | 1 | NPAS3,KMT2D,KMT2A,KMT2E,FO |

| | | | | | | |
|---|---|---|---|---|---|---|
| Regulation of transcription, DNA_dependent | 3770 | 102 | 75 | 1 | 1 | NPAS3,KMT2D,KMT2A,KMT2E,FO[illegible] |
| Regulation of transcription, DNA_dependent | 3770 | 102 | 75 | 1 | 1 | NPAS3,KMT2D,KMT2A,KMT2E,FO[illegible] |
| Positive regulation of phosphorylation | 697 | 18.9 | 7 | 1 | 1 | PRKD1,F2,RPTOR,CHRNA3,FGFR[illegible] |
| Regulation of protein kinase activity | 698 | 18.9 | 7 | 1 | 1 | RPTOR,PPM1E,CHRNA3,FGFR1,P[illegible] |
| Immune response | 1430 | 38.7 | 21 | 1 | 1 | HLA-B,HLA-DMA,PRKD1,HFE,CBL[illegible] |
| Intracellular protein kinase cascade | 1140 | 30.8 | 15 | 1 | 1 | SYNGAP1,VRK2,PRKD1,CTR9,FG[illegible] |
| Regulation of defense response | 519 | 14.1 | 4 | 1 | 1 | MEF2C,CD55,MICA,ELMO1 |
| Carbohydrate metabolic process | 1040 | 28.2 | 13 | 1 | 1 | GALNT12,HDAC4,GPD2,EXT1,PRK[illegible] |
| Regulation of transferase activity | 768 | 20.8 | 8 | 1 | 1 | RPTOR,PPM1E,CHRNA3,FGFR1,P[illegible] |
| Macromolecule localization | 2240 | 60.7 | 38 | 1 | 1 | AKAP6,RERE,SYNGAP1,HLA-DMA[illegible] |
| Positive regulation of signal transduction | 998 | 27 | 12 | 1 | 1 | KMT2D,DGKI,PRKD1,F2,RPTOR,C[illegible] |
| Regulation of RNA metabolic process | 3900 | 106 | 77 | 1 | 1 | NPAS3,KMT2D,KMT2A,KMT2E,FO[illegible] |
| Catabolic process | 2560 | 69.4 | 45 | 1 | 1 | PTPRN2,ERLIN1,DGKI,RPA3,ARH[illegible] |

| | | | | | | |
|---|---|---|---|---|---|---|
| Cell surface receptor signaling pathway | 3340 | 90.4 | 63 | 1 | 1 | NREP,MAGI1,GPR26,PDE1C,SEMA |
| Positive regulation of phosphate metabolic process | 789 | 21.4 | 8 | 1 | 1 | PRKD1,F2,RPTOR,CHRNA3,FGFR |
| Regulation of nucleobase_containing compound metabolic process | 4540 | 123 | 92 | 1 | 1 | PTPRN2,NPAS3,KMT2D,KMT2A,D |
| Immune system process | 2720 | 73.6 | 48 | 1 | 1 | KMT2A,HLA-B,KMT2E,FOXP1,HLA- |
| DNA metabolic process | 1240 | 33.7 | 16 | 1 | 1 | MYO18A,RPA3,KMT2E,FOXP1,WR |
| Regulation of protein modification process | 1250 | 33.8 | 16 | 1 | 1 | PSMA5,PRKD1,F2,ANAPC4,CTR9, |
| Cofactor metabolic process | 331 | 8.97 | 1 | 1 | 1 | MTHFD1L |
| Negative regulation of cellular process | 4110 | 111 | 80 | 1 | 1 | SEMA3E,KMT2D,MYO18A,RPA3,K |
| Regulation of gene expression | 4480 | 121 | 89 | 1 | 1 | NPAS3,KMT2D,KMT2A,KMT2E,FO |
| Organ development | 3290 | 89.2 | 60 | 1 | 1 | SLC8A3,KMT2A,HLA-B,KMT2E,FO |
| Lipid biosynthetic process | 776 | 21 | 7 | 1 | 1 | AGMO,FADS2,PRKD1,PRKAG2,PI |
| Organic acid metabolic process | 1430 | 38.7 | 19 | 1 | 1 | MTHFD1L,MSRA,PSMA5,AGMO,F |
| Regulation of signal transduction | 2440 | 66 | 40 | 1 | 1 | NREP,KMT2D,DGKI,ARHGAP15,S |

| | | | | | | |
|---|---|---|---|---|---|---|
| Steroid metabolic process | 361 | 9.78 | 1 | 1 | 1 | PRKAG2 |
| Regulation of protein phosphorylation | 987 | 26.7 | 10 | 1 | 1 | PRKD1,F2,RPTOR,PPM1E,CHRNA |
| Carboxylic acid metabolic process | 1270 | 34.5 | 15 | 1 | 1 | MTHFD1L,MSRA,PSMA5,AGMO,FA |
| Response to chemical stimulus | 3830 | 104 | 71 | 1 | 1 | NREP,SLC8A3,PDE1C,SEMA3E,KM |
| Apoptotic process | 2130 | 57.7 | 32 | 1 | 1 | MYO18A,KMT2A,SYNGAP1,SEMA3 |
| Apoptotic process | 2130 | 57.7 | 32 | 1 | 1 | MYO18A,KMT2A,SYNGAP1,SEMA3 |
| Regulation of phosphorylation | 1070 | 28.9 | 11 | 1 | 1 | PRKD1,F2,RPTOR,PPM1E,CHRNA |
| Regulation of response to stimulus | 3360 | 91 | 59 | 1 | 1 | NREP,KMT2D,KMT2A,HLA-B,DGKI |
| Programmed cell death | 2160 | 58.4 | 32 | 1 | 1 | MYO18A,KMT2A,SYNGAP1,SEMA3 |
| System development | 4950 | 134 | 97 | 1 | 1 | NRGN,NREP,SLC8A3,SEMA3E,KM |
| Macromolecule modification | 3430 | 92.9 | 60 | 1 | 1 | FKBP9,KDM5D,PTPRN2,GALNT12, |
| Protein modification process | 3290 | 89.2 | 56 | 1 | 1 | FKBP9,KDM5D,PTPRN2,GALNT12, |
| Phosphorylation | 1680 | 45.4 | 21 | 1 | 1 | VRK2,RSRC1,PRKD1,F2,RPTOR,P |
| Negative regulation of biological process | 4590 | 124 | 86 | 1 | 1 | SEMA3E,KMT2D,MYO18A,RPA3,KM |
| Protein phosphorylation | 1480 | 40.2 | 17 | 1 | 1 | VRK2,RSRC1,PRKD1,F2,RPTOR,P |
| Defense response | 1510 | 40.9 | 17 | 1 | 1 | HLA-B,PRKD1,MDK,F2,HDAC4,CIT |

| | | | | | | |
|---|---|---|---|---|---|---|
| Positive regulation of cellular process | 4780 | 129 | 84 | 1 | 1 | NPAS3,KMT2D,KMT2A,DGKI,KMT2 |
| Cellular lipid metabolic process | 1200 | 32.6 | 9 | 1 | 1 | AGMO,FADS2,PRKD1,GPD2,PRKA |
| Transcription, DNA_dependent | 4830 | 131 | 85 | 1 | 1 | NPAS3,KMT2D,KMT2A,KMT2E,REI |
| Transcription, DNA_dependent | 4830 | 131 | 85 | 1 | 1 | NPAS3,KMT2D,KMT2A,KMT2E,REI |
| Multicellular organismal development | 5720 | 155 | 106 | 1 | 1 | NRGN,NREP,SLC8A3,SEMA3E,KM |
| Anatomical structure development | 5830 | 158 | 108 | 1 | 1 | NRGN,NREP,SLC8A3,SEMA3E,KM |
| RNA biosynthetic process | 4930 | 134 | 85 | 1 | 1 | NPAS3,KMT2D,KMT2A,KMT2E,REI |
| Regulation of cellular metabolic process | 6120 | 166 | 114 | 1 | 1 | PTPRN2,NPAS3,KMT2D,KMT2A,D( |
| Lipid metabolic process | 1730 | 46.8 | 15 | 1 | 1 | AGMO,FADS2,PRKD1,PLCL1,GPD2 |
| Positive regulation of biological process | 5500 | 149 | 95 | 1 | 1 | NPAS3,KMT2D,KMT2A,DGKI,KMT2 |
| Cellular nitrogen compound biosynthetic process | 5610 | 152 | 95 | 1 | 1 | MTHFD1L,NPAS3,KMT2D,KMT2A,K |
| Establishment of localization | 4910 | 133 | 78 | 1 | 1 | SLC8A3,KMT2A,AKAP6,DPP6,RER |
| Transport | 4830 | 131 | 76 | 1 | 1 | SLC8A3,KMT2A,AKAP6,DPP6,RER |

| | | | | | | |
|---|---|---|---|---|---|---|
| Cellular protein metabolic process | 4540 | 123 | 69 | 1 | 1 | FKBP9,KDM5D,PTPRN2,GALNT12, |
| Response to stress | 4150 | 112 | 60 | 1 | 1 | NREP,SLC8A3,HLA-B,MSRA,DGKI, |
| Signal transductio n | 6310 | 171 | 110 | 1 | 1 | NRGN,NREP,MAGI1,GPR26,PDE1( |
| Regulation of metabolic process | 6920 | 187 | 125 | 1 | 1 | PDE1C,PTPRN2,NPAS3,KMT2D,KI |
| RNA metabolic process | 6010 | 163 | 101 | 1 | 1 | NPAS3,KMT2D,KMT2A,KMT2E,REI |
| Protein metabolic process | 5570 | 151 | 85 | 1 | 1 | FKBP9,KDM5D,PTPRN2,GALNT12, |
| Macromole cule biosyntheti c process | 6540 | 177 | 106 | 1 | 1 | NPAS3,GALNT12,KMT2D,KMT2A,F |
| Macromole cule biosyntheti c process | 6540 | 177 | 106 | 1 | 1 | NPAS3,GALNT12,KMT2D,KMT2A,F |
| Nucleobas e_containi ng compound metabolic process | 8180 | 221 | 136 | 1 | 1 | PTPRN2,NPAS3,KMT2D,MYO18A,I |
| Nucleobas e_containi ng compound metabolic process | 8180 | 221 | 136 | 1 | 1 | PTPRN2,NPAS3,KMT2D,MYO18A,I |
| Nitrogen compound metabolic process | 9380 | 254 | 148 | 1 | 1 | MTHFD1L,PTPRN2,NPAS3,KMT2D |
| Biosyntheti c process | 8090 | 219 | 124 | 1 | 1 | MTHFD1L,NPAS3,GALNT12,KMT2I |
| Macromole cule metabolic process | 12000 | 326 | 171 | 1 | 1 | FKBP9,KDM5D,PTPRN2,NPAS3,G/ |
| Cellular biosyntheti c process | 7800 | 211 | 121 | 1 | 1 | MTHFD1L,NPAS3,GALNT12,KMT2I |

| | | | | | | |
|---|---|---|---|---|---|---|
| Cellular macromolecule metabolic process | 10800 | 293 | 156 | 1 | 1 | FKBP9,KDM5D,PTPRN2,NPAS3,GA |
| Heterocycle metabolic process | 8430 | 228 | 139 | 1 | 1 | MTHFD1L,PTPRN2,NPAS3,KMT2D |
| Cellular response to stimulus | 8010 | 217 | 127 | 1 | 1 | NRGN,NREP,SLC8A3,MAGI1,GPR2 |
| Cellular aromatic compound metabolic process | 8460 | 229 | 139 | 1 | 1 | MTHFD1L,PTPRN2,NPAS3,KMT2D |

APT,PRKD1,F2,ZNF536,TCF4,CNTN4,CHRNA3,FGFR1,XRCC6,AMIGO1,MEF2C,ZSWIM6,DBN1,SOX5,CTN

MA3A,MAPT,F2,CITED1,CHRNA3,AMIGO1,RHOJ,CTNNB1,NGEF,TWF2,BCL11A,PTPRO,SEMA3F,NTRK3

IZ1,FGFR1,MEF2C,MAGI2,PALB2,TWF2,ESRRB,BCL11A,FOXO3,SEMA3F,NTRK3,WNT3,GLI3

SEMA3A,FEZ1,MAPT,CUL3,CNTN4,SEMA6D,EXT1,KIF5C,CITED1,CHRNA3,CACNA1C,FGFR1,AMIGO1,ME

SEMA3A,FEZ1,MAPT,CNTN4,SEMA6D,EXT1,KIF5C,CACNA1C,FGFR1,AMIGO1,CACNB2,ANK3,CACNA1I,T

SEMA3A,RTN1,GIGYF2,FEZ1,ESR2,MAPT,PRKD1,F2,ZNF536,TCF4,CNTNAP2,CNTN4,SEMA6D,EXT1,KIF

SEMA3A,FEZ1,MAPT,PRKD1,CNTNAP2,CNTN4,SEMA6D,EXT1,KIF5C,CHRNA3,CACNA1C,FGFR1,AMIGO

NTN4,SEMA6D,EXT1,KIF5C,CACNA1C,FGFR1,CACNB2,ANK3,CACNA1I,PTPRO,SEMA3F,TENM2,WNT3,B

l,FOXP1,SEMA3A,RTN1,GRIN2A,GIGYF2,FEZ1,ESR2,MAPT,PRKD1,F2,ZNF536,TCF4,CNTNAP2,CNTN4,S

GIGYF2,ESR2,PTCH1,MDK,F2,NPAS1,CNTNAP2,CNTN4,EXT1,CITED1,CKB,LDB1,FGFR1,GRIK1,CADM2,

SEMA3A,RTN1,GIGYF2,FEZ1,MAPT,PRKD1,ZNF536,TCF4,CNTNAP2,CNTN4,SEMA6D,EXT1,KIF5C,CHRN

SEMA3A,FEZ1,MAPT,PRKD1,CNTNAP2,CNTN4,SEMA6D,EXT1,KIF5C,CHRNA3,CACNA1C,FGFR1,AMIGO

XP1,SEMA3A,HLA-DMA,FEZ1,MAPT,PTCH1,PRKD1,F2,ZNF536,TCF4,CTR9,HDAC4,ZFPM2,CNTN4,CITEI

K,CNTNAP2,CNTN4,EXT1,CITED1,CKB,LDB1,FGFR1,CADM2,CTNNB1,PRKG1,SRR,CTTNBP2,PPP2R2B,

YNGAP1,FOXP1,SEMA3A,FEZ1,ESR2,MAPT,PRKD1,F2,ZNF536,TCF4,CACNA2D2,CTR9,CUL3,CNTNAP2,

PT,PRKD1,TCF4,FGFR1,XRCC6,AMIGO1,MEF2C,SOX5,CTNNB1,TWF2,BCL11A,TSHZ3,FOXO3,RGS6,NT

XP1,SEMA3A,MAPT,PRKD1,F2,CITED1,CHRNA3,FGFR1,AMIGO1,RHOJ,MAGI2,CTNNB1,NGEF,TWF2,BC

A,GRIN2A,GIGYF2,MAPT,PRKD1,MDK,PLCL1,CHRNA5,CNTNAP2,CHRNA3,CACNA1C,MEF2C,GRIK1,PTF

536,CTR9,HDAC4,ZFPM2,CNTN4,CITED1,LDB1,CTNNB1,MED27,ESRRB,BCL11A,SEMA3F,NTRK3,WNT3

RIN2A,TSNARE1,CHRNA5,CACNA2D2,KCNG2,CNTNAP2,PCLO,GABBR1,BSN,CNTN4,CHRNA3,CACNA1

AM22,SYNGAP1,FOXP1,SEMA3A,RTN1,GRIN2A,GIGYF2,FEZ1,ESR2,MAPT,PTCH1,PRKD1,MDK,F2,ZNF5

MYO9B,CUL3,ARFGEF2,GMIP,RHOJ,RASGRP4,NGEF,GBF1,NTRK3,PREX1,DOCK4,RHOA,PIK3R2

2E,RERE,PRKD1,CTR9,SGF29,HDAC4,L3MBTL4,LDB1,ARID1B,KDM4A,JMJD1C,RBL2,KDM3B,HDAC9,TL

SYNGAP1,GRIN2A,MAPT,PKP4,MYO9B,HDAC4,NEDD4L,CBLB,PRKAG2,GMIP,NGEF,FURIN,PRKG1,RGS6

F4,CTR9,HDAC4,ZFPM2,ZMIZ1,LDB1,XRCC6,SLC30A9,MEF2C,SOX5,CTNNB1,ESRRB,MAML3,BCL11A,FC

SR2,MAPT,WRN,PTCH1,F2,CACNA2D2,RPTOR,ZMIZ1,CACNA1C,FGFR1,MEF2C,MAGI2,RASGRP4,PALE

H1,F2,ZNF536,CTR9,HDAC4,ZFPM2,CNTN4,CITED1,LDB1,CTNNB1,NGEF,MED27,ESRRB,BCL11A,SEMA3

E1,CHRNA5,CACNA2D2,KCNG2,PCLO,GABBR1,BSN,CNTN4,CHRNA3,CACNA1C,MEF2C,GRIK1,DBN1,CT

N,PRKD1,PKP4,MYO9B,FGFR1,MEF2C,GMIP,MAGI2,RASGRP4,RGS6,NTRK3,PREX1,DOCK4,RHOA

CTNNB1,MICA,BCL11A,HLA-DRB5,HLA-DRB1,HLA-DQB1,BTN2A2,BCL11B,GLI3,PIK3R2,BTN3A1

)AC4,CBLB,HLA-DQA1,MEF2C,CTNNB1,MICA,BCL11A,HLA-DRB5,HLA-DRB1,HLA-DQB1,BTN2A2,BCL11B

SYNGAP1,WRN,EFL1,PKP4,PDE8B,MYO9B,GABBR1,PFAS,ATP2A2,GUCY1A2,PRKAG2,GMIP,RHOJ,NGE

PM1E,CARMIL1,RHOJ,DBN1,PRKG1,TWF2,NTRK3,TNXB,ELMO1,PREX1,SORBS1,RHOA,TRIOBP

O9B,CUL3,PPM1E,CARMIL1,RHOJ,DBN1,PRKG1,TWF2,NTRK3,TNXB,ELMO1,PREX1,SORBS1,RHOA,TRI

.PT,PRKD1,TCF4,ZFPM2,FGFR1,XRCC6,AMIGO1,MEF2C,SOX5,CTNNB1,TWF2,BCL11A,TSHZ3,FOXO3,R

SYNGAP1,WRN,EFL1,PKP4,PDE8B,MYO9B,GABBR1,PFAS,ATP2A2,GUCY1A2,PRKAG2,GMIP,RHOJ,NGE

NEDD4L,FGFR1,AMIGO1,MAGI2,CTNNB1,TWF2,BCL11A,TENM2,NTRK3,WNT3,SORBS1,RHOA,TRIOBP

,MYO9B,CUL3,DOCK8,ARFGEF2,GMIP,RHOJ,RASGRP4,NGEF,GBF1,NKIRAS1,ELMO1,PREX1,DOCK4,RH

SYNGAP1,WRN,PRKD1,PKP4,ITIH3,MYO9B,FGFR1,MEF2C,GMIP,MAGI2,RASGRP4,NGEF,FURIN,PRKG1

SYNGAP1,WRN,EFL1,PKP4,PDE8B,MYO9B,GABBR1,PFAS,ATP2A2,NT5C2,GUCY1A2,PRKAG2,GMIP,RH

PP4,HDAC4,CBLB,HLA-DQA1,MEF2C,CTNNB1,MICA,BCL11A,HLA-DRB5,HLA-DRB1,HLA-DQB1,PREX1,B

IGAP15,SYNGAP1,FOXP1,SEMA3A,HLA-DMA,FEZ1,MAPT,PTCH1,PRKD1,F2,ZNF536,TCF4,CTR9,HDAC4

I,PTCH1,F2,CACNA2D2,CACNA1C,FGFR1,MEF2C,TWF2,BCL11A,SEMA3F,NTRK3,WNT3,RHOA

KP1,ESRRG,PRKD1,MDK,TCF4,CTR9,RPTOR,HDAC4,ZFPM2,ZNF398,ZMIZ1,CITED1,LDB1,ARID1B,XRCC

KP1,ESRRG,PRKD1,MDK,TCF4,CTR9,RPTOR,HDAC4,ZFPM2,ZNF398,ZMIZ1,CITED1,LDB1,ARID1B,XRCC

2E,RERE,PRKD1,CTR9,SGF29,HDAC4,L3MBTL4,LDB1,ARID1B,KDM4A,JMJD1C,RBL2,KDM3B,HDAC9,TL

MA,BANK1,DPP4,F2,HDAC4,CBLB,CALU,HLA-DQA1,MEF2C,CTNNB1,PRKG1,MICA,BCL11A,HLA-DRB5,H

P1,ESRRG,PRKD1,MDK,TCF4,CTR9,RPTOR,HDAC4,ZFPM2,ZNF398,ZMIZ1,CITED1,LDB1,ARID1B,XRC

3,KMT2E,RERE,WRN,PRKD1,CTR9,SGF29,HDAC4,L3MBTL4,LDB1,ARID1B,MSH5,XRCC6,CTNNB1,KDM4 [illegible]D4L,ATP2A2,CACNA1C,SLC30A9,CTNNB1,CACNB2,ANK3,CACNA1I,CNNM2,KCNN2,SLC39A8,ATP2B2,[illegible]

-DMA,TET2,CTR9,HDAC4,LDB1,MEF2C,RASGRP4,CTNNB1,BCL11A,FOXO3,WNT3,BCL11B,GLI3,HDAC9

-DMA,TET2,CTR9,HDAC4,CBLB,LDB1,XRCC6,MEF2C,RASGRP4,CTNNB1,BCL11A,FOXO3,WNT3,BCL11E

SYNGAP1,WRN,EFL1,PKP4,PDE8B,FHIT,MYO9B,GABBR1,PFAS,ATP2A2,NT5C2,GUCY1A2,PRKAG2,GMII

-DMA,TET2,CTR9,HDAC4,LDB1,MEF2C,RASGRP4,CTNNB1,BCL11A,FOXO3,WNT3,BCL11B,GLI3,HDAC9

CK8,ZFPM2,CALU,ATP2A2,GUCY1A2,CARMIL1,PRKG1,PTPRO,JMJD1C,ATP2B2,NEURL1,RHOA,PIK3R2

RIN2A,FEZ1,WRN,PRKD1,F2,HFE,TNRC6A,RPTOR,CNTN4,COL11A1,SEMA6D,EXT1,KIF5C,CACNA1C,F

TCF4,CTR9,HDAC4,ZFPM2,NEDD4L,CITED1,LDB1,FGFR1,XRCC6,MEF2C,SOX5,CTNNB1,TSHZ3,FOXO3

TCF4,CTR9,HDAC4,ZFPM2,NEDD4L,CITED1,LDB1,FGFR1,XRCC6,MEF2C,SOX5,CTNNB1,TSHZ3,FOXO3

RIN2A,GIGYF2,TSNARE1,MDK,BBS9,CHRNA5,CACNA2D2,KCNG2,CNTNAP2,PKD1L3,PCLO,GABBR1,BS

1,THOC7,NEDD4L,SND1,ZSCAN12,XRCC6,ZSCAN31,ZKSCAN3,FURIN,MICA,KDM4A,POLR2E,SRPK2,NT

XP1,ESRRG,PRKD1,MDK,TCF4,CTR9,RPTOR,HDAC4,ZFPM2,ZNF398,ZMIZ1,CITED1,LDB1,ARID1B,GUCY

1,CTR9,CUL3,ZFPM2,COL11A1,EXT1,ZMIZ1,CITED1,LDB1,FGFR1,MEF2C,PALB2,SOX5,CTNNB1,ESRRB,

E,KCNG2,CHRNA3,MAGI2,SNAP91,CTNNB1,SYT1,KCNB1,SRR,TWF2,CADPS,BCL11A,HLA-DRB5,HLA-D

Z,CUL3,PCLO,PPM1E,CARMIL1,RHOJ,DBN1,CTNNB1,PRKG1,TWF2,CCDC88C,NTRK3,TNXB,SYNE1,EL

TCF4,CTR9,HDAC4,ZFPM2,NEDD4L,CITED1,LDB1,FGFR1,XRCC6,MEF2C,SOX5,CTNNB1,TSHZ3,FOXO3

EMA3A,FEZ1,ESR2,MAPT,PRKD1,DPP4,F2,ANAPC4,CTR9,PPM1E,NEDD4L,CITED1,CHRNA3,FGFR1,AMI

IGAP1,FOXP1,SEMA3A,HLA-DMA,GRIN2A,FEZ1,MAPT,PTCH1,PRKD1,F2,ZNF536,TCF4,CACNA2D2,KCN

MX2,F2,HFE,NEDD4L,ATP2A2,CKB,CHRNA3,CACNA1C,AMIGO1,GRIK1,CD55,CTNNB1,ANK3,HLA-DRB5,H

)1L3,NEDD4L,ATP2A2,CHRNA3,CACNA1C,PRKAG2,SLC30A9,CTNNB1,CACNB2,ANK3,CACNA1I,CNNM2,

)1L3,NEDD4L,ATP2A2,CHRNA3,CACNA1C,PRKAG2,SLC30A9,CTNNB1,CACNB2,ANK3,CACNA1I,CNNM2,

2,HFE,NEDD4L,ATP2A2,CKB,CHRNA3,CACNA1C,AMIGO1,GRIK1,CD55,CTNNB1,ANK3,CNNM2,CLCN3,AT

HRNA3,FGFR1,PRKAG2,MEF2C,RASGRP4,CTNNB1,NGEF,FURIN,FOXO3,NTRK3,PREX1,SORBS1,FOXO6

SD4,RSU1,MAPT,EFL1,PKP4,SPAG16,TCF4,HFE,KCNG2,CUL3,PPM1E,NEDD4L,CHRNA3,LDB1,MSH5,CA

SNAI1,SYNGAP1,WRN,EFL1,PKP4,PDE8B,MYO9B,NT5C2,CNOT1,ETF1,GMIP,RHOJ,NGEF,PRKG1,DPYD,

)1,FGFR1,PRKAG2,MEF2C,CD55,CTNNB1,PSPH,FOXO3,NTRK3,SORBS1,FOXO6,RHOA,PIK3R2,HDAC9

,PTCH1,TCF4,CTR9,TNRC6A,HDAC4,GABBR1,ZFPM2,NEDD4L,FXR1,CITED1,LDB1,FGFR1,XRCC6,MEF2

SNARE1,PKP4,PDE8B,CHRNA5,CACNA2D2,KCNG2,PCLO,GABBR1,BSN,CNTN4,CHRNA3,CACNA1C,FGF

SEMA3A,FEZ1,ESR2,MAPT,PTCH1,PRKD1,SPAG16,CTR9,CUL3,ZFPM2,CNTN4,COL11A1,SEMA6D,EXT1,

,PTCH1,TCF4,CTR9,TNRC6A,HDAC4,GABBR1,ZFPM2,NEDD4L,FXR1,CITED1,LDB1,FGFR1,XRCC6,MEF2

TCF4,CTR9,HDAC4,GABBR1,ZFPM2,NEDD4L,CITED1,LDB1,FGFR1,XRCC6,MEF2C,SOX5,CTNNB1,TSHZ

H1,PRKD1,TCF4,CTR9,HDAC4,ZFPM2,NEDD4L,ZMIZ1,CITED1,LDB1,FGFR1,XRCC6,SLC30A9,MEF2C,SO

1,F2,PDE8B,CACNA2D2,KCNG2,PCLO,NEDD4L,CHRNA3,CACNA1C,PRKAG2,GRIK1,MAGI2,CTNNB1,SYT

D2,KCNG2,PCLO,CALU,ARFGEF2,CHRNA3,CACNA1C,GRIK1,SYT1,KCNB1,FURIN,CADPS,HLA-DRB5,BT

D2,KCNG2,PCLO,CALU,ARFGEF2,CHRNA3,CACNA1C,SYT1,KCNB1,FURIN,CADPS,HLA-DRB5,BTN3A2,

PT,EFL1,TCF4,HFE,KCNG2,CHRNA3,MAGI2,SNAP91,CTNNB1,SYT1,KCNB1,SRR,TWF2,CADPS,BCL11A,

KD1,MDK,DPP4,F2,CUL3,FGFR1,CARMIL1,ZSWIM6,MAGI2,CTNNB1,PRKG1,PTPRO,PDE4B,SEMA3F,NT[illegible]

TCH1,F2,HFE,NEDD4L,ATP2A2,CKB,CHRNA3,CACNA1C,AMIGO1,GRIK1,CD55,CTNNB1,ANK3,CNNM2,F[illegible]

R2,UTY,PTCH1,F2,ANAPC4,TCF4,CTR9,USP4,TNRC6A,HDAC4,PPM1E,GABBR1,ZFPM2,NEDD4L,FXR1,

SMA5,ESR2,UTY,PTCH1,F2,ANAPC4,TCF4,CTR9,USP4,TNRC6A,HDAC4,PPM1E,GABBR1,ZFPM2,NEDD4

RG,PTCH1,PRKD1,TAF1C,TCF4,CTR9,HDAC4,ZFPM2,NEDD4L,NR1D2,ZMIZ1,CITED1,LDB1,FGFR1,XRCC

,ANAPC4,CUL3,RPTOR,FGFR1,PRKAG2,MAD1L1,MAGI2,CTNNB1,RFC2,RBL2,PPP1R13B,SRPK2,PPP2R

OR,NEDD4L,CITED1,CHRNA3,FGFR1,PRKAG2,MEF2C,MAGI2,RASGRP4,CTNNB1,NGEF,PTPRF,FURIN,

22,GRIN2A,GIGYF2,WRN,TMX2,PTCH1,F2,HFE,NEDD4L,ATP2A2,CKB,CHRNA3,CACNA1C,LDB1,XRCC6

A5,WRN,PRKD1,PKP4,ANAPC4,MYO9B,RPTOR,MMP16,CHRNA3,FGFR1,PRKAG2,MEF2C,GMIP,MAGI2,

AI1,GRIN2A,MDK,CHRNA5,RPTOR,NEDD4L,CITED1,CHRNA3,FGFR1,PRKAG2,MEF2C,SOX5,CD55,CTNN

KMT2A,MSL2,RPA3,KMT2E,RERE,FOXP1,MAPT,WRN,EFL1,KIZ,PRKD1,ANAPC4,CTR9,SGF29,CUL3,PCL

EMA3A,GRIN2A,GIGYF2,UTY,TSNARE1,MDK,BBS9,CHRNA5,CACNA2D2,KCNG2,CNTNAP2,PKD1L3,PCL

2E,DPP6,ARHGAP15,GNAI1,ADAM22,SYNGAP1,SEMA3A,GRIN2A,GIGYF2,MAPT,WRN,TSNARE1,TMX2,F

ZNF536,CUL3,NEDD4L,CBLB,CITED1,MAGI2,CTNND1,CD55,PTPRO,SEMA3F,RGS6,WNT3,TRIM33,GLI3,

TCH1,CACNA2D2,CTR9,HDAC4,ZFPM2,COL11A1,EXT1,ATP2A2,CITED1,LDB1,FGFR1,CREB5,MEF2C,MA

A,PRKD1,MDK,F2,KLC1,DOCK8,HDAC4,ZFPM2,DYSF,CALU,ATP2A2,GUCY1A2,CARMIL1,CD55,PRKG1,J

PP4,CTR9,CBLB,LDB1,RAET1E,HLA-DQA1,MEF2C,CD55,CTNNB1,ABCB9,MICA,FOXO3,HLA-DRB5,HLA-D

SYNGAP1,PSMA5,WRN,PRKD1,PKP4,ANAPC4,ITIH3,MYO9B,RPTOR,PPM1E,GABBR1,MMP16,CHRNA3,[illegible]

1A1,ZMIZ1,CITED1,FGFR1,MEF2C,MAGI2,PALB2,CTNNB1,TSHZ3,WNT3,THRB,ATP2B2,BCL11B,GLI3

)1,DPP4,F2,CUL3,HDAC4,CBLB,ZMIZ1,FGFR1,MEF2C,MAGI2,CTNNB1,FOXO3,HLA-DRB5,HLA-DRB1,SRI

P4,F2,ANAPC4,USP4,CUL3,NEDD4L,MMP16,ATP23,FURIN,KEL,RNF111,LONRF2,DPP8,IMMP2L,PPP2R5(

AP1,FOXP1,SEMA3A,FEZ1,ESR2,MAPT,PTCH1,PRKD1,F2,SPAG16,CTR9,CUL3,ZFPM2,CNTN4,COL11A1, [illegible]CNG2,PKD1L3,NEDD4L,ATP2A2,CHRNA3,CACNA1C,SLC13A1,PRKAG2,SLC30A9,GRIK1,CTNNB1,SYT1,

[illegible]1,DPP4,MRM2,F2,CUL3,HDAC4,CBLB,ZMIZ1,CITED1,FGFR1,MEF2C,DBN1,MAGI2,RASGRP4,PALB2,CT[illegible]

L3,PCLO,DYSF,NEDD4L,CALU,ARFGEF2,CACNA1C,MAGI2,SNAP91,CTNNB1,SYT1,ANK3,CADPS,GBF1,

IGAP15,SYNGAP1,FOXP1,PSMA5,ESR2,WRN,PTCH1,PRKD1,PKP4,ANAPC4,ITIH3,MYO9B,RPTOR,HDAC

IGAP1,GRIN2A,VRK2,ESR2,SPAG16,HFE,KLC1,CNTNAP2,ZFPM2,THOC7,NEDD4L,SND1,ZMIZ1,CACNA1

N9,HFE,CTR9,KLC1,THOC7,NEDD4L,SND1,CITED1,ZSCAN12,XRCC6,ZSCAN31,MEF2C,ZKSCAN3,CD55,

KP1,PSMA5,ESRRG,PRKD1,MDK,PKP4,F2,ANAPC4,TCF4,CTR9,RPTOR,HDAC4,ZFPM2,CBLB,ZNF398,ZM

GAP15,GNAI1,CISD2,SYNGAP1,PSMA5,PXDNL,MAPT,WRN,EFL1,PKP4,PDE8B,ANAPC4,USP4,MYO9B,CL

KP1,PSMA5,ESRRG,PRKD1,MDK,F2,ANAPC4,TCF4,CTR9,RPTOR,HDAC4,ZFPM2,ZNF398,ZMIZ1,CITED1,

,GRIN2A,UTY,TSNARE1,PTCH1,EXOC4,USP4,SGF29,CNTNAP2,CBLB,ARFGEF2,DBN1,MAGI2,CTNNB1,A

RPTOR,HDAC4,CBLB,CHRNA3,LDB1,FGFR1,PRKAG2,CTNNB1,FURIN,RNF111,NTRK3,RHOA

ARE1,PTCH1,EXOC4,SGF29,CBLB,ARFGEF2,ANK3,ABCB9,CADPS,POM121C,HLA-DRB5,BTN3A2,PPP2R2

IGAP1,GRIN2A,VRK2,ESR2,SPAG16,HFE,KLC1,CNTNAP2,ZFPM2,THOC7,NEDD4L,SND1,ZMIZ1,CACNA1

PT,WRN,TSNARE1,PTCH1,PRKD1,F2,DYNC1I1,CUL3,THOC7,CBLB,ATP2A2,ARFGEF2,CITED1,CACNA1C

EZ1,RSRC1,MAPT,WRN,TSNARE1,PTCH1,PRKD1,EXOC4,F2,PDE8B,CACNA2D2,DYNC1I1,KCNG2,USP4

MDK,PKP4,ANAPC4,CUL3,RPTOR,MPHOSPH9,MSH5,PRKAG2,MAD1L1,MAGI2,CTNNB1,NEK4,STAG1,AN

1,MAPT,WRN,TSNARE1,PTCH1,PRKD1,EXOC4,F2,PDE8B,CACNA2D2,DYNC1I1,KCNG2,CUL3,PCLO,THC

5,VRK2,RSU1,PRKD1,PLCL1,F2,CTR9,MYO9B,CUL3,DOCK8,RPTOR,PCLO,HDAC4,CBLB,ARFGEF2,CACI

,PTCH1,EXOC4,CBLB,ARFGEF2,ANK3,ABCB9,CADPS,POM121C,HLA-DRB5,BTN3A2,PPP2R2B,HLA-DRB

A5,TET2,KIZ,PTCH1,MDK,PKP4,ANAPC4,CUL3,RPTOR,MPHOSPH9,DMTF1,FGFR1,MSH5,PRKAG2,MAD1

KD1,F2,RPTOR,CBLB,CITED1,FGFR1,HLA-DQA1,MEF2C,RASGRP4,CD55,CTNNB1,ABCB9,MICA,HLA-DF

-B,KMT2E,GNAI1,GRIN2A,FEZ1,VRK2,PTCH1,PRKD1,MDK,CHRNA5,CTR9,RPTOR,HDAC4,NEDD4L,CITE

WRN,PRKD1,HFE,TNRC6A,MSH5,XRCC6,SLC30A9,MEF2C,ATP23,PALB2,MICA,FOXO3,RFC2,POLR2E,P

5,ESR2,WRN,CUL3,CITED1,FGFR1,MEF2C,PALB2,CTNNB1,FURIN,FOXO3,PPP2R2B,PPP1R13B,SRPK2,

2,ANAPC4,CTR9,USP4,TNRC6A,RPTOR,HDAC4,PPM1E,NEDD4L,CBLB,FXR1,CHRNA3,LDB1,FGFR1,PRK

5,ESR2,WRN,CUL3,CITED1,FGFR1,MEF2C,PALB2,CTNNB1,FURIN,FOXO3,PPP2R2B,PPP1R13B,SRPK2,

4,CTR9,USP4,TNRC6A,RPTOR,HDAC4,PPM1E,FXR1,CHRNA3,FGFR1,PRKAG2,ETF1,FURIN,CPEB1,FOX

KP1,NCOA5,ESR2,ESRRG,UTY,PTCH1,PRKD1,MDK,PDE8B,ZNF536,TAF1C,TCF4,CTR9,SGF29,RPTOR,H

KP1,NCOA5,ESR2,ESRRG,UTY,PTCH1,PRKD1,MDK,PDE8B,ZNF536,TAF1C,TCF4,CTR9,SGF29,RPTOR,H

KP1,NCOA5,ESR2,ESRRG,UTY,PTCH1,PRKD1,MDK,PDE8B,ZNF536,TAF1C,TCF4,CTR9,SGF29,RPTOR,H

B,CITED1,RAET1E,HLA-DQA1,MEF2C,CD55,ABCB9,TANK,MICA,HLA-DRB5,BTN3A2,HLA-DRB1,SRPK2,HL

,GRIN2A,UTY,TSNARE1,PTCH1,EXOC4,USP4,SGF29,CNTNAP2,SPNS1,THOC7,CBLB,KIF5C,ARFGEF2,P

KP1,NCOA5,ESR2,ESRRG,UTY,PTCH1,PRKD1,MDK,PDE8B,ZNF536,TAF1C,TCF4,CTR9,SGF29,RPTOR,H

GAP15,GNAI1,CISD2,SYNGAP1,GRIN2A,PSMA5,PXDNL,MAPT,WRN,EFL1,PKP4,PDE8B,ANAPC4,USP4,M

3E,HLA-B,DGKI,GNAI1,SEMA3A,GRIN2A,GIGYF2,VRK2,GPR141,PTCH1,PRKD1,F2,CTR9,TNRC6A,CUL3

KI,KMT2E,ARHGAP15,SYNGAP1,FOXP1,NCOA5,ESR2,ESRRG,UTY,PTCH1,PRKD1,MDK,PKP4,PDE8B,

DMA,PSMA5,BANK1,TET2,PRKD1,DPP4,F2,DYNC1I1,HFE,CTR9,KLC1,HDAC4,CBLB,CITED1,LDB1,RAET

T2E,ADAM22,SYNGAP1,FOXP1,SEMA3A,GIGYF2,PSMA5,ESR2,UTY,PTCH1,PRKD1,DPP4,F2,PDE8B,Z

P1,GIGYF2,NCOA5,ESR2,ESRRG,UTY,TET2,PTCH1,PRKD1,MDK,PKP4,PDE8B,ZNF536,TAF1C,TCF4,CT

P1,SEMA3A,HLA-DMA,ESR2,UTY,TET2,PTCH1,MDK,CACNA2D2,CTR9,CNTNAP2,HDAC4,ZFPM2,CNTN

DS2,PFAS,EXT1,CKB,GATB,PRKAG2,CARMIL1,CTNNB1,ST3GAL3,SRR,DPYD,IVD,PSPH,TNXB,PPA2

NGAP1,VRK2,ESR2,PTCH1,PRKD1,PKP4,F2,ZNF536,MYO9B,CUL3,RPTOR,NEDD4L,CBLB,ARFGEF2,CI

/IT2D,KMT2A,HLA-B,KMT2E,GNAI1,FOXP1,SEMA3A,GRIN2A,FEZ1,VRK2,PXDNL,RSRC1,PTCH1,PRKD1,F

3A,GRIN2A,PSMA5,ESR2,MAPT,WRN,PRKD1,CUL3,FXR1,CITED1,FGFR1,MEF2C,PALB2,CTNNB1,NGEF,I

3A,GRIN2A,PSMA5,ESR2,MAPT,WRN,PRKD1,CUL3,FXR1,CITED1,FGFR1,MEF2C,PALB2,CTNNB1,NGEF,I

,ARHGAP15,SYNGAP1,SEMA3A,HLA-DMA,VRK2,ESR2,PTCH1,PRKD1,MDK,PKP4,F2,ZNF536,MYO9B,CU

3A,GRIN2A,PSMA5,ESR2,MAPT,WRN,PRKD1,CUL3,FXR1,CITED1,FGFR1,MEF2C,PALB2,CTNNB1,NGEF,I

T2A,HLA-B,KMT2E,ADAM22,SYNGAP1,FOXP1,SEMA3A,RTN1,HLA-DMA,GRIN2A,GIGYF2,FEZ1,ESR2,MA

KMT2D,KMT2A,MSL2,MSRA,KMT2E,PSMA5,VRK2,RSRC1,TET2,PRKD1,MRM2,F2,ANAPC4,CTR9,USP4,S

KMT2D,KMT2A,MSL2,MSRA,KMT2E,PSMA5,VRK2,RSRC1,PRKD1,F2,ANAPC4,CTR9,USP4,SGF29,CUL3,I

PM1E,CHRNA3,FGFR1,PRKAG2,IMPA2,NEK4,PRKG1,PIP5K1B,RBL2,POLR2E,CCDC88C,SRPK2,NTRK3,T

/IT2E,ADAM22,SYNGAP1,FOXP1,SEMA3A,GRIN2A,GIGYF2,PSMA5,ESR2,MAPT,UTY,PTCH1,PRKD1,DPF

PM1E,CHRNA3,FGFR1,PRKAG2,NEK4,PRKG1,POLR2E,CCDC88C,SRPK2,NTRK3,TNXB,TLK1

ED1,HLA-DQA1,MEF2C,CD55,TANK,MICA,HLA-DRB5,HLA-DRB1,SRPK2,HLA-DQB1,ELMO1,HDAC9

2E,FOXP1,HLA-DMA,GRIN2A,FEZ1,PSMA5,ESR2,MAPT,ESRRG,PRKD1,MDK,DPP4,PKP4,F2,ANAPC4,TCF

RE,FOXP1,NCOA5,ESR2,ESRRG,UTY,ZSCAN9,PTCH1,PRKD1,MDK,PDE8B,ZNF536,TAF1C,NPAS1,TCF4,

RE,FOXP1,NCOA5,ESR2,ESRRG,UTY,ZSCAN9,PTCH1,PRKD1,MDK,PDE8B,ZNF536,TAF1C,NPAS1,TCF4,

T2D,KMT2A,HLA-B,KMT2E,ADAM22,RERE,CISD2,SYNGAP1,FOXP1,SEMA3A,RTN1,HLA-DMA,GRIN2A,GI

T2D,KMT2A,HLA-B,KMT2E,ARHGAP15,ADAM22,SYNGAP1,FOXP1,SEMA3A,RTN1,HLA-DMA,GRIN2A,GIG

RE,FOXP1,NCOA5,ESR2,ESRRG,UTY,ZSCAN9,PTCH1,PRKD1,MDK,PDE8B,ZNF536,TAF1C,NPAS1,TCF4,

GKI,KMT2E,ARHGAP15,CISD2,SYNGAP1,FOXP1,GIGYF2,PSMA5,NCOA5,ESR2,MAPT,ESRRG,UTY,PTCH

2E,FOXP1,HLA-DMA,GRIN2A,FEZ1,PSMA5,ESR2,MAPT,ESRRG,PTCH1,PRKD1,MDK,DPP4,PKP4,F2,ANAF

KMT2E,RERE,FOXP1,NCOA5,ESR2,ESRRG,WRN,UTY,ZSCAN9,PTCH1,PRKD1,MDK,PDE8B,ZNF536,TAF1

E,HLA-DMA,GRIN2A,FEZ1,RSRC1,MAPT,WRN,UTY,TSNARE1,PTCH1,PRKD1,EXOC4,SLC22A23,F2,PDE8

E,HLA-DMA,GRIN2A,FEZ1,RSRC1,MAPT,WRN,TSNARE1,PTCH1,PRKD1,EXOC4,SLC22A23,F2,PDE8B,CH

KMT2D,KMT2A,ERLIN1,MSL2,MSRA,KMT2E,MRPL33,HLA-DMA,GIGYF2,PSMA5,VRK2,RSRC1,PRKD1,F2

RPA3,GNAI1,GRIN2A,PSMA5,VRK2,PXDNL,WRN,PRKD1,MDK,DPP4,F2,HFE,KLC1,TNRC6A,DOCK8,HDA

C,SEMA3E,KMT2D,HLA-B,DGKI,KMT2E,ARHGAP15,GNAI1,SYNGAP1,SEMA3A,RTN1,GRIN2A,GIGYF2,PS

MT2A,DGKI,KMT2E,ARHGAP15,CISD2,SYNGAP1,FOXP1,GRIN2A,GIGYF2,PSMA5,NCOA5,ESR2,MAPT,ES

RE,FOXP1,PSMA5,RSRC1,NCOA5,ESR2,ESRRG,UTY,ZSCAN9,PTCH1,PRKD1,MDK,TYW1B,MRM2,PDE8

KMT2D,KMT2A,ERLIN1,MSL2,MSRA,KMT2E,DPP6,ADAM22,MRPL33,HLA-DMA,GRIN2A,GIGYF2,PSMA5,

RPA3,KMT2E,MRPL33,RERE,FOXP1,GIGYF2,NCOA5,ESR2,ESRRG,WRN,UTY,ZSCAN9,PTCH1,PRKD1,MD

RPA3,KMT2E,MRPL33,RERE,FOXP1,GIGYF2,NCOA5,ESR2,ESRRG,WRN,UTY,ZSCAN9,PTCH1,PRKD1,MD

KMT2A,DGKI,RPA3,KMT2E,ARHGAP15,GNAI1,RERE,SYNGAP1,FOXP1,PSMA5,RSRC1,NCOA5,ESR2,ESR

KMT2A,DGKI,RPA3,KMT2E,ARHGAP15,GNAI1,RERE,SYNGAP1,FOXP1,PSMA5,RSRC1,NCOA5,ESR2,ESR

,MYO18A,KMT2A,MSRA,DGKI,RPA3,KMT2E,ARHGAP15,GNAI1,RERE,SYNGAP1,FOXP1,GRIN2A,PSMA5,

D,KMT2A,RPA3,KMT2E,MRPL33,RERE,FOXP1,GIGYF2,AGMO,NCOA5,ESR2,ESRRG,WRN,UTY,FADS2,ZS

ALNT12,KMT2D,MYO18A,KMT2A,ERLIN1,MSL2,MSRA,RPA3,KMT2E,DPP6,ADAM22,MRPL33,RERE,FOXP

D,KMT2A,RPA3,KMT2E,MRPL33,RERE,FOXP1,GIGYF2,AGMO,NCOA5,ESR2,ESRRG,WRN,UTY,FADS2,ZS

ALNT12,KMT2D,MYO18A,KMT2A,ERLIN1,MSL2,MSRA,RPA3,KMT2E,MRPL33,RERE,FOXP1,HLA-DMA,GIG

,MYO18A,KMT2A,DGKI,RPA3,KMT2E,ARHGAP15,GNAI1,RERE,SYNGAP1,FOXP1,GRIN2A,PSMA5,RSRC1

26,PDE1C,SEMA3E,KMT2D,KMT2A,HLA-B,DGKI,RPA3,KMT2E,ARHGAP15,GNAI1,SYNGAP1,SEMA3A,RTN

,MYO18A,KMT2A,DGKI,RPA3,KMT2E,ARHGAP15,GNAI1,RERE,SYNGAP1,FOXP1,GRIN2A,PSMA5,RSRC1

NNB1,NGEF,TWF2,BCL11A,PTPRO,SEMA3F,RGS6,NTRK3,WNT3,PREX1,TENM4,BCL11B,GLI3,RHOA

EF2C,CTNNB1,NGEF,CACNB2,ANK3,CACNA1I,TWF2,ESRRB,BCL11A,PTPRO,SEMA3F,TENM2,NTRK3,W

5C,CHRNA3,CACNA1C,LDB1,FGFR1,XRCC6,AMIGO1,MEF2C,ZSWIM6,DBN1,SOX5,CTNNB1,NGEF,NTM,

1,ZSWIM6,DBN1,NGEF,CACNB2,ANK3,PRKG1,CACNA1I,TWF2,BCL11A,PTPRO,SEMA3F,TENM2,NTRK3,

EMA6D,EXT1,KIF5C,CHRNA3,CACNA1C,LDB1,FGFR1,XRCC6,AMIGO1,MEF2C,ZSWIM6,DBN1,SOX5,CT

,SOX5,CTNNB1,PRKG1,SRR,CTTNBP2,PPP2R2B,NTRK3,WNT3,MACROD2,ATP2B2,TENM4,BCL11B,GLI

A3,CACNA1C,LDB1,FGFR1,AMIGO1,MEF2C,ZSWIM6,DBN1,SOX5,NGEF,NTM,CACNB2,ANK3,PRKG1,CA

1,MEF2C,ZSWIM6,DBN1,NGEF,NTM,CACNB2,ANK3,PRKG1,CACNA1I,TWF2,BCL11A,PTPRO,SEMA3F,TE

D1,CHRNA3,LDB1,FGFR1,XRCC6,AMIGO1,MEF2C,ZSWIM6,DBN1,SOX5,CTNNB1,NGEF,MED27,TWF2,ES

HDAC4,CNTN4,COL11A1,SEMA6D,EXT1,KIF5C,CITED1,CHRNA3,CACNA1C,LDB1,FGFR1,XRCC6,AMIGC

C,AMIGO1,MEF2C,GRIK1,DBN1,CTNNB1,SYT1,CACNB2,KCNB1,ANK3,CADPS,SHISA9,KCNN2,NOVA1,AT

36,NPAS1,TCF4,CNTNAP2,HDAC4,CNTN4,SEMA6D,EXT1,KIF5C,CITED1,CKB,CHRNA3,CACNA1C,TTLL7

,ZFPM2,CNTN4,CBLB,CITED1,CHRNA3,LDB1,FGFR1,XRCC6,AMIGO1,MEF2C,RHOJ,ZSWIM6,DBN1,MAG

:6,SLC30A9,CREB5,MEF2C,ZKSCAN3,SOX5,CTNNB1,ESRRB,MAML3,BCL11A,FOXO3,RNF111,ZMIZ2,NF

:6,SLC30A9,CREB5,MEF2C,ZKSCAN3,SOX5,CTNNB1,ESRRB,MAML3,BCL11A,FOXO3,RNF111,ZMIZ2,NF

C6,SLC30A9,CREB5,MEF2C,ZKSCAN3,SOX5,CTNNB1,ESRRB,MAML3,BCL11A,FOXO3,RNF111,POLR2E,Z

GFR1,RASGRP4,CD55,CACNB2,ANK3,PRKG1,CACNA1I,PSPH,PTPRO,HLA-DRB5,PDE4B,SEMA3F,HLA-D

N,CNTN4,COL11A1,CHRNA3,CACNA1C,FGFR1,AMIGO1,MEF2C,GRIK1,DBN1,CTNNB1,SYT1,CACNB2,K

Y1A2,XRCC6,SLC30A9,CREB5,MEF2C,ZKSCAN3,SOX5,CTNNB1,ESRRB,MAML3,BCL11A,FOXO3,RNF111

GO1,ETF1,RHOJ,MAD1L1,DBN1,MAGI2,CTNNB1,NGEF,TWF2,PPP1R16B,BCL11A,PTPRO,SEMA3F,TENM

G2,CTR9,HDAC4,ZFPM2,CNTN4,CBLB,ATP2A2,ARFGEF2,CITED1,CHRNA3,CACNA1C,LDB1,FGFR1,XRC

RMIL1,MAGI2,CADM2,CTNND1,SNAP91,CTNNB1,SYT1,KCNB1,ANK3,SRR,TWF2,PPP1R16B,CADPS,BCL

R1,MEF2C,GRIK1,DBN1,CTNNB1,SYT1,CACNB2,KCNB1,CADPS,HLA-DRB5,SHISA9,KCNN2,HLA-DRB1,N

,ZMIZ1,KIF5C,CITED1,CHRNA3,CACNA1C,LDB1,FGFR1,AMIGO1,MEF2C,ZSWIM6,MAGI2,PALB2,CTNNB1

X5,CTNNB1,MED27,ESRRB,MAML3,BCL11A,FOXO3,LCORL,RNF111,ZMIZ2,TENM2,NFIX,THRB,BRWD1,T

1,CACNB2,KCNB1,ANK3,HLA-DRB5,KCNN2,HLA-DRB1,ATP2B2,BTN2A2,SORBS1,GLI3,RHOA

CITED1,LDB1,FGFR1,PRKAG2,XRCC6,MEF2C,SOX5,CTNNB1,FURIN,TSHZ3,FOXO3,KDM4A,TENM2,NFIX

L,SND1,FXR1,CITED1,LDB1,FGFR1,PRKAG2,XRCC6,MEF2C,SOX5,CD55,CTNNB1,FURIN,BCL11A,TSHZ3

C6,SLC30A9,CREB5,MEF2C,SOX5,CTNNB1,MED27,ESRRB,MAML3,BCL11A,FOXO3,LCORL,RNF111,POLI

,AMIGO1,MEF2C,GRIK1,CD55,CTNNB1,ANK3,CNNM2,FOXO3,RFC2,HLA-DRB5,PDE4B,HLA-DRB1,CLCN3

B1,FURIN,SRR,PSPH,FOXO3,RNF111,NTRK3,TRIM33,SORBS1,FOXO6,RHOA,PIK3R2,HDAC9

.O,HDAC4,PPM1E,L3MBTL4,DYSF,KIF5C,ARFGEF2,LDB1,GATB,ARID1B,MSH5,XRCC6,CARMIL1,RHOJ,M

O,HDAC4,GABBR1,BSN,CNTN4,COL11A1,NEDD4L,ATP2A2,CHRNA3,CACNA1C,FGFR1,AMIGO1,MEF2C,

PTCH1,F2,PDE8B,CACNA2D2,HFE,KCNG2,USP4,KLC1,DOCK8,RPTOR,PCLO,ZFPM2,CNTN4,NEDD4L,CA

GI2,PALB2,CTNND1,SOX5,CTNNB1,CACNB2,TSHZ3,FOXO3,PTPRO,SEMA3F,WNT3,ATP2B2,TENM4,BCI

[illegible]GFR1,PRKAG2,MEF2C,GMIP,MAGI2,RASGRP4,NGEF,FURIN,PRKG1,RBL2,ZFYVE1,PPP2R2B,RGS6,NT[illegible]

SEMA6D,EXT1,ZMIZ1,KIF5C,CITED1,CHRNA3,CACNA1C,LDB1,FGFR1,AMIGO1,MEF2C,RHOJ,ZSWIM6,M

NNB1,FURIN,ESRRB,FOXO3,HLA-DRB5,HLA-DRB1,SRPK2,NTRK3,WNT3,BTN2A2,PPP2R5C,TENM4,TRA

:4,PPM1E,GABBR1,NEDD4L,MMP16,CHRNA3,FGFR1,PRKAG2,MEF2C,GMIP,MAGI2,RASGRP4,NGEF,ANI

C,CD55,CTNNB1,FURIN,MICA,FOXO3,KDM4A,POLR2E,PPP2R2B,SRPK2,NTRK3,IMMP2L,THRB,ATP2B2,

FURIN,SRR,MICA,KDM4A,HLA-DRB5,PDE4B,POLR2E,HLA-DRB1,SRPK2,NTRK3,ELMO1,THRB,ZKSCAN8

IIZ1,CITED1,CHRNA3,LDB1,ARID1B,FGFR1,GUCY1A2,PRKAG2,XRCC6,SLC30A9,CREB5,MEF2C,ZKSCAI

JL3,HDAC4,NEDD4L,NT5C2,CNOT1,PRKAG2,ETF1,GMIP,RHOJ,NGEF,FURIN,PRKG1,DPYD,IVD,PDE4B,F

,CHRNA3,LDB1,ARID1B,FGFR1,GUCY1A2,PRKAG2,XRCC6,SLC30A9,CREB5,MEF2C,ZKSCAN3,MAGI2,SC

NK3,ABCB9,TWF2,CADPS,POM121C,HLA-DRB5,BTN3A2,PPP2R2B,HLA-DRB1,SYNE1,IMMP2L,BTN2A2,

C,MSH5,CD55,CTNNB1,FURIN,MICA,FOXO3,KDM4A,POLR2E,PPP2R2B,SRPK2,NTRK3,IMMP2L,THRB,AT

,SGF29,CUL3,CNTNAP2,PCLO,THOC7,CBLB,CALU,ATP2A2,ARFGEF2,CITED1,CHRNA3,CACNA1C,PRKA

)C7,CBLB,CALU,ATP2A2,ARFGEF2,CITED1,CHRNA3,CACNA1C,PRKAG2,SYT1,KCNB1,ANK3,FURIN,CAD

NA1C,FGFR1,GUCY1A2,PRKAG2,MEF2C,GMIP,RHOJ,MAGI2,RASGRP4,CTNNB1,NGEF,PLCL2,TANK,FO

L1,MAGI2,CTNNB1,NEK4,STAG1,ANK3,RFC2,RBL2,PARD3B,PPP1R13B,SRPK2,PPP2R5C,RHOA,TLK1

D1,CHRNA3,FGFR1,HLA-DQA1,PRKAG2,MEF2C,SOX5,CD55,CTNNB1,FURIN,SRR,TWF2,PSPH,FOXO3,

DAC4,TCF20,ZFPM2,NEDD4L,SND1,ZNF398,NR1D2,ZNF823,ZMIZ1,DMTF1,CITED1,LDB1,ARID1B,ZNF63

IDAC4,TCF20,ZFPM2,NEDD4L,SND1,ZNF398,NR1D2,ZNF823,ZMIZ1,DMTF1,CITED1,LDB1,ARID1B,ZNF63

IDAC4,TCF20,ZFPM2,NEDD4L,SND1,ZNF398,NR1D2,ZNF823,ZMIZ1,DMTF1,CITED1,LDB1,ARID1B,ZNF63

RKAG2,DBN1,MAGI2,CTNNB1,ANK3,ABCB9,TWF2,CADPS,POM121C,HLA-DRB5,BTN3A2,PPP2R2B,HLA-

IDAC4,TCF20,ZFPM2,NEDD4L,SND1,ZNF398,NR1D2,ZNF823,ZMIZ1,DMTF1,CITED1,LDB1,ARID1B,ZNF63

YO9B,CUL3,HDAC4,NEDD4L,CBLB,GPD2,NT5C2,CNOT1,PRKAG2,ETF1,GMIP,RHOJ,IMPA2,NGEF,FURIN

,PKD1L3,RPTOR,GABBR1,NEDD4L,CBLB,ITGA11,CITED1,CHRNA3,LDB1,FGFR1,HLA-DQA1,PRKAG2,ME

NF536,TAF1C,TCF4,CTR9,SGF29,MYO9B,RPTOR,HDAC4,TCF20,GABBR1,ZFPM2,NEDD4L,SND1,ZNF39

1E,HLA-DQA1,XRCC6,MEF2C,RASGRP4,CD55,CTNNB1,ABCB9,TANK,MICA,BCL11A,FOXO3,PTPRO,HL

IF536,ANAPC4,TCF4,CTR9,USP4,TNRC6A,CUL3,RPTOR,HDAC4,PPM1E,GABBR1,ZFPM2,CNTN4,NEDD4

R9,USP4,SGF29,TNRC6A,RPTOR,HDAC4,TCF20,ZFPM2,NEDD4L,SND1,ZNF398,NR1D2,FXR1,ZNF823,Z

,COL11A1,FXR1,EXT1,ITGA11,ATP2A2,ZMIZ1,CITED1,CKB,LDB1,FGFR1,CREB5,MEF2C,MAGI2,RASGR

TED1,FGFR1,GMIP,RHOJ,MAGI2,RASGRP4,CTNND1,CTNNB1,NGEF,FURIN,PTPRO,GBF1,RNF111,RGS6

EF1,MDK,CHRNA5,CTR9,PKD1L3,RPTOR,HDAC4,PPM1E,CNTN4,PFAS,NEDD4L,SEMA6D,EXT1,KIF5C,C

FOXO3,TMEM219,PPP2R2B,PPP1R13B,SRPK2,ELMO1,PREX1,PPP2R5C,TRAIP,BCL11B,AMBRA1,CSRN

FOXO3,TMEM219,PPP2R2B,PPP1R13B,SRPK2,ELMO1,PREX1,PPP2R5C,TRAIP,BCL11B,AMBRA1,CSRN

L3,RPTOR,NEDD4L,CBLB,ARFGEF2,CITED1,RAET1E,FGFR1,HLA-DQA1,MEF2C,GMIP,RHOJ,MAGI2,RAS

FOXO3,TMEM219,PPP2R2B,PPP1R13B,SRPK2,ELMO1,PREX1,PPP2R5C,TRAIP,BCL11B,AMBRA1,CSRN

PT,UTY,TET2,PTCH1,PRKD1,MDK,F2,ZNF536,NPAS1,TCF4,CACNA2D2,CTR9,CNTNAP2,HDAC4,ZFPM2,

GF29,CUL3,RPTOR,HDAC4,PPM1E,NEDD4L,EXT1,CHRNA3,TTLL7,LDB1,FGFR1,PRKAG2,ETF1,MGAT5

RPTOR,HDAC4,PPM1E,NEDD4L,EXT1,CHRNA3,TTLL7,LDB1,FGFR1,PRKAG2,ETF1,MGAT5B,NEK4,ST3G

4,F2,PDE8B,ZNF536,ANAPC4,TCF4,CTR9,USP4,TNRC6A,CUL3,RPTOR,HDAC4,PPM1E,GABBR1,ZFPM2,

F4,CTR9,CUL3,RPTOR,HDAC4,PPM1E,ZFPM2,NEDD4L,ZNF398,ZMIZ1,CITED1,CHRNA3,LDB1,ARID1B,FC

CTR9,SGF29,RPTOR,HDAC4,L3MBTL4,TCF20,ZFPM2,NEDD4L,SND1,ZNF398,NR1D2,ZNF823,ZMIZ1,DM

CTR9,SGF29,RPTOR,HDAC4,L3MBTL4,TCF20,ZFPM2,NEDD4L,SND1,ZNF398,NR1D2,ZNF823,ZMIZ1,DM

GYF2,FEZ1,ESR2,MAPT,WRN,UTY,TET2,PTCH1,PRKD1,MDK,F2,ZNF536,NPAS1,TCF4,CACNA2D2,CTR9

YF2,FEZ1,ESR2,MAPT,UTY,TET2,PTCH1,PRKD1,MDK,F2,SPAG16,ZNF536,NPAS1,TCF4,CACNA2D2,CTF

CTR9,SGF29,RPTOR,HDAC4,L3MBTL4,TCF20,ZFPM2,NEDD4L,SND1,ZNF398,NR1D2,ZNF823,ZMIZ1,DM

1,PRKD1,MDK,PKP4,F2,PDE8B,ZNF536,TAF1C,ANAPC4,TCF4,CTR9,USP4,SGF29,TNRC6A,MYO9B,RPT

PC4,TCF4,CACNA2D2,CTR9,CUL3,RPTOR,HDAC4,PPM1E,ZFPM2,NEDD4L,CBLB,ZNF398,ATP2A2,ZMIZ1

1C,NPAS1,TCF4,CTR9,SGF29,RPTOR,HDAC4,L3MBTL4,TCF20,GABBR1,ZFPM2,PFAS,NEDD4L,SND1,ZN

3B,CHRNA5,CACNA2D2,DYNC1I1,HFE,KCNG2,SGF29,CUL3,SLC6A9,PKD1L3,PCLO,SPNS1,DYSF,THOC7

HRNA5,CACNA2D2,DYNC1I1,HFE,KCNG2,CUL3,SLC6A9,PKD1L3,PCLO,SPNS1,DYSF,THOC7,NEDD4L,CE

,ANAPC4,CTR9,USP4,SGF29,TNRC6A,CUL3,RPTOR,HDAC4,PPM1E,NEDD4L,FXR1,EXT1,CHRNA3,TTLL

C4,ZFPM2,DYSF,NEDD4L,CALU,ATP2A2,CITED1,MSH5,HLA-DQA1,GUCY1A2,XRCC6,SLC30A9,CARMIL1

MA5,VRK2,RSU1,ESR2,GPR141,PTCH1,PRKD1,MDK,PLCL1,F2,CHRNA5,ZNF536,CTR9,TNRC6A,MYO9B,

RRG,WRN,UTY,TET2,PTCH1,PRKD1,MDK,PKP4,F2,PDE8B,ZNF536,TAF1C,ANAPC4,TCF4,ITIH3,CTR9,U

,ZNF536,TAF1C,NPAS1,TCF4,CTR9,SGF29,RBM6,RPTOR,HDAC4,L3MBTL4,TCF20,ZFPM2,THOC7,NED

RK2,RSRC1,PTCH1,PRKD1,DPP4,F2,ANAPC4,CTR9,USP4,SGF29,TNRC6A,CUL3,RPTOR,HDAC4,PPM1

K,F2,PDE8B,ZNF536,TAF1C,NPAS1,TCF4,CTR9,FHIT,SGF29,TNRC6A,RPTOR,HDAC4,L3MBTL4,TCF20,

K,F2,PDE8B,ZNF536,TAF1C,NPAS1,TCF4,CTR9,FHIT,SGF29,TNRC6A,RPTOR,HDAC4,L3MBTL4,TCF20,

RG,WRN,UTY,TET2,EFL1,ZSCAN9,PTCH1,PRKD1,MDK,TYW1B,PKP4,MRM2,PDE8B,ZNF536,TAF1C,NP

RG,WRN,UTY,TET2,EFL1,ZSCAN9,PTCH1,PRKD1,MDK,TYW1B,PKP4,MRM2,PDE8B,ZNF536,TAF1C,NP

RSRC1,NCOA5,ESR2,ESRRG,WRN,UTY,TET2,EFL1,ZSCAN9,PTCH1,PRKD1,MDK,TYW1B,PKP4,MRM2,P

CAN9,PTCH1,PRKD1,MDK,F2,PDE8B,ZNF536,TAF1C,NPAS1,TCF4,HFE,CTR9,FHIT,SGF29,TNRC6A,RP

1,HLA-DMA,GRIN2A,GIGYF2,PSMA5,VRK2,RSRC1,NCOA5,ESR2,ESRRG,WRN,UTY,TET2,ZSCAN9,PTCH

CAN9,PTCH1,PRKD1,MDK,PDE8B,ZNF536,TAF1C,NPAS1,TCF4,CTR9,FHIT,SGF29,TNRC6A,RPTOR,HD

YF2,PSMA5,VRK2,RSRC1,NCOA5,ESR2,ESRRG,WRN,UTY,TET2,ZSCAN9,PTCH1,PRKD1,MDK,TYW1B,

,NCOA5,ESR2,ESRRG,WRN,UTY,TET2,EFL1,ZSCAN9,PTCH1,PRKD1,MDK,TYW1B,PKP4,MRM2,PDE8B,

1,GRIN2A,GIGYF2,FEZ1,PSMA5,VRK2,PXDNL,RSU1,ESR2,WRN,GPR141,PTCH1,PRKD1,MDK,PLCL1,F2

,NCOA5,ESR2,ESRRG,WRN,UTY,TET2,EFL1,ZSCAN9,PTCH1,PRKD1,MDK,TYW1B,PKP4,MRM2,PDE8B,

CACNB2,ANK3,PRKG1,CACNA1I,TWF2,BCL11A,PTPRO,SEMA3F,PPP2R2B,RGS6,TENM2,NTRK3,WNT3,

NNB1,NGEF,NTM,CACNB2,ANK3,PRKG1,CACNA1I,TWF2,BCL11A,PTPRO,SEMA3F,PPP2R2B,RGS6,TENI

CNA1I,TWF2,BCL11A,PTPRO,SEMA3F,PPP2R2B,RGS6,TENM2,NTRK3,WNT3,PREX1,ATP2B2,BCL11B,G

SRRB,BCL11A,TSHZ3,FOXO3,PTPRO,SEMA3F,RGS6,NTRK3,WNT3,PREX1,TENM4,BCL11B,FOXO6,GLI3

)1,MEF2C,ZSWIM6,DBN1,MAGI2,SOX5,CTNNB1,NGEF,NTM,CACNB2,ANK3,MED27,PRKG1,CACNA1I,TW [illegible],LDB1,ARID1B,FGFR1,XRCC6,AMIGO1,MEF2C,GRIK1,ZSWIM6,DBN1,CADM2,SOX5,CTNNB1,NGEF,NTM

I2,SOX5,CTNNB1,NGEF,MED27,TWF2,ESRRB,BCL11A,TSHZ3,FOXO3,PTPRO,SEMA3F,RGS6,NTRK3,W

C6,AMIGO1,MEF2C,GRIK1,ZSWIM6,DBN1,MAGI2,SOX5,CTNNB1,NGEF,FURIN,PRKG1,TWF2,BCL11A,TS

.11A,PTPRO,HLA-DRB5,HLA-DRB1,TENM2,CCDC88C,SRPK2,TNXB,PREX1,SORBS1,NEURL1,RHOA,TRI[illegible]

,NGEF,CACNB2,ANK3,CACNA1I,TWF2,ESRRB,BCL11A,PTPRO,SEMA3F,TENM2,SRPK2,NTRK3,WNT3,A

R2E,ZMIZ2,TENM2,NFIX,THRB,BRWD1,TRIM33,BCL11B,FOXO6,CSRNP3,MED8,GLI3,RHOA,HDAC9

IAD1L1,DBN1,SNAP91,CTNNB1,NEK4,STAG1,PRKG1,TWF2,CADPS,KDM4A,JMJD1C,RFC2,GBF1,RBL2,P

GRIK1,DBN1,CALD1,CTNNB1,SYT1,CACNB2,KCNB1,ANK3,CADPS,TSHZ3,PTPRO,KEL,SHISA9,KCNN2,N

LU,ATP2A2,CKB,CHRNA3,CACNA1C,LDB1,GUCY1A2,XRCC6,AMIGO1,CARMIL1,MEF2C,GRIK1,RHOJ,DB

MAGI2,PALB2,CTNND1,CTNNB1,NGEF,CACNB2,ANK3,CACNA1I,TWF2,ESRRB,BCL11A,TSHZ3,PTPRO,SE

K3,FURIN,PRKG1,FOXO3,RBL2,ZFYVE1,PPP2R2B,RGS6,NTRK3,TNXB,PREX1,DOCK4,FOXO6,CSRNP3,F

N3,MAGI2,SOX5,CTNNB1,ANK3,FURIN,ESRRB,MAML3,BCL11A,FOXO3,RNF111,POLR2E,ZMIZ2,NFIX,SR

OX5,CTNNB1,FURIN,ESRRB,MAML3,BCL11A,FOXO3,RNF111,POLR2E,ZMIZ2,NFIX,NTRK3,THRB,SORBS

\G2,DBN1,MAGI2,CTNNB1,SYT1,KCNB1,ANK3,FURIN,CADPS,GBF1,HLA-DRB5,BTN3A2,PPP2R2B,HLA-D

)PS,GBF1,HLA-DRB5,BTN3A2,PPP2R2B,HLA-DRB1,IMMP2L,BTN2A2,GLI3,RHOA,BTN3A1,TLK1

XO3,GBF1,NKIRAS1,RGS6,TENM2,SRPK2,NTRK3,TNXB,ELMO1,PREX1,PPP2R5C,DOCK4,RHOA,PIK3R2

HLA-DRB5,PDE4B,RNF111,HLA-DRB1,NTRK3,HLA-DQB1,WNT3,TRIM33,SORBS1,FOXO6,RHOA,PIK3R2,

8,FGFR1,CNOT1,XRCC6,SLC30A9,CREB5,MEF2C,ZNF568,ZNF19,TFB1M,ZKSCAN3,CTNND1,SOX5,CTN

8,FGFR1,CNOT1,XRCC6,SLC30A9,CREB5,MEF2C,ZNF568,ZNF19,TFB1M,ZKSCAN3,CTNND1,SOX5,CTN

8,FGFR1,CNOT1,XRCC6,SLC30A9,CREB5,MEF2C,ZNF568,ZNF19,TFB1M,ZKSCAN3,CTNND1,SOX5,CTN

8,FGFR1,CNOT1,XRCC6,SLC30A9,CREB5,MEF2C,ZNF568,ZNF19,TFB1M,ZKSCAN3,CTNND1,SOX5,CTN

I,PRKG1,DPYD,IVD,PDE4B,RNF111,RGS6,MANBA,NTRK3,PREX1,PPP2R5C,DOCK4,AMBRA1

:F2C,GRIK1,MAGI2,RASGRP4,CTNND1,CTNNB1,NGEF,PTPRF,FURIN,MICA,MAML3,FOXO3,PTPRO,HLA-

8,NR1D2,ZNF823,ZMIZ1,DMTF1,CITED1,LDB1,ARID1B,ZNF638,FGFR1,CNOT1,GUCY1A2,XRCC6,SLC30A

\-DRB5,PDE4B,BTN3A2,HLA-DRB1,SRPK2,HLA-DQB1,ELMO1,DPP8,WNT3,PREX1,BTN2A2,BCL11B,GLI3

L,CBLB,FXR1,CITED1,LDB1,FGFR1,PRKAG2,XRCC6,MEF2C,GRIK1,MAD1L1,MAGI2,PALB2,CTNND1,SOX

MIZ1,DMTF1,CITED1,LDB1,ARID1B,ZNF638,FGFR1,CNOT1,XRCC6,SLC30A9,ETF1,CREB5,MEF2C,ZNF56

P4,CADM2,PALB2,CTNND1,SOX5,CTNNB1,CACNB2,PRKG1,SRR,CTTNBP2,BCL11A,TSHZ3,FOXO3,PTPF

ITED1,CHRNA3,CACNA1C,FGFR1,HLA-DQA1,PRKAG2,MEF2C,SOX5,CD55,CTNNB1,SYT1,CACNB2,ANK

GRP4,CTNND1,CD55,CTNNB1,NGEF,ABCB9,FURIN,PRKG1,MICA,PTPRO,GBF1,HLA-DRB5,SEMA3F,RN

CNTN4,COL11A1,CBLB,SEMA6D,FXR1,EXT1,ITGA11,ATP2A2,ZMIZ1,KIF5C,CITED1,CKB,CHRNA3,CACN

,NEK4,ST3GAL3,PTPRF,FURIN,PRKG1,BCL11A,RFT1,PTPRO,KDM4A,RNF111,POLR2E,CCDC88C,MAN

AL3,PTPRF,FURIN,PRKG1,BCL11A,RFT1,PTPRO,KDM4A,RNF111,POLR2E,CCDC88C,MANBA,SRPK2,N

CNTN4,NEDD4L,CBLB,SND1,FXR1,ATP2A2,CITED1,CACNA1C,LDB1,FGFR1,PRKAG2,XRCC6,MEF2C,G

FR1,HLA-DQA1,GUCY1A2,PRKAG2,XRCC6,AMIGO1,SLC30A9,CREB5,MEF2C,GRIK1,ZKSCAN3,MAGI2,

TF1,CITED1,ZSCAN12,LDB1,ARID1B,ZNF638,FGFR1,CNOT1,XRCC6,SLC30A9,CREB5,ZSCAN31,MEF2C,

TF1,CITED1,ZSCAN12,LDB1,ARID1B,ZNF638,FGFR1,CNOT1,XRCC6,SLC30A9,CREB5,ZSCAN31,MEF2C,

,CUL3,CNTNAP2,HDAC4,ZFPM2,CNTN4,COL11A1,CBLB,SEMA6D,FXR1,EXT1,ITGA11,ATP2A2,ZMIZ1,

9,CUL3,CNTNAP2,HDAC4,ZFPM2,CNTN4,COL11A1,CBLB,SEMA6D,FXR1,EXT1,ITGA11,ATP2A2,ZMIZ1,

TF1,CITED1,ZSCAN12,LDB1,ARID1B,ZNF638,FGFR1,CNOT1,XRCC6,SLC30A9,CREB5,ZSCAN31,MEF2C,

R,HDAC4,PPM1E,TCF20,GABBR1,ZFPM2,NEDD4L,SND1,ZNF398,NR1D2,FXR1,ZNF823,ZMIZ1,DMTF1,

,ARFGEF2,CITED1,CHRNA3,LDB1,ARID1B,FGFR1,HLA-DQA1,GUCY1A2,PRKAG2,XRCC6,AMIGO1,SLC3

F398,NR1D2,ZNF823,ATP2A2,ZMIZ1,DMTF1,CITED1,ZSCAN12,LDB1,ARID1B,ZNF638,FGFR1,CNOT1,GU

,NEDD4L,CBLB,CALU,ATP2A2,KIF5C,ARFGEF2,CITED1,CHRNA3,CACNA1C,SLC13A1,PRKAG2,SLC30A

LB,CALU,ATP2A2,KIF5C,ARFGEF2,CITED1,CHRNA3,CACNA1C,SLC13A1,PRKAG2,SLC30A9,GRIK1,MA

7,LDB1,GATB,FGFR1,PRKAG2,ETF1,MGAT5B,NEK4,ST3GAL3,PTPRF,FURIN,PRKG1,CPEB1,BCL11A,FO

,MEF2C,ATP23,GRIK1,PALB2,CD55,PRKG1,TANK,MICA,FOXO3,JMJD1C,RFC2,HLA-DRB5,POLR2E,PPP2

CUL3,DOCK8,CNTNAP2,PKD1L3,RPTOR,PCLO,HDAC4,GABBR1,NEDD4L,CBLB,NR1D2,EXT1,ITGA11,AT

SP4,SGF29,TNRC6A,MYO9B,RPTOR,HDAC4,PPM1E,TCF20,GABBR1,ZFPM2,NEDD4L,CBLB,SND1,ZNF3

D4L,SND1,ZNF398,NR1D2,ZNF823,ZMIZ1,DMTF1,CITED1,ZSCAN12,LDB1,GATB,ARID1B,ZNF638,PRPF3,

E,COL11A1,NEDD4L,CBLB,FXR1,MMP16,EXT1,CHRNA3,TTLL7,LDB1,GATB,FGFR1,PRKAG2,ETF1,ATP2

ZFPM2,NEDD4L,SND1,ZNF398,NR1D2,FXR1,ZNF823,EXT1,ZMIZ1,DMTF1,CITED1,ZSCAN12,LDB1,GATB,

ZFPM2,NEDD4L,SND1,ZNF398,NR1D2,FXR1,ZNF823,EXT1,ZMIZ1,DMTF1,CITED1,ZSCAN12,LDB1,GATB,

AS1,TCF4,CTR9,FHIT,SGF29,RBM6,MYO9B,RPTOR,HDAC4,L3MBTL4,TCF20,GABBR1,ZFPM2,PFAS,THO

AS1,TCF4,CTR9,FHIT,SGF29,RBM6,MYO9B,RPTOR,HDAC4,L3MBTL4,TCF20,GABBR1,ZFPM2,PFAS,THO

PDE8B,ZNF536,TAF1C,NPAS1,TCF4,ITIH3,CTR9,FHIT,SGF29,RBM6,MYO9B,RPTOR,HDAC4,L3MBTL4,TC

TOR,HDAC4,L3MBTL4,TCF20,GABBR1,ZFPM2,PFAS,NEDD4L,GPD2,SND1,ZNF398,NR1D2,FXR1,ZNF823

1,PRKD1,MDK,TYW1B,DPP4,PKP4,MRM2,F2,PDE8B,ZNF536,TAF1C,NPAS1,ANAPC4,TCF4,ITIH3,CTR9,

AC4,L3MBTL4,TCF20,GABBR1,ZFPM2,PFAS,NEDD4L,SND1,ZNF398,NR1D2,FXR1,ZNF823,EXT1,ATP2A2

MRM2,F2,PDE8B,ZNF536,TAF1C,NPAS1,ANAPC4,TCF4,CTR9,USP4,FHIT,SGF29,RBM6,TNRC6A,CUL3,RP

ZNF536,TAF1C,NPAS1,TCF4,CTR9,FHIT,SGF29,RBM6,MYO9B,RPTOR,HDAC4,L3MBTL4,TCF20,GABBR1

,CHRNA5,ZNF536,HFE,CTR9,TNRC6A,MYO9B,CUL3,DOCK8,CNTNAP2,PKD1L3,RPTOR,PCLO,HDAC4,PP

ZNF536,TAF1C,NPAS1,TCF4,CTR9,FHIT,SGF29,RBM6,MYO9B,RPTOR,HDAC4,L3MBTL4,TCF20,GABBR1

F2,ESRRB,BCL11A,FOXO3,PTPRO,SEMA3F,PPP2R2B,RGS6,TENM2,NTRK3,WNT3,PREX1,ATP2B2,TEN[illegible]

,CACNB2,ANK3,PRKG1,CACNA1I,SRR,CTTNBP2,TWF2,BCL11A,PTPRO,SEMA3F,PPP2R2B,RGS6,TENM

HZ3,FOXO3,PTPRO,HLA-DRB5,SHISA9,SEMA3F,HLA-DRB1,RGS6,NTRK3,WNT3,THRB,PREX1,ATP2B2,

'PP2R2B,CCDC88C,SRPK2,NTRK3,TNXB,SYNE1,ELMO1,IMMP2L,CLCN3,PREX1,ATP2B2,BRWD1,KDM3E

N1,CD55,CTNNB1,SYT1,KCNB1,ANK3,FURIN,PRKG1,TWF2,CNNM2,CADPS,BCL11A,FOXO3,PTPRO,JM

EMA3F,TENM2,SRPK2,NTRK3,WNT3,THRB,ATP2B2,BRWD1,TENM4,BCL11B,FOXO6,GLI3,NEURL1,RHOA

NB1,PCGF3,MED27,ESRRB,MAML3,BCL11A,TSHZ3,FOXO3,LCORL,KDM4A,JMJD1C,ZNF600,RNF111,RB

NB1,PCGF3,MED27,ESRRB,MAML3,BCL11A,TSHZ3,FOXO3,LCORL,KDM4A,JMJD1C,ZNF600,RNF111,RB

NB1,PCGF3,MED27,ESRRB,MAML3,BCL11A,TSHZ3,FOXO3,LCORL,KDM4A,JMJD1C,ZNF600,RNF111,RB

NB1,PCGF3,MED27,ESRRB,MAML3,BCL11A,TSHZ3,FOXO3,LCORL,KDM4A,JMJD1C,ZNF600,RNF111,RB

-DRB5,RNF111,SORCS3,HLA-DRB1,RGS6,CCDC88C,NTRK3,HLA-DQB1,WNT3,PREX1,TRIM33,SORBS1,[illegible]

A9,CREB5,MEF2C,ZNF568,ZNF19,GMIP,TFB1M,ZKSCAN3,CTNND1,SOX5,CTNNB1,NGEF,PCGF3,MED27,

X5,CTNNB1,NGEF,FURIN,MED27,PRKG1,TWF2,ESRRB,BCL11A,TSHZ3,FOXO3,PTPRO,KDM4A,RFC2,HL

68,ZNF19,TFB1M,ZKSCAN3,CTNND1,SOX5,CD55,CTNNB1,PCGF3,ANK3,MED27,ESRRB,CPEB1,MAML3,[illegible]

RO,SEMA3F,PPP2R2B,NTRK3,IMMP2L,WNT3,THRB,MACROD2,ATP2B2,TENM4,BCL11B,FOXO6,GLI3,NE

3,FURIN,CACNA1I,SRR,TWF2,PSPH,FOXO3,PTPRO,HLA-DRB5,PDE4B,SEMA3F,RNF111,HLA-DRB1,TEN

F111,HLA-DRB1,RGS6,NTRK3,TNXB,HLA-DQB1,ELMO1,WNT3,PREX1,DOCK4,TRIM33,GLI3,RHOA,PIK3R

A1C,TTLL7,LDB1,ARID1B,FGFR1,XRCC6,AMIGO1,CREB5,MEF2C,GRIK1,ZSWIM6,DBN1,MAGI2,RASGRP

3A,SRPK2,NTRK3,GALNT1,TNXB,GALNT10,TRIM33,MED8,NEURL1,PUS7,TRIOBP,HDAC9,RNF144A,CDK

RIK1,MAD1L1,MAGI2,PALB2,CTNND1,SOX5,CD55,CTNNB1,NGEF,FURIN,MED27,PRKG1,TWF2,ESRRB,B

RASGRP4,SOX5,CTNNB1,ANK3,FURIN,TWF2,ESRRB,MAML3,BCL11A,TSHZ3,FOXO3,HLA-DRB5,RNF111

ZNF568,ZNF19,TFB1M,ZKSCAN3,CTNND1,SOX5,CTNNB1,PCGF3,MED27,ESRRB,MAML3,BCL11A,TSHZ3

ZNF568,ZNF19,TFB1M,ZKSCAN3,CTNND1,SOX5,CTNNB1,PCGF3,MED27,ESRRB,MAML3,BCL11A,TSHZ3

F5C,CITED1,CKB,CHRNA3,CACNA1C,TTLL7,LDB1,ARID1B,FGFR1,XRCC6,AMIGO1,CREB5,MEF2C,GRIK1

KIF5C,CITED1,CKB,CHRNA3,CACNA1C,TTLL7,LDB1,ARID1B,FGFR1,XRCC6,AMIGO1,CREB5,MEF2C,GRI

ZNF568,ZNF19,TFB1M,ZKSCAN3,CTNND1,SOX5,CTNNB1,PCGF3,MED27,ESRRB,MAML3,BCL11A,TSHZ3

CITED1,CHRNA3,LDB1,ARID1B,ZNF638,FGFR1,CNOT1,GUCY1A2,PRKAG2,XRCC6,SLC30A9,ETF1,CREB5

0A9,CREB5,MEF2C,GRIK1,ZKSCAN3,MAGI2,RASGRP4,SOX5,CD55,CTNNB1,CACNB2,ANK3,ABCB9,FURI

CY1A2,PRKAG2,XRCC6,SLC30A9,CREB5,ZSCAN31,MEF2C,ZNF568,ZNF19,TFB1M,ZKSCAN3,CTNND1,S

9,GRIK1,MAGI2,SNAP91,CTNNB1,SYT1,CACNB2,KCNB1,ANK3,ABCB9,FURIN,CACNA1I,CNNM2,CADPS,P

GI2,SNAP91,CTNNB1,SYT1,CACNB2,KCNB1,ANK3,ABCB9,FURIN,CACNA1I,CNNM2,CADPS,POM121C,SL

XO3,RFT1,PTPRO,KDM4A,RNF111,POLR2E,CCDC88C,MANBA,SRPK2,NTRK3,GALNT1,TNXB,GALNT10,

2R2B,HLA-DRB1,SRPK2,NTRK3,TNXB,HLA-DQB1,ELMO1,ATP2B2,PPP2R5C,FOXO6,RHOA,PIK3R2,HDAC

P2A2,ARFGEF2,CITED1,CHRNA3,CACNA1C,LDB1,FGFR1,HLA-DQA1,GUCY1A2,PRKAG2,MEF2C,GMIP,

98,NR1D2,FXR1,MMP16,ZNF823,ZMIZ1,DMTF1,CITED1,CHRNA3,LDB1,ARID1B,ZNF638,FGFR1,CNOT1,G

FGFR1,CNOT1,XRCC6,SLC30A9,ETF1,CREB5,ZSCAN31,MEF2C,ZNF568,ZNF19,TFB1M,ZKSCAN3,CTNN

3,MGAT5B,CD55,CTNNB1,NEK4,ST3GAL3,PTPRF,FURIN,PRKG1,CPEB1,BCL11A,FOXO3,RFT1,PTPRO,K

ARID1B,ZNF638,FGFR1,CNOT1,XRCC6,SLC30A9,ETF1,CREB5,ZSCAN31,MEF2C,ZNF568,ZNF19,MGAT5

ARID1B,ZNF638,FGFR1,CNOT1,XRCC6,SLC30A9,ETF1,CREB5,ZSCAN31,MEF2C,ZNF568,ZNF19,MGAT5

C7,NEDD4L,SND1,ZNF398,NR1D2,ZNF823,ATP2A2,ZMIZ1,DMTF1,CITED1,ZSCAN12,LDB1,GATB,ARID1B

C7,NEDD4L,SND1,ZNF398,NR1D2,ZNF823,ATP2A2,ZMIZ1,DMTF1,CITED1,ZSCAN12,LDB1,GATB,ARID1B

F20,GABBR1,ZFPM2,PFAS,THOC7,NEDD4L,SND1,ZNF398,NR1D2,ZNF823,EXT1,ATP2A2,ZMIZ1,DMTF1,

,EXT1,ATP2A2,ZMIZ1,DMTF1,CITED1,ZSCAN12,LDB1,GATB,ARID1B,ZNF638,FGFR1,CNOT1,GUCY1A2,P

JSP4,FHIT,SGF29,RBM6,TNRC6A,CUL3,RPTOR,HDAC4,PPM1E,L3MBTL4,TCF20,ZFPM2,COL11A1,THOC

,ZMIZ1,DMTF1,CITED1,ZSCAN12,LDB1,GATB,ARID1B,ZNF638,FGFR1,CNOT1,GUCY1A2,PRKAG2,XRCC

PTOR,HDAC4,PPM1E,L3MBTL4,TCF20,ZFPM2,COL11A1,THOC7,NEDD4L,SND1,ZNF398,NR1D2,FXR1,ZN

,ZFPM2,PFAS,THOC7,NEDD4L,SND1,ZNF398,NR1D2,ZNF823,ATP2A2,ZMIZ1,DMTF1,CITED1,ZSCAN12,L

PM1E,GABBR1,NEDD4L,CBLB,NR1D2,EXT1,ITGA11,ATP2A2,ARFGEF2,CITED1,CHRNA3,CACNA1C,LDB1

,ZFPM2,PFAS,THOC7,NEDD4L,SND1,ZNF398,NR1D2,ZNF823,ATP2A2,ZMIZ1,DMTF1,CITED1,ZSCAN12,L

JD1C,RFC2,HLA-DRB5,KEL,PDE4B,SHISA9,SEMA3F,HLA-DRB1,CCDC88C,NTRK3,SYNE1,WNT3,THRB,C

L2,ZMIZ2,TENM2,NFIX,ZNF615,THRB,BRWD1,KDM3B,TRIM33,BCL11B,FOXO6,CSRNP3,GLI3,RHOA,AFF

L2,ZMIZ2,TENM2,NFIX,ZNF615,THRB,BRWD1,KDM3B,TRIM33,BCL11B,FOXO6,CSRNP3,GLI3,RHOA,AFF

L2,ZMIZ2,TENM2,NFIX,ZNF615,THRB,BRWD1,KDM3B,TRIM33,BCL11B,FOXO6,CSRNP3,GLI3,RHOA,AFF

L2,POLR2E,ZMIZ2,TENM2,NFIX,SRPK2,ZNF615,THRB,BRWD1,KDM3B,TRIM33,BCL11B,FOXO6,CSRNP3

PRKG1,ESRRB,MAML3,BCL11A,TSHZ3,FOXO3,LCORL,KDM4A,JMJD1C,ZNF600,RNF111,RBL2,POLR2E,[illegible]

A-DRB5,SEMA3F,HLA-DRB1,RGS6,PPP1R13B,TENM2,NFIX,SRPK2,NTRK3,WNT3,THRB,BTN2A2,PPP2R[illegible]

[illegible]CL11A,TSHZ3,FOXO3,LCORL,KDM4A,JMJD1C,ZNF600,RNF111,RBL2,ZMIZ2,TENM2,CCDC88C,NFIX,SR ıM2,NTRK3,HLA-DQB1,WNT3,DOCK4,TRIM33,SORBS1,BCL11B,FOXO6,GLI3,RHOA,PIK3R2,AFF3,HDAC9

4,CADM2,PALB2,CTNND1,SOX5,CTNNB1,NGEF,NTM,CACNB2,ANK3,PRKG1,CACNA1I,SRR,CTTNBP2,TW

CL11A,TSHZ3,FOXO3,PTPRO,KDM4A,RFC2,HLA-DRB5,SEMA3F,HLA-DRB1,RGS6,PPP1R13B,TENM2,NF

,POLR2E,ZMIZ2,PPP2R2B,HLA-DRB1,RGS6,PPP1R13B,TENM2,NFIX,SRPK2,NTRK3,HLA-DQB1,WNT3,TH

3,FOXO3,LCORL,KDM4A,JMJD1C,ZNF600,RNF111,RBL2,POLR2E,ZMIZ2,TENM2,NFIX,GATAD2B,ZNF615,

3,FOXO3,LCORL,KDM4A,JMJD1C,ZNF600,RNF111,RBL2,POLR2E,ZMIZ2,TENM2,NFIX,GATAD2B,ZNF615,

1,ZSWIM6,DBN1,MAGI2,RASGRP4,CADM2,PALB2,CTNND1,SOX5,CTNNB1,NGEF,NTM,CACNB2,ANK3,PF

K1,RHOJ,ZSWIM6,DBN1,MAGI2,RASGRP4,CADM2,PALB2,CTNND1,SOX5,CTNNB1,NGEF,NTM,CACNB2,A

3,FOXO3,LCORL,KDM4A,JMJD1C,ZNF600,RNF111,RBL2,POLR2E,ZMIZ2,TENM2,NFIX,GATAD2B,ZNF615,

5,MEF2C,ZNF568,ZNF19,GMIP,TFB1M,ZKSCAN3,MAGI2,CTNND1,SOX5,CTNNB1,NGEF,PCGF3,FURIN,M

N,TWF2,ESRRB,MICA,MAML3,BCL11A,TSHZ3,FOXO3,HLA-DRB5,RNF111,POLR2E,ZMIZ2,PPP2R2B,HLA

OX5,CTNNB1,PCGF3,FURIN,MED27,ESRRB,DPYD,MAML3,BCL11A,TSHZ3,FOXO3,LCORL,KDM4A,JMJD

'OM121C,SLC35F2,RFT1,GBF1,HLA-DRB5,KCNN2,BTN3A2,PPP2R2B,HLA-DRB1,ELMO1,IMMP2L,CLCN3,

C35F2,RFT1,GBF1,HLA-DRB5,KCNN2,BTN3A2,PPP2R2B,HLA-DRB1,ELMO1,IMMP2L,CLCN3,SLC39A8,AT

GRIK1,RHOJ,MAGI2,RASGRP4,CTNND1,IMPA2,CTNNB1,NGEF,ANK3,PTPRF,FURIN,PRKG1,CACNA1I,PP

UCY1A2,PRKAG2,XRCC6,SLC30A9,ETF1,CREB5,MEF2C,ZNF568,ZNF19,GMIP,TFB1M,ZKSCAN3,MAGI2,

D1,SOX5,CTNNB1,PCGF3,MED27,ESRRB,CPEB1,MAML3,BCL11A,TSHZ3,FOXO3,LCORL,KDM4A,JMJD1

DM4A,KEL,RNF111,LONRF2,POLR2E,CCDC88C,MANBA,SRPK2,NTRK3,GALNT1,TNXB,GALNT10,DPP8,I

B,TFB1M,ZKSCAN3,CTNND1,SOX5,CTNNB1,ST3GAL3,PCGF3,MED27,ESRRB,CPEB1,MAML3,BCL11A,T

B,TFB1M,ZKSCAN3,CTNND1,SOX5,CTNNB1,ST3GAL3,PCGF3,MED27,ESRRB,CPEB1,MAML3,BCL11A,T

,NT5C2,ZNF638,PRPF3,FGFR1,MSH5,CNOT1,GUCY1A2,PRKAG2,XRCC6,SLC30A9,ETF1,CREB5,ZSCAN

,NT5C2,ZNF638,PRPF3,FGFR1,MSH5,CNOT1,GUCY1A2,PRKAG2,XRCC6,SLC30A9,ETF1,CREB5,ZSCAN

CITED1,CKB,ZSCAN12,LDB1,GATB,ARID1B,NT5C2,ZNF638,PRPF3,FGFR1,MSH5,CNOT1,GUCY1A2,PRK

PRKAG2,XRCC6,SLC30A9,ETF1,CREB5,ZSCAN31,MEF2C,ZNF568,ZNF19,MGAT5B,TFB1M,ZKSCAN3,CTN

7,NEDD4L,CBLB,SND1,ZNF398,NR1D2,FXR1,MMP16,ZNF823,EXT1,ZMIZ1,DMTF1,ARFGEF2,CITED1,CHI

6,SLC30A9,ETF1,CREB5,ZSCAN31,MEF2C,ZNF568,ZNF19,MGAT5B,TFB1M,ZKSCAN3,CTNND1,SOX5,IMI

F823,EXT1,ZMIZ1,DMTF1,ARFGEF2,CITED1,CHRNA3,TTLL7,ZSCAN12,LDB1,GATB,ARID1B,ZNF638,PRF

.DB1,GATB,ARID1B,NT5C2,ZNF638,PRPF3,FGFR1,MSH5,CNOT1,GUCY1A2,PRKAG2,XRCC6,SLC30A9,E

,FGFR1,MSH5,HLA-DQA1,GUCY1A2,PRKAG2,XRCC6,SLC30A9,MEF2C,ATP23,GMIP,GRIK1,RHOJ,MAGI

.DB1,GATB,ARID1B,NT5C2,ZNF638,PRPF3,FGFR1,MSH5,CNOT1,GUCY1A2,PRKAG2,XRCC6,SLC30A9,E

ZMIZ2,RGS6,TENM2,NFIX,SRPK2,NTRK3,ZNF615,THRB,PREX1,BRWD1,DOCK4,KDM3B,TRIM33,BCL11B

PK2,NTRK3,ZNF615,WNT3,THRB,BRWD1,KDM3B,TRIM33,BCL11B,FOXO6,CSRNP3,GLI3,RHOA,AFF3,H[illegible]

WF2,BCL11A,TSHZ3,FOXO3,PTPRO,SEMA3F,PPP2R2B,RGS6,TENM2,SRPK2,NTRK3,IMMP2L,WNT3,THR

IX,SRPK2,NTRK3,WNT3,THRB,BTN2A2,PPP2R5C,TRIM33,BCL11B,FOXO6,AMBRA1,GLI3,RHOA,TRIOBP,

HRB,PPP2R5C,TENM4,TRAIP,SORBS1,BCL11B,FOXO6,AMBRA1,CSRNP3,GLI3,RHOA,PIK3R2,TRIOBP,HI

,THRB,BRWD1,KDM3B,TRIM33,BCL11B,FOXO6,CSRNP3,MED8,GLI3,ZKSCAN8,RHOA,AFF3,HDAC9,ZNF

,THRB,BRWD1,KDM3B,TRIM33,BCL11B,FOXO6,CSRNP3,MED8,GLI3,ZKSCAN8,RHOA,AFF3,HDAC9,ZNF

RKG1,CACNA1I,SRR,CTTNBP2,TWF2,ESRRB,BCL11A,TSHZ3,FOXO3,PTPRO,SEMA3F,RNF111,PPP2R2E

ANK3,MED27,PRKG1,CACNA1I,SRR,CTTNBP2,TWF2,ESRRB,BCL11A,TSHZ3,FOXO3,PTPRO,SEMA3F,PI

,THRB,BRWD1,KDM3B,TRIM33,BCL11B,FOXO6,CSRNP3,MED8,GLI3,ZKSCAN8,RHOA,AFF3,HDAC9,ZNF

ED27,PRKG1,ESRRB,CPEB1,MAML3,BCL11A,TSHZ3,FOXO3,LCORL,KDM4A,JMJD1C,ZNF600,RNF111,RI

-DRB1,RGS6,PPP1R13B,TENM2,NFIX,SRPK2,NTRK3,HLA-DQB1,WNT3,THRB,ATP2B2,PPP2R5C,TENM4

1C,ZNF600,RNF111,RBL2,POLR2E,ZMIZ2,TENM2,NFIX,GATAD2B,ZNF615,THRB,ATP2B2,BRWD1,KDM3E

P1R16B,ESRRB,PLCL2,TANK,MICA,MAML3,FOXO3,PTPRO,GBF1,HLA-DRB5,PDE4B,NKIRAS1,RNF111,S

RASGRP4,CTNND1,SOX5,CD55,CTNNB1,NGEF,PCGF3,ANK3,FURIN,MED27,PRKG1,ESRRB,CPEB1,MAM

C,ZNF600,RNF111,RBL2,POLR2E,ZMIZ2,TENM2,NFIX,SRPK2,GATAD2B,ZNF615,NOVA1,THRB,PPA2,BRW

MMP2L,PPA2,PPP2R5C,TRIM33,MED8,GLI3,NEURL1,RHOA,TRIOBP,NAALADL2,HDAC9,RNF144A,TLK1

SHZ3,FOXO3,RFT1,LCORL,KDM4A,JMJD1C,RFC2,ZNF600,RNF111,RBL2,POLR2E,ZMIZ2,TENM2,NFIX,GA

SHZ3,FOXO3,RFT1,LCORL,KDM4A,JMJD1C,RFC2,ZNF600,RNF111,RBL2,POLR2E,ZMIZ2,TENM2,NFIX,GA

31,MEF2C,ATP23,ZNF568,ZNF19,GMIP,RHOJ,TFB1M,ZKSCAN3,PALB2,CTNND1,SOX5,CTNNB1,NGEF,PC

31,MEF2C,ATP23,ZNF568,ZNF19,GMIP,RHOJ,TFB1M,ZKSCAN3,PALB2,CTNND1,SOX5,CTNNB1,NGEF,PC

AG2,XRCC6,SLC30A9,ETF1,CARMIL1,CREB5,ZSCAN31,MEF2C,ATP23,ZNF568,ZNF19,GMIP,RHOJ,TFB1M

IND1,SOX5,IMPA2,CTNNB1,ST3GAL3,PCGF3,FURIN,MED27,SRR,ESRRB,DPYD,CPEB1,MAML3,BCL11A,T

RNA3,TTLL7,ZSCAN12,LDB1,GATB,ARID1B,ZNF638,PRPF3,FGFR1,MSH5,CNOT1,PRKAG2,XRCC6,SLC3

PA2,CTNNB1,ST3GAL3,PCGF3,FURIN,MED27,SRR,ESRRB,DPYD,CPEB1,MAML3,BCL11A,TSHZ3,PSPH,F

F3,FGFR1,MSH5,CNOT1,PRKAG2,XRCC6,SLC30A9,ETF1,CREB5,ZSCAN31,MEF2C,ATP23,ZNF568,ZNF1

TF1,CARMIL1,CREB5,ZSCAN31,MEF2C,ATP23,ZNF568,ZNF19,GMIP,RHOJ,TFB1M,ZKSCAN3,PALB2,CTN

2,RASGRP4,PALB2,CTNND1,SOX5,IMPA2,CTNNB1,NGEF,ANK3,PTPRF,FURIN,PRKG1,CACNA1I,TWF2,P

TF1,CARMIL1,CREB5,ZSCAN31,MEF2C,ATP23,ZNF568,ZNF19,GMIP,RHOJ,TFB1M,ZKSCAN3,PALB2,CTN

,RGS6,TENM2,SRPK2,NTRK3,IMMP2L,WNT3,THRB,MACROD2,PREX1,ATP2B2,TENM4,BCL11B,FOXO6,

PP2R2B,RGS6,TENM2,SRPK2,NTRK3,SYNE1,IMMP2L,WNT3,THRB,MACROD2,PREX1,ATP2B2,BRWD1,T

BL2,POLR2E,ZFYVE1,ZMIZ2,PPP2R2B,RGS6,TENM2,CCDC88C,NFIX,SRPK2,NTRK3,TNXB,ZNF615,THRE

,TRAIP,SORBS1,BCL11B,FOXO6,AMBRA1,CSRNP3,GLI3,RHOA,PIK3R2,BTN3A1,TRIOBP,HDAC9

ORCS3,PPP2R2B,HLA-DRB1,RGS6,TENM2,CCDC88C,SRPK2,NTRK3,TNXB,HLA-DQB1,ELMO1,WNT3,PF

/L3,BCL11A,TSHZ3,FOXO3,LCORL,KDM4A,JMJD1C,ZNF600,RNF111,RBL2,POLR2E,ZFYVE1,ZMIZ2,PPP2

VD1,KDM3B,TRIM33,BCL11B,FOXO6,CSRNP3,MED8,GLI3,ZKSCAN8,RHOA,AFF3,PUS7,HDAC9,ZNF800,(

ALNT1,GALNT10,GATAD2B,ZNF615,THRB,PPA2,BRWD1,KDM3B,TRIM33,SORBS1,BCL11B,FOXO6,CSRN

ALNT1,GALNT10,GATAD2B,ZNF615,THRB,PPA2,BRWD1,KDM3B,TRIM33,SORBS1,BCL11B,FOXO6,CSRN

CGF3,MED27,PRKG1,ESRRB,DPYD,CPEB1,MAML3,BCL11A,TSHZ3,FOXO3,LCORL,KDM4A,JMJD1C,RFC

CGF3,MED27,PRKG1,ESRRB,DPYD,CPEB1,MAML3,BCL11A,TSHZ3,FOXO3,LCORL,KDM4A,JMJD1C,RFC

/I,ZKSCAN3,PALB2,CTNND1,SOX5,CTNNB1,ST3GAL3,NGEF,PCGF3,FURIN,MED27,PRKG1,SRR,ESRRB,

TSHZ3,PSPH,FOXO3,RFT1,LCORL,KDM4A,JMJD1C,RFC2,ZNF600,PIP5K1B,RNF111,RBL2,POLR2E,ZMIZ

0A9,ETF1,CREB5,ZSCAN31,MEF2C,ATP23,ZNF568,ZNF19,MGAT5B,TFB1M,ZKSCAN3,MAGI2,PALB2,CTN

OXO3,RFT1,LCORL,KDM4A,JMJD1C,RFC2,ZNF600,PIP5K1B,RNF111,RBL2,POLR2E,ZMIZ2,TENM2,NFIX

I9,MGAT5B,TFB1M,ZKSCAN3,MAGI2,PALB2,CTNND1,SOX5,CTNNB1,NEK4,ST3GAL3,PCGF3,PTPRF,FUI

IND1,SOX5,CTNNB1,NGEF,PCGF3,MED27,PRKG1,ESRRB,DPYD,CPEB1,MAML3,BCL11A,TSHZ3,FOXO3,

PP1R16B,ESRRB,PLCL2,TANK,MICA,MAML3,FOXO3,PTPRO,RFC2,GBF1,HLA-DRB5,PDE4B,NKIRAS1,RI

IND1,SOX5,CTNNB1,NGEF,PCGF3,MED27,PRKG1,ESRRB,DPYD,CPEB1,MAML3,BCL11A,TSHZ3,FOXO3,

[illegible],PREX1,BTN2A2,BRWD1,DOCK4,KDM3B,TRIM33,SORBS1,BCL11B,FOXO6,CSRNP3,GLI3,RHOA,AFF3,H

REX1,PPP2R5C,TENM4,DOCK4,TRAIP,TRIM33,SORBS1,FOXO6,GLI3,NEURL1,RHOA,PIK3R2,BTN3A1,HD

2R2B,RGS6,TENM2,CCDC88C,NFIX,SRPK2,NTRK3,TNXB,ZNF615,WNT3,THRB,PREX1,BTN2A2,BRWD1,

2,ZNF600,PDE4B,RNF111,RBL2,POLR2E,ZMIZ2,RGS6,TENM2,NFIX,SRPK2,NTRK3,GATAD2B,ZNF615,N

2,ZNF600,PDE4B,RNF111,RBL2,POLR2E,ZMIZ2,RGS6,TENM2,NFIX,SRPK2,NTRK3,GATAD2B,ZNF615,N

DPYD,CPEB1,IVD,MAML3,BCL11A,TSHZ3,PSPH,FOXO3,LCORL,KDM4A,JMJD1C,RFC2,ZNF600,PDE4B,R

2,TENM2,NFIX,GALNT1,GALNT10,GATAD2B,ZNF615,THRB,PPA2,ATP2B2,BRWD1,KDM3B,TRIM33,SORE

ND1,SOX5,CD55,CTNNB1,NEK4,ST3GAL3,PCGF3,ANK3,PTPRF,FURIN,MED27,PRKG1,ESRRB,CPEB1,M

,GALNT1,GALNT10,GATAD2B,ZNF615,THRB,PPA2,ATP2B2,BRWD1,KDM3B,TRIM33,SORBS1,BCL11B,F

RIN,MED27,PRKG1,ESRRB,CPEB1,MAML3,BCL11A,TSHZ3,FOXO3,RFT1,LCORL,PTPRO,KDM4A,JMJD1C

LCORL,KDM4A,JMJD1C,RFC2,ZNF600,PDE4B,RNF111,RBL2,POLR2E,ZMIZ2,RGS6,TENM2,NFIX,SRPK2,

NF111,POLR2E,SORCS3,PPP2R2B,HLA-DRB1,RGS6,TENM2,CCDC88C,SRPK2,NTRK3,TNXB,HLA-DQB1,

LCORL,KDM4A,JMJD1C,RFC2,ZNF600,PDE4B,RNF111,RBL2,POLR2E,ZMIZ2,RGS6,TENM2,NFIX,SRPK2,

OVA1,THRB,MACROD2,PPA2,PREX1,ATP2B2,BRWD1,DOCK4,KDM3B,TRIM33,BCL11B,FOXO6,CSRNP3,N

OVA1,THRB,MACROD2,PPA2,PREX1,ATP2B2,BRWD1,DOCK4,KDM3B,TRIM33,BCL11B,FOXO6,CSRNP3,N

NF111,RBL2,POLR2E,ZMIZ2,RGS6,TENM2,NFIX,SRPK2,NTRK3,GATAD2B,ZNF615,NOVA1,THRB,MACRO

IAML3,BCL11A,TSHZ3,FOXO3,RFT1,LCORL,PTPRO,KDM4A,JMJD1C,RFC2,ZNF600,KEL,RNF111,RBL2,L

,RFC2,ZNF600,RNF111,RBL2,POLR2E,ZMIZ2,TENM2,CCDC88C,MANBA,NFIX,SRPK2,NTRK3,GALNT1,TN

NTRK3,GATAD2B,ZNF615,NOVA1,THRB,MACROD2,PPA2,PREX1,ATP2B2,BRWD1,DOCK4,KDM3B,TRIM3

ELMO1,WNT3,PREX1,PPP2R5C,TENM4,DOCK4,TRAIP,TRIM33,SORBS1,FOXO6,GLI3,NEURL1,RHOA,PIK

NTRK3,GATAD2B,ZNF615,NOVA1,THRB,MACROD2,PPA2,PREX1,ATP2B2,BRWD1,DOCK4,KDM3B,TRIM3

[illegible]D2,PPA2,PREX1,ATP2B2,BRWD1,DOCK4,KDM3B,TRIM33,BCL11B,FOXO6,CSRNP3,MED8,GLI3,ZKSCA[illegible]

[illegible]NRF2,POLR2E,ZMIZ2,TENM2,CCDC88C,MANBA,NFIX,SRPK2,NTRK3,GALNT1,TNXB,GALNT10,GATAD2

IXB,GALNT10,GATAD2B,ZNF615,NOVA1,IMMP2L,THRB,PPA2,PPP2R5C,BRWD1,KDM3B,TRIM33,SORBS

33,BCL11B,FOXO6,CSRNP3,MED8,GLI3,ZKSCAN8,RHOA,AFF3,PUS7,HDAC9,ZNF800,CDKAL1,SREK1IP1

33,BCL11B,FOXO6,CSRNP3,MED8,GLI3,ZKSCAN8,RHOA,AFF3,PUS7,HDAC9,ZNF800,CDKAL1,SREK1IP1

:B,ZNF615,DPP8,NOVA1,IMMP2L,WNT3,THRB,PPA2,PPP2R5C,BRWD1,KDM3B,TRIM33,SORBS1,BCL11E

1,BCL11B,FOXO6,CSRNP3,MED8,GLI3,ZKSCAN8,NEURL1,RHOA,AFF3,PUS7,TRIOBP,HDAC9,RNF144A,[illegible]

[illegible],FOXO6,CSRNP3,MED8,GLI3,ZKSCAN8,NEURL1,RHOA,AFF3,PUS7,TRIOBP,NAALADL2,HDAC9,RNF144[illegible]

A,ZNF
